# Principles underlying implementation of *nearly*-homeostatic biological networks

**DOI:** 10.1101/2025.04.02.646789

**Authors:** Zhe F. Tang, David R. McMillen

## Abstract

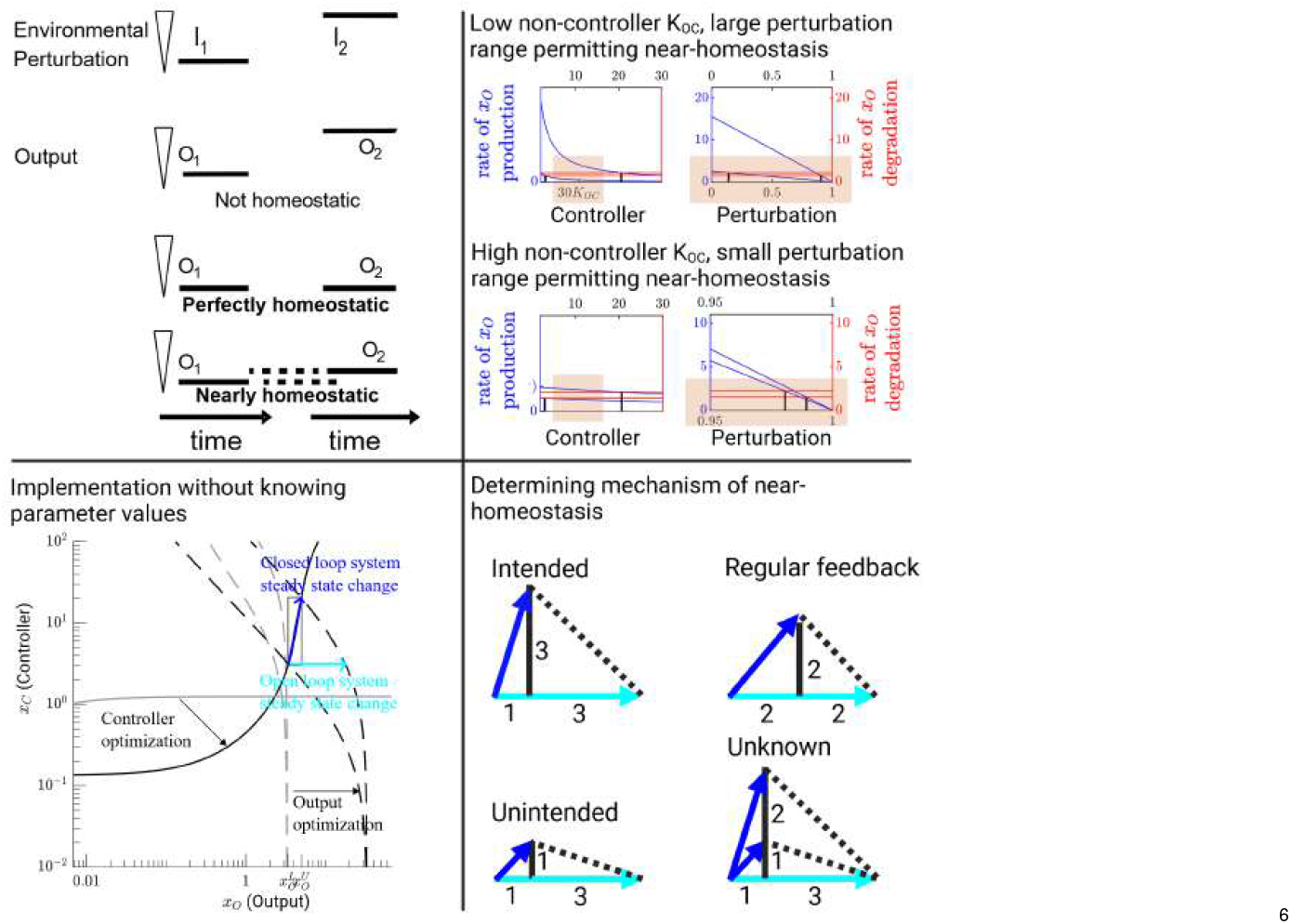

A nearly-homeostatic biological system keeps the steady-state output (like internal body temperature) within a narrow range, regardless of different persistent levels of environmental perturbations (like external temperatures). A nearly-homeostatic system can guarantee performance of a therapeutic device in different patient contexts or can be a detector that generates a response only upon encountering an anomaly. We have developed the inverse homeostasis perspective to determine the impact of each parameter on the system’s homeostatic performance, which has allowed us to vastly widen the region of implementable parameter sets supporting near-homeostasis compared to the predominant approach of minimizing the controller’s integration leakiness. To implement nearly-homeostatic systems without measuring parameter values, we have used feedback-free auxiliary systems to accurately approximate steady-state response curves of key components of the feedback system and adjusted those curves to attain desired characteristics. Our approach has not only discovered a new mechanism of near-homeostasis, but also serves as a new framework to reinterpret whether the homeostatic performance observed in previously published systems arises from the mechanisms originally proposed to explain the behaviour.

## INTRODUCTION

A homeostatic system like internal body temperature maintenance is able keep the steady state system output (like internal body temperature) at a fixed set point, regardless of different persistent levels of environmental perturbations (like external temperatures)^1,2^ (Figure 1A). A nearly homeostatic system maintain the steady state output within some narrow range, regardless of different persistent levels of environmental perturbation (Figure 1A).

**Figure 1:**
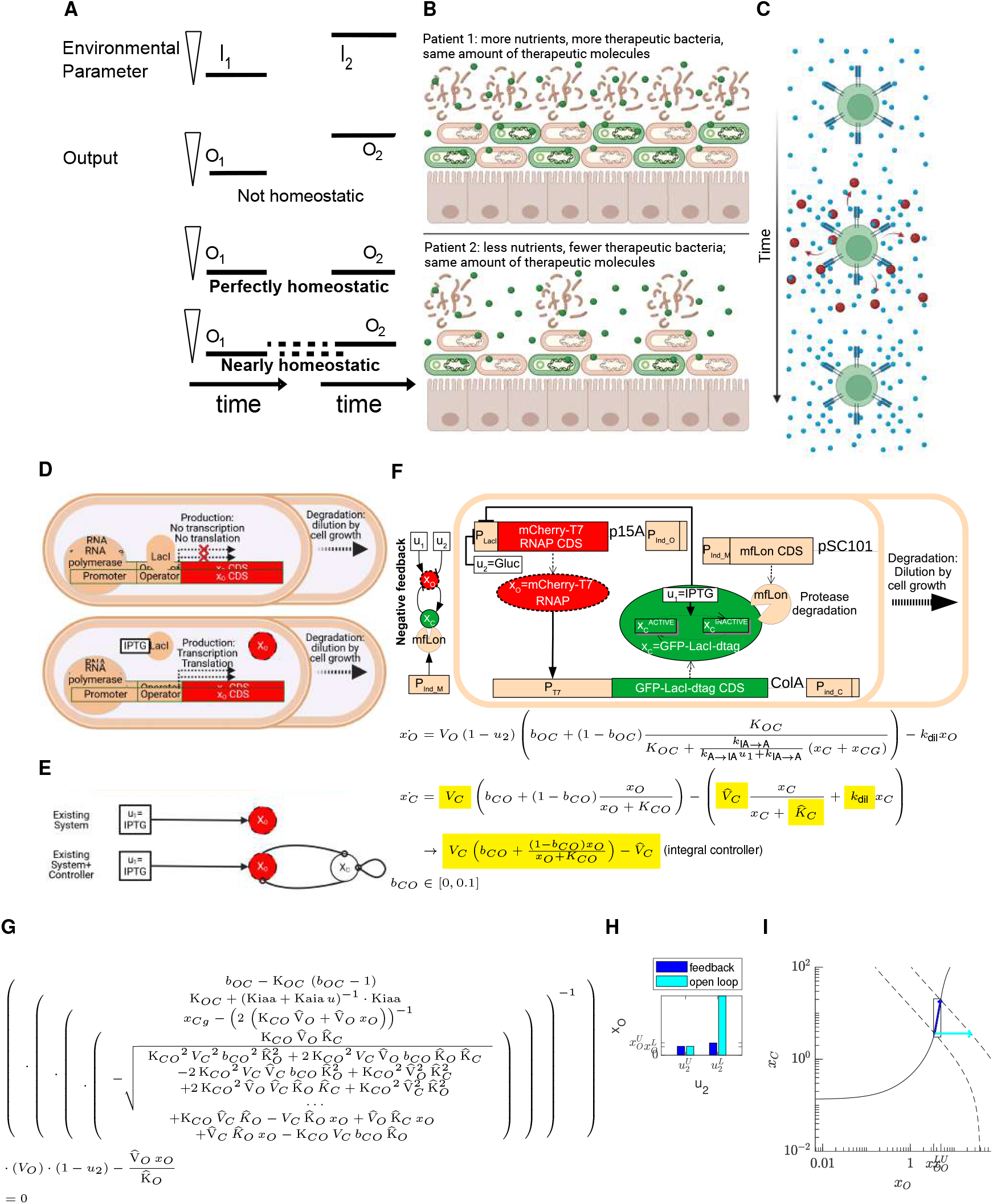
An introduction to homeostatic systems. (A) Definition of homeostasis and near-homeostasis. (B) A homeostatic system can ensure therapeutic molecular level is independent of nutrient state or microbiotic competitive landscape. (C) A homeostatic system can limit aberrant immune response by limiting response of T cell (red cytotoxic molecules) to transient changes in stimuli (blue molecules) rather than persistent stimuli ^3,4^. It would be useful to engineer CAR-T cells to have such a property as well. (D) A basic transcriptional element to be modeled. (E) Definition of implementing a system to ensure near-homeostasis of the output *x*_*O*_’s steady state. (F) Reducing leaky integral control to result in perfect integral control. The output *x*_*O*_ (mCherry-tagged T7 RNA polymerase) up-regulates expression of the controller *x*_*C*_ (EGFP-tagged LacI), which in turn down-regulates the output. IPTG *u*_1_ ≥ 0 and the glucose-induced fraction of repressed promoters *u*_2_ ∈ [0, 1] are the two perturbations. All regulatory parameters except the basal expression parameters *b*_*OC*_, *b*_*CO*_ ∈ [0, 0.1] are limited to the range of [0, 10]. The controller exists in an EGFP-labelled version supplied on a plasmid, *x*_*C*_, but a non labeled version of LacI *x*_*CG*_ also exists in the genome. *V*_*O*_ and *V*_*C*_ are maximal production rates of *x*_*O*_ and *x*_*C*_, respectively. *b*_*OC*_ and *b*_*CO*_ are fractions of maximal production rate of *x*_*O*_ and *x*_*C*_ induced by their corresponding regulators at base line, respectively. *K*_*OC*_ and *K*_*CO*_ are interaction affinity values for the *x*_*O*_’s promoter to *x*_*C*_ pair and the *x*_*C*_’s promoter to *x*_*O*_ pair, respectively. 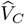 is the maximal mfLon-based protease degradation rate for *x*_*C*_, and 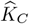 is the protease’s affinity value towards its mfLon-tagged substrate *x*_*C*_. (G) Implicit expression of the perturbation to steady state map for the feedback system in Figure 1F. (H) Conventional comparison between feedback system and open loop system using matched output only. (I) Steady state response curve analysis adds another avenue to compare performance of the open and closed loop system. The solid curve represents the steady state of the controller in response to different steady state output levels. Each dashed curve represents the steady state of the output in response to different steady state controller levels, at a particular perturbation. The blue and cyan arrows respectively represent the changes in steady state for the feedback system and the open loop system between two fixed perturbations.

It would be useful to construct synthetic nearly-homeostatic systems to guarantee performance of a therapeutic device in different patient contexts or to generate a response only upon detection of an anomaly. For example, we can engineer bacteria to secret therapeutic molecules (shown in green) in a manner that responds primarily to a sensed disease state such as gut inflammation, but is not affected by other perturbations such as different nutrient states and different microbiotic competitive landscapes between patients as well as over time within an individual patient (Figure 1B). We can also build a homeostatic system whose output increases from its basal level only when the perturbation is changing rather than staying the same, effectively acting like a statistical anomaly detector (Figure 1C).

First order ordinary differential equations are useful descriptors of nearly-homeostatic biological systems. We will introduce our approach with a running mathematical example that will also be implemented experimentally. Suppose our existing system *A*_*E*_ is described by a set of first order Ordinary Differential Equations (ODEs):

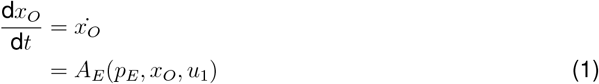

In this example (Figure 1D), perturbation *u*_1_ = *IPTG* activates transcription of the system output *x*_*O*_, by deactivating the LacI repressor *x*_*C*_ + *x*_*CG*_. Here, *x*_*C*_ and *x*_*CG*_ denotes plasmid encoded and genomically encoded LacI, respectively. This first order ODE says that the current rate of change in the system output *x*_*O*_ depends on a fixed set of non-negative regulatory parameters *p*_*E*_, the current level of the system output itself, and some fixed IPTG perturbation level *u*_1_. The rate of change of the system output is its production rate subtracted by its degradation rate

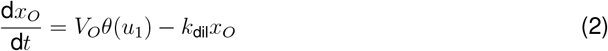

The production rate of the system output is simply the maximal production rate *V*_*O*_ multiplied by fraction of available promoters dependent on the perturbation *u*_1_,

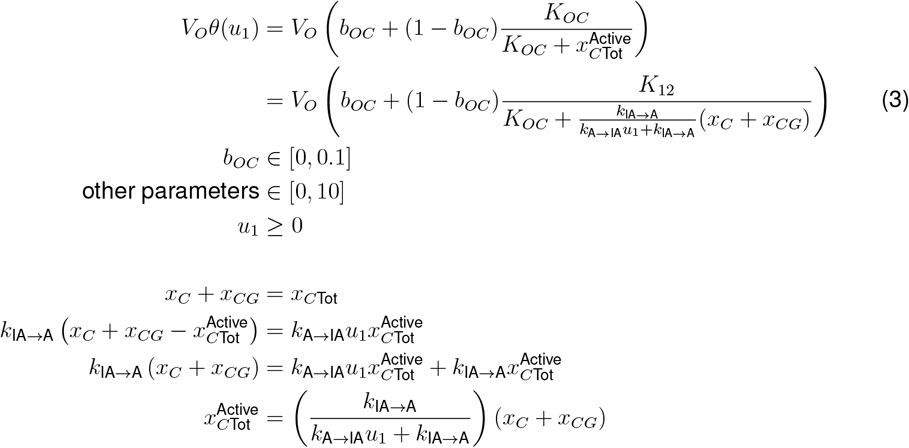

The degradation rate of the system output is dilution of the system output due to cell growth, which assumes that the proteins are not being rapidly degraded by other processes such as proteolysis. The maximal production rate *V*_*O*_, cell dilution constant *k*_dil_, and any constants within the function *θ*(*u*_1_) are all regulatory parameter coordinates of parameter set *p*_*E*_. At steady state, the rates of all system components equal zero

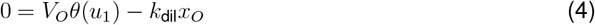

Thus, we can find the steady state of the system output at each perturbation level 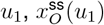, by algebraically isolating *x*_*O*_

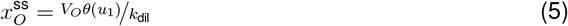

Empirically, we define the near-homeostasis of the system output (also known as the homeostatic performance of the system) as the difference in system output steady state at two fixed perturbations (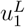 and 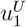), further normalized by the scale of the perturbation change 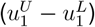 and the system output level 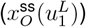:

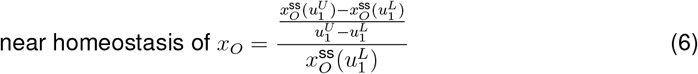

To make the output *x*_*O*_ of the existing system nearly-homeostatic, we want to find a controller architecture composed of how the controller works internally *A*_*C*_ and how the controller acts upon the existing system *A*_*CO*_, and implement a set of controller parameters *p*_*C*_

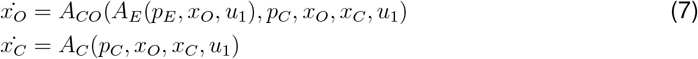

such that whenever a persistent perturbation IPTG is within the desired range 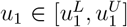, the steady state output is also within the desired range 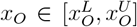. For example, the controller architecture may be a single component (*x*_*C*_ in Figure 1E) that provides feedback for the system output based on the system output value.

Since it is difficult to measure parameter values experimentally due to identifiability, multiple local minima, and in-vitro to in-vivo transferability limitations^5,6^ (supplementary information SI1), multiple approaches were used to implement different nearly-homeostatic systems with severe inherent limitations for each approach. A conceptually straightforward approach is to explicitly determine the effect of parameters on the mapping from perturbations to steady state outputs and then change those parameters in desired directions until near-homeostasis is attained^7^, but that approach cannot be readily applied to complicated perturbation to output steady state maps. Even our simple negative feedback system (Figure 1F) has a complicated perturbation to output steady state map (Figure 1G). The predominant approach to implement nearly-homeostatic systems without measuring parameter values is getting the controller to behave as closely to a perfect-integral controller as possible^8–11^, by increasing 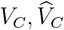 relative to *k*_dil_ and decreasing 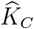 to zero (Figure 1F). However, this approach results in near-homeostasis if and only if the ideal integral controller combined with the non-controller component results in asymptotically stable steady states at all perturbations within desired range; and it is possible that the ideal combined feedback system does not permit asymptotically stable steady states^8^. Since a perfectly ideal integral controller cannot be implemented in practice, all we can do without more advanced tools is to keep minimizing integration leakiness and hope that the system output becomes nearly-homeostatic.

In addition, an open question in the process of implementing nearly-homeostatic systems experimentally is whether the observed near-homeostasis is supported by the intended mechanism. Implementations of homeostatic systems have demonstrated near-homeostasis of the output in the feedback system compared to open loop system at similar output values^9,10,12^ (Figure 1H), but the experiments do not necessarily demonstrate that the actual mechanism of near-homeostasis is as the same as the intended mechanism. In our example system, we can plot the steady state of the controller in response to constant output levels according to the controller’s rate equation (solid curve in Figure 1I). For the output, we can fix a perturbation value and plot the steady state of the output in response the constant controller levels according to the output’s rate equation (one dashed curve for each of 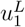 and 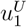 in Figure 1I). The intersection between the controller’s steady state response curve and the output’s steady state response curve at a particular perturbation is the steady state of the feedback system at that level of perturbation. Compared to the open loop system at two fixed perturbations (cyan arrow in Figure 1I), our near-integral controller minimizes the change in output by making the controller response curve increasingly vertical, resulting large changes in the controller steady states^1,13,14^ (blue arrow in Figure 1I). Since a large fold change in controller steady state is not necessarily the only way to attain the same change in output steady state at the fixed perturbations, controller steady states in addition to the output steady state should be measured. If the intended mechanism of near-homeostasis is different from the actual mechanism like the one we will eventually discover (Figure 5F), the network would exhibit unpredictable compensation behavior and have unpredictable interactions with the rest of the organism.

In this study, we devise and experimentally validate a theoretical framework to analyze and implement nearly homeostatic systems that does not measure parameter values, minimizes reliance on trial and error, and provides tests to verify the implemented mechanism of near-homeostasis. We have unexpectedly discovered a novel mechanism of near-homeostasis that already exists in nature, which could serve as a basis for synthetic programmable nearly-homeostatic systems.

## RESULTS AND DISCUSSION

### The inverse homeostasis perspective widens the region of implementable parameter sets supporting near-homeostasis by analyzing non-controller leakiness parameters

The integration leakiness reduction approach to implement nearly-homeostatic systems without measuring parameter values not only failed to consistently enable near-homeostasis, but also required very small leakiness to consistently attain near-homeostasis. Prior characterizations of the leaky integration reduction approach revealed that reducing integration leakiness eventually results in near-homeostasis if the feedback system containing the perfect integral controller permits a stable steady state^8^. In this study, we want to know often decreasing integration leakiness will eventually lead to near-homeostasis and at what integration leakiness near-homeostasis can be attained, if we stumble upon a feedback system having the same qualitative regulatory interactions as Figure 1F but has a random set of interaction strengths. We first randomly generated 100 sets of controller parameters in our example system (Figure 1F: parameters present in *x*_*C*_’s rate equation), such that for each controller parameter set and for each integration leakiness, there exists a randomly generated open loop system that results in at least 2-fold change in output steady state. For each set of controller parameters, a particular non-zero integration leakiness

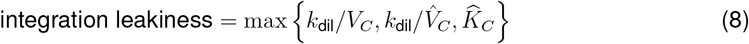

was then set to the desired values by multiplying non-dilution integration-leakiness parameters 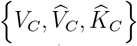 by a constant factor and zero integration leakiness was attained by setting *k*_dil_ = 0 and 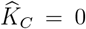. Each parameter set for the closed loop system was formed by combining the controller parameter set with its qualifying open loop parameter set. A system is considered nearly-homeostatic (or have satisfactory homeostatic performance) if the steady state output would only increase by less than 1/6 or 1/3 of the steady state output increase in log units in the the corresponding open loop system. For example, if the steady state output of the open loop system increases by 2 fold (log(2) log units), then the corresponding nearly-homeostatic closed loop system requires its steady state output to increase by less than 10^1/6 log(2)^ = 2^1/6^ fold (1/6 log(2) log units) or 10^1/3 log(2)^ = 2^1/3^ fold (1/3 log(2) log units). Consistent with prior observations^8^, reducing integration leakiness of randomly generated parameter sets only resulted in perfect homeostasis *∼* 20 *−* 60% of the time depending on which perturbation was changed (Figure 2A). In addition, even if one can guarantee that reducing integration leakiness from each non-zero value to a perfect integral controller permits stable steady states, different controller parameter sets with the same integration leakiness resulted in vastly different degrees of near-homeostasis depending on other regulatory parameter values (Figure 2B: different colors at each non-zero integration leakiness). Thus, if we only optimize integration leakiness of the controller to attain near-homeostasis, integration leakiness often needs to be quite small (Figure 2B: ≤ 0.001 integration leakiness) for a satisfactory percentage of randomly generated combined system to be nearly-homeostatic.

**Figure 2:**
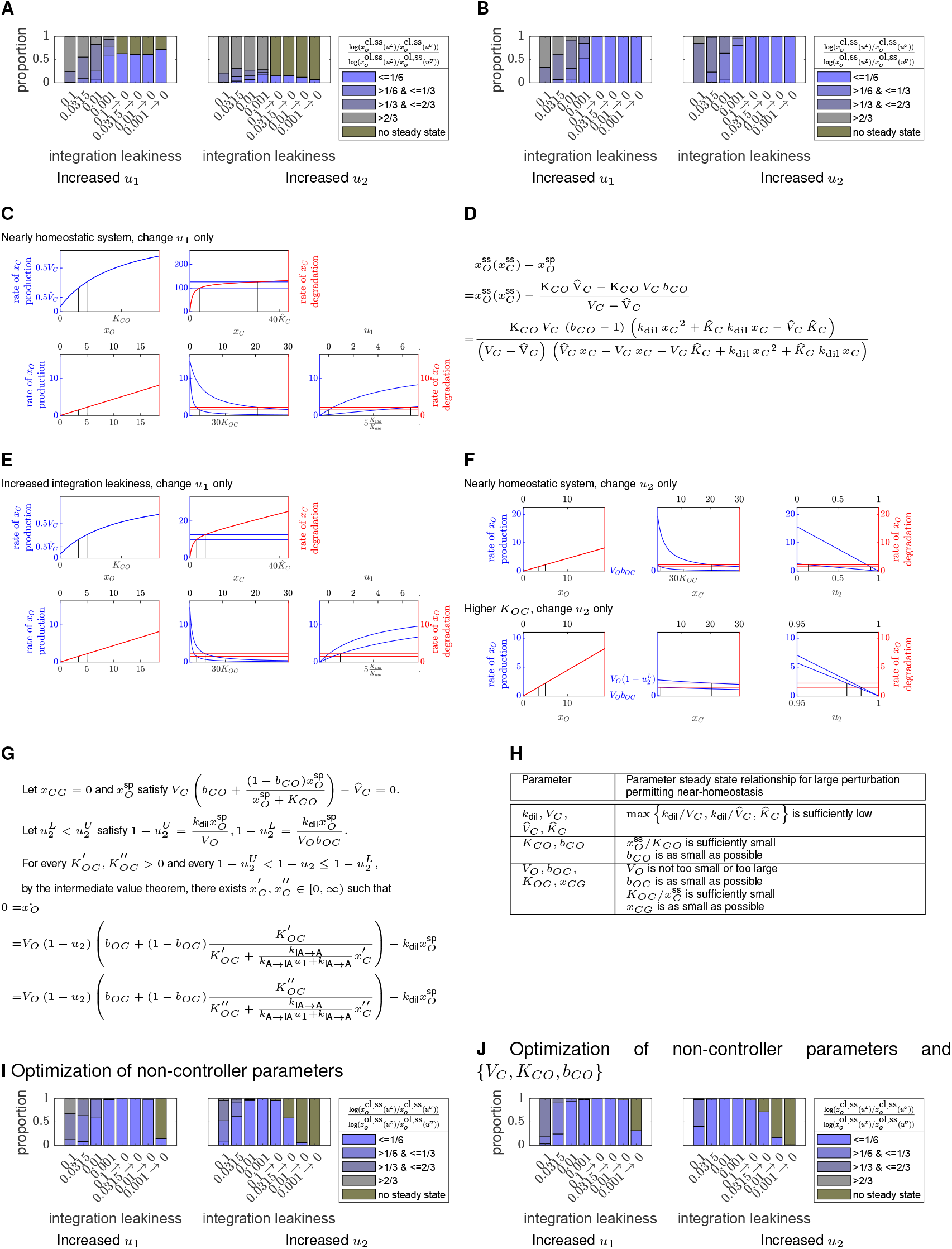
The inverse homeostasis perspective ensures near-homeostasis at far less stringent integral leakiness. (A,B) Degree of near-homeostasis at different integration leakiness without (A) and with (B) the assumption of the perfect integral controller permitting a stable steady state. (C,E,F) Inverse homeostasis plots associated with various parameter sets. *x*_*CG*_ = 0. (D) Algebraically computed distance between the controller’s response curve 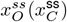 and the perfect integral controller’s response curve 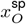 was minimized to improve near-homeostasis of the output^14^. It is not obvious that how increased 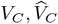 would change the algebraic distance even though we know the changes would improve near-homeostasis by decreasing integration leakiness. Decreasing *b*_*CO*_ increases the magnitude of algebraic distance but also still improves near-homeostasis, according to the inverse homeostasis plots. (G) Perturbation range permitting perfect homeostasis under the same conditions as Figure 2F. (H) Parameter steady-state relationships that enhance near-homeostasis. (I,J) Degree of near-homeostais at different integration leakiness when non-controller parameters (I) or non-controller-leakiness parameters (non-controller parameters ∪ {*V*_*C*_, *K*_*CO*_, *b*_*CO*_}; J) are optimized to maximally cover the desired perturbation range. In order for a perfect integral controller to attain a steady state, the maximal controller production rate *V*_*C*_ is always greater than the maximal protease degradation of the controller 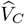, so optimization of *V*_*C*_, *K*_*CO*_, *b*_*CO*_ does not affect integration leakiness. Further, the inverse homeostasis perspective makes it unnecessary to verify whether the perfect integral controller paired with non-controller parameters admits a steady state. In fact, when non-controller parameters are purely optimized for the perturbation range permitting near-homeostasis to maximally cover the desired range, it is possible for the leaky integral controller but not the perfect integral controller to permit a stable steady state.

To find out what causes one set of parameters to result in better near-homeostasis than another parameter set despite having the same integration leakiness, we have developed the “inverse homeostasis perspective”. For illustration purposes, we will apply the inverse homeostasis perspective to perturbation *u*_1_ in our example system, initially keeping perturbation *u*_2_ fixed. The inverse homeostasis perspective maximizes the fold difference in perturbation *u*_1_ that will induce the desired small fold difference in the system output *x*_*O*_ at steady state. This perspective is the conceptual inverse of the more common approach in which one seeks to minimize the fold difference in the steady state system output *x*_*O*_ at two fixed perturbations 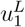 and 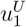.

The primary tool in applying the inverse homeostasis perspective is a set of inverse homeostasis plots, in order to understand how a small fold change in the output steady state from 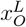 to 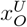 is induced by a large fold change in perturbation *u*_1_ from 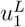 to 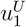. To generate inverse homeostasis plots for our example system that visualize the mechanism of near-homeostasis, we selectively plot production rates (always in blue) and degradation rates (always in red) of system components *x*_*O*_, *x*_*C*_ as functions of the system output *x*_*O*_, the controller state *x*_*C*_, the input (*u*_1_ or *u*_2_), and use as anchor points the steady states of a nearly-homeostatic feedback system at both ends of the perturbation range: 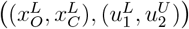 and 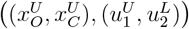. We first consider the production rate of the controller *x*_*C*_ as a function of the output *x*_*O*_ (Figure 2C: top left plot, in blue), and mark the desired range of steady state system outputs, 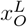 to 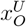, with black vertical lines. These output steady states defines specific steady-state production rates of the controller species, as indicated by where the vertical lines intersect with the blue controller production rate curve. We then create a new plot of the controller’s production and degradation rates as a function of its own state, *x*_*C*_ (Figure 2C: top middle plot): the controller production rates corresponding to the two output steady states 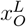 and 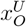 appear as horizontal blue lines, while the controller’s degradation rate appears as a red curve. The controller production rates as a function of the controller appear as horizontal lines because every rate curve must fix the non-varying component of its anchoring steady state (*x*_*O*_ in this case) and the controller *x*_*C*_’s production rate equation is independent of the varying component *x*_*C*_. Where the controller’s production and degradation rates intersect defines the controller’s steady state values 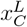 and 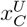, marked with black vertical lines. We can see that a small fold change in the output steady state from 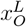 to 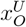 is induced by a small fold increase in steady state production and degradation rate of the controller *x*_*C*_, which is in turn induced by a large fold increase in controller steady state from 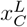 to 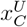.

We then proceed to generate plots that examine the production and degradation rates of output species *x*_*O*_, and see how the small fold increase in the output steady state from 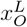 to 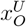 and the large fold increase in controller steady state from 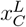 to 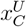 is induced by a large increase in perturbation from 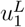 to 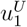. Plotting the degradation rate of the output species *x*_*O*_ against itself (Figure 2C: lower left plot, in red) allows us to identify the output’s steady state degradation rates corresponding to the output’s steady states 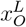 and 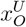 (marked with vertical black lines). We then plot the output’s production and degradation rates against the controller species, *x*_*C*_ (Figure 2C: lower middle plot; blue and red curves respectively). The output’s steady state degradation rates is transferred across from the lower left plot, appearing as a pair of red horizontal lines. For the two production rate curves (in blue) for the output species *x*_*O*_ as a function of *x*_*C*_, one uses fixed 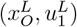 but allows *x*_*C*_ to vary and the other uses fixed 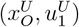 but allows *x*_*C*_ to vary. Finally, we create a plot of the output species’ production and degradation rates against the input value *u*_1_ (Figure 2C: lower right plot; blue and red curves respectively). We can see that a small fold increase in output steady state from 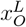 to 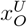 is induced by a small fold increase in output degradation rate at steady state, and a large fold increase in controller steady state from 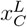 to 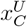 increases the distance between the two output production rate curves as a function of perturbation *u*_1_. The intersection of the output’s production rate and degradation rate curves as a function of perturbation *u*_1_ shows a large change increase in *u*_1_ from 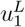 to 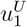, as marked by black vertical lines.

The inverse homeostasis plots make it straightforward to determine the effect of every parameter on perturbation range permitting near-homeostasis, not just the effect of controller parameters (parameters in *x*_*C*_’s rate equation in Figure 1F). Even the simplified algebraic approach^14^ that avoids computing the extremely complicated perturbation to steady state map (Figure 1G) is less intuitive than the graphical insights offered by inverse homeostasis plots (Figure 2D). Once we relax the idealized assumption of perfect integral control, we discovered that it is not just controller parameters that affect homeostatic performance and the details of the output system has a large impact as well. Consistent with the leaky integration reduction approach, increased integration leakiness by decreasing the controller’s maximum production rate *V*_*C*_ and maximum enzymatic degradation rate 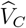 relative to dilution constant *k*_dil_ resulted in much smaller perturbation *u*_1_’s range permitting near-homeostasis (Figure 2E). On the other hand, decreasing affinity of controller toward the promoter of the output by increasing the non-controller parameter *K*_*OC*_ drastically decreases perturbation *u*_2_’s range permitting near-homeostasis, with perturbation *u*_1_ now fixed (Figure 2F). If the a perfect integral controller is present, then perturbation *u*_2_’s range permitting perfect-homeostasis does not change regardless of the value of regulatory parameter *K*_*OC*_ (Figure 2G).

We used the inverse homeostasis plots to articulate a set of parameter steady state relationships that are expected to maximize the range of perturbations the system can handle, and thus vastly widens the region of implementable parameter sets supporting near-homeostasis compared to the integration leakiness reduction approach. Qualitative characteristics of the parameter steady state relationships to maximize homeostatic performance is summarized in Figure 2H, where only the first row minimizes integration leakiness but the following rows tune non-integration leakiness parameters. Using parameters selected according to these rules vastly widens the parameter space supporting near-homeostasis. To show this, we repeated the simulations of Figure 2A, but picking non-controller parameters to optimize perturbation range permitting near-homeostasis to maximally cover the desired range in addition to merely ensuring perturbation sensitivity of the open loop system. The optimization of non-controller parameters noticeably increased the proportion of systems having satisfactory homeostatic performance, enabling at least a 10 fold increase in integration leakiness while still maintaining an equal or greater proportion of parameter sets supporting near-homeostasis (Figure 2I compared to Figure 2A). Among controllers with the same integration leakiness, the inverse homeostasis perspective also identified controllers that resulted in better homeostatic performance (Figure 2H: *V*_*C*_, *K*_*CO*_, *b*_*CO*_). Even if one can guarantee that integration leakiness minimization always results in asymptotically steady states, optimization of the three controller parameters *V*_*C*_, *K*_*CO*_, *b*_*CO*_ on top of non-controller parameters enabled at least 10 to 31.5 fold increase in integration leakiness while still maintaining homeostatic performance.

### The inverse homeostasis perspective enables algorithmic implementation of nearly-homeostatic systems without measuring parameter values

The inverse homeostasis perspective can be applied to identify alterations to an existing system or its controller that would improve its homeostatic performance. Since direct parameter value measurements are not generally feasible in biological systems, we have considered which experimentally observable signals will enable us to determine when the necessary parameter steady state relationships are met to support near-homeostasis. We have found that the shapes of key system components’ steady state response curves map back to both desirable and undesirable parameter steady-state relationships. For our feedback system, we want the controller’s steady state response curve to increase sharply across the desired output range 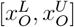, and we want the output’s steady state response curves to have a slope close to *−*1 across the acting range of the controller 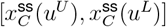 (Figure 1I). Ensuring that key system components’ steady state response curves attain the desired shapes guarantees near-homeostasis.

Before moving on to experimental validation of the approach, we will consider a hypothetical example demonstrating how to determine a sequence of parameter calibrations to shift a set of experimentally measurable steady state response curves to the desired shapes. Suppose we implement an initial randomly generated set of controller *x*_*C*_’s parameters that is hidden from us, resulting in an undesirably flat shape of the controller’s steady states in response to different constant output *x*_*O*_ levels (Figure 3A: gray curve). We select inverse homeostasis plots for the controller rate equation, hypothesize that activator affinity *K*_*CO*_ is too small relative to ideal system output range defined by 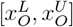, and iteratively determine the expected controller steady states at different output steady states (Figure 3B: top plot). We know our hypothesis is likely correct when the expected steady state response curve generated by the hypothesized inverse homeostasis plots matches the simulated experimental response curve.

**Figure 3:**
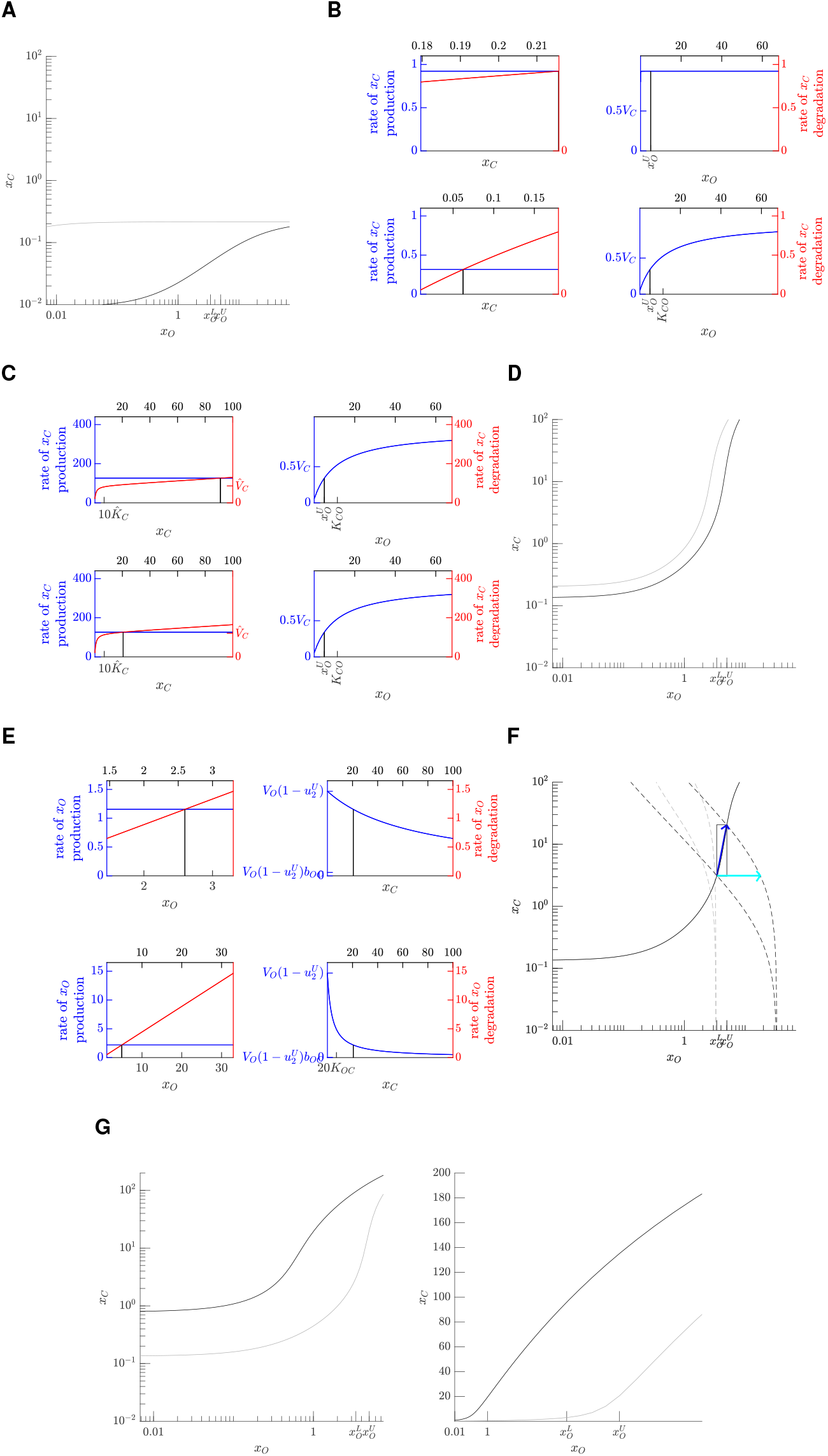
Implementation of a near-homeostatic system by obtaining the desired steady state response curves. Please see Figure 1I for a description of plotting conventions. (A,B,C,D,E,F) Implementation of a near-homeostatic system from a randomly generated set of parameters using steady state response curves (A,D,F) and inverse homeostasis plots (B,C,E). For the inverse homeostasis plots, the vertical black lines indicate the expected steady state value of the component in the first panel in response to the persistent value of the component in the second panel. (G) Comparison of log scale and linear scale when plotting steady state response curves. The inverse homeostasis plots imply that only a logarithmic scale is useful for continuously improving near-homeostasis by directly reflecting fold changes in the steepness of the response curve. In our system, according to the inverse homeostasis perspective, a larger fold change in controller steady state in response to change in the output has greater potential for near-homeostasis. This larger fold change in controller steady state directly translates to greater steepness of the controller response curve on the log scale. Thus, the gray controller response curve have far greater potential for near-homeostasis in the desired output range 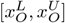 compared to the black controller response curve by having a steeper curve in that output range (left plot). However, when linear scale is used to plot steady state response curves, the gray controller response curve restricted to the desired output range 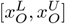 is noticeably less steep than the black controller response curve restricted to the same range and the “overall” slope of the two response curves appear very similar within the plotted output range (right plot).

We then increase activator affinity *K*_*CO*_ (Figure 3B: bottom plots) in an effort to shift the controller’s response curve towards the desired steeper response (like the black curve shown in Figure 3A), which would further validate our hypothesis that *K*_*CO*_ is indeed too small. The controller’s response curve now has a slope of close to 1 over the desired output range. However, the desired slope over the desired output range 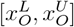 should be much steeper. This lack of the desired steepness is consistent with the controller degradation rate curve in the desired output range 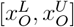 being largely dominated by first order protein dilution rather than saturated protease degradation (Figure 3B: bottom plot).

We then increase the maximal controller transcription rate *V*_*C*_ at a faster rate than increasing maximal protease induced controller degradation rate 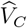, to get the former to be hopefully larger than the latter and to increase the dominance of the maximal protease degradation over protein dilution (Figure 3C: top plot). If our guess is correct, we should observe the slope of the controller response curve becoming much larger than 1. The desired slope does arise, but the output range of ultrasensitivity for the controller response curve is lower than the desired system output range 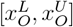(Figure 3D: gray curve)

We then increase the controller’s maximal protease degradation 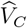 slightly (Figure 3C: bottom plot) to delay the controller’s ultrasensitive response to larger output values and thus shift the steep section of the controller’s response curve into the desired range of output values (Figure 3D: black curve).

We then need to ensure that the system output *x*_*O*_’s regulatory parameters are also within desired ranges (Figure 2H). Starting with an arbitrary set of regulatory parameters for the output system (Figure 3E: top plot), for each of the extreme ends of the perturbation range (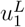 to 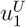), we plot the steady state of the output in response to different controller levels (Figure 3F: gray dashed curves). The plot now includes steady state response curves for both the controller and the output, as well as a black rectangle indicating the desired ranges of controller and system output values. We aim to tune the output response curves at the extreme ends of the perturbation range to intersect with the controller response curve in the black box, and to have the open loop system be at least 4 fold more sensitive to perturbation on the log scale. The initial output response curves in gray has several undesired characteristics: the steady state controller value is too large at each output steady state, the steady state output value is too small at each controller value, and the steepest slope of the output response is also shallower than *−*1 such that open loop has suboptimal sensitivity to perturbation. To shift the system output *x*_*O*_’s response curves toward lower controller *x*_*C*_ levels, higher output *x*_*O*_ levels, and sharper slopes, we consult the top plots in Figure 3E and hypothesize the need to decrease *K*_*OC*_, increase *V*_*O*_, and decrease *b*_*OC*_, respectively, resulting in the ideal black dashed curves shown in (Figure 3F).

In summary, the most current steady state response curves on the log scale are the only observable indicators of homeostatic performance of the current system, but the inverse homeostasis plots allow us to hypothesize the parameter steady state relationships that gives rise to the observable indicators and move those indicators to the desired state. In previous reports^1,13,14^, steady state response curves were plotted on linear horizontal and vertical axis, rather than using logarithmic axes. While linear axes are sufficient for illustrating mechanism of nearly-perfect homeostasis, it can be difficult to determine which of two response curves has greater potential for near-homeostasis (Figure 3G).

### Steady state response curves of key system components can be closely approximated without feedback

A key assumption of our theoretical framework is that steady state response curves of key system components within the negative feedback system can be approximated by auxiliary systems that include only selected aspects of the complete system. The perturbation-dependent steady states of the negative feedback system only reveal a portion of the controller’s steady state response curve. Steady state response curves restricted to a limited window make it hard to estimate parameter steady state relationships. For our system, only the part of the controller response curve overlapping with the dark blue arrow is revealed by characterization of the feedback system (Figure 3E). Auxiliary systems with no feedback must be constructed to measure the full steady state response curves of the controller and of the system output. To characterize the controller response curve, we need to independently generate the system output at different levels and observe the steady state of the controller at each constant output level (Figure 4A). To characterize the output response curve at a particular perturbation, we need to independently generate the controller at different levels and observe the steady state of the output at each controller level (Figure 4C). This output auxiliary system is also known as the open loop system. Since the output and the controller in the auxiliary systems no longer form any feedback loop, it is not clear whether the steady state response curves characterized using the auxiliary system would indeed approximate the steady state behavior of the negative feedback system, even if genetic contexts are kept similar between the auxiliary systems (Figure 4A, 4C) and the negative feedback system (Figure 1F).

**Figure 4:**
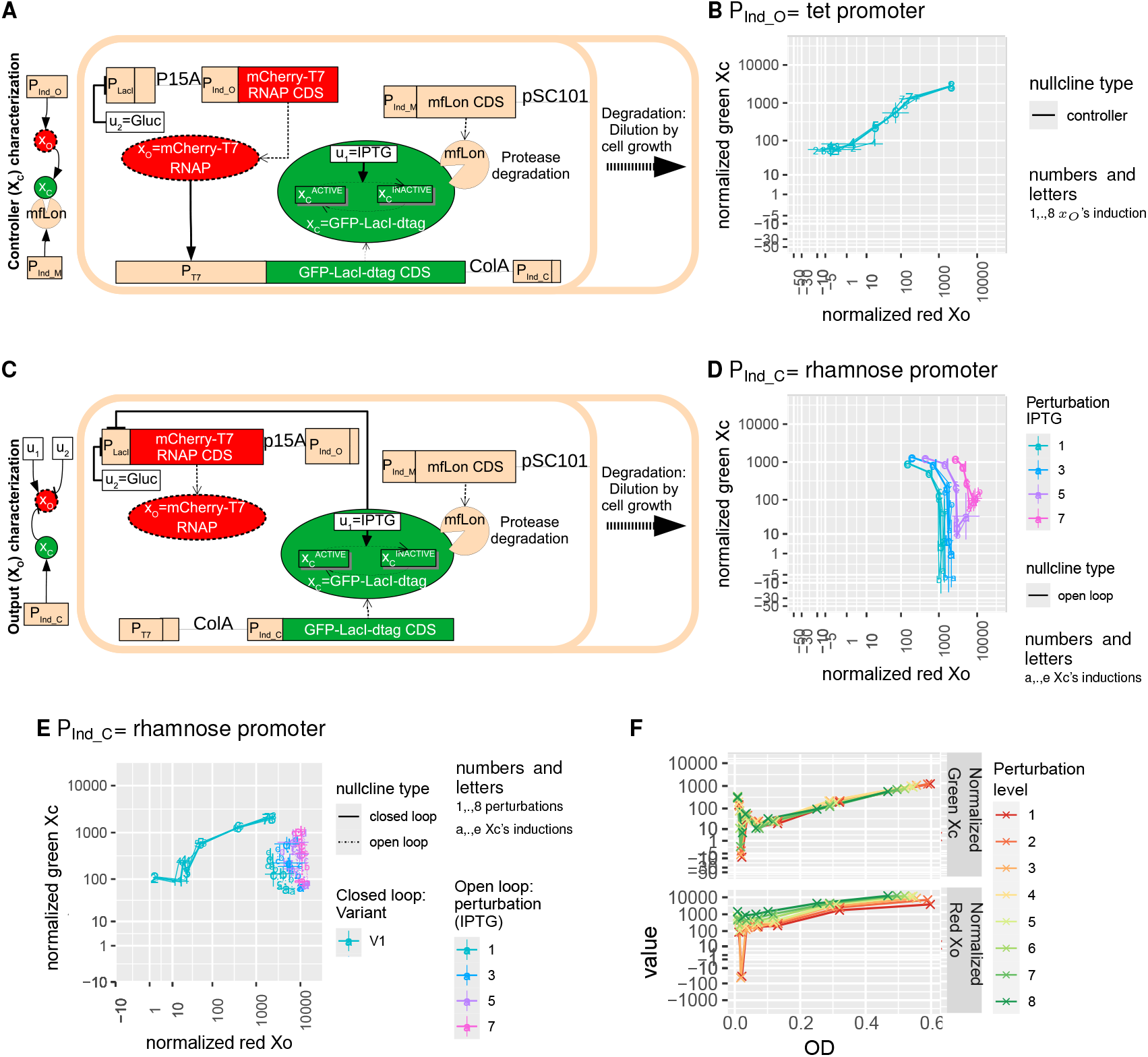
Initial implementation of the T7-LacI negative feedback system. (A) Schematic of the controller auxiliary system. Promoter P_Ind_O_ aims to INDependently generate persistent levels of the Output *x*_*O*_, but its exact identity may change to depending on the limitations discovered. (B) Verifying regulation of the controller by the output. (C) Schematic of the output auxiliary system (also known as the open loop system). Promoter P_Ind_C_ aims to INDependently generate persistent levels of the Controller *x*_*C*_, but its exact identity may change depending on the limitations discovered. Since transcriptional elements have been reported to be dependent on their sequence context, we aimed to keep the genetic sequences surrounding each transcriptional element as similar as possible between the negative feedback system and each auxiliary system. For instance, compared to the negative feedback network, we kept the lac promoter but removed its downstream open reading frame in the controller auxiliary system and did the same thing to the T7 promoter in the output auxiliary system. (D) Verifying regulation of the output by the controller. (E) Initial attempted overlap between the feedback response curve and output auxiliary system’s approximation of the output response curve. For the open loop auxiliary system, the overnight culture contained maximum amount of rhamnose (1 mg/mL). (F) Expression of the open loop auxiliary system across different *E. coli* growth stages under maximal rhamnose induction for both the overnight culture and on measurement day (1 mg/mL). The different inducer levels here refer to different IPTG levels. Please see STAR METHODS for more details. Overnight culture typically reaches OD600 of ≈ 1.

We have experimentally negative feedback system (Figure 1F), the controller auxiliary system with independent output *x*_*O*_ generation performed by the tetracycline inducible promoter (Figure 4A: P_Ind_O_ =tet promoter), and the output auxiliary system with independent controller *x*_*C*_ generation performed by the rhamnose inducible promoter (Figure 4C: P_Ind_C_ = P_rhaBCD_ ^15^). Prior publications^10,16^ placed a fluorescent reporter next to the system output in a polycistronic gene to visualize system output levels. In contrast, we have made both the system output and the controller into fluorescent fusion proteins to visualize the levels of all key system components and to avoid assuming expression linearity between a system component and its adjacent fluorescent reporter (Figure 1F, 4A, 4C).

Since the controller-encoding plasmid within the controller auxiliary system can be co-transformed with the output-encoding plasmid within the output auxiliary system to result in the full feedback system, we can determine whether both the output and controller elements behave as expected using only two tests. Using the controller auxiliary system, we verified that the mCherry-T7 RNA polymerase was able to transcribe from its native promoter by observing that increasing the system output increased the level of the controller (Figure 4B); this is an experimental version of the controller response curves shown in Figure 3A. Using the output auxiliary system, we confirmed that EGFP-tagged LacI was able to repress its cognate promoter by observing that increased controller expression decreased the level of the system output (Figure 4D), in an experimental version of the output response cuves shown as dashed lines in Figure 3F.

In our initial implementation, the output response curve approximated with the auxiliary system is far away from output response curve of the negative feedback system (Figure 4E). If the output response curve of the auxiliary system accurately approximates the output response curve of the negative feedback system, then the steady state of the negative feedback system with no perturbation (Figure 4E: #1 data points of the feedback system) should lie on the output response curve of the auxiliary system in absence of perturbation. If the steady state response curves of key system components cannot be approximated by auxiliary systems, then reducing integration leakiness would be the only viable alternative to implement nearly-homeostatic systems. Since reducing integration leakiness does not consistently result in near-homeostasis and integration leakiness often needs to be unnecessarily low for satisfactory homeostatic performance, we investigated the root cause of the poor approximation by the open loop auxiliary system. We found that dilution of maximally rhamnose induced (1 mg/mL) overnight culture into fresh media containing maximal rhamnose induction resulted in dramatic decrease in the expression of rhamnose inducible reporter during the logarithmic phase of *E. coli* growth (Figure 4F). This observation suggests that rhamnose induction of the controller is not stable across different stages of *E. coli* growth, which could lead to unexpectedly lower repression of the system output; to address this, we replaced the rhamnose-inducible promoter with a series of constitutive promoters of different strengths. Thus, the poor approximation of the output response curve by the auxiliary system does not likely stem from a fundamental issue with our approach, but rather stems from the specific biology involved in our initial design of the output auxiliary system.

Under the stable expression of the controller that is driven by a series of constitutive promoters of different strengths, we found that the approximated output response curve robustly predicted the steady state coordinate of the negative feedback system at the same perturbation. For each constitutive controller expression, we measured how the steady state output responds to different levels of perturbation (Figure 5A: dashed lines in the left panel). Alternatively, for each perturbation, we can connect different steady state output and constitutive controller levels determined by the open loop system to form an approximation of the output response curve (Figure 5A dashed lines in the middle panel). Since the numbers for the closed loop system also represent different perturbations, we see that the output response curve approximations at the different perturbations are close (within 10^1/4^ = 1.8 fold) to the steady states of the five different feedback variants at the corresponding perturbations (Figure 5A middle panel). For instance, the output approximation at perturbation 5 lies within 1.8 fold (0.25 log10 units) to perturbation-5 steady states for all five feedback variants L1 to L5. Compared to induction of the rhamnose promoter (Figure 4F), the expression levels of the controller downstream of constitutive promoters are much stable across different growth stages (Figure 5B).

**Figure 5:**
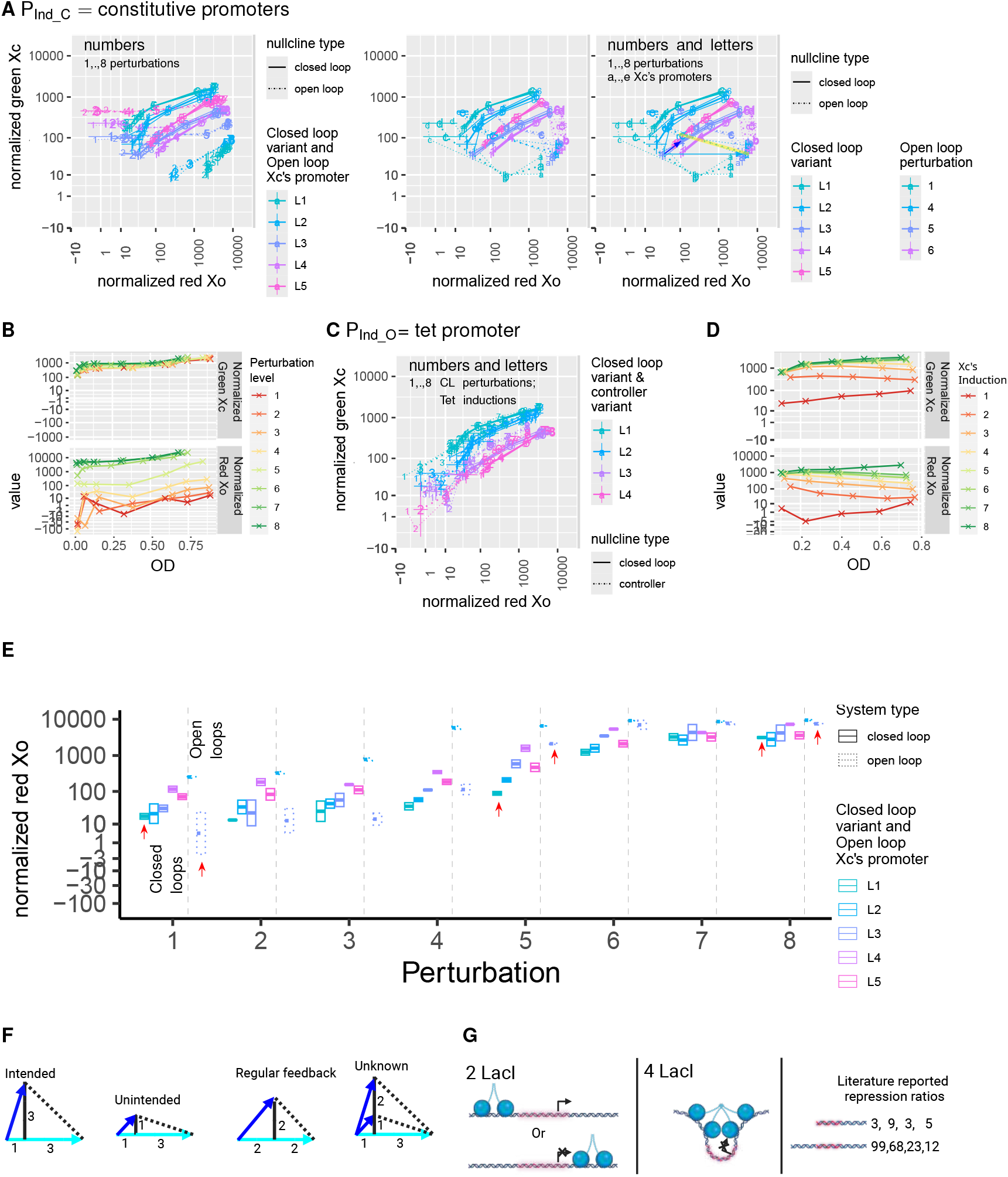
Implementation of a nearly-homeostatic system based on ultrasensitivity of the output response curve. (A) Overlapping of feedback response curves for 5 variants of the feedback system against response curves derived from open loop auxiliary systems at 5 different constitutive controller levels. (B) Expression of a strong constitutively expressed controller at different growth stages. (C) Overlapping of feedback response curves of different feedback system variants against controller response curve approximations for matching variants of both systems. Please see STAR METHODS for more details. (D) Expression of the controller auxiliary system at different growth stages. Specific values of the inducer ATC are present in STAR METHODS. (E) Conventional comparison (Figure 1H) between 5 variants of the feedback system and two open loop auxiliary systems that maximally matches the feedback system variants’s steady state output levels. (F) Simplifying characterization of nearly-homeostatic systems using triangles. The plotting convention here is the same as Figure 1I. (G) The molecular mechanism underlying ultrasensitivity of the output response curve. The LacI repressible promoter upstream of the system output contains two LacI operators that flank both upstream and downstream of the UP element + RNA polymerase core promoter. Doubling the amount of LacI from 50% operator occupancy to 100% operator occupancy not only increases the number of promoters with downstream LacI repression, but also increases the effectiveness of the repression via LacI-tetramer induced DNA looping. Although DNA looping ^17^ by the LacI tetramer in the presence of flanking LacI operators has previously been described to enhance repression efficiency ^18,19^, we are not aware of any studies on the quantitative characterization of LacI’s repression cooperativity and utilization of the cooperative repression in downstream applications. Such DNA looping induced repression cooperativity could be used to create a programmable nearly-homeostatic feedback network with the same topology as the current system. The output T7 activator could be replaced with a CRISPR activator ^20,21^, and controller LacI could be replaced with a multimer-formation-capable CRISPR repressor ^22,23^. To enable multimer formation of the CRISPR repressor, the guide RNA can be modified to contain MS2 hairpins ^21^ that recruit MS2 Coat Proteins coupled with dimerization domains.

We found that the controller response curve could be approximated by the controller auxiliary system, and changing controller parameter results in matching changes in both the steady state response curve of the feedback system and the controller response curve approximation (Figure 5C). For our system, the accuracy of approximation by the auxiliary system is within 0.25 log10 units or 1.8 fold. The R632S variant of our mCherry-T7 (variant L1 and L2) resulted in noticeably more accurate approximation compared to the K631R variant of our mCherry-T7 (variant L3 and L4). Here, constitutive expression of the system output was not required for good controller response curve approximation, probably because 1) the tet inducible promoter was able to maintain much more stable expression of the output compared to the rhamnose-inducible promoter (Figure 5D) and because 2) inducible production of the controller likely reached steady state faster than inducible repression of the system output.

### We discovered a novel non-controller mechanism of near-homeostasis from nature and falsified the proposed zeroth-order-degradation based mechanism of near-homeostasis

Once we verify that the output response curve can be approximated by the open loop auxiliary systems, we can then verify that the system output of closed loop system system indeed changes less than the open loop system between two fixed perturbations (Figure 5E). Traditionally, the output steady state of the feedback system at the initial perturbation is matched to the output steady state of an open loop variant at the same initial perturbation^12^. The output steady state of the feedback system L1 at perturbations 5 to 8 is 2.8 to 20 fold less than that of the matched open loop system L3 (Figure 5E: red arrows). However, between open loop variants L2 and L3, it is difficult to tune constitutive promoters of the controller for the output steady state of the open loop system to match the output steady states of L2-L5 feedback systems at perturbation 1 (Figure 5E). Fortunately, the validated output response curve approximation allows us to predict the behavior of the open loop system at more than the few observed discrete controller levels. As shown in the right panel of Figure 5A, the interpolated open loop equivalent of the L3 feedback system would result in *>* 100 fold increase in output steady state as perturbation 1 is increased to perturbation 4. The L3 feedback system itself would only result in 3.15 fold increase in output steady state for the same increase in perturbation.

According to the right panel of Figure 5A, we have discovered a novel non-controller mechanism of near-homeostasis from nature. We and others have extensively explored how changing parameter steady-state relationships in the controller rate equations enhances near-homeostasis^1^, One such example to improve near-homeostasis is to make the controller response curve more vertical (Figure 5F: panel 1), but the mfLon-producing feedback variant L5 does not yet have desired vertical controller response curve (Figure 5A). Our novel theoretical framework tells us any parameter in the feedback system can dramatically change perturbation range permitting near-homeostasis or equivalently change near-homeostasis of the steady state output at two fixed perturbations (Figure 2H). Here, we show that making the output response curve more horizontal by having a small negative slope also enhances near-homeostasis. The horizontal output response curve is encoded by the output rate equation alone, hence the name non-controller mechanism of near-homeostasis. At low controller levels, the output response curve at perturbation 4 has a slope of *> −*0.5/1.5 = *−*1/3 on the log scale and the controller response curve have a slope of ≈ 1 on the log scale. The relative output change of the feedback system compared to the open loop system is 1/(1 + 3) = 1/4 on the log scale (Figure 5F: panel 2). In contrast, for a regular feedback system whose controller response curve have a slope of 1 and output response curve having a slope of *−*1, the relative output change of the feedback system compared to the open loop system is 1/(1 + 1) = 1/2 on the log scale (Figure 5F: panel 3).

Traditional implementations of feedback systems only measures the output steady state but not the controller steady state^9,10,12^, so it is difficult to tell whether enhanced near-homeostasis is due a vertical controller response curve or flat output response curve (Figure 5F: panel 4). The new non-controller mechanism of near-homeostasis is a result of LacI tetramer induced DNA looping (Figure 5G), which could serve as basis for a programmable nearly-homeostatic system.

### Our theoretical framework cast doubt on whether the intended mechanism of near-homeostasis has been implemented in other feedback systems

In the last few sections, our theoretical framework was useful in diagnosing the direction of parameter change required to improve near-homeostasis without knowing the actual parameter values, and in distinguishing between multiple possible mechanisms of near-homeostasis. Although the theoretical framework should be applicable to all implementations of nearly-homeostatic systems, the accuracy of auxiliary system’s approximation of steady state response curves and the capability to verify implementation of the intended rather than an alternate near-homeostasis mechanism remains to be determined for other feedback systems.

We implemented Sun et al^16^‘s negative feedback network (Figure 6A) and its associated auxiliary networks, by designing our own genetic constructs but making the genetic elements to be as close to the original work as possible. In this feedback system, the output cpf1 and crRNA are both under the control of T7 inducible and CymR repressible promoter. crRNA and cpf1 together repress transcription of the controller T7 RNA polymerase. T7 RNA polymerase also activates its own transcription. The system output is made measurable by placing a sfGFP reporter downstream of the crRNA and the controller is made measurable by placing a mCherry reporter downstream of T7 RNA polymerase. We also constructed a mathematical model based on the network topology

**Figure 6:**
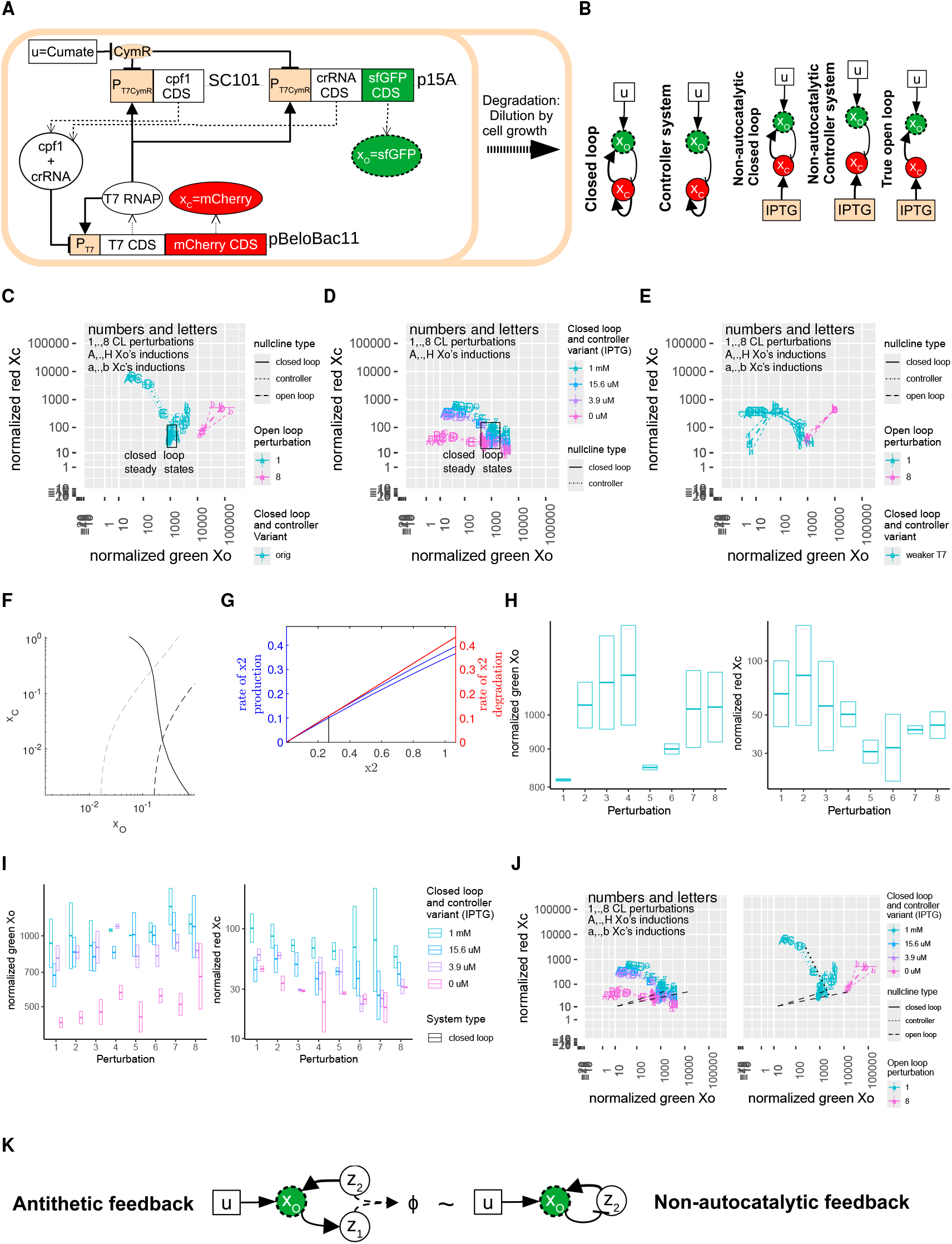
Verifying intended mechanism of near-homeostasis requires measuring steady state response curves of key system components. (A) Schematic of Sun et al ^16^‘s autocatalytic feedback system. (B) Schematics of different variants of Sun et al ^16^‘s feedback system as well as schematics of the controller and the output auxiliary systems. (C) Overlapping the feedback response curve with the controller response curve approximation and output response curve approximation for the original system. (D) Overlapping the feedback response curve with the controller response curve approximation for matching variants of non-autocatalytic systems. (E) Overlapping the feedback response curve with the controller response curve approximation and output response curve approximation for the autocatalytic system with shifted output response curve. (F,G) Expected steady state response curves and inverse homeostasis plot of the autocatalytic system with the intended mechanism of near-homeostasis. (H) Output and controller steady state at different perturbations for the original autocatalytic system. (I) Output and controller steady state at different perturbations for different variants of the non-autocatalytic system. (J) An alternative mechanism of near-homeostasis for Sun et al ^16^‘s autocatalytic feedback system. Black dashed lines represent extensions of the observed output response curves below the basal controller level in the absence of controller induction. The black dotted line represents the possible position of the actual controller response curve rather than an empirical approximation. (K) Similarity between the antithetic feedback system and the non-autocatalytic system.

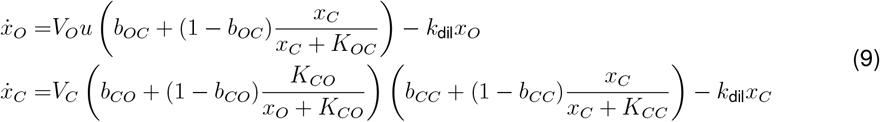

In addition to the original feedback system, we designed and implemented the controller auxiliary system by making the system output no longer dependent on controller levels (Figure 6B). We also designed and implemented the open loop / output auxiliary system by replacing the cpf1 containing plasmid with an empty vector and placing the controller under the control of IPTG inducible promoter (Figure 6B).

For multiple variants of this feedback system, the auxiliary systems accurately approximated the steady state response curves of both the controller and the output. The original version of the feedback system from Sun et al^16^ showed steady states of the feedback system in close proximity (≤ 3.15 fold or ≤ 0.5 log10 unit) with the controller response curve approximation and the 0-perturbation output response curve approximation (Figure 6C). For the feedback systems without autocatalysis but at different maximal controller expression levels, the steady states of the feedback system lay on the controller approximation (Figure 6D). Furthermore, when the original T7 polymerase was changed (R632S) to have lower affinity to its cognate promoter, the output response curves shifted toward lower output levels (Figure 6E). The steady states of the feedback system at minimal and maximal perturbation still lies on the output response curves with matching perturbation (Figure 6E). The steady state response curve of the feedback system at different perturbations also overlaps with the controller response curve approximation (Figure 6E).

Our experiments cast doubt on whether the observed perfect homeostasis in Figure 6C is actually due to autocatalysis of the controller, rather than due ultrasensitive output response curve. If the network is nearly-homeostatic via the intended mechanism of autocatalysis, the controller steady state response curve is expected to show an sharp drop (Figure 6F: solid curve). Here, the inverse homeostasis plot shows how a small fold increase in system output induces a small downward rotation in the production rate curve of the controller, which in turn results in a large drop in controller steady state from ≈ 0.27 to ≈ 0.02 (Figure 6G). However, while the output steady state is perfectly homeostatic for the original feedback system, the controller steady state is also unexpectedly nearly-perfectly-homeostatic rather than showing a sharp drop (Figure 6H). In addition, the original autocatalytic network and the non-autocatalytic networks with IPTG induction had similar degree of near-homeostasis in both the output and the controller (Figure 6H, 6I). This observation is surprising since the autocatalytic network is expected to show a higher degree of near-homeostasis than the non-catalytic network. The ≈ 2 fold increase in output steady state for the non-autocatalytic network without IPTG induction (Figure 6I) tells us how the output response curve should be extended at lower controller levels (Figure 6J: left panel). The extended output response curves explains near-homeostasis of both the output steady state and the controller steady state as well as the slight decrease in controller steady state in many of the feedback variants (Figure 6J: right panel).

Our finding of the autocatalytic network using the ultrasensitive output-response-curve mechanism of near-homeostasis may also hold true for the antithetic integral controllers^10,12^. The output *x*_*O*_ of the anthetic integral control system activates controller *z*_1_, controller *z*_1_ and controller *z*_2_ annihilate each other, the annihilation of the two controllers reduced the amount of activation of the system output by controller *z*_2_ (Figure 6K). Since the output effectively deactivates the second controller *z*_2_, both the antithetic feedback system and Sun et al^16^‘s autocatalytic system have the output deactivating the key controller component (*z*_2_, *x*_*C*_) and the key controller component activating the output. The similarity in the interaction between the output and the key controller component in both feedback systems raises the possibility that the antithetic feedback system was not using the intended mechanism to achieve perfect homeostasis. Thus, it would be worthwhile to measure the active amount of controller *z*_2_ at two different perturbation levels to see whether the controller *z*_2_ steady state barely (Figure 6J) or drastically (Figure 6F) changes while the output is nearly-homeostatic.

In a broader context, network implementation by component characterization may allow us to build ever more complex biological regulatory systems. A more complex biological regulatory system is expected to have greater information processing ability and exhibit more fine-tuned behaviors, which are likely required in therapeutic settings. Such networks may include coupling of multiple feedback networks and integration of logic gates within a feedback network. Component characterization would allow us to accurately pin-point the undesired component behavior that leads to the undesired network behavior. However, a key obstacle to the network implementation by component characterization approach is whether component behavior in absence of the network reflects that component’s behavior within the network. Our work shows that under sufficiently similar genetic contexts and making comparisons using the logarithmic scale rather than the linear scale, component characterization using the auxiliary system satisfactorily approximates component behavior within the network.

## ACKNOWLEDGMENTS

This work was supported by the Natural Sciences and Engineering Research Council (NSERC) of Canada, Discovery grant number RGPIN-2017-06795, and by the New Frontiers in Research Fund (NFRF), Exploration grant numbers NFRFE-2019-00742 and NFRFE-2023-00505. The authors thank Alexander Duggan and Matthew Newman for valuable feedback on the manuscript.

## AUTHOR CONTRIBUTIONS

Z.T. conceived and validated the inverse homeostasis perspective. Z.T. and D.R.M conceived and validated the proposed approach to experimentally implement nearly-homeostatic systems without measuring parameter values. Z.T. and D.R.M wrote the paper.

## RESOURCE AVAILABILITY

### Lead contact

Requests for further information and resources should be directed to and will be fulfilled by the lead contact, David R. McMillen.

### Material availability

Key plasmids generated in this study will be deposited to Addgene.

### Data and code availability

All code used to generate the comparisons between our approach and integration leakiness, the theoretical steady state response curves, and the inverse homeostasis plots are contained in the Supplementary Information zip folder.

## STAR METHODS

## Abbreviations and definitions

ATC: Anhydrotetracycline
CL: closed loop
Error bar: 1 standard deviation of the plotted values. If plotted values are on a log scale, then standard deviation of the log transformed values is computed.
Mean: each mean value takes the average of measurements from two distinct colonies.
Near-homeostasis of *x*_*O*_ at perturbations *u*^*L*^ and: 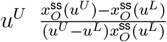
ORF: open reading frame
pLac: IPTG responsive Lac promoter from E coli’s genome; BBa_R0010
RBS: ribosomal binding site
T7: T7 RNA polymerase
UTR: upstream untranslated region of the RNA transcript

## Computational simulations

We built a software package BioSystem Suite in Matlab to perform simulation of a regulatory network and wrap all information related to each regulatory network into a single BioSystem or subclass (e.g. GNetwork) object. We will provide an illustrating example as part of our software overview. Each rate equation for each system component can be specified using a symbolic expression within the “sys” property

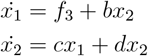

Rate equations may contain symbolic variables that are defined by symbolic expressions within the “flux” property. All symbolic expressions, including those within the “flux” property, can contain symbolic variables that are further defined by the “flux” property

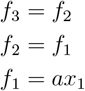

In addition, rate equations may contain symbolic variables whose numerical values can be specified in the “param” property (*a, b, c*). Symbolic variables whose numerical values are not specified in the “param” property (*d*) are automatically determined to be a member of “inputvars” or equivalently perturbations. The time-dependent perturbation values can be specified in the “ut” property. The output of the system can be specified by a symbolic expression within the “output_label” and “output” property

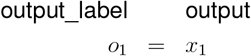

The range of allowable values for each system component is specified in the “xrange” property. The range of allowable values for each parameter is specified in the “prange” property. The initial conditions is specified in the “x0_map” property.

The software package allows rapid change in initial condition, system component ranges, parameter values, parameter value ranges, and time-dependent perturbation values by setting the “locked” property to true. To simulate system component levels across time, Matlab requires the system to be collapsed into a single set of equations when the property “num_sim_only” is false

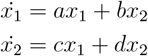

or the expressions arranged in sequentially executable manner

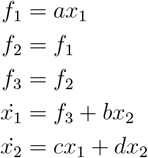

when the property “num_sim_only” is true. The numerically computable function used in simulations is stored in the “numeric_form” and “numeric_form_full” properties. The numerically computable output function is stored in the “numf_output” and “numf_output_full” property. However, collapsing the system or arranging expressions for each simulation takes an unnecessarily long amount of time. We implemented the “locked” property that stores the collapsed system or arranged expressions to prevent recomputation and blocks attempts to change the system in a way that invalidates the stored numerical functions.

The software package contains convenient features for both analytical and numerical analysis of the regulatory network. Features relevant for the purpose of the current study are:

- “convert_to_process” removes numeric symbolic variables from the “param” property such that those variables become perturbations.
- “get_feasible” randomly sets parameter values in the “param” property such that the supplied constraint is satisfied. In particular, supplying an always-true constraint allows one to completely randomize parameter values in the “param” property.
- “simulate_sys” determines the time-dependent system component values of the regulatory system.
- “find_steadystate” finds steady states of the regulatory system, and “find_dose_response” does the same but at different persistent perturbation levels.
- “compute_invhomeo” computes inverse homeostasis plots based on supplied steady state values.
- “compute_sscurves” finds the steady state curve of one system component in response to another system component, at different perturbation levels. Inverse homeostasis plots are also generated to explain the shape of the plotted steady state response curves.

One can see how the above features in combination with different simulation parameters are used to generate different figures in this study (Table 1). Additional example usages of those figures can be found in relevant tests in the BioSystemTests unit testing framework.

**Table 1:**
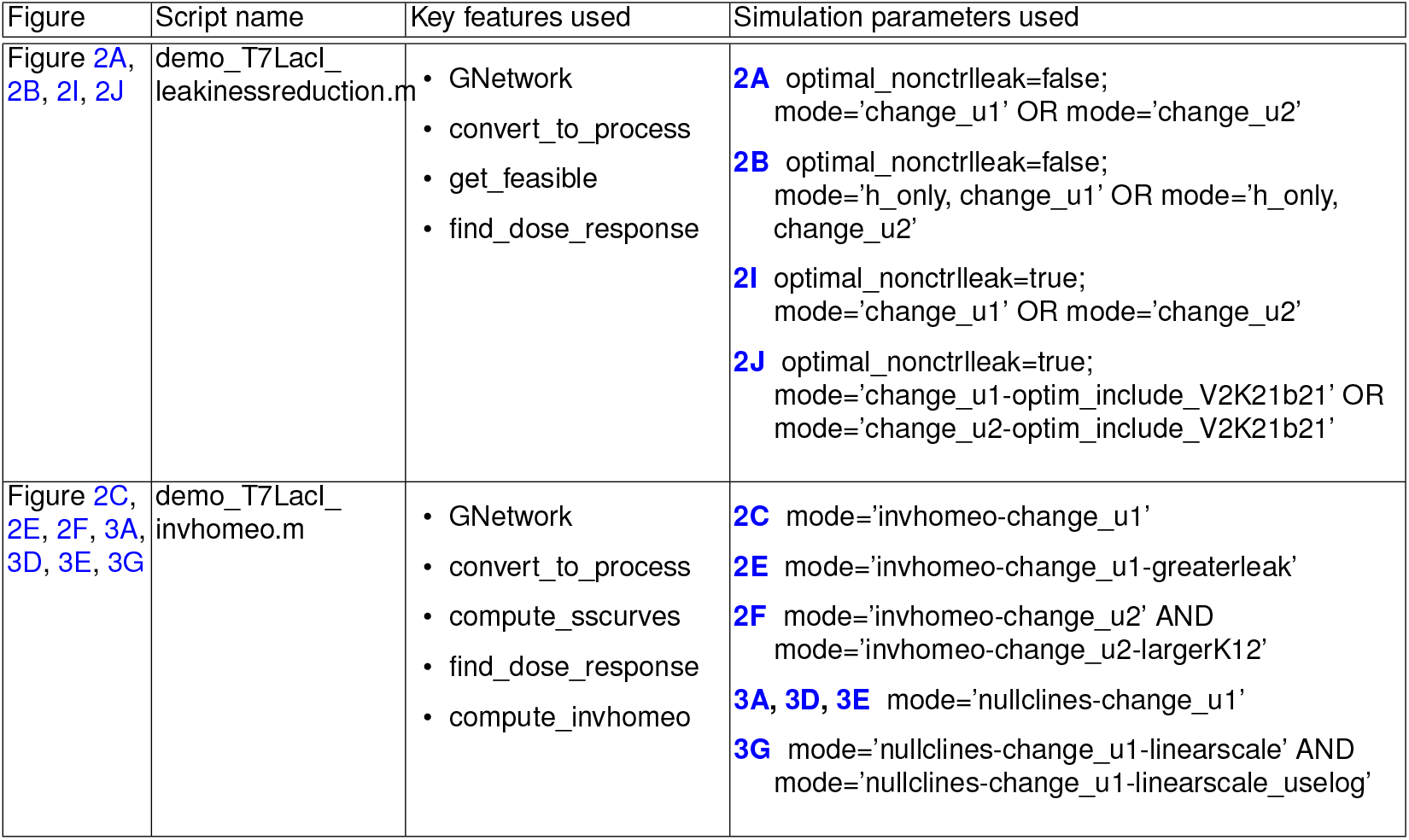
Simulation parameters used to generate different figures.

### Plasmid construction

Plasmids for different system components (Table 2) were built using fragment amplification followed by Gibson Assembly. For each plasmid construct, overlapping DNA fragments were first amplified from appropriate sources using Touchdown Polymerase Chain Reaction (TD-PCR)^25,26^. The error rate of PCR products were minimized^27,28^ using Q5 DNA polymerase (New England Biolabs). PCR products were first visualized on a 1.2% agarose gel to check for approximately correct band sizes and then were purified using the NEB Monarch DNA Cleanup Kit (New England Biolabs). The right set of purified DNA fragments were then assembled at appropriate overlapping sequences using DNA Hifi Assembly Kit (New England Biolabs). Assembly reactions were then transformed into DH5α chemically-competent cells prepared using Inoue’s method^29^ and plated onto LB Agar plates containing the appropriate antibiotics (35*µg/*mL chloramphenicol, 100*µg/*mL carbenicillen, 50*µg/*mL kanamycin, 100*µg/*mL spectinomycin) to select for colonies with assembled constructs. After 18-24h incubation of the LB Agar plates containing the appropriate transformation mixtures at 37ºC, multiple colonies for each assembled construct were inoculated into separate LB media containing appropriate antibiotic. After another 24h rigorous shaking of the liquid cultures at 37ºC, plasmid from each inoculated culture was extracted using Monarch Miniprep Kit (New England Biolabs) and sequenced by Eurofins Genomics at the intended joining regions between different DNA fragments, in order to select for at least one clone with a correctly assembled construct.

**Table 2:**
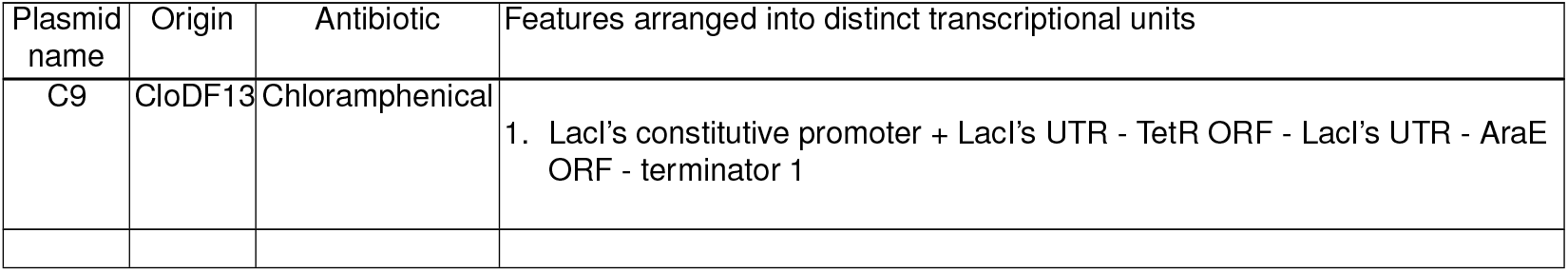
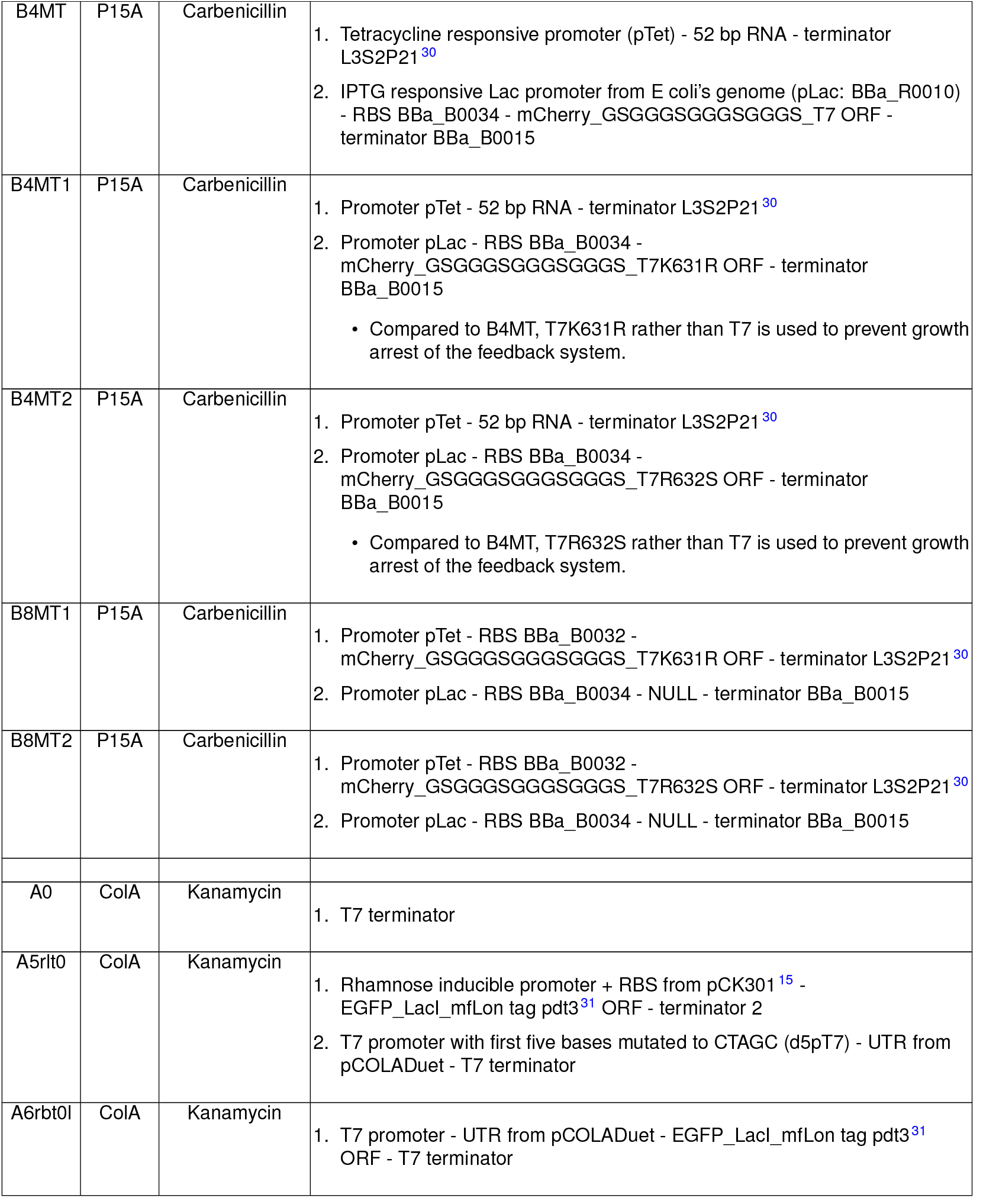
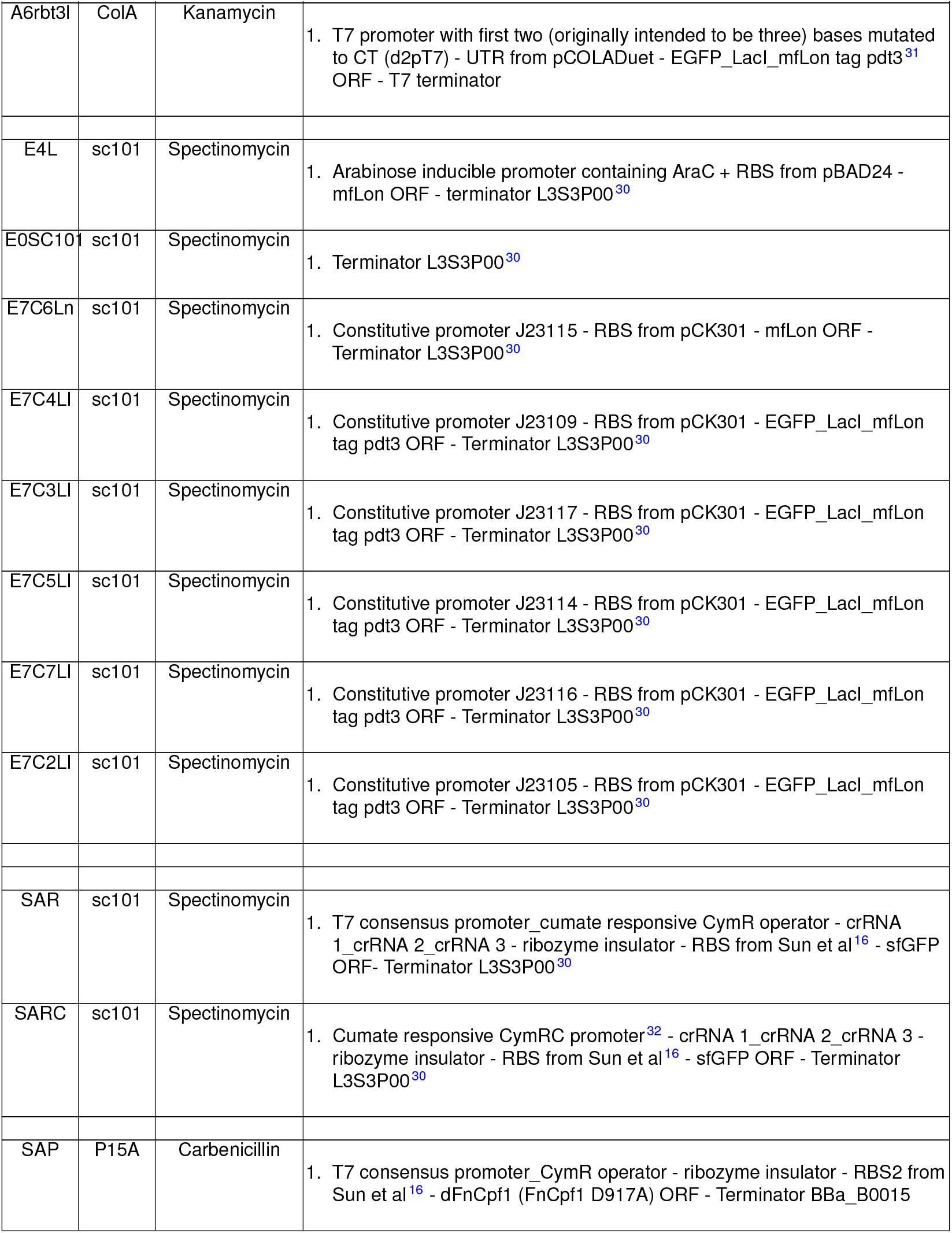
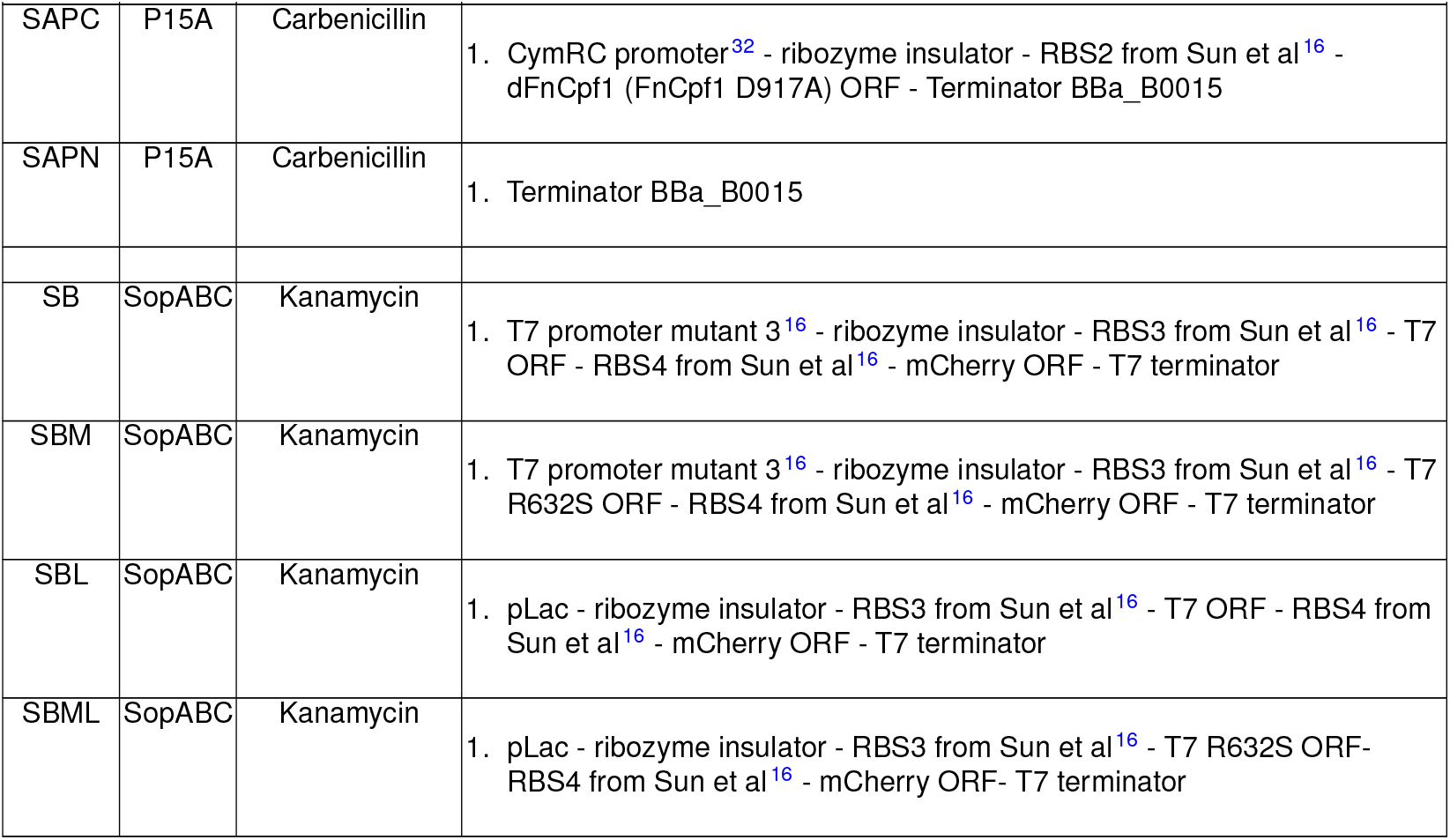
Key features of different plasmid constructs used in this study.

### Characterization of regulatory networks

Cell-density-normalized fluorescence-protein-specific expression values were measured for appropriate combinations of plasmids (Table 3). For each combination of plasmids, electro-competent cells (MG1655 for our feedback system, sAJM1506^32^ for Sun et al’s feedback system) were prepared^33^, transformed using high voltage electroporation and plated onto LB Agar plates containing the appropriate antibiotics (35*µg/*mL chloramphenicol, 100*µg/*mL carbenicillen, 50*µg/*mL kanamycin, 100*µg/*mL spectinomycin) to select for clones containing all plasmids in that combination. Whenever simultaneous transformation with all plasmids did not yield any colonies, we transformed the cells with fewer plasmids and prepared competent cells for transformation with the rest of the plasmids. After 24-48h incubation of the LB Agar plates at 37ºC, 2 colonies were separately inoculated into EZ rich media (no glucose, 0.25% v/v glycerol; teknova) containing appropriate antibiotics (4AB: 20*µg/*mL chloramphenicol + 25*µg/*mL carbenicillen + 15*µg/*mL kanamycin + 25*µg/*mL spectinomycin; 3AB: 50*µg/*mL carbenicillen + 25*µg/*mL kanamycin + 50*µg/*mL spectinomycin) and inducers. After another 18-24h incubation of liquid culture at 37ºC with rigorous shaking to aerate the culture, each liquid culture was diluted 4000 fold (unless otherwise indicated) into EZ rich media (no glucose, 0.25% glycerol) containing appropriate antibiotics and different levels of inducers, in 96 well plates with clear bottoms. The 96 well plates were incubated at 37ºC with rigorous shaking in a microplate shaker (800 rpm, 3 mm orbit; VWR). At different time points of cell growth until all cultures are saturated or 10-13 hours post culture-inoculation, fluorescence (EGFP: 489 *±* 5nm excitation/509 *±* 5nm emission; mCherry: 587 *±* 5nm excitation/610 *±* 5nm emission, fixed z-position and gain) and absorbance (OD600 reflects density of cells, maximum value of 1 for this instrument) measurements were performed twice using TECAN Infinite M1000 Pro plate reader. Normalized expression

**Table 3:** Experimental conditions used to generate different figures.

| Figure | Plasmid combination | Experimental conditions |
| --- | --- | --- |
| 4B | C9 + B8MT2 + A6rbt3l + E4L | No inducer for overnight culture<br>Measurement day inducers: 0, 3.125, 6.25, 12.5, 20, 30, 50, 400 ng/mL ATC (Milipore Sigma) |
| 4D | C9 + B4MT + A5rlt0 + E4L | No inducer for overnight culture<br>Dilution on measurement day: 1:200<br>Measurement day inducers: 0, 0.0156, 0.0625, 0.25, 1 mg/mL rhamnose (Milipore Sigma); for each rhamnose concentration, 0, 1.95, 3.9, 7.8, 15.6, 31.2, 62.5, 125 µM IPTG (Milipore Sigma) |
| 4E, 4F | C9 + B4MT2 + A5rlt0 + E0SC101 | 1 mg/mL rhamnose in overnight culture<br>Dilution on measurement day: 1:400<br>Measurement day inducers: 0, 0.0625, 0.25, 1 mg/mL rhamnose; for each rhamnose concentration, 0, 1.95, 3.9, 7.8, 15.6, 31.2, 62.5, 125 µM IPTG |

Table 3: Experimental conditions used to generate different figures
| Figure | Plasmid combination | Experimental conditions |
| --- | --- | --- |
| 5A,<br>5B,<br>5E | Negative feedback:<br>C9 + B4MT2 + A6rbt0I + E0SC101 (L1)<br>C9 + B4MT2 + A6rbt3I + E0SC101 (L2)<br>C9 + B4MT1 + A6rbt0I + E0SC101 (L3)<br>C9 + B4MT1 + A6rbt3I + E0SC101 (L4)<br>C9 + B4MT2 + A6rbt3I + E7C6Ln (L5)<br>Open loop auxiliary system:<br>C9 + B4MT2 + A0 + E7C4LI (L1)<br>C9 + B4MT2 + A0 + E7C3LI (L2)<br>C9 + B4MT2 + A0 + E7C5LI (L3)<br>C9 + B4MT2 + A0 + E7C7LI (L4)<br>C9 + B4MT2 + A0 + E7C2LI (L5) | No inducer for overnight culture<br>Measurement day inducers: 0, 1.95, 3.9, 7.8, 15.6, 31.2, 62.5, 125 μM IPTG |
| 5C,<br>5D | Negative feedback:<br>C9 + B4MT2 + A6rbt0I + E0SC101 (L1)<br>C9 + B4MT2 + A6rbt3I + E0SC101 (L2)<br>C9 + B4MT1 + A6rbt0I + E0SC101 (L3)<br>C9 + B4MT1 + A6rbt3I + E0SC101 (L4)<br>Controller auxiliary system:<br>C9 + B8MT2 + A6rbt0I + E0SC101 (L1)<br>C9 + B8MT2 + A6rbt3I + E0SC101 (L2)<br>C9 + B8MT1 + A6rbt0I + E0SC101 (L3)<br>C9 + B8MT1 + A6rbt3I + E0SC101 (L4) | No inducer for overnight culture<br>Measurement day inducers: 0, 1.95, 3.9, 7.8, 15.6, 31.2, 62.5, 125 μM IPTG for the feedback systems;<br>0, 3.125, 6.25, 12.5, 20, 30, 50, 400 ng/mL ATC for the controller systems |
| 6C,<br>6H | Negative feedback system:<br>SAR + SAP + SB<br>Controller auxiliary system:<br>SARC + SAPC + SB<br>Open loop auxiliary system:<br>SAR + SAPN + SBL | No inducer for overnight culture<br>Measurement day inducers: 0, 0.488, 0.975, 1.95, 3.9, 7.8, 15.6, 1000 μM cuminic acid (Milipore Sigma)<br>Open loop system is also characterized in the presence and absence of 1 mM IPTG. |
| 6D,<br>6I | Negative feedback system:<br>SAR + SAP + SBL<br>Controller auxiliary system:<br>SARC + SAPC + SBL | No inducer for overnight culture<br>Measurement day inducers: 0, 0.488, 0.975, 1.95, 3.9, 7.8, 15.6, 1000 μM cuminic acid (Milipore Sigma)<br>Each system is also characterized in the additional presence of 0, 3.9, 15.6, 1000 μM IPTG. |
| 6E | Negative feedback system:<br>SAR + SAP + SBM<br>Controller auxiliary system:<br>SARC + SAPC + SBM<br>Open loop auxiliary system:<br>SAR + SAPN + SBML | No inducer for overnight culture<br>Measurement day inducers: 0, 0.244, 0.488, 0.975, 1.95, 3.9, 7.8, 15.6 μM cuminic acid (Milipore Sigma)<br>Open loop system is also characterized in the presence and absence of 1 mM IPTG. |

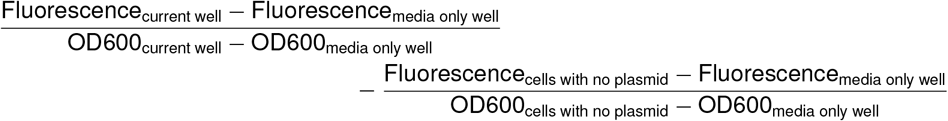

is computed to reflect the amount of EGFP or mCherry per cell in arbitrary units. The above formula first corrects for media-based contribution to fluorescence and absorbance to arrive at fluorescence per cell for both the current well and wells containing cells with no plasmid. The formula then corrects for the amount of non-EGFP or non-mCherry proteins per cell that contribute to the detected fluorescence value per cell. Expression values were averaged between replicate measurements to minimize variability for weakly-expressing cells. To construct steady state response curve of one system component in response to another system component, expression values of fluorescent proteins at a particular inducer concentration were derived from the time point of the highest OD below 0.4 (OD of overnight culture is ≈ 1). Graphics were generated using the GGPlot2 package^34^ in the R programming environment.

Note that the experimental steady state response curves were plotted on the log10 modulus scale rather than the log10 scale. The log modulus scale

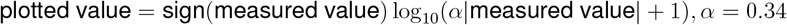

transforms both positive and negative measured values to a hybrid logarithmic scale. Every two sufficiently large, measured values (≥ 3.4*/α* in absolute value) that are ten fold apart are plotted as two points that are nearly a fixed distance apart (0.9 to 1) on the graph. The value of *α* is set such that measurements below the minimum absolute value of 3.4*/α* are close to the noise range of the instrument. The log modulus scale behaves the same as the log scale for sufficiently high measurement values, but also includes the ability to represent negative or barely detectable measurement values.

## Supplementary Information

### SI1 Challenges in experimental parameter estimation

One obstacle to implementing nearly-homeostatic systems is the set of difficulties involved in measuring and targeting parameter values experimentally; and this obstacle precludes direct implementation of simulated parameter values of nearly-homeostatic systems. For the existing system in Figure 1D, we can directly observe the level of the output *x*_*O*_ a function of time by making *x*_*O*_ a fluorescent fusion protein, but parameter estimation remains a challenge because some parameters are not identifiable or multiple local minima are often present during least squared parameter-estimation^5,6^. One way to make all parameters identifiable and ensure a unique global minimum during parameter estimation is to decouple cellular processes, but cellular processes proceed in parallel and cannot be decoupled in a simple way. In our example, we could first calculate the dilution constant *k*_dil_ from the cell’s average doubling rate, and then use the calculated *k*_dil_ to fit production parameters *V*_*O*_, *b*_*OC*_, *K*_*OC*_, *k*_IA→A_*/k*_A→IA_. However, distinguishing LacI’s affinity toward the promoter *K*_*OC*_ from IPTG’s affinity toward LacI *k*_IA→A_*/k*_A→IA_ would require two separate sets of measurements under in-vitro conditions that may not reflect the in-vivo conditions within the cell.

