## Supplemental Data 1 for "Principles underlying implementation of *nearly*-homeostatic biological networks": BioSystem.m

```
%% Simulation of symbolic biochemical system
% Given a symbolic biochemical system. This program joins together various modules that are used to 
% analyze biochemical systems: BioSystem_Base and BioSystem_Homeostasis.
% 
classdef BioSystem < BioSystem_Base & BioSystem_Homeostasis
  methods
    function [obj]=BioSystem(dx,flux,param,x0_map,prange,xrange)
      % constructor method that is the same as BioSystem_Base
      if nargin<=5
        xrange=containers.Map2(dx.keys,repmat({[-Inf,Inf]},1,dx.Count));
        if nargin<=4
          if param.Count>=1
            prange=containers.Map2(param.keys,repmat({[-10,10]},1,param.Count));
          else
            prange=containers.Map2;
          end
        end
      end
      obj@BioSystem_Base(dx,flux,param,x0_map,prange,xrange);
      obj@BioSystem_Homeostasis(dx,flux,param,x0_map,prange,xrange);
    end
  end
end
```
