## Supplemental Data 1 for "Principles underlying implementation of *nearly*-homeostatic biological networks": BioSystem.html

Simulation of symbolic biochemical system 

### Simulation of symbolic biochemical system

Given a symbolic biochemical system. This program analytically computes certain properties and verifies those properties through simulation.

#### Contents

- Specifications
- Design

#### Specifications

This program allows manipulation and simulation of invariant systems of symbolic differential equations. Specifically, it can

1. call and substitute any part of the symbolic system. Common subexpressions is represented by the same symbol and can be substituted togather. Also replace symbols with expressions. Additional modifications can be performed, such as apply conservation, take limits, convert from full system to partial system with inputs, combine multiple systems togather
2. generate feasible parameter sets based on symbolic constraints.
3. compute analytical properties of the system such as Jacobian, output sensitivity, eigenvalues at steady states
4. simulate the symbolic system after conversion to numeric form: simulate for a time-varying input; until it reaches steady state; finding steady state of dynamical system at multiple parameter sets; visualization of solution trajectories to all togather; visualization of trajectory of a component from multiple simulations all together.
5. generate inverse homeostasis plots for the system; generate steady state curves of key system components in response to another system component or perturbation.
6. generate augmented system to find near-homeostasis supporting parameter sets using modified gradient descent algorithms described in homeostasis-backbone.lyx
7. convert the dynamical system with only velocity vectors into a dynamical system with acceleration vector.

#### Design

To perform all specified functions, the program need to store symbolic process in a centralized object and enable it perform actions.

##### Static properties of the dynamical system

- **sys** and **flux** stores the complete dynamical system
- **param** stores parameters values
- **xrange** specifies range of state variables that simulations must obey at all times.
- **prange** specifies ideal parameter range of the dynamical system. This is especially useful for finding feasible parameter sets for a given constraint. When setting this property, range for one parameter is sufficient. Range for other parameters will be derived from the supplied range.
- **x0\_map** specifies initial conditions for simulation
- **output** specifies output of the dynamical system
- **ut** specifies input to the dynamical system
- **int\_tol** integration tolerances to try during simulation
- **int\_tol\_absdiff** is the additional exponent added to int\_tol for AbsTol
- **num\_sim\_only** decides whether to turns off analytical computation of the jacobian and eigenvalues

Using the above basic properties, we can calculate other useful properties for analysis and simulation of the system.

- **statevars** computes state variables of the current dynamical system
- **inputvars** computes missing parameters as inputs to the current dynamical system
- **jacobn** computes numerical jacobian function that evaluates at the set of all real numbers and tries to return individual solution components back to threshold.
- **eigenvalue** uses the jacobian to construct eigenvalue function
- **vecfun\_form** compute complete symbolic vector function expression of by combining information in **sys** and **flux**. This property assumes that no flux has matrix form.
- **numeric\_form\_full** uses **vecfun\_form** to generate a function that can be evaluated numerically for simulation.
- **numeric\_form\_full\_wrapped** generates a numeric form that evaluates at the set of all real numbers and tries to return individual solution components back to threshold but still intend to stay outside the threshold. When the solution component is outside the threshold, dynamics within the original system can draw the solution component back into desired xrange.
- **gen\_numf** generates a function file that is the numeric form of the function. The original function may depend on nested fluxes. These nested fluxes are not expanded to save MatlabFunction from taking a long time to optimize the function.
- **numf\_output** is numeric form of the output function.
- **order\_fluxvars** find a order of flux variables such that the earlier variables do not contain later variables.

To avoid recomputation of most of the above properties, the property **locked** can be set to true.

##### Changing the dynamical system

- **replace** call and substitute a symbol in the dynamical system
- **apply\_conservation** reduces dynamical system by eliminating state variables that can be infered by other state variables.
- **convert\_to\_process** makes add user-specified variables as additional inputs to the dynamical system.
- **take\_limit** compute limit process as certain parameters and fluxes approaches a particular value.
- **convert\_to\_acceleration** convert the dynamical system with only velocity vectors into a dynamical system with acceleration vector.

##### Generate feasible parameter sets

- **gen\_constrs** converts a symbolic set of inequilities into numerical constraints that can be evaluated
- **satisfy\_constrs** determines whether the current parameter set satisfies the constraint.
- **get\_feasible** finds a partial set of parameters that satisfies constraints when combined with a fixed coordinates. At each iteration, a random parameter of chosen coordinates is generated and tested whether it togather with rest of the fixed parameters satisfies the supplied constraint.
- **get\_feasible\_sw** finds a partial set of parameters that satisfied constraints when combined with a fixed coordinate vector in a stepwise fashion.
- **get\_feasibles** finds a number of feasible partial parameter sets satisfying supplied constraints when combined with sets of fixed parameter coodinates. Each feasible partial parameter set is either found at once (**get\_feasible**) or found in specified series of steps.
- **get\_infeasible** calibrates a selected parameter until one of the constraints is not satisfied.

##### Analytical and numerical analysis of the dynamical system

- **output\_sensitivity** computes the symbolic homeostasis indicator of the dynamical system
- **output\_numrelsensitivity** computes the numerical relative sensitivity of the dynamical system using fast matrix inverses.
- **output\_sensitivity\_nolaplaceexp** computes the sensitivity of the dynamical system without using Laplace expansion.
- **find\_ssbranch** computes the symbolic steady state of the system or tries to find steady states numerically.
- **compute\_invhomeo** compute inverse homeostasis plots for the current dynamical system. For each system component, the production rate and degradation rate curves of that system component is plotted against each system component and perturbation. All inverse homeostasis plots for each system component will have the same YAxis scale. The user can specify lower and upper limits of system component levels to plot. The user can also specify which production curves to plot and which to not plot. They can also specify scaling factor for the production and degradation curves.
- **compute\_sscurves** finds the steady state curve of a system component in response to another system component according to a selected subset of rate equations, at different persistent variable levels. A persistent variable may be a regulatory parameter, a fixed system component, or perturbation. Inverse homeostasis plots are also drawn for the last parameter set to show how steady state relationships can be mapped to shapes of nullcline.
- **compute\_invhomeo\_lvl2** uses **compute\_sscurves** to systematically plot how one system component responds to another system component at different persistent variable levels, for each unique pair of system components.
- **gen\_augsys4nearh** generate augmented system to find near-homeostasis supporting parameter sets using modified gradient descent algorithm described in homeostasis-backbone.lyx
- **lm\_allset\_f\_gen** generates a stopping function that takes t,y and outputs whether the current simulation should be stopped. Stopping criteria is satisfied if output allset is fulfilled or rsenso1 doesn't change for a user-defined number of iterations. This generated function is only valid if the simulation does not change p, ut, numeric\_form\_full, and numf\_output\_full, in order to maintain maximum simulation speed. Stored information is frozen in place such that the same stop condition can be regenerated again.

##### Simulation of the dynamical system

- **simulate\_sys** simulate the dynamical system possible during time-varying inputs. The solution storage is updated in real time rather than after the simulation is complete. A useful feature of this method is that the input function is only evaluated during the simulation. When u(t) utilizes past solution values, the aforementioned feature allows us to check u(t) generated during simulation process is the same as u(t) generated after the simulation process. In addition, this function assumes that rates are zero outside of the box bounded by xrange. The simulator is set up such that out of range solution trajectory is forced near the range boundary but still intend to stay outside the range.
- **simulate\_sys\_adam** finds uses gradient descent for a very bumpy objective function.
- **view\_solution** uses **simulate\_sys** to simulate the dynamical system and views the solution of selected outputs all togather in a single plot. To enable viewing of different concentration scales in the same plot, a custom scale including the log modulus scale could be used.
- **find\_steadystate** finds steady state of the dynamical systen. It computes time trajectory using **simulate\_sys** and take state variable values in a time window when the variables stop changing (i.e. fluctuation <=1e-4 relatively or absolutely, whichever one comes first).
- **find\_dose\_response** uses **find\_steadystate** to find steady state of the dynamical system at different inputs.
- **find\_homeostasis** finds steady state of dynamical system at different multiple sets of parameters. For each parameter, whether generated at random or supplied, **find\_dose\_response** is called to find steady steady state that that parameter set; various metrics important to homeostasis are also computed.
- **current\_sol** finds current solution in progress.

###### Properties of homeostatic processes

- **enrich** increase percentage of homeostatic stimulations belonging to a specific category and simulate the corresponding limit process.

1. Based on instructions given, create a limit process.
2. Either alter parameter ranges of the processes OR define favored parameters by a
   set symbolic inequalities
3. Generate favored parameters for the actual process. Certain parameters can be
   fixed while letting others vary
4. For both limit process and actual process, simulate steady states for all favored
   parameters

##### General operators

- **combine** concatnates a series of BioSystem objects into a single BioSystem object after checking for conflicting parameter values (param), fluxes (flux), system rate equations (sys), state variable initial values, (x0\_map), state variable ranges (xrange), and parameter value ranges (prange). Outputs need to be respecified.
- **findnhp\_fmincon** uses the original system A with a single output, augmented system A\_aug that tracks progress of discretization and fmincon to find and verify nhp, as well as output any troubleshooting graphs. The signature of this static method is the same as **findnhp**
- **findnhp** uses the original system A and the gradient descent based augmented system to find and verify nhp, as well as output any troubleshooting graphs.

1. Stage 0: a near-homeostasis supporting continuous topological parameter and parameter set
   is found.
2. Stage 1: Every topological parameter very close to a discrete value is first discretized
   in a manner that satisfy additional constraints after discretization.
3. Stage 2.1: Rest of the topological parameters are discretized to the closet discrete
   values.
4. Stage 2.2a: Undiscretized topological parameters from stage 1 is discretized in manner
   that is biased toward the closest discrete topological parameter value. Multiple trials
   of this stage can be run.
5. Stage 2.2b: Some undiscretized topological parameters from stage 1 is discretized in manner
   that is biased toward the closest discrete topological parameter value. Multiple trials
   of this stage can be run.
6. Stage 3: Undiscretized topological parameters from stage 2.2b is discretized in a manner
   that is biased toward the closest discrete topological parameter value. Multiple trials
   of this stage can be run.

- **util\_topotree\_inc** increases position of topology parameters to be discretized according to the manner described below. Currently, this function is not committed to repository, as it might not be very useful.

1. The first parameter is discretized to lower value and nhdt&p finding attempt is
   made
2. The first parameter is discretized to upper value and nhdt&p finding attempt is
   made, if the previous attempt is not successful
3. The second parameter is discretized to lower value and nhdt&p finding attempt is
   made, if the previous attempt is successful
4. The third parameter is discretized to lower value and nhdt&p finding attempt is
   made, if the previous attempt is successful
5. The third parameter is discretized to upper value and nhdt&p finding attempt is
   made, if the previous attempt is not successful
6. The second parameter is discretized to upper value and nhdt&p finding attempt is
   made, if the previous attempt is not successful

##### Misc helpers

- **fullexp** expands symbolic expressions in the first arguments based on expression pairs recursively up to ten times or until relevant expressions are fully expanded
- **view\_solutions** visualizes solution trajectories of a single output from multiple simulations in one plot and tiles plots of different outputs in one figure. The time axis of the solution trajectories should be able to be coordinated.
- **classify\_findnhp** classifies a table of parameter searches into different categories and displays relative proportion of weighted searches for each category. For each row, each combination of non-weight columns that are all true gives rise to its own category.
- **fmincon\_genfunc** generates relative sensitivity function and nonlinear constraint function for findnhp\_fmincon based on a subset of sorted poptx. The functions are generated from the augmented system.
- **map\_Aaug2A** maps a numeric function of A\_aug (p',x',u) to a function that takes in (p,x,u,p\_extra) of A or vice versa. p\_extra is the vector of extra parameters of the first BioSystem object that does not appear in the second BioSystem object.
- **subsd1** performs substitution of first derivative with the actual function for a cell array of symbolic expressions.

Published with MATLAB® R2022b
