## Supplemental Data 1 for "Principles underlying implementation of *nearly*-homeostatic biological networks": apply_conservation.m

```
function apply_conservation(obj,conserv_vars,total_var,elim_var)
%given conservation law, outputs the equivalent system/plant after 
% conservation law is applied.
%input:
%conserv_vars - {<var_name>,coefficient;...}; conservation law can include
% non-state variables
%total_var - <var_name>; name of total variable
%elim_var - <var_name>; name of variable to be eliminated

%check for correct input
if size(conserv_vars,2)~=2
    error('form of conserv_vars is not correct');
end
conserv_vars=conserv_vars(ismember(conserv_vars(:,1),obj.statevars),:);

%check that conservation law actually holds
coeff=[conserv_vars{:,2}];%row vector
result=coeff*obj.vecfun_form(ismember(obj.statevars,conserv_vars(:,1)));
if result~=0
    error(['proposed conservation law does not hold. Leftover terms: ',...
        char(numTostr(result))]);
end
%check elim_var is a variable in conserv_vars
if ~ismember(elim_var,conserv_vars(:,1))
    error('elim_var must be a member of conserv_vars');
end
%total_var cannot be part of state variables
if ismember(total_var,obj.statevars)
    error('total_var must not be a member of state variables!');
end
    
%applying conservation law
obj.param(total_var)=NaN;
obj.x0_map.remove(elim_var);
obj.xrange.remove(elim_var);
obj.sys.remove(elim_var);
obj.sys=sub_4_param(obj.sys,containers.Map(elim_var,...
    solve(coeff*sym(conserv_vars(:,1))==sym(total_var),sym(elim_var))));
obj.flux=sub_4_param(obj.flux,containers.Map(elim_var,...
    solve(coeff*sym(conserv_vars(:,1))==sym(total_var),sym(elim_var))));
end
```
