## Supplemental Data 1 for "Principles underlying implementation of *nearly*-homeostatic biological networks": BioSystem_Base.m

```
%% Simulation of symbolic biochemical system
% Given a symbolic biochemical system. This program analytically computes
% certain properties required for simulation and simulates the dynamical system.
%% Specifications
% This program allows manipulation and simulation of invariant systems of symbolic
% differential equations. Specifically, it can:
%
% # call and substitute any part of the symbolic system. Common subexpressions is represented
%   by the same symbol and can be substituted togather. Also replace symbols
%   with expressions. Additional modifications can be performed, such as apply conservation,
%   take limits, convert from full system to partial system with inputs,
%   combine multiple systems togather
% # generate feasible parameter sets based on symbolic constraints.
% # compute analytical properties of the system such as Jacobian, output
%   sensitivity, eigenvalues at steady states
% # simulate the symbolic system after conversion to numeric form:
%   simulate for a time-varying input; until it reaches steady state;
%   finding steady state of dynamical system at multiple parameter sets;
%   visualization of solution trajectories to all togather; visualization
%   of trajectory of a component from multiple simulations all together.
% # tries to make the solution trajectory reach its final steady state faster, converting
%   the dynamical system with only velocity vectors into a dynamical system
%   with acceleration vector.
%
%% Design
% To perform all specified functions, the program need to store symbolic process in a
% centralized object and enable it perform actions.
classdef BioSystem_Base < matlab.mixin.Copyable
  %%
  % <html><h3> Static properties of the dynamical system</h3></html>
  %
  % * *sys* and *flux* stores the complete dynamical system, but sys.values must only contain scalar
  %   symbolic expressions. 
  % * *param* stores parameters values
  % * *xrange* specifies range of state variables that simulations must obey at all times.
  % * *prange* specifies ideal parameter range of the dynamical system. This is especially
  %   useful for finding feasible parameter sets for a given constraint. When setting this
  %   property, range for one parameter is sufficient. Range for other parameters will be
  %   derived from the supplied range.
  % * *x0_map* specifies initial conditions for simulation
  % * *output* specifies output of the dynamical system
  % * *ut* specifies input to the dynamical system
  % * *int_tol* integration tolerances to try during simulation
  % * *int_tol_absdiff* is the additional exponent added to int_tol for AbsTol
  % * *num_sim_only* decides whether to turns off analytical computation of the
  %   jacobian and eigenvalues
  %
  % Using the above basic properties, we can calculate other useful
  % properties for analysis and simulation of the system.
  %
  % * *statevars* computes state variables of the current dynamical system
  % * *inputvars* computes missing parameters as inputs to the current
  %   dynamical system
  % * *jacobn* computes numerical jacobian function that evaluates at the
  %   set of all real numbers and tries to return individual solution
  %   components back to threshold.
  % * *eigenvalue* uses the jacobian to construct eigenvalue function
  % * *vecfun_form* compute complete symbolic vector function expression of
  %   by combining information in *sys* and *flux*. This property assumes
  %   that no flux has matrix form.
  % * *numeric_form_full* uses *vecfun_form* to generate a function that can be
  %   evaluated numerically for simulation.
  % * *numeric_form_full_wrapped* generates a numeric form that evaluates at the
  %   set of all real numbers and tries to return individual solution
  %   components back to threshold but still intend to stay outside the threshold.
  %   When the solution component is outside the threshold, dynamics within
  %   the original system can draw the solution component back into desired
  %   xrange.
  % * *gen_numf* generates a function file that is the numeric form of the
  %   function. The original function may depend on nested fluxes. These
  %   nested fluxes are not expanded to save MatlabFunction from taking a
  %   long time to optimize the function.
  % * *numf_output* is numeric form of the output function.
  % * *order_fluxvars* find a order of flux variables such that the earlier variables do not
  %   contain later variables.
  %
  % To avoid recomputation of most of the above properties, the property *locked* or *sol_locked* 
  % can be set to true.
  properties
    % number of integration iterations to check whether max run time has been reached.
    iter_check=1e4
    % Maximum number of tries used to find steady state of the system.
    ss_iter_limit=3
    % ode solver to use; solver input must be the same as ode15s
    odesolver=@ode15s
    % a function having inputs t,y and output a scalar indicating whether simulation
    % should be prematurely stopped. This function will be called once after
    % each solution iteration.
    allset_f=@(t,y)0;
    s_tm=[] % timetracker during solution finding
  end
  properties(Dependent)
    % dynamics of the system. map container; {<var_name>, <symbolic expr of fluxes,
    % parameters,and inputs>}; Values of the map container can be characters when
    % setting this property. Compared to struct, map container has native support
    % for simultanuous multi-field extraction.
    sys
    %fluxes within the system; map container; {<flux_name>, <symbolic expr>}
    % Values of the map container can be characters when setting this property.
    % Fluxes can refer to itself.
    flux
    % map container; {<param_name>, <param_value>}.
    param
    % output of the system; symbolic column vector. Consists of state variables, fluxes and
    % parameter values. Value can be a cell array of strings as well when setting this
    % property.
    output
    % column cell array of characters representing label for output of the system;
    output_label
    % map container; {<start variable>,<value>}. Keys match sys.keys when retrieved.
    x0_map
    % container map; {'<x1>',[<from>,<to>]\[;...\]}. Keys match sys.keys when retrieved.
    xrange
    % container map; {'<param1>',[<from>,<to>][;...]}. Keys match parameters when retrieved.
    prange
    % input function of time; function handle; input is scalar; output is a column vector
    ut
    % cell array of characters with 2 columns where first column contains function derivatives in
    % the form of 'D([n1],<func>)' with no spacing and the second column contains names of functions
    % corresponding to the derivatives.
    deriv_pairs

    x0 %initial state to simulate from; column vector
    statevars %names of all state variables inside the system; row cell array
    inputvars %names of all input variables inside the system; row cell array
    %Jacobian of the substituted system; function handle; @(x,u)...;
    jacobn
    %Jacobian of the substituted system; function handle; @(p,x,u)...;
    %If num_sim_only is false, then compute exact jacobian. If num_sim_only is true,
    %returns numerically determined jacobian.
    jacobn_full
    %function that evaluates eigenvalue; function handle; @(x,u) ...;
    %outputs an error when num_sim_only is true
    eigenvalue
    %Symbolic vector expression of the current system. Flux
    %variables that are present in flux expressions are first substituted
    %up to 5 times or until no flux variables are present. The substituted
    %fluxes are then substituted into the system.
    vecfun_form
    %numeric form of the class instance used for simulation;
    %function handle - @(x,u)f(p,x,u); The underlying function is stored as a
    %file. x and u are both column vectors. p stays fixed until the next numeric_form generation.
    numeric_form
    %numeric form of the class instance used for simulation;
    %function handle - @(p,x,u)f(p,x,u); The underlying function is stored as a
    %file. x and u are both column vectors.
    numeric_form_full
    % numeric_form_full but able to be evaluate at outside of xrange.
    numeric_form_full_wrapped
    %numeric form of output function. @(<statevars column vector>,
    % <possibly inputs>) <returns column vector>; The underlying function
    % is stored as a file.
    numf_output
    %numeric form of output function. @(p,<statevars column vector>,
    % <possibly inputs>) <returns column vector>; The underlying function
    % is stored as a file.
    numf_output_full
    % numf_output_full but able to be evaluated at outside of xrange
    numf_output_full_wrapped
    % Finds current solution in progress. Valid outside of a simulation only if
    % sol_locked is true. either [] or a structure / ref1_array with methods 'read'(r,c), 
    % 'mod'(r,c,rep), 'subset'(r,c) and 'size'(). The functions queries an underlying matrix where 
    % each column represents a separate time point and the first row indicates the times. 
    % Should only be writable by simulate_sys. The each column of the array stores solution at one
    % time point. The first row denotes the times of the solution. 
    current_sol
    % Returned last index of current_sol that is not nan
    lastsimindex


    % vector of relative integration tolerances in log10 units to try. Absolute tolerance
    % adds int_tol_absdiff on top of each element
    int_tol
    int_tol_absdiff
    % Indicates whether simulation is purely numerical based. True turns off
    % analytical computation of the jacobian and outputs numeric form of the rate equations
    % into a file to use fluxes to avoid repeated calculations of the same
    % thing. As such, the variable storing the numeric rate equation points
    % to a file name, which means subsequent updates to the system can result in update
    % to that file. In this mode, flux values can be a matrix.
    % If num_sim_only is off, numeric form of rate equations is stored as a true
    % function in memory, so updating the system cannot result in update to the variable
    % already storing the numeric form. In addition, when num_sim_only is off, each flux
    % value must be a scalar: subs(str2sym('f(rates)'),sym('rates'),sym('x',2))
    % unexpectedly returns [ f(x1_1), f(x1_2); f(x2_1), f(x2_2)]. Here, f=extract_array.
    %
    num_sim_only
    % logical vector or scalar indicating whether out of bound solution component
    % would prematurely stop the simulation. If logical vector, then the length has to
    % be the same as number of state variables.
    crash_sim
    % logical scalar; when both the current property is true and num_sin_only is true, 
    % numeric_form and numf_output generates mex files. 
    generate_mex 
  end
  properties(Dependent)
    % Locking the dynamical system
    %
    % The Jacobian, numerical jacobian, numerical form,
    % numerical form of output and input variables are all supposed to be implemented as
    % dependent properties. However, retrieval of these properties as dependents is slow.
    %
    % Design
    %
    % When the system is not locked with respect to sys, flux, output, output_label,
    % parameter keys, paremeter range keys, state variable range keys, x0 keys, 
    % num_sim_only, deriv_pairs, generate_mex,
    % relavant properties (jacobn, numeric_form, numf_output) are recalculated each time they 
    % are retrieved. When the system is locked, those properties should not be recalculated 
    % each time they are retrieved. When previously unlocked, locking the system initiates 
    % recalculation of relevant properties.
    %
    % Each of the aforementioned properties have storage and as well as dependent
    % variant. If the system is unlocked, performs clear previous calculated property in
    % storage if need to be cleared. If the system is locked, does not clear previous
    % calculated property in storage. If storage is empty, calculate property and put
    % calculated property in storage before final retrieval. If locking status changes,
    % delete the calculated property in storage all at once.
    %
    % Locked dynamical system should prevent modifications to relevant properties.
    % This is accomplished by having a storage for each above property. When system is locked,
    % cannot modify storage (box and content inside the box) for sys and flux, but allow
    % conditonal modification of parameter values but not keys. To allow parameter key
    % modification in event of storage box modification, we keep track of an intended set of
    % keys to compare contents of the modified storage box to. If storage is modified but
    % content keys are locked, then it is up the content to detect key changes.
    % When system is unlocked, allow modifications of the above properties.
    locked
    % Locking the solution so multiple analysis of the same solution can be
    % performed without resimulation.
    %
    % Design
    %
    % When the solution is locked, the system is first locked but with the following additional
    % locks: parameter values, parameter range's values, sv range's values, x0's values, 
    % ut, integration tolerances (2), crash_sim. For handle based property values, the
    % contents of the handle is completely locked as well. For current_sol, the underlying storage
    % is cleared immediate after simulation when sol_locked is false, to prevent
    % further changes in the system that makes the solution invalid.
    % If locking status changes, clear all relevant calculated properties in storage
    %
    % In addition to the aforementioned properties, simulate_sys will
    % retrieve stored solution if solution lock is in place and resimulation
    % does not have to be performed; otherwise, simulate_sys will perform
    % resimulation and update storage. Solution storage should ONLY be
    % changed by simulate_sys and sol_locked, and read by current_sol.
    sol_locked
  end
  properties (Hidden, Access='private') % storage for dependent properties
    s_locked=false
    s_sol_locked=false

    s_sys % storage for sys
    s_flux % storage for flux
    s_param % storage for param
    s_output % storage for output
    s_output_label % storage for output_label
    s_x0_map % storage for x0_map
    s_xrange % storage for xrange
    s_prange % storage for prange
    s_deriv_pairs={} % storage for derivative pairs
    s_inputvars='true empty' % storage for inputvars
    s_generate_mex=false; % storage for generate_mex
    s_param_keys % intended keys for s_param
    s_ut % storage for ut

    s_solution=[] % storage for solution;
    s_int_tol=-3:-2:-7
    s_int_tol_absdiff=[-3,-3,-3]
    s_num_sim_only=false
    s_crash_sim=true; % default value ensures fastest simulation time under most scenarios.
    

    s_jacobn='true empty' %@(param,x,u) size(x) x size(x) matrix function
    % Cell array of multiples of two elements {Function,full file name of the function}.
    % Function has the form @(param,x,u) f(param,x,u) outputs m x 1 vector;
    % m=number of state variables. The first function is the primary function to be used. 
    % When mex file generation is active, then {[2,4]} corresponds to the locations of the mex file
    % and source directory, respectively. 
    s_numeric_form=repmat({@()error('Unitialized function!'),''},1,3);
    % Cell array of multiples of two elements {Function,full file name of the function}.
    % Function has the form @(param,x[,u]) out; outputs and inputs are all
    % column vectors. The first function is the primary function to be used. 
    % When mex file generation is active, then {[2,4]} corresponds to the locations of the mex file
    % and source directory, respectively.
    s_numf_output=repmat({@()error('Unitialized function!'),''},1,3);
  end

  methods(Hidden) %% internal methods - hiding non-sealed handle class methods
    function varargout=addlistener(varargin)
      varargout=cell(1,nargout);
      [varargout{1:end}]=addlistener@handle(varargin{:});
    end
    function varargout=eq(varargin)
      varargout=cell(1,nargout);
      [varargout{1:end}]=eq@handle(varargin{:});
    end
    function varargout=findobj(varargin)
      varargout=cell(1,nargout);
      [varargout{1:end}]=findobj@handle(varargin{:});
    end
    function varargout=findprop(varargin)
      varargout=cell(1,nargout);
      [varargout{1:end}]=findprop@handle(varargin{:});
    end
    function varargout=ge(varargin)
      varargout=cell(1,nargout);
      [varargout{1:end}]=ge@handle(varargin{:});
    end
    function varargout=gt(varargin)
      varargout=cell(1,nargout);
      [varargout{1:end}]=gt@handle(varargin{:});
    end
    function varargout=le(varargin)
      varargout=cell(1,nargout);
      [varargout{1:end}]=le@handle(varargin{:});
    end
    function varargout=lt(varargin)
      varargout=cell(1,nargout);
      [varargout{1:end}]=lt@handle(varargin{:});
    end
    function varargout=ne(varargin)
      varargout=cell(1,nargout);
      [varargout{1:end}]=ne@handle(varargin{:});
    end
    function varargout=notify(varargin)
      varargout=cell(1,nargout);
      [varargout{1:end}]=notify@handle(varargin{:});
    end
  end
  methods %% internal methods - accessor methods
    function value=get.lastsimindex(obj)
      if isa(obj.s_tm,'timetracker')
        value=obj.s_tm.curr_iter;
      else
        value=size(obj.current_sol,2);
      end
    end
    function value=get.current_sol(obj)
      % Output
      % out - matrix where each column corresponds to a different time and first
      % row corresponds to the time vector. The time and state matrix are bundled
      % togather to save on matrix retrieval time.
      if isa(obj.s_solution,'struct') || isa(obj.s_solution,'ref1_array')
        size_ofsolution=obj.s_solution.size();
        value=obj.s_solution.read(1:size_ofsolution(1),1:size_ofsolution(2));
      else
        error('current solution has not been initialized!');
      end
    end
    function value=get.sys(obj)
      value=obj.s_sys;
    end
    function set.sys(obj,value)
      if obj.locked
        error('BioSystem:sys','BioSystem object is locked!');
      elseif isa(value,'containers.Map')
        obj.s_sys=ensure_form(map2map2(value),'symbolic');
      else
        error('container map expected!');
      end
    end
    function value=get.flux(obj)
      value=obj.s_flux;
    end
    function set.flux(obj,value)
      if obj.locked
        error('BioSystem:flux','BioSystem object is locked!');
      elseif isa(value,'containers.Map')
        obj.s_flux=ensure_form(map2map2(value),'symbolic');
      else
        error('container map expected!');
      end
    end
    function value=get.param(obj)
      value=obj.s_param;
    end
    function set.param(obj,value)
      if (obj.locked && ...
          (value.Count~=length(obj.s_param_keys) || ...
          ~all(strcmp(obj.s_param_keys,value.keys)))) ...
          || obj.sol_locked
        error('BioSystem:param','BioSystem object is locked!');
      elseif isa(value,'containers.Map')
        obj.s_param=map2map2(value);
        obj.s_param.locked_k=obj.locked;
        if obj.sol_locked
          obj.s_param.locked_a=true;
        end
      else
        error('container map expected!');
      end
    end
    function value=get.output(obj)
      value=obj.s_output;
    end
    function set.output(obj,value)
      %ensure form of output is set to symbolic
      value=ensure_form(value,'symbolic');
      if obj.locked
        error('BioSystem:output','System output cannot be changed when it is locked!');
      elseif (size(value,2)==1)
        obj.s_output=value;
      else
        error('output:IncorrectValue','Output must be a column vector or cell array!');
      end
    end
    function value=get.output_label(obj)
      value=obj.s_output_label;
      if numel(value)~=size(obj.output,1)
        error('BioSystem:output_label:format',...
          'output_label does not have the same number of elements as output!\n');
      end
    end
    function set.output_label(obj,value)
      %ensure form of output is set to symbolic
      value=ensure_form(value(:),'character');
      if obj.locked
        error('BioSystem:output_label','output_label cannot be changed when it is locked!');
      else
        obj.s_output_label=value;
      end
    end
    function value=get.x0_map(obj)
      k_x=obj.sys.keys; k_x0=obj.s_x0_map.keys;
      if length(k_x)~=length(k_x0) || ~all(strcmp(k_x,k_x0))
        error('BioSystem:getx0_map',...
          ['x0_map does not have exact state variables as sys!\n',...
          'sys.keys are %s\nx0_map.keys are %s.'],...
          strjoin(k_x,','),strjoin(k_x0,','));
      end
      value=obj.s_x0_map;
    end
    function set.x0_map(obj,value)
      if isa(value,'containers.Map')
        if (obj.locked && ...
            (value.Count~=length(obj.statevars) || ...
            ~all(strcmp(obj.statevars,value.keys)))) ...
            || obj.sol_locked
          error('BioSystem:x0_map','x0_map keys are locked!');
        else
          obj.s_x0_map=map2map2(value);
          obj.s_x0_map.locked_k=obj.locked;
          if obj.sol_locked
            obj.s_x0_map.locked_a=true;
          end
        end
      else
        error('container map expected!');
      end
    end
    function value=get.xrange(obj)
      k_x=obj.sys.keys; k_xr=obj.s_xrange.keys;
      if length(k_x)~=length(k_xr) || ~all(strcmp(k_x,k_xr))
        error('BioSystem:getxrange',...
          ['xrange does not have exact state variables as sys!\n',...
          'sys.keys are %s\nxrange.keys are%s.'],...
          strjoin(k_x,','),strjoin(k_xr,','));
      end
      value=obj.s_xrange;
    end
    function set.xrange(obj,value)
      if ~isa(value,'containers.Map')
        error('container map expected!');
      end
      if (obj.locked && ...
          (value.Count~=obj.xrange.Count || ...
          ~all(strcmp(obj.xrange.keys,value.keys)))) ...
          || obj.sol_locked
        error('BioSystem:xrange','xrange is locked!');
      else
        obj.s_xrange=map2map2(value);
        obj.s_xrange.locked_k=obj.locked;
        if obj.sol_locked
          obj.s_xrange.locked_a=true;
        end
      end
    end
    function value=get.prange(obj)
      k_param=obj.param.keys;
      k_prange=obj.s_prange.keys;
      if length(k_param)~=length(k_prange) || ~all(strcmp(k_param,k_prange))
        error('BioSystem:getprange',...
          ['prange does not have exact state variables as param!\n',...
          'param.keys are %s\nprange.keys are %s'],...
          strjoin(k_param,','),strjoin(k_prange,','));
      end
      value=obj.s_prange;
    end
    function set.prange(obj,value)
      if (obj.locked && ...
          (value.Count~=obj.prange.Count || ...
          ~all(strcmp(obj.prange.keys,value.keys)))) ...
          || obj.sol_locked
        error('BioSystem:prange','prange is locked!');
      elseif isa(value,'containers.Map')
        obj.s_prange=ensure_form(map2map2(value),'numeric');
        obj.s_prange.locked_k=obj.locked;
        if obj.sol_locked
          obj.s_prange.locked_a=true;
        end
        k=obj.s_prange.keys;
        for i=1:length(k)
          if ~all(size(obj.s_prange(k{i}))==[1,2])
            error('Form of value not correct!');
          end
        end
      else
        error('container map expected!');
      end
    end
    function value=get.deriv_pairs(obj)
      value=obj.s_deriv_pairs;
    end
    function set.deriv_pairs(obj,value)
      if obj.locked
        error('BioSystem:deriv_pairs',...
          'Property deriv_pairs cannot be set because the system is locked!');
      else
        obj.s_deriv_pairs=value;
      end
    end
    function value=get.int_tol(obj)
      value=obj.s_int_tol;
    end
    function value=get.int_tol_absdiff(obj)
      value=obj.s_int_tol_absdiff;
    end
    function set.int_tol(obj,value)
      if obj.sol_locked
        error('BioSystem:int_tol','int_tol is solution locked!');
      else
        obj.s_int_tol=value;
      end
    end
    function set.int_tol_absdiff(obj,value)
      if obj.sol_locked
        error('BioSystem:int_tol_absdiff','int_tol_absdiff is solution locked!');
      else
        obj.s_int_tol_absdiff=value;
      end
    end
    function value=get.num_sim_only(obj)
      value=obj.s_num_sim_only;
    end
    function set.num_sim_only(obj,value)
      if obj.locked
        error('BioSystem:num_sim_only','num_sim_only is locked!');
      else
        obj.s_num_sim_only=value;
      end
    end
    function value=get.crash_sim(obj)
      value=obj.s_crash_sim;
    end
    function set.crash_sim(obj,value)
      if obj.sol_locked
        error('BioSystem:crash_sim','crash_sim is solution locked!');
      else
        obj.s_crash_sim=value;
      end
    end
    function value=get.generate_mex(obj)
      value=obj.s_generate_mex;
    end
    function set.generate_mex(obj,value)
      if obj.locked
        error('BioSystem:generate_mex','generate_mex is locked!');
      else
        obj.s_generate_mex=value;
      end
    end
    function value=get.ut(obj)
      value=obj.s_ut;
    end
    function set.ut(obj,value)
      if obj.sol_locked
        error('BioSystem:ut','ut is solution locked!');
      else
        obj.s_ut=value;
      end
    end
    function set.locked(obj,value)
      if ~isa(value,'logical')
        error('lock property must be either true of false');
      end

      % If locking status changes, clear all relevant calculated properties in
      % storage
      if obj.s_locked~=value
        obj.s_inputvars='true empty';
        obj.s_jacobn='true empty';
        obj.clear_numf('s_numeric_form');
        obj.clear_numf('s_numf_output');
      end

      % locking and unlocking
      if value==true
        %need to lock all stored dependent properties
        %update storage of dependent properties
        obj.s_locked=true; % should be first
        obj.inputvars; %should be second
        obj.jacobn;
        obj.numeric_form;
        obj.numf_output;
        obj.sys.locked_a=true;
        obj.flux.locked_a=true;
        obj.param.locked_k=true;
        obj.x0_map.locked_k=true;
        obj.xrange.locked_k=true;
        obj.prange;
        obj.prange.locked_k=true;
        obj.s_param_keys=obj.param.keys;
      elseif value==false
        obj.s_locked=false; % stored property should be updated first
        obj.sol_locked=false;
        obj.sys.locked_a=false;
        obj.flux.locked_a=false;
        obj.param.locked_k=false;
        obj.x0_map.locked_k=false;
        obj.xrange.locked_k=false;
        obj.prange.locked_k=false;
      end
    end
    function value=get.locked(obj)
      value=obj.s_locked;
    end
    function set.sol_locked(obj,value)
      if ~isa(value,'logical')
        error('lock property must be either true of false');
      end

      % If locking status changes, clear all relevant calculated properties in
      % storage
      if obj.s_sol_locked~=value
        obj.s_solution=[];
      end

      %locking and unlocking
      if value==true
        obj.locked=true;
        obj.param.locked_a=true;
        obj.prange.locked_a=true;
        obj.x0_map.locked_a=true;
        obj.xrange.locked_a=true;
        obj.s_sol_locked=true;
      else
        obj.s_sol_locked=false;
        obj.param.locked_a=false;
        obj.x0_map.locked_a=false;
        obj.xrange.locked_a=false;
        obj.prange.locked_a=false;
        if obj.locked
          obj.locked=true;
        end
      end
    end
    function value=get.sol_locked(obj)
      value=obj.s_sol_locked;
    end
    function set.x0(obj,value)
      obj.x0_map=containers.Map2(obj.statevars,value);
    end
    function value = get.x0(obj)
      value=cell2mat(obj.x0_map.values(obj.statevars)).';
    end
    function[value]=get.statevars(obj)
      %finds the names of all state variables inside the system
      value=keys(obj.sys);
    end
    function[value]=get.inputvars(obj)
      %finds the names of all input variables inside the system
      if obj.locked==false
        obj.s_inputvars='true empty';
      end
      if strcmp(obj.s_inputvars,'true empty')
        sys=obj.sys.values;
        flux=obj.flux.values;
        sysflux_cchar=cellfun(@(x)evalc('disp(x)'),[sys,flux],'UniformOutput',false);
        sysflux_cchar=regexprep(sysflux_cchar,...
          {'<strong>','</strong>','[ \f\t\v]+'},{'','',''});
        value=cellfun(@(x)symvar(x),sysflux_cchar,'UniformOutput',false);
        funcs=symvarfun(sysflux_cchar);
        value=unique(vertcat(value{:}).');
        value=setdiff(value,funcs.funcname);
        value=setdiff(value,obj.statevars);
        value=setdiff(value,obj.flux.keys);
        value=setdiff(value,obj.param.keys);
        obj.s_inputvars=value;
      end
      value=obj.s_inputvars;
    end
    function value=get.vecfun_form(obj)
      %find symbolic vector expression of the current system.
      value=BioSystem.fullexp(obj.sys,obj.flux);
      value=value.values;
      value=[value{:}].';%make sure it's a column vector
    end
    function value=get.jacobn(obj)
      value=obj.jacobn_full;
      params=cell2mat(obj.param.values).';
      switch isempty(obj.inputvars)
        case true
          value=@(x)value(params,x);
        case false
          value=@(x,u)value(params,x,u);
      end
    end
    function value=get.jacobn_full(obj)
      %finds Jacobian of the substituted system
      if obj.locked==false
        obj.s_jacobn='true empty';
      end
      if strcmp(obj.s_jacobn,'true empty')
        if ~obj.num_sim_only
          Jacobn=jacobian(obj.vecfun_form,sym(obj.statevars));
          Jacobn=obj.subsd1({Jacobn},obj.deriv_pairs);
          Jacobn=Jacobn{1};
          if isempty(obj.inputvars)
            obj.s_jacobn=matlabFunction(Jacobn,'vars',...
              {sym(obj.param.keys).',sym(obj.statevars).'});
          else
            obj.s_jacobn=matlabFunction(Jacobn,'vars',...
              {sym(obj.param.keys).',sym(obj.statevars).',sym(obj.inputvars).'});
          end
        else
          numf_full=obj.numeric_form_full;
          numf_full_wrapped=obj.numeric_form_full_wrapped;
          thres=repmat(sqrt(eps),obj.sys.Count,1);
          % numjac sometimes tries values outside of xrange, so I try to
          % prevent that from happening
          % numf_4jac=numf_full;
          numf_4jac=numf_full_wrapped;
          if isempty(obj.inputvars)
            obj.s_jacobn=@(p,x)numjac(...
              @(t,y)numf_4jac(p,y),nan,x(:,1),numf_4jac(p,x(:,1)),convert2_absthres(thres,x),[],false);
          else
            obj.s_jacobn=@(p,x,u)numjac(...
              @(t,y)numf_4jac(p,y,u),nan,x(:,1),numf_4jac(p,x(:,1),u),convert2_absthres(thres,x),[],false);
          end
        end
      end
      value=obj.s_jacobn;
    end
    function value=get.eigenvalue(obj)
      %find eigenvalue function based on current state of the system
      Jacobn=obj.jacobn;
      if isempty(obj.inputvars)
        value=@(x) eig(Jacobn(x));
      else
        value=@(x,u) eig(Jacobn(x,u));
      end
    end
    eigvalues=eval_eig(obj,x,u);
    function value=get.numeric_form(obj)
      value=obj.numeric_form_full;
      params=cell2mat(obj.param.values).';
      switch isempty(obj.inputvars)
        case true
          value=@(x)value(params,x);
        case false
          value=@(x,u)value(params,x,u);
      end
    end
    function value=get.numeric_form_full(obj)
      if obj.locked==false
        % remove previously stored information
        obj.clear_numf('s_numeric_form');
      end

      if isempty(obj.s_numeric_form{2})
        % update storage with generated function, if appropriate
        obj.s_numeric_form{2}=what('.');
        obj.s_numeric_form{2}=new_fname(...
          obj.s_numeric_form{2}.path,'BioSys_numf_','.m');
        func=obj.sys.values;
        func=[func{:}].';
        obj.s_numeric_form{1}=gen_numf(obj,func,obj.s_numeric_form{2});

        if obj.generate_mex
          % generate mex files
          obj.gen_mex('s_numeric_form');
        end
      end

      value=obj.s_numeric_form{1};
    end
    function value=get.numeric_form_full_wrapped(obj)
      max_reltol=max(10.^obj.int_tol);
      max_abstol=max(10.^(obj.int_tol+obj.int_tol_absdiff));
      xrg=cell2mat(obj.xrange.values.');
      xband=max(abs(xrg).*max_reltol,repmat(max_abstol,size(xrg))*100);
      numf_full=obj.numeric_form_full;
      if isempty(obj.inputvars)
        value=@(p,x)BioSystem.numf_full_wrapper(numf_full,'rates',xrg,xband,p,x);
      else
        value=@(p,x,u)BioSystem.numf_full_wrapper(numf_full,'rates',xrg,xband,p,x,u);
      end
    end
    function value=get.numf_output(obj)
      value=obj.numf_output_full;
      params=cell2mat(obj.param.values).';
      if isempty(obj.inputvars)
        value=@(x)value(params,x);
      else
        value=@(x,u)value(params,x,u);
      end
    end
    function value=get.numf_output_full(obj)
      if obj.locked==false
        % remove previously stored information
        obj.clear_numf('s_numf_output');
      end

      % update storage with generated function, if appropriate
      if isempty(obj.s_numf_output{2})
        obj.s_numf_output{2}=what('.');
        obj.s_numf_output{2}=new_fname(...
          obj.s_numf_output{2}.path,'BioSys_numfout_','.m');
        func=obj.output;
        obj.s_numf_output{1}=gen_numf(obj,func,obj.s_numf_output{2});

        if obj.generate_mex
          % generate mex files
          obj.gen_mex('s_numf_output');
        end
      end

      value=obj.s_numf_output{1};
    end
    function value=get.numf_output_full_wrapped(obj)
      max_reltol=max(10.^obj.int_tol);
      max_abstol=max(10.^(obj.int_tol+obj.int_tol_absdiff));
      xrg=cell2mat(obj.xrange.values.');
      xband=max(abs(xrg).*max_reltol,repmat(max_abstol,size(xrg))*100);
      numf_out_full=obj.numf_output_full;
      if isempty(obj.inputvars)
        value=@(p,x)BioSystem.numf_full_wrapper(numf_out_full,'other',xrg,xband,p,x);
      else
        value=@(p,x,u)BioSystem.numf_full_wrapper(numf_out_full,'other',xrg,xband,p,x,u);
      end
    end
  end
  methods(Hidden,Access=private)
    % helper methods 
    clear_numf(obj,prop); % clear s_numeric_form_full or s_numf_output_full.
    gen_mex(obj,prop); % generate mex files for a partially filled property prop according specs.
  end
  methods(Hidden,Access='protected') %% internal methods - object methods
    %%application specific methods
    function simplify(obj)
      %simplify expression in flux and sys properties
      for j=obj.sys.keys
        obj.sys(j{:})=simplify(obj.sys(j{:}));
      end
      for j=obj.flux.keys
        obj.flux(j{:})=simplify(obj.flux(j{:}));
      end
    end
  end

  %%User intended methods
  methods (Access='protected')
    cpObj=copyElement(obj)
  end

  %%
  % <html><h3> Changing the dynamical system </h3></html>
  % 
  % * *replace* call and substitute a symbol in the dynamical system
  % * *apply_conservation* reduces dynamical system by eliminating state
  %   variables that can be infered by other state variables. 
  % * *convert_to_process* makes add user-specified variables as additional inputs 
  %   to the dynamical system. 
  % * *take_limit* compute limit process as certain parameters and fluxes approaches
  %   a particular value.
  % * *convert_to_acceleration* converts dynamical system with only velocity vectors into a dynamical 
  %   system with both velocity and acceleration vectors. This conversion process also allows the
  %   velocity vector to be modified by the normal vector of the solution trajectory, for coordinates 
  %   with both velocity and acceleration vectors. 
  % 
  % <html><h3> Internal verifications </h3></html>
  % * *verify_mex* verfies consistency between mex function and original function for numeric_form
  %   and numf_output. The verification process is useful when locked, num_sim_only, and
  %   generate_mex all equal to true. 
  % 
  % <html><h3> Generate feasible parameter sets </h3></html>
  %
  % * *gen_constrs* converts a symbolic set of inequilities into numerical
  %   constraints that can be evaluated
  % * *satisfy_constrs* determines whether the current parameter set
  %   satisfies the constraint.
  % * *get_feasible* finds a partial set of parameters that satisfies constraints 
  %   when combined with a fixed coordinates. At each iteration, a random parameter of 
  %   chosen coordinates is generated and tested whether it togather with rest of the 
  %   fixed parameters satisfies the supplied constraint.
  % * *get_feasible_sw* finds a partial set of parameters that satisfied constraints when
  %   combined with a fixed coordinate vector in a stepwise fashion. 
  % * *get_feasibles* finds a number of feasible partial parameter sets satisfying
  %   supplied constraints when combined with sets of fixed parameter coodinates. Each 
  %   feasible partial parameter set is either found at once (*get_feasible*) or found in
  %   specified series of steps. 
  % * *get_infeasible* calibrates a selected parameter until one of the
  %   constraints is not satisfied.
  %
  % <html><h3> Simulation of the dynamical system </h3></html>
  %
  % * *simulate_sys* simulate the dynamical system possible during
  %   time-varying inputs. The solution storage is updated in real time
  %   rather than after the simulation is complete. A useful feature of this
  %   method is that the input function is only evaluated during the
  %   simulation. When u(t) utilizes past solution values, the aforementioned
  %   feature allows us to check u(t) generated during simulation process is
  %   the same as u(t) generated after the simulation process. In addition,
  %   this function assumes that rates are zero outside of the box bounded by
  %   xrange. The simulator is set up such that out of range solution
  %   trajectory is forced near the range boundary but still intend to stay
  %   outside the range.
  % * *current_sol* finds current solution in progress.
  % * *simulate_sys_adam* finds uses gradient descent for a very bumpy objective function.
  % * *view_solution* uses *simulate_sys* to simulate the dynamical system
  %   and views the solution of selected outputs all togather in a single
  %   plot. To enable viewing of different concentration scales in the same
  %   plot, a custom scale including the log modulus scale could be used.
  % * *find_steadystate* finds steady state of the dynamical systen. It computes
  %   time trajectory using *simulate_sys* and take state variable values in
  %   a time window when the variables stop changing (i.e. fluctuation <=1e-4 relatively
  %   or absolutely, whichever one comes first).
  % * *find_dose_response* uses *find_steadystate* to find steady state of
  %   the dynamical system at different inputs.
  methods
    function [obj]=BioSystem_Base(dx,flux,param,x0_map,prange,xrange)
      %constructor method: all inputs will have to be container.Map and is internally 
      %converted to containers.Map2.
      %input:
      %dx - dynamics of the system. map or map2 container;
      % {<var_name>, <symbolic expr / fluxes>}
      %flux - fluxes within the system; map or map2 container;
      % {<flux_name>, <symbolic expr>}
      %param - parameter values; map or map2 container; {<param_name>, <param_value>}
      %x0_map - initial state to simulate from; map or map2 container;
      %prange - OPTIONAL: if no specified, all parameter ranges from -10 to 10. 
      % container map or map2; same form as the property;
      %xrange - OPTIONAL: if not specified, all state space ranges from -Inf to Inf;
      % map or map2 container.
      obj.sys=map2map2(dx);
      obj.flux=map2map2(flux);
      obj.param=map2map2(param);
      obj.x0_map=map2map2(x0_map);
      if nargin<=5
        xrange=containers.Map2(obj.sys.keys,repmat({[-Inf,Inf]},1,obj.sys.Count));
        if nargin<=4
          if obj.param.Count>=1
            prange=containers.Map2(obj.param.keys,repmat({[-10,10]},1,obj.param.Count));
          else
            prange=containers.Map2;
          end
        end
      end
      obj.prange=prange;
      obj.xrange=xrange;
    end
    function delete(obj)
      obj.clear_numf('s_numeric_form');
      obj.clear_numf('s_numf_output');
      obj.delete@matlab.mixin.Copyable;
    end
    function disp(obj)
      %displays BioSystem object; overwrites default display behavior
      
      %system dynamics
      value=[obj.sys.keys;obj.sys.values];
      for j=1:size(value,2)
        value{1,j}=['d/dt(',value{1,j},')'];
        value{2,j}=char(value{2,j});
      end
      fprintf('system dynamics:\n');
      fprintf('%s=%s\n',value{:});
      %outputs
      fprintf('\nsystem output:\n');
      value=obj.output;
      for j=1:size(value,1);
        fprintf('%s\n',char(value(j)));
      end
      %flux dynamics
      value=[obj.flux.keys;obj.flux.values];
      for j=1:numel(value)
        value{j}=regexprep(evalc('disp(value{j})'),'\n','');
      end
      fprintf('\nflux dynamics:\n');
      fprintf('%s=%s\n',value{:});
      %parameters
      value=[obj.param.keys;obj.param.values];
      fprintf('\nparameters:\n');
      fprintf('%s=%d\n',value{:});
      %inputs
      value=obj.inputvars;
      fprintf('\ninput variables\n');
      fprintf('%s\n',value{:});      
      %initial conditions
      value=[obj.x0_map.keys;obj.x0_map.values];
      fprintf('\ninitial condition:\n');
      fprintf('%s=%d\n',value{:});
    end
    % changing the dynamical system
    replace(obj,var,exp);
    apply_conservation(obj,conserv_vars,total_var,elim_var);
    convert_to_process(obj,inputvars);
    take_limit(obj,var,value,generic);
    convert_to_acceleration(obj,sv_add_v,fric_var,normalv_coeff)
    % internal verifications
    [pass,sv,sv_mex,out,out_mex]=verify_mex(obj);
    % Generate (un)feasible parameter sets and tests whether the current
    % parameter set satisfies given constraints
    l=get_feasible(obj,inequalities,range,nmax);
    l=get_feasible_sw(obj,inequalities,range,nmax,steps);
    get_feasibles(obj,inequalities,range,params,fname,steps,varargin);
    get_infeasible(obj,inequalities,param,dir,stepsize,limit,almost);
    L=satisfy_constrs(obj,constrs);
    constrs=gen_constrs(obj,constrs);
    % Simulation of the dynamical system
    [response,fluct,o,o_fluct]=find_dose_response(...
      obj,input_range,cutoff_time,max_runtime,resp_cor);
    [response,fluct,o,o_fluct,ss_found]=find_steadystate(obj,cutoff_time,max_runtime);
    [t,y,u_hat]=simulate_behavior(obj,Tu_hat,offset,amp,omega,type,tStart,Fs,noise);%redebug
    [t,y,full_y]=view_solution(obj,vars_disp,tend,time_series,time_limit,trans_y,view);
    [t,x]=simulate_sys_adam(obj,tcut,max_runtime,stepsize,decayrates,min_vdenom);%no unit test
    [t,y]=simulate_sys(obj,tcut,sim_limit);
    [t,y]=simulate_sys_myode15s_halft(obj,tcut,sim_limit);
    [t,y]=simulate_sys_myode15s_eulerheuns(obj,tcut,sim_limit);
  end

  % <html><h3> General operators </h3></html>
  %
  % * *combine* concatnates a series of BioSystem objects into a single
  % BioSystem object after checking for conflicting parameter values (param),
  % fluxes (flux), system rate equations (sys), state variable initial values,
  % (x0_map), state variable ranges (xrange), and parameter value ranges (prange).
  % Outputs need to be respecified.
  % 
  % <html><h3> Misc helpers </h3></html>
  %
  % * *fullexp* expands symbolic expressions in the first arguments based
  % on expression pairs recursively up to ten times or until relevant
  % expressions are fully expanded
  % * *view_solutions* visualizes solution trajectories of a single output from
  % multiple simulations in one plot and tiles plots of different outputs
  % in one figure. The time axis of the solution trajectories should be
  % able to be coordinated. 
  % * *subsd1* performs substitution of first derivative with the actual function for a cell array
  % of symbolic expressions.
  methods(Static=true)
    C = combine(varargin);
    [cell_sym_out] = subsd1(cell_sym,d1pairs,charout);
    view_solutions(data,output_labels,outputs_plotted,tcut);
    function [newexp] = fullexp(exp,flux)
      % Convert expression of fluxes to full expression without flux variables. 
      % INPUT
      % exp - container map of scalar symbolic expressions
      % flux - container map of symbolic expressions; keys must NOT be composite expressions
      % OUTPUT
      % exp - container map of symbolic expressions, fully substituted. 
      newexp=[containers.Map2();exp];
      for i=1:10
        newexp_dep=compute_dependencies(newexp);
        newexp_dep_v=newexp_dep.values;
        fluxvars_unsub=intersect(vertcat(newexp_dep_v{:}),flux.keys);
        if isempty(fluxvars_unsub)
          break;
        else
          newexp=sub_4_param(newexp,flux,newexp_dep);
        end
      end
    end
  end
  methods(Static=true,Hidden)
    [out] = numf_full_wrapper(numf,type,xrange,size_out,p,x,u)
  end
end
```
