## Supplemental Data 1 for "Principles underlying implementation of *nearly*-homeostatic biological networks": clear_numf.m

```
function clear_numf(obj,prop)
% INPUTS
% obj - BioSystem object
% prop - a character vector specifying the name of the property to clear (remove all files and
%   directories and reset the property); prop must have the same form as s_numeric_form. 

val=obj.(prop);
for i=1:2:numel(val)
  if ~isempty(val{i+1})
    val{i}=[];
    if isfolder(val{i+1})
      dir=val{i+1};
      for j=1:10
        try
          rmdir(dir,'s');
          break;
        catch err
          pause(0.1);
          warning([err.identifier,' has occurred!']);
        end
      end
    else
      delete(val{i+1});
    end
    val{i+1}='';
  end
end
obj.(prop)=val;
end
```
