## Supplemental Data 1 for "Principles underlying implementation of *nearly*-homeostatic biological networks": combine.m

```
function C = combine(varargin)
% Inputs
% vargin - variable number of BioSystem objects to be combined
% Output
% C - combined BioSystem object
loop_num=nargin;
if loop_num < 2
  error('combine:too_few_inputs',...
    'There must be at least two BioSystem objects to be combined!');
end

comb=struct; confl=struct;
properties={'prange','param','x0_map','xrange','sys','flux'};
for p_l = properties
  p=p_l{:};
  comb.(p)=varargin{1}.(p);
  confl.(p)={};
end

for i=2:loop_num
  for p_l=properties
    p=p_l{:};
    confl.(p)=intersect(comb.(p).keys,varargin{i}.(p).keys);
    for conf_name_l=confl.(p)
      conf_name=conf_name_l{:};
      if ~isequal(comb.(p)(conf_name),varargin{i}.(p)(conf_name))
        error('combine:conflict',['Object',i,...
          'has a conflicting property value as previously combined objects!\n']);
      end
    end  
    comb.(p)=[comb.(p);varargin{i}.(p)];
  end
end
C=BioSystem(comb.sys,comb.flux,comb.param,comb.x0_map,comb.prange,comb.xrange);
end
```
