## Supplemental Data 1 for "Principles underlying implementation of *nearly*-homeostatic biological networks": convert_to_acceleration.m

```
function convert_to_acceleration(obj,sv_add_v,fric_var)
% INPUTS
% obj - the BioSystem object
% sv_add_v - cell array of state variables to add velocity state and acceleration vectors to 
% fric_var - character vector; name of friction coefficient; if the system does not 
%  already have one, fric_var will be the new friction coefficient
sv_p=strcat(obj.statevars);
sv_v=strcat(sv_add_v,'_v');
if any(ismember([sv_v,{fric_var}],sv_p)) % additional unit test
  error('BioSystem:convert_to_acceleration:statevars',...
    ['Original state variable concatnated with "_v" ',fric_var,...
    ' cannot be another original state!']);
end
% generating new velocity state for accelerate vector
obj.xrange=[obj.xrange;containers.Map2(sv_v,repmat({[-inf,inf]},size(sv_v)))];
obj.x0_map=[obj.x0_map;containers.Map2(sv_v,repmat({0},size(sv_v)))];

% velocity vectors becoming acceleration vectors
obj.sys=[obj.sys;containers.Map2(sv_add_v,num2cell(str2sym(sv_v))); 
  containers.Map2(sv_v,obj.sys.values(sv_add_v))];

% friction to avoid stable oscillation
for i=1:numel(sv_v)
  obj.sys(sv_v{i})=obj.sys(sv_v{i})-str2sym([fric_var,'*',sv_v{i}]);
end
if ~any(strcmp(fric_var,obj.param.keys))
  obj.prange('f')=[0,inf];
  obj.param('f')=0.1;
end
end
```
