## Supplemental Data 1 for "Principles underlying implementation of *nearly*-homeostatic biological networks": convert_to_process.m

```
function convert_to_process(obj, inputvars)
%%converts current system into process with indicated inputvars
%input: 
%inputvars - cell array of variable names to make into inputs

obj.prange.remove(inputvars(ismember(inputvars,obj.prange.keys)));
obj.param.remove(inputvars(ismember(inputvars,obj.param.keys)));
obj.xrange.remove(inputvars(ismember(inputvars,obj.xrange.keys)));
obj.x0_map.remove(inputvars(ismember(inputvars,obj.x0_map.keys)));
obj.sys.remove(inputvars(ismember(inputvars,obj.sys.keys)));
end
```
