## Supplemental Data 1 for "Principles underlying implementation of *nearly*-homeostatic biological networks": convert_to_xpp.m

```
function convert_to_xpp(obj,fname)
%convert MatLab dynamical system to XPP .ode file
%input:
%fname - string of file name to be outputted.

dx=obj.sys;
flux=obj.flux;
param=obj.param;
init=obj.x0_map;

fid=fopen(fname,'w');
fprintf(fid,'#section 0: numerical simulation parameters\n');
fprintf(fid,'@ bound=1e6, meth=gear\n');

fprintf(fid,'\n#section 1: dynamical system\n');
for k=keys(dx)
    fprintf(fid,'d%s/dt=%s\n',k{:},char(numTostr(dx(k{:}))));
end

fprintf(fid,'\n#section 2: parameters\n');
for k=keys(param)
    fprintf(fid,'param %s=%s\n',k{:},char(numTostr(param(k{:}))));
end

fprintf(fid,'\n#section 3: flux\n');
for k=keys(flux)
    fprintf(fid,'%s=%s\n',k{:},char(numTostr(flux(k{:}))));
end

fprintf(fid,'\n#section 4: initial conditions\n');
for k=keys(init)
    fprintf(fid,'init %s=%s\n',k{:},char(numTostr(init(k{:}))));
end

fprintf(fid,'done');
fclose(fid);
end
```
