## Supplemental Data 1 for "Principles underlying implementation of *nearly*-homeostatic biological networks": copyElement.m

```
function[cpObj]=copyElement(obj)
% create a copy of BioSystem handle object
% The copied object is in unlocked form, such that multiple copies of the object
% do have the same container map as its properties. 
cpObj=copyElement@matlab.mixin.Copyable(obj);
% Ensure that the files of cpObj is not the same as obj
cpObj.s_numeric_form=repmat({@()error('Unitialized function!'),''},1,3); 
cpObj.s_numf_output=repmat({@()error('Unitialized function!'),''},1,3); 
% 
cpObj.locked=false;
p={'sys','flux','param','prange','x0_map','xrange'};
for j=1:length(p)
  if isa(cpObj.(p{j}),'containers.Map')
    orig=cpObj.(p{j});
    if orig.Count>0
      cpObj.(p{j})=containers.Map2(orig.keys,orig.values);
    else
      cpObj.(p{j})=containers.Map2;
    end
  end
end
end
```
