## Supplemental Data 1 for "Principles underlying implementation of *nearly*-homeostatic biological networks": eval_eig.m

```
function eigvalues=eval_eig(obj,x,u)
%evaluates eigenvalue of the numeric form of system at indicated points.
%input:
%x - matrix; each column vector is a state space coordinate
%u - matrix; each column vector is an input
eig_func=obj.eigenvalue;
eigvalues=zeros(size(x));

if isempty(obj.inputvars)
    for j=1:size(x,2)
        eigvalues(:,j)=eig_func(x(:,j));
    end
else
    for j=1:size(x,2)
        eigvalues(:,j)=eig_func(x(:,j),u(:,j));
    end
end
end
```
