## Supplemental Data 1 for "Principles underlying implementation of *nearly*-homeostatic biological networks": find_dose_response.m

```
function [response,fluct,o_response,o_fluct]=find_dose_response(...
  obj,input_range,cutoff_time,max_runtime,resp_cor)
%%finding dose response curve of a BioSystem
%input:
%input_range: different inputs must be in different rows; input coordinates
%   should be consistently increasing for optimal speed.
%cutoff_time:[cutoff_start,cutoff_finish] asymptotic time window
%max_runtime (optional): 1x1 matrix; the maximum amount of time per run.
%resp_cor (optional): a function handle that takes state variables as a column 
% vector and output an identically sized column vector that corrects  
% certain states. By default, the program uses subsequent
% steady states to try to save time, but those steady states at times 
% will cause simulation to fail (e.g. center of a dirac-delta function). 
%output:
%response - different steady states are in different rows as well
%fluct - how much max response fluctuate away from min response value in
% the final time window
%o_response - different outputs are in different rows
%o_fluct - how much max response output fluctuate away from min response output

%checking for errors in input
if size(input_range,2)~=length(obj.inputvars)
  error('different inputs must be in different rows');
elseif ~(all(size(cutoff_time)==[1,2]) || all(size(cutoff_time)==[2,1]))
  error('format of cutoff_time is wrong');
end
if nargin<=4
  if nargin<=3
    max_runtime=20;
  end
  resp_cor=@(x)x;
end
n_output=length(obj.output);
n_statevars=obj.sys.Count;
old_x0=obj.x0;

response=NaN(size(input_range,1),n_statevars);
fluct=NaN(size(input_range,1),n_statevars);
o_response=NaN(size(input_range,1),n_output);
o_fluct=NaN(size(input_range,1),n_output);
for j=1:size(input_range,1)
  uc=input_range(j,:).';
  obj.ut=@(t)uc;
  [response(j,:),fluct(j,:),o_response(j,:),o_fluct(j,:),ss_found]=...
    find_steadystate(obj,cutoff_time,max_runtime);
  if ss_found; obj.x0=resp_cor(response(j,:).'); end
end
obj.x0=old_x0;
end
```
