## Supplemental Data 1 for "Principles underlying implementation of *nearly*-homeostatic biological networks": find_steadystate.m

```
function [response,fluct,o_response,o_fluct,ss_found]=find_steadystate(...
  obj,cutoff_time,max_runtime)
% Finding steady state of the system.
%input:
%cutoff_time:[cutoff_start,cutoff_finish] asymptotic time window
%max_runtime: 1x1 matrix; the maximum amount of time per run.
%output:
%response - row vector of steady states
%fluct - how much max response fluctuate away from min response value in
% the final time window
%o_response - row vector of outputs
%o_fluct - how much max response output fluctuate away from min response output
%ss_found - logical scalar of whether steady state is found
n_output=length(obj.output);
n_statevars=obj.sys.Count;
out_func=obj.numf_output;
ss_found=false;
response=NaN(1,n_statevars);
fluct=response;
o_response=NaN(1,n_output);
o_fluct=o_response;


counter=1;
reach_eowindow_once=false;
reach_eocurwindow=false;
while counter %counter to increase cutoff time frame
  new_cutoff=cutoff_time+cutoff_time(2)*(5^(counter-1)-1);
  try
    [t,y]=simulate_sys(obj,[0,new_cutoff(2)],max_runtime);
    reach_eocurwindow=(max(t)>=new_cutoff(2));
    if ~reach_eocurwindow
      warning('BioSystem:find_steadystate:insufficientsimtime',...
        'Simulation does not have enough time to reach desired time window!');
      if counter==1
        break;
      else
        % the previous iteration should have reached desired time window,
        % so take simulation data from the previous time window.
        prev_cutoff=cutoff_time+cutoff_time(2)*(5^(counter-2)-1);
        sel=t>=prev_cutoff(1); % must have at least one element
        sel_ind=find(sel);
        if any(t(sel_ind)>=prev_cutoff(2)) % ensure that prev_cutoff(2) is reached
          if numel(sel_ind)==1
            sel_ind=[sel_ind-1,sel_ind];
          else
            if numel(sel_ind(t(sel_ind)<=prev_cutoff(2)))<=1
              % do nothing
            else
              sel_ind=sel_ind(t(sel_ind)<=prev_cutoff(2));
              sel_ind=[sel_ind;min(sel_ind(end)+1,numel(t))];
            end
          end
        else
          error('This scenario should not have happend!')
        end
        new_cutoff=prev_cutoff;
      end
    else % take data from current time windowv
      reach_eowindow_once=true;
      sel=t>=new_cutoff(1);
      sel_ind=find(sel);
      if numel(sel_ind)>=2
        % sel_ind have at least one element within desired time window case 1
        % Do nothing
      elseif numel(sel_ind)==1
        % sel_ind have at least one element within desired time window case 2
        sel_ind=[sel_ind-1,sel_ind];
      else 
        error('This scenario should not have happened!');
      end
    end
    t=t(sel_ind,:); y=y(sel_ind,:);
    y=real(y);
  catch err
    msgid=err.identifier;   msgstring=getReport(err);
    fprintf('%s: %s\n',msgid,msgstring);
    break;
  end
  
  o_y=NaN(size(y,1),n_output);
  if isempty(obj.inputvars)
    for k=1:size(y,1)
      o_y(k,:)=out_func(y(k,:).');
    end
  else
    for k=1:size(y,1)
      o_y(k,:)=out_func(y(k,:).',obj.ut(t(k)));
    end
  end
  
  max_y=max(y,[],1);
  min_y=min(y,[],1);
  response=y(end,:);
  fluct=max_y-min_y;
  max_oy=max(o_y,[],1);
  min_oy=min(o_y,[],1);
  o_response=o_y(end,:);
  o_fluct=max_oy-min_oy;
  if ~all(abs((max_y-min_y)./(0.5*(max_y+min_y)))<1e-4 |...
      max_y-min_y<1e-4)
    fprintf(['BioSystem:find_dose_response:',...
      'time window %d-%d overlaps with transient state\n'],...
      new_cutoff(1),new_cutoff(2));
    if counter==obj.ss_iter_limit
      warning('BioSystem:find_steadystate',...
        ['simulation unable to reach steady state',...
        ' for the longest time frame']);
      break;
    elseif ~reach_eocurwindow && reach_eowindow_once 
      break;
    else
      counter=counter+1;
    end
  else
    break;
  end
end
end
```
