## Supplemental Data 1 for "Principles underlying implementation of *nearly*-homeostatic biological networks": gen_constrs.m

```
function[constrs]=gen_constrs(obj,constrs)
%generate a numerical function version of symbolic constraint that can be
%   used to quickly test if the object satisfy given constraints on
%   parameters of the BioSystem object. Specifically, the symbolic
%   expression is converted to a numeric function.
%inputs:
%constrs - cell array of strings or symbolic vector of inequalities
%output:
%constrs - a function that takes in parameter values - as a colum vector in
%   the order of parameter keys - and outputs logical column vector indicating which
%   constraints are satisfied.  

%input checking
if size(constrs,2)~=1;error('Constraints must be in a column array or column cell array!');end; 

constrs=ensure_form(constrs,'symbolic');
symbols=symvar(constrs);
l=ismember(symbols,obj.param.keys);
if all(l)
    constrs=matlabFunction(constrs,'vars',{sym(obj.param.keys).'});
else
    error('gen_constrs:InvalidConstraint', ...
        ['constraint can only contain parameters of the BioSystem!\n',...
        '... non-parameter symbols are: ',char(symbols(~l))]);
end
end
```
