## Supplemental Data 1 for "Principles underlying implementation of *nearly*-homeostatic biological networks": gen_mex.m

```
function gen_mex(obj,prop)
% INPUTS
% obj - BioSystem object
% prop - a character vector specifying the name of the property to generate mex files for; 
%   prop must have the same form as s_numeric_form. The first two elements are assumed to contain
%   the original matlab function file. After mex file generation, the first four elements contain
%   information related to the mex files and and last two elements contain information related to
%   the original matlab function. 

obj.(prop)(5:6)=obj.(prop)(1:2);
func_f=obj.(prop){end};
func_f=regexprep(func_f,'^.*[/\\]([^/\\]+).m$','$1');
numel_iv=numel(obj.inputvars);
if numel_iv>0
  argspecs=['{ zeros(',num2str(obj.param.Count),',1,''double''),',...
    'zeros(',num2str(obj.sys.Count),',1,''double''),',...
    'zeros(',num2str(numel_iv),',1,''double'') }'];
else
  argspecs=['{ zeros(',num2str(obj.param.Count),',1,''double''),',...
    'zeros(',num2str(obj.sys.Count),',1,''double'') }'];
end
mex_f=func2mex(func_f,argspecs);
obj.(prop){1}=eval(['@',mex_f]);
mex_fd=[what('.').path,'/codegen/mex/',func_f];
obj.(prop){4}=mex_fd;
mex_f=[what('.').path,'/',mex_f,'.',mexext];
obj.(prop){2}=mex_f;
end
```
