## Supplemental Data 1 for "Principles underlying implementation of *nearly*-homeostatic biological networks": gen_numf.m

```
function [fhandle] = gen_numf(obj,func,fname)
% Input
% func - symbolic matrix containing state variables, parameters, input
%  variables, and fluxes
% fname - name of file to store the function; must end with .m; the
%  directory of the function file must be the current working directory.
% Output
% fhandle - handle of the numeric function with input params,sv[,input] and
%  output out is a vertical vector. Input is a function with one input t. 

params=obj.param.keys; 
sv=obj.statevars;
input=obj.inputvars;
if ~isempty(intersect(params,sv)) || ~isempty(intersect(params,input)) 
  error('BioSystem:gen_numf',...
    'parameters, state variables, and input variables must not overlap!\n')
elseif any(ismember({'params','sv','input'},[params,sv,input]))
  error('BioSystem:gen_numf',...
    '"param","sv","input" must not be present in the system!\n')
end

%% substituting deriv_pairs into func, flux
func_subbed=obj.subsd1(sym2cell(func),obj.deriv_pairs,true);
if obj.flux.Count>0
  flux=containers.Map2(obj.flux.keys,obj.subsd1(obj.flux.values,obj.deriv_pairs,true));
else
  flux=containers.Map2;
end

%% calculate flux variable dependencies
start_fv=intersect(ensure_form(symvar(func),'character'),obj.flux.keys);
start_fv=start_fv.';
ordered_fluxv=order_fluxvars(obj,start_fv);

%% write numeric function
f=fopen(fname,'w');
si=regexp(fname,'[/\\][^/\\]+.m$');
func_name=fname(si+1:(end-2));
% function heading
fprintf(f,'function out = %s (params,sv',func_name);
if ~isempty(input)
  fprintf(f,',input');
end
fprintf(f,')\n');
% params, sv, input extraction
cs=[params;num2cell(1:numel(params))];
fprintf(f,'%s = params(%d,:);\n',cs{:});
cs=[sv;num2cell(1:numel(sv))];
fprintf(f,'%s = sv(%d,:);\n',cs{:});
if ~isempty(input)
  cs=[input;num2cell(1:numel(input))];
  fprintf(f,'%s = input(%d,:);\n',cs{:});
end
% flux expressions
for i=1:numel(ordered_fluxv)
  fv=ordered_fluxv{i};
  v_fv=flux(fv);
  v_fv=regexprep(v_fv,{'<strong>','</strong>',';'},{'','',';\n'});
  % tic; for i=1:1e9 x=1+1; end; toc; utilized the same amount of time as 
  % tic; for i=1:1e9 x=[1+1,;]; end; toc; on Matlab 2022
  fprintf(f,'%s=%s;\n',fv,v_fv);
end
% writing func
fprintf(f,'out = [\n');
func_subbed_t=func_subbed.';
fprintf(f,[repmat('%s,',1,size(func_subbed_t,1)),';\n'],func_subbed_t{:});
fprintf(f,'];\n');
% function ending
fprintf(f,'end\n');
fclose(f);

%% return output
fhandle=str2func(func_name);
end
```
