## Supplemental Data 1 for "Principles underlying implementation of *nearly*-homeostatic biological networks": get_feasible.m

```
function l=get_feasible(obj,inequalities,range,nmax)
%Generates a most likely NEW set of randomized parameters satisfying a set of inequalities.
%If a feasible set of parameters cannot be found, a false value will be
%returned. 
%INPUTS
%inequalities - cell array of strings specifying parameter inequalities or
%   vector of symbolic inequalities; or a numeric function that takes
%   parameter values as a column vector and returns a column logical
%   vector. 
%range - range of paramter sampling; container map: {<param_name|parameter index>,
% [<start>, <end>]}; keys must be a subset of property params; OR cell
% array of free parameters names/indices to change
%nmax - OPTIONAL; maximum number of random parameter sets to try
%OUTPUTS
%l - true or false for ability to find a feasible set of parameters.
if nargin==3; nmax=Inf; end;
if ~isa(inequalities,'function_handle')
    inequalities=obj.gen_constrs(inequalities);
end
if isa(range,'cell')
  range=containers.Map(range,obj.prange.values(range));
end
if strcmp(range.KeyType,'char')
  range_keys=range.keys;
  [~,range_index]=ismember(range_keys,obj.param.keys);
else
  range_index=cell2mat(range.keys);
  range_keys=obj.param.keys; range_keys=range_keys(range_index);
end
pvalues=cell2mat(obj.param.values);

%actual work
range=prange_convert(range);
i=1;
while true
  if i>nmax
    l=false;
    break;
  else
    param_values=rand_param(range,1);pvalues(range_index)=param_values;
    satis=inequalities(pvalues.');
    if(all(satis))
      l=true;
      obj.param=[obj.param; containers.Map2(range_keys,param_values)];
      break;
    end
  end
  i=i+1;
end
end
```
