## Supplemental Data 1 for "Principles underlying implementation of *nearly*-homeostatic biological networks": get_feasible_sw.m

```
function [ l ] = get_feasible_sw( obj,inequalities,range,steps )
%Stepwise generate a randomized set of parameters satisfying a set of inequalities.
%If a feasible set of parameters cannot be found, a false value will be
%returned. 
%INPUTS
%inequalities - cell array of strings specifying parameter inequalities or
%   vector of symbolic inequalities; or a numeric function that takes
%   parameter values as a column vector and returns a column logical
%   vector. 
%range - range of paramter sampling; container map: {<parameter index>,
% [<start>, <end>]}; keys must contain all parameters specified in steps
%steps - n x 3 cell array; Each entry in first column is a
% vector of indices for constraints; each entry in second column is a cell array of
% parameter indices; each entry in third column is the maximum number of attempts for that
% iteration. 
%OUTPUTS
%l - true or false for ability to find a feasible set of parameters.

%input checking
if ~isa(inequalities,'function_handle')
    inequalities=obj.gen_constrs(inequalities);
end
%actual work
maxstep=size(steps,1); 
counter=1;
while true
  for i=1:maxstep
    idx=steps{i,1}; prange=containers.Map(steps{i,2},range.values(steps{i,2}));
    l=get_feasible(obj,@(p)select(inequalities(p),idx),prange,steps{i,3});
    if ~l; break; end;
  end
  if i==maxstep && l
    if inequalities(cell2mat(obj.param.values).')
      break;
    else
      error('BioSystem:get_feasible_sw',...
        'All inequalities are supposed to be satisfied after the last step!');
    end
  else
    fprintf('Pid=%d: search failed at step=%d in cycle=%d.\n',...
      feature('getpid'),i,counter);
  end
  counter=counter+1;
end
end

function out=select(in,idx)
%INPUT
%idx - linear index. 
out=in(idx);
end
```
