## Supplemental Data 1 for "Principles underlying implementation of *nearly*-homeostatic biological networks": get_feasibles.m

```
function get_feasibles (obj,constrs,range,params,fname,steps,varargin)
% INPUT
% A - BioSystems process; parameters cannot contain 'ID'
% constrs - numeric function that takes a column vector of system
%  parameters and return a logical vector OR a symbolic matrix of inequalities
%  (slow) OR a cell array of strings (slowest)
% range - map container of parameter ranges; same format as input to
%  get_feasible; must contain free parameters.
% params - table of parameters whose coordinates are to be fixed: each row is 
% a separate set of parameters. OR if no coordinates are to be fixed, a number 
% denoting number of parameter sets to generate. Column names are parameter names. 
% fname - name of file to write parameters to
% steps - OPTIONAL; steps to take to find each feasible parameter set. 
%   n x 3 cell array; Each entry in first column is a
%   vector of indices for constraints; each entry in second column is a cell array of
%   parameter indices; each entry in third column is the maximum number of attempts for that
%   iteration. If steps is an empty array, normal parameter search algorithm will be used. 
% varargin - additional information needed by get.feasible(_sw) in subclasses. 
% OUTPUT
% file of feasible parameters with columns {obj.param.keys,'ID'}

% input checking
allparams=obj.param.keys;
if ismember('ID',allparams) 
  error('BioSystem:get_feasibles','Parameters cannot contain the "ID" column');
elseif exist(fname,'file') 
  error('BioSystem:enrich',[fname,' already exists!']);
end
if nargin==5; steps={}; end;
if ~isa(constrs,'function_handle')
    constrs=obj.gen_constrs(constrs);
end
if strcmp(range.KeyType,'char')
  [~,p_idx]=ismember(range.keys,allparams);
  range=containers.Map(p_idx,range.values);
end
%initial setup of fixed vs free parameters
if isa(params,'table')
  np=height(params);
  [~,fixed_params]=ismember(params.Properties.VariableNames,allparams);
  fixed_params=fixed_params(fixed_params>0);
else
  np=params;
  fixed_params=[];
end
free_params=setdiff(1:length(allparams),fixed_params); 
if ~isempty(steps) && any(ismember(fixed_params,cell2mat(steps{:,2})))
  error('BioSystem:get_feasibles','A fixed parameter is part of the stepwise algorithm!');
elseif isempty(free_params)
  error('BioSystem:get_feasibles','There has to be at least one free parameter!');
end
fixed_params=num2cell(fixed_params); free_params=num2cell(free_params);
%generate sets of feasible partial parameters. 
fp=fopen(fname,'w'); fprintf(fp,[strjoin([allparams,{'ID'}],','),'\n']); fclose(fp);
for j=1:np
  for k=cell2mat(fixed_params)
    obj.param(allparams{k})=params{j,allparams{k}};
  end
  prange=containers.Map(free_params,range.values(free_params));
  if isempty(steps)
    get_feasible(obj,constrs,prange,Inf,varargin{:});
  else
    get_feasible_sw(obj,constrs,prange,steps,varargin{:});
  end
  dlmwrite(fname,[cell2mat(obj.param.values),j],'-append','delimiter',',');
end
end
```
