## Supplemental Data 1 for "Principles underlying implementation of *nearly*-homeostatic biological networks": get_infeasible.m

```
function get_infeasible(obj,inequalities,param,dir,stepsize,limit,almost)
%Generates an infeasible set of parameters based on a feasible set
%inputs:
%inequalities - cell array of strings specifying parameter inequalities or
%   vector of symbolic inequalities; or a numeric function that takes
%   parameter values as a column vector and returns a column logical
%   vector. 
%param - name of parameter to change
%dir - up | down; direction of parameter change
%stepsize - size of parameter change. 
%limit - a single number denoting limit of parameter change. If limit is
%   reached without breaking feasibility, a warning is given. 
%almost - OPTIONAL. true or false. If true, then the parameter just before
%   breaking the constraints will be the output. 

%input checking
if ~isa(inequalities,'function_handle')
    inequalities=obj.gen_constrs(inequalities);
end
if ~all(inequalities(cell2mat(obj.param.values.'))); 
    warning('BioSystem:get_infeasible','Parameter is already infeasible!');
    return;
end

%get infeasible
while all(inequalities(cell2mat(obj.param.values.')))
    if strcmp(dir,'up')
        if obj.param(param)<=limit
            obj.param(param)=obj.param(param)+stepsize;
        else
            warning('BioSystem:get_infeasible','Parameter limit has been reached!');
            return;
        end
    elseif strcmp(dir,'down')
        if obj.param(param)>=limit
            obj.param(param)=obj.param(param)-stepsize;
        else
            warning('BioSystem:get_infeasible','Parameter limit has been reached!');
            return;
        end
    else
        error('get_infeasible: direction of parameter change must be either up or down!')
    end
end

%become feasible again
if nargin==7 && almost==true
    if strcmp(dir,'up')
        obj.param(param)=obj.param(param)-stepsize;
    elseif strcmp(dir,'down')
        obj.param(param)=obj.param(param)+stepsize;
    end
end
end
```
