## Supplemental Data 1 for "Principles underlying implementation of *nearly*-homeostatic biological networks": numf_full_wrapper.m

```
function [out] = numf_full_wrapper(numf,type,xrange,xband,p,x,u)
% Obtain modified rate of change / output:
% If x is out of xrange, set x to xrange boundary where xrange is
% exceeded to prevent very slow complex number formation (e.g. (-1e5)^0.5) and 
% compute the rate of change, and then set rates to prevent solution trajectory 
% from further exceeding range and to return solution component toward range. 
% A out of bound solution component can be drawn back to within the bound under 
% suitable conditions for the original dynamical system. 
% - Only results in ~2% slow down in BioSystemTests.simulate_sys_simpnumjac with ode23, 
% compared to no wrapper. 
% INPUTS
% numf - full numeric form that output rates
% type - type of numf, 'rates'|'other'. If 'rate', then rates are corrected
% to prevent solution from become further exceeding xrange. If 'other',
% then the output is not corrected but only x is. 
% xrange - N by 2 matrix
% xband - N by 2 matrix specifying the band width for which applicable
%  rates is zero
% p,x,u - arguments to numf, u is optional

l_lowrexcd=x<xrange(:,1);
l_urexcd=x>xrange(:,2);
if any(l_lowrexcd) || any(l_urexcd)
  % set x to xrange boundary where xrange is exceeded
  x_new=x;
  x_new(l_lowrexcd)=xrange(l_lowrexcd,1);
  x_new(l_urexcd)=xrange(l_urexcd,2);
  if nargin<=6 % minimal additional time
    out=numf(p,x_new);
  else
    out=numf(p,x_new,u);
  end

  if strcmp(type,'rates')
    % set xr to prevent solution trajectory from further exceeding range
    l_lowrexcd_n_moreexcd=l_lowrexcd & out<0;
    past_lar=xrange(:,1)-xband(:,1)-x;
    out(l_lowrexcd_n_moreexcd)=extract_farray( max(past_lar,0) , l_lowrexcd_n_moreexcd );
    l_urexcd_n_moreexcd=l_urexcd & out>0;
    past_ur=x-(xrange(:,2)+xband(:,2));
    out(l_urexcd_n_moreexcd)=extract_farray( min(-past_ur,0) , l_urexcd_n_moreexcd );
  end
else
  % just call original numf to noticeably enhance speed
  if nargin<=6 % minimal additional time
    out=numf(p,x);
  else
    out=numf(p,x,u);
  end
end
end
```
