## Supplemental Data 1 for "Principles underlying implementation of *nearly*-homeostatic biological networks": numjac_gen_broyden.m

```
function [outf] = numjac_gen_broyden(obj)
% Generate numeric jacobian function that utilize Broyden's method.
% In phase 1, update prevsimindex Jacobian to lastsimindex if applicable and 
% Jacobian to stored J if applicable. In phase 2, retrieve from stored
% Jacobian. If nan is passed to x, then reset stored_J to Jacobian from
% lastsimindex. If a matrix of x_numel by x_numel+1 is passed to x, then
% update first x_numel column stored_J and end-1 column to stored_x and end
% column to stored_rate
% Output
% outf - a function handle @(p,x,u) that outputs the jacobian matrix
mode='Broy'; % 'Broy' || 'BFGS' || 'fmincon'

x_numel=obj.sys.Count;
thres=repmat(1e-12,x_numel,1);
inputvars=obj.inputvars;
numf_full_wrapped=obj.numeric_form_full_wrapped;
ut=obj.ut;

% variables for Broyden update
lastsol=nan(x_numel+1,1);
prev_simindex=0;
prev_J=nan(x_numel,x_numel); 
prev_x=nan(x_numel,1);
prev_rate=nan(x_numel,1);
stored_J=prev_J; 
stored_x=prev_x;
stored_rate=prev_rate;

% variables for fmincon update
D0=zeros(x_numel,x_numel)+1e-8;
obj_f=@(D,J0)sqrt(sum(D.^2,"all"));
cons_f=@(D,J0,dx,drates)deal([],(J0+D)*dx-drates);
opts.ConstraintTolerance=1e-8;
opts=optimoptions('fmincon');
% opts.Display="off";

outf=@numjac_broyden;

  function [J] = numjac_broyden(p,x,u)
    lastsimindex=obj.lastsimindex;
    cursimindex=lastsimindex+1;
    if lastsimindex==0 % reset Jacobian
      lastsol=nan(x_numel+1,1);
      prev_simindex=0;
      prev_J=nan(x_numel,x_numel);
      prev_x=nan(x_numel,1);
      prev_rate=nan(x_numel,1);
      stored_J=prev_J;
      stored_x=prev_x;
      stored_rate=prev_rate;
    end

    %% update prevsimindex Jacobian to lastsimindex and Jacobian to stored J
    if isnan(x(1))
      % if resetting stored_J to prev_J
      stored_J=prev_J;
      stored_x=prev_x;
      stored_rate=prev_rate;
    elseif all(size(x)==[x_numel,x_numel+2])
      % forced updating of stored_J
      stored_J=x(:,1:x_numel);
      stored_x=x(:,end-1);
      stored_rate=x(:,end);
    % elseif true % always numjac
    elseif lastsimindex<=1 || (cursimindex-prev_simindex)>24 || mod(cursimindex,10000)==0
    % elseif lastsimindex<=1 || (cursimindex-prev_simindex)>24 || mod(cursimindex,inf)==0
      % determine jacobian for last_simindex and for current x for storage using numjac 
      % if simulation just starting or jacobian hasn't been updated for a while
      if isempty(inputvars)
        rate=numf_full_wrapped(p,x);
        J=numjac(@(t,y)numf_full_wrapped(p,y),nan,x,rate,thres,[],false);
      else
        rate=numf_full_wrapped(p,x,u);
        J=numjac(@(t,y)numf_full_wrapped(p,y,ut(t)),nan,x,rate,thres,[],false);
      end
      if true % default
      % if false
        % start with a scaled identity matrix instead
        J=1e-12*eye(x_numel);
        %         %%% drate=alpha*J*dx: made the simulated solution very inaccurate
        %         rate2=numf_full(p,x+thres);
        %         alpha=norm(rate2-rate)/norm(J*thres);
        %         J=alpha*J;
      end
      stored_J=J;
      stored_x=x;
      stored_rate=rate;

      if lastsimindex-prev_simindex>0 % update J for last_simindex
        lastsol=obj.current_sol(:,lastsimindex);
        if any(lastsol(2,:)~=x)
          if isempty(inputvars)
            rate=numf_full_wrapped(p,lastsol(2:end));
            J=numjac(@(t,y)numf_full_wrapped(p,y),nan,lastsol(2:end),rate,thres,[],false);
          else
            rate=numf_full_wrapped(p,lastsol(2:end),u);
            J=numjac(@(t,y)numf_full_wrapped(p,y,ut(t)),nan,lastsol(2:end),rate,thres,[],false);
          end
        end
        % update Jacobian for lastsimindx
        prev_J=J;
        prev_simindex=lastsimindex;
        prev_x=lastsol(2:end);
        prev_rate=rate;
      end
    elseif lastsimindex-prev_simindex>0
      % estimate jacobian of last known completed solution using Broyden
      % method using previous stored Jacobian as a reference. Put into storage 
      % for the last completed solution
      lastsol=obj.current_sol(:,lastsimindex);
      if isempty(inputvars)
        rate=numf_full_wrapped(p,lastsol(2:end));
      else
        rate=numf_full_wrapped(p,lastsol(2:end),ut(lastsol(1)));
      end

      if true % using stored Jacobian as a reference
        % To prevent using a bad stored_J as a reference, need to reset
        % stored_J as soon as bad stored_J is detected. 
        dx=lastsol(2:end)-stored_x;
        drate=rate-stored_rate;
        ref_J=stored_J;
        ref_x=stored_x;
        ref_rate=stored_rate;
      else
        % using prev_simindex Jacobian as a reference
        dx=lastsol(2:end)-prev_x;
        drate=rate-prev_rate;
        ref_J=prev_J;
        ref_x=prev_x;
        ref_rate=prev_rate;
      end

      % update Jacobian for lastsimindx
      prev_simindex=lastsimindex;
      if any(abs(dx)>1e-12)
        prev_J=update_J(ref_J,dx,drate,mode);
        prev_x=lastsol(2:end);
        prev_rate=rate;
      else
        prev_J=ref_J;
        prev_x=ref_x;
        prev_rate=ref_rate;
      end
    end
    if ~any(isnan(x),'all') && size(x,2)==1 && any(abs(x-stored_x)>1e-12)
      % update storage only if someone is just probing for Jacobian based on
      % latest stored Jacobian
      
      if isempty(inputvars)
        rate=numf_full_wrapped(p,x);
      else
        rate=numf_full_wrapped(p,x,u);
      end
      dx=x-stored_x;
      drate=rate-stored_rate;
      J=update_J(stored_J,dx,drate,mode);
      stored_J=J;
      stored_rate=rate;
      stored_x=x;
    end

    %% estimiate Jacobian for the current x based on stored Jacobian
    if isnan(x(1))
      J=nan(x_numel,x_numel);
    else
      J=stored_J;
    end
  end
end
```
