## Supplemental Data 1 for "Principles underlying implementation of *nearly*-homeostatic biological networks": order_fluxvars.m

```
function [fluxvars] = order_fluxvars(obj,start_fv)
% find a order of flux variables such that the earlier variables do not
% contain later variables.  
% Input
% obj - BioSystem object
% start_fv - column cell array of flux variables to determine other flux variables 
% that the currently referenced flux variables depends upon. 
% Output 
% fluxvars - ordered column cell array of flux variables

dependencies = compute_dependencies(obj.flux);
fluxvars = topologicalSort(dependencies,start_fv);
end
```
