## Supplemental Data 1 for "Principles underlying implementation of *nearly*-homeostatic biological networks": replace.m

```
function replace(obj,var,exp)
%Let's define a hiararchy of  symbols: flux(3)>param+inputvars+statevars(1).
% This function replaces a symbol of class flux, param, input, statevars by
% expression containing symbols of equal or lower class. This prevents undefined
% cases like having f1=a*x1 and replacing a with f1. If the symbol to be
% replaced is not part of the new expression and is a parameter or flux or sys, then
% that symbol is removed. All new symbols are automatically categorized to inputvars class
%input:
%var: char vector
%exp: symbolic expression or a string

%% input checking
cls_var=identify_class(obj,var);
expvars=symvar(char(exp)); varinexp=any(strcmp(symvar(exp),var));
cls_exp=zeros(size(expvars));
for j=1:length(expvars)
  cls_exp(j)=identify_class(obj,expvars{j});
end
if cls_var==0
  error('The symbol to be replaced must be a flux variable, parameter, input, or state variable!');
elseif max(cls_exp)>cls_var
  error('replace:ClassIncompatibility', ...
    'A symbol can only be replaced symbols of equal or lower class!');
end

%% basic clean up.
if ~varinexp && obj.param.isKey(var)
  obj.prange.remove(var);
  obj.param.remove(var);
elseif ~varinexp && obj.flux.isKey(var)
  obj.flux.remove(var);
elseif ~varinexp && obj.sys.isKey(var)
  obj.xrange.remove(var);
  obj.x0_map.remove(var);
  obj.sys.remove(var);
end

%% symbol substitution.
obj.flux=sub_4_param(obj.flux,...
  containers.Map(var,ensure_form(exp,'symbolic')));
obj.sys=sub_4_param(obj.sys,...
  containers.Map(var,ensure_form(exp,'symbolic')));
end

function[cls]=identify_class(obj,var)
if obj.flux.isKey(var)
  cls=3;
elseif obj.param.isKey(var) ||  ismember(var,obj.inputvars)
  cls=2;
elseif ismember(var,obj.statevars)
  cls=2;
else
  cls=0;
end
end
```
