## Supplemental Data 1 for "Principles underlying implementation of *nearly*-homeostatic biological networks": satisfy_constrs.m

```
function[L]=satisfy_constrs(obj,constrs)
    %see if parameters of the instance satisfies each constraint
    %inputs:
    %constrs - symbolic matrix of constraints OR cell array of characters 
    %output:
    %L - vector indicating whether each constraint is satisfied. 
    constrs=obj.gen_constrs(constrs);
    L=constrs(cell2mat(obj.param.values).');
end
```
