## Supplemental Data 1 for "Principles underlying implementation of *nearly*-homeostatic biological networks": simulate_behavior.m

```
function [t,y,u_hat]=simulate_behavior(obj,...
    Tu_hat,offset,amp,omega,type,tStart,Fs,noise)
%simulate behaviour of plant given periodic input of a single variable
%Inputs:
%Tu_hat - coordinate transformation from u_hat to u
%offset - offset of periodic u_hat
%amp - amplitude of u
%omega - frequency of signal in Hz; CANNOT be 0. 
%type - generate type of periodic signal (square or cos)
%tStart - start of asymptotic time t;
%Fs - sampling frequency
%noise - amount of noise to be applied to the output
%Outputs:
%t - vector of times; matrix; each row is a time point
%y - vector of corresponding output of the plant;matrix;
% each row is a time point
%u_hat - vector of corresponding input values as different times;matrix;
% each row is a time point

%generate time varying signal
u_hat=gen_signal(offset,amp,omega,type);

obj.ut=@(t)Tu_hat(u_hat(t));
[t,y]=obj.simulate_sys([tStart,tStart+omega^-1*10]);
u_hat=u_hat(t);

%pass simulated data through a discrete and probabilistic filter
[t,y,u_hat]=filter_output(t,y,u_hat,Fs,noise);
end

function[t_sample,y_sample,u_sample]=filter_output(t,y,u,Fs,noise)
%Input: Fs - sampling frequency
%noise - noise level to apply to output value 
%tStart - estimated time that the system reach asymptotic state
t_sample=(0:1/Fs:max(t)).';
y_sample=interp1(t,y,t_sample);
u_sample=interp1(t,u,t_sample);
y_sample=y_sample+(rand(size(y_sample))-0.5).*2.*noise;
end

function[u]=gen_signal(offset,amp,omega,type)
if strcmp(type,'cos')
    u=@(t) offset+amp*cos(2*pi*omega*t);
elseif strcmp(type,'square')
    u=@(t) offset+amp*bump(cos(2*pi*omega*t));
end
end

function [x]=bump(x)
    x(x>=0)=1;
    x(x<0)=-1;
end
```
