## Supplemental Data 1 for "Principles underlying implementation of *nearly*-homeostatic biological networks": simulate_sys.m

```
function[t,y]=simulate_sys(obj,tcut,sim_limit)
%simulate behaviour of plant or closed loop system
%Inputs: 
%tcut - time frame to take the output [<start_time>,<end_time>]
%sim_limit - simulation time limit in seconds. OPTIONAL. sim_limit of 0
%            indicates retrieval of stored solution when sol_locked=true.
%outputs: 
%t - column vector of time; time is translated to start at 0
%y - matrix of simulated trajectory; rows correspond to different times;
%   columns correspond to different state variables

if nargin==2; sim_limit=Inf; end

if ~(all(size(tcut)==[1,2]) || all(size(tcut)==[2,1]))
    error('format of cutoff_time is wrong');
end

%simulation
if isa(obj.s_solution,'struct') || isa(obj.s_solution,'ref1_array')
  s_solution_m=obj.current_sol;
else
  s_solution_m=[];
end
if ~isempty(s_solution_m);s_solution_m=s_solution_m(:,~isnan(s_solution_m(1,:)));end
if ~obj.sol_locked || isempty(s_solution_m) || ...
    (s_solution_m(1,end)<tcut(2) && sim_limit~=0)
  p=cell2mat(obj.param.values.');
  numeric_form_full_wrapped=obj.numeric_form_full_wrapped;
  % numeric_form_full_wrapped=obj.numeric_form_full;
  numjac_full=obj.jacobn_full;
  if isempty(obj.inputvars)
    ode=@(t,x)numeric_form_full_wrapped(p,x);
    jacobn=@(t,x)numjac_full(p,x);
  else
    if isempty(obj.ut)
      error('an input function must be specified for the system');
    end
    ut=obj.ut;
    ode=@(t,x)numeric_form_full_wrapped(p,x,ut(t));
    jacobn=@(t,x)numjac_full(p,x,ut(t));
  end
  
  min_inttol=min(obj.int_tol);
  xrange=cell2mat(obj.xrange.values.');
  xmins=xrange(:,1);
  xmaxs=xrange(:,2);
  for i=1:numel(obj.int_tol) %increase tolerance if simulation fails
    j=obj.int_tol(i);
    j_add=obj.int_tol_absdiff(i);
    tm=timetracker;
    crash_sim=obj.crash_sim;
    allset_f=obj.allset_f;
    lastwarn('');
    obj.s_solution=ref1_array(obj.sys.Count+1,0);
    ode_prefs=odeset('Jacobian',jacobn,'Refine',1,'RelTol',1*10^j,'AbsTol',1*10^(j+j_add),...
      'OutputFcn',@(t,y,flag)stopsim(t,y,flag,obj,tm,sim_limit,...
      xmins-1e16.*~crash_sim,xmaxs+1e16.*~crash_sim,allset_f));
    % latest speed comparison against no OutputFcn done on 2024-07-24 to ensure less than 
    % 10% slow down
    %
    odesolver=obj.odesolver;
    obj.s_tm=tm;
    tm.start;
    [t,y]=odesolver(ode,[0,tcut(2)],obj.x0,ode_prefs);
    tm.pause;
    obj.s_tm=[];
    %
    obj.s_solution.mod(1:(obj.sys.Count+1),1:size(y,1),[t.';y.']);
    s_solution_m=obj.current_sol;
    obj.s_solution.subset(1:(obj.sys.Count+1),~isnan(s_solution_m(1,:)));
    s_solution_m=obj.current_sol;
    t=s_solution_m(1,:).';
    y=s_solution_m(2:end,:).';
    
    warn_msg=lastwarn;
    if isempty(warn_msg)
      break;
    elseif ~isempty(regexp(warn_msg,'tolerances','once')) && j==min_inttol
      error('BioSystem:simulate_sys',...
        ['something''s wrong with your system.\n',...
        'simulation cannot be completed at most strict relative tol!']);
    elseif ~isempty(regexp(warn_msg,'exceeds range','once')) && j==min_inttol
      error('BioSystem:simulate_sys:traj',warn_msg);
    % else % NO LONGER USED BECAUSE RelTol increase warning cause crash
    %  error('BioSystem:simulate_sys:ode15s',warn_msg);
    end
  end
else
  t=s_solution_m(1,:).';
  y=s_solution_m(2:end,:).';
end
sel=t>=tcut(1) & t<=tcut(2);
t=t(sel,:);
y=y(sel,:);
if ~obj.sol_locked
  obj.s_solution=[];
end
end

function status=stopsim(t,y,flag,obj,tm,sim_limit,y_min,y_max,allset_f)
% Input
% y_min,y_min - sv by 1 column vector. 
% allset_f - a function having inputs t,y and output a scalar indicating
%  whether simulation should be stopped.
% As long as ode15s is used, t and y will always be a single column
% vector

if ~strcmp(flag,'done')
  iter_check=obj.iter_check;
  i=tm.inc_iter;
  m_i=mod(i,iter_check);
  % Real time recording of solutions
  if m_i==1
    s_solution_tmp=obj.current_sol;
    s_solution_tmp_ncol=size(s_solution_tmp,2);
    obj.s_solution.mod(':',(1:iter_check)+s_solution_tmp_ncol,NaN(1+size(y,1),iter_check));
  end
  obj.s_solution.mod(':',i,[t(1);y(:,1)]); % turning off only saves 3.85% of time
  
  excdd_xi=find(y(:,1)<y_min | y(:,1)>y_max,1);
  if ~isempty(excdd_xi)
    % checking to see if current solution exceeded limits
    status=1;
    warning(['Simulated trajectory for statvar ',num2str(excdd_xi),' = ',...
      num2str(y(excdd_xi,1)),' exceeds range at time = ',num2str(t(1)),'!']);
  elseif allset_f(t,y)
    status=1;
  elseif m_i==0 && tm.elapsed>=sim_limit
    % Check to see whether time limit has been reached
    status=1;
  else
    status=0;
  end
else
  status=0;
end
end
```
