## Supplemental Data 1 for "Principles underlying implementation of *nearly*-homeostatic biological networks": simulate_sys_myode15s_eulerheuns.m

```
function [t,x] = simulate_sys_myode15s_eulerheuns(obj,cutoff_time,max_runtime)
% Step size changes based on the difference between Euler's solution and
% Heun's solution. 
% INPUTS
% obj - BioSystems object; Assumes that A.ut is constant
% cutoff_time - two element row vector that determine iteration range to
%  determine whether steady state has been reached
% max_runtime - scalar number that specify number of seconds the steady
%  state finding should take.
% OUTPUT
% t - column vector;
% x - matrix where each row is a state vector
stepsize=10^(obj.int_tol(1)+1);
RelTol=10^obj.int_tol(1);
AbsTol=10^(obj.int_tol(1)-3);

%% generated ode form
numf_full=obj.numeric_form_full;
p=cell2mat(obj.param.values.');
iv=obj.inputvars;
x0=obj.x0;
if isempty(iv)
  ode=@(t,x)numf_full(p,x);
else
  u=obj.ut(0);
  ode=@(t,x)numf_full(p,x,u(t));
end

%% adaptive step algorithm
iter_check=obj.iter_check;
t=0;
x=x0;
tm=timetracker;
tm.start;
obj.s_solution=[t;x]; 
while 1
  i=tm.inc_iter; % solver step
  g=ode(t,x);

  while 1
    e_t=t+stepsize;
    e_x=x+stepsize.*g;
    e_g=ode(e_t,e_x);
    h_t=e_t;
    h_x=x+stepsize/2.*(g+e_g);
    diff_abs=abs(e_x-h_x);
    ratio_diffabs=diff_abs./AbsTol;
    ratio_diffrel=diff_abs./abs(h_x)./RelTol;
    ratio_diff=max(min(ratio_diffabs,ratio_diffrel));
    stepsize_recm=1/(1.1*max(ratio_diff,[],'all'));
    stepsize_recm=min(2,stepsize_recm);
    if any(ratio_diff>1)
      stepsize=stepsize*stepsize_recm;
    else
      % current stepsize satisfactory, record step
      t=h_t;
      x=h_x;
      stepsize=stepsize*stepsize_recm;
      break;
    end
  end

  % solution storage
  m_i=mod(i,iter_check);
  if m_i==1
    s_solution_tmp=obj.s_solution;
    s_solution_ncol=size(s_solution_tmp,2);
    s_solution_ids=1:(s_solution_ncol*size(s_solution_tmp,1));
    obj.s_solution=NaN(1+size(x,1),s_solution_ncol+iter_check);
    obj.s_solution(s_solution_ids)=s_solution_tmp;
  end
  obj.s_solution(:,i+1)=[t;x]; 

  % see if steady state finding is complete. 
  if tm.elapsed>max_runtime || t>=cutoff_time(2)
    break;
  end
end
obj.s_solution=obj.s_solution(:,~isnan(obj.s_solution(1,:))); % eliminate nan cols
im=obj.s_solution;
t=im(1,:).';
x=im(2:end,:).';
sel=t>=cutoff_time(1) & t<=cutoff_time(2);
t=t(sel,:);
x=x(sel,:);
end
```
