## Supplemental Data 1 for "Principles underlying implementation of *nearly*-homeostatic biological networks": simulate_sys_myode15s_halft.m

```
function [t,x] = simulate_sys_myode15s_halft(obj,cutoff_time,max_runtime)
% Step size changes based on Euler's method solution difference between current step size
% and half step size. 
% INPUTS
% obj - BioSystems object; Assumes that A.ut is constant
% cutoff_time - two element row vector that determine iteration range to
%  determine whether steady state has been reached
% max_runtime - scalar number that specify number of seconds the steady
%  state finding should take.
% OUTPUT
% t - column vector;
% x - matrix where each row is a state vector
stepsize=10^(obj.int_tol(1)+1);
RelTol=10^obj.int_tol(1);
AbsTol=10^(obj.int_tol(1)-3);
stepsize_multiplier=2;

%% generated ode form
numf_full=obj.numeric_form_full;
p=cell2mat(obj.param.values.');
iv=obj.inputvars;
x0=obj.x0;
if isempty(iv)
  ode=@(t,x)numf_full(p,x);
else
  u=obj.ut(0);
  ode=@(t,x)numf_full(p,x,u(t));
end

%% adaptive step algorithm
iter_check=obj.iter_check;
t=0;
x=x0;
tm=timetracker;
tm.start;
obj.s_solution=[t;x]; 
while 1
  i=tm.inc_iter; % solver step
  g=ode(t,x);

  % attempt to double step size
  ls_stepsize=stepsize_multiplier*stepsize;
  ls_t=t+ls_stepsize;
  ls_x=x+ls_stepsize.*g;
  os_g=g;
  os_t=t;
  os_x=x;
  for j=1:stepsize_multiplier
    os_t=os_t+stepsize;
    os_x=os_x+stepsize.*os_g;
    if j~=stepsize_multiplier
      os_g=ode(os_t,os_x);
    end
  end
  diff_abs=abs(os_x-ls_x);
  if all(diff_abs<AbsTol | diff_abs./abs(os_x)<RelTol)
    % if larger step size still resulted in low solution difference
    stepsize=ls_stepsize;
    t=ls_t;
    x=ls_x;
  else % decrease step size
    while 1;
      os_t=t+stepsize;
      os_x=x+stepsize.*g;
      ss_stepsize=stepsize/stepsize_multiplier;
      ss_g=g;
      ss_t=t;
      ss_x=x;
      for j=1:stepsize_multiplier
        ss_t=ss_t+ss_stepsize;
        ss_x=ss_x+ss_stepsize.*ss_g;
        if j~=stepsize_multiplier
          ss_g=ode(os_t,os_x);
        end
      end
      diff_abs=abs(ss_x-os_x);
      if all(diff_abs<AbsTol | diff_abs./abs(ss_x)<RelTol)
        % if smaller step size still resulted in low solution difference
        t=os_t;
        x=os_x;
        break;
      else
        % if smaller step size resulted in high solution difference
        stepsize=ss_stepsize;
      end
    end
  end

  % solution storage
  m_i=mod(i,iter_check);
  if m_i==1
    s_solution_tmp=obj.s_solution;
    s_solution_ncol=size(s_solution_tmp,2);
    s_solution_ids=1:(s_solution_ncol*size(s_solution_tmp,1));
    obj.s_solution=NaN(1+size(x,1),s_solution_ncol+iter_check);
    obj.s_solution(s_solution_ids)=s_solution_tmp;
  end
  obj.s_solution(:,i+1)=[t;x]; 

  % see if steady state finding is complete. 
  if tm.elapsed>max_runtime || t>=cutoff_time(2)
    break;
  end
end
obj.s_solution=obj.s_solution(:,~isnan(obj.s_solution(1,:))); % eliminate nan cols
im=obj.s_solution;
t=im(1,:).';
x=im(2:end,:).';
sel=t>=cutoff_time(1) & t<=cutoff_time(2);
t=t(sel,:);
x=x(sel,:);
end
```
