## Supplemental Data 1 for "Principles underlying implementation of *nearly*-homeostatic biological networks": subsd1.m

```
function [cell_out] = subsd1(cell_sym,d1pairs,charout)
% Substitution of first derivative with the actual function for all elements in cell_sym.
% Currently, diff(fv_symfunmatrix(y,2*y),y) does not result in expected expression, so elements that
% are symmatrix or symfunmatrix will be kept the same. 
% INPUTS
% cell_sym: cell array of symbolic expressions
% d1pairs: 2 column cell array of character vectors specifying partial derivative of a possibly vector 
%   valued function in the form of {'D([n],<function name>)',<derivative function name>}. No spacing
%   is assumed for entries.
% charout: logical scalar indicating whether the output is a cell array of characters or cell array
%   of symbolic expressions. 
% OUTPUTS
% cell_out: cell array of symbolic expression where custom partial derivatives have been
%   specified.
if nargin<=2
  charout=false;
end

%% convert all elements of cell_sym to character vectors and indicate which elements are symmatrix 
%  or symfunmatrix
cell_char=cell(size(cell_sym));
idx_symmatrix=[];
for i=1:numel(cell_char)
  exp=cell_sym{i};
  if isa(exp,'sym')
    cell_char{i}=char(exp);
  else
    cell_char{i}=evalc('disp(exp)');
  end
  if isa(exp,'symmatrix') || isa(exp,'symfunmatrix')
    idx_symmatrix(end+1)=i;
  end
end
cell_char=regexprep(cell_char,{'\n','\s+'},{'',''});
idx_reg=setdiff(1:numel(cell_sym),idx_symmatrix);

%% convert diff(f(...,xi,...),xi) to D([n],f)(...) and find position of xi by counting number of 
%  "," when the number of unmatched forward bracket is at 2 and before appearance of xi
expr4_diff='diff\((\w+)\((?:(?!\2).)*?(\w+)(?:(?!\2).)*?\)(,)\2\)';
cell_char_rep=regexprep(cell_char,{expr4_diff},{'D([n],$1)'});
for i=idx_reg
  [idx_start,idx_end]=regexp(cell_char{i},expr4_diff);
  for j=1:numel(idx_start)
    cellchari_substr=extract_farray(cell_char{i},idx_start(j):idx_end(j));
    [tokens,idx_tokens]=regexp(cellchari_substr,expr4_diff,'tokens','tokenExtents');
    tokens=tokens{:}; % to cell array
    idx_tokens=idx_tokens{:}; % to array
    [name_func,name_var,~]=deal(tokens{:});
    args=extract_farray(cellchari_substr,idx_tokens(1,end)+1:idx_tokens(3,1)-1);
    idx_fbr=regexp(cellchari_substr,'\(');
    idx_bbr=regexp(cellchari_substr,'\)');
    idx_comma=regexp(cellchari_substr,',');
    count_idx=zeros(1,numel(cellchari_substr));
    count_idx(idx_fbr)=1;
    count_idx(idx_bbr)=-1;
    sum_idx=cumsum(count_idx);
    idx_var=idx_tokens(2);
    n=sum(sum_idx(idx_comma(idx_comma<=idx_var))==2) + 1;
    cell_char_rep{i}=regexprep(cell_char_rep{i},{['D\(\[n\],',name_func,'\)']},...
      {['D([',num2str(n),']',',',name_func,')',args]},'once');
  end
end

%% substitute D([n],f) with the true function that compute the indicated partial derivative
if numel(d1pairs)>=2
  cell_char_rep=replace(cell_char_rep,d1pairs(:,1),d1pairs(:,2));
end

if charout
  cell_out=cell_char_rep;
else
  %% convert cell array of character vectors to cell array of symbolic expressions after creating all
  %  symbolic functions with the right number of arguments
  func_info=symvarfun(cell_char_rep);
  for i=1:height(func_info)
    fname=func_info.funcname{i};
    if ~isempty(regexp(fname,'<strong>','once'))
      fname_nob=regexprep(fname,'<strong>|</strong>','');
    else
      fname_nob=fname;
    end
    numargs=func_info.numargs(i);
    str_args=strjoin(strcat('x',arrayfun(@num2str,1:numargs,'UniformOutput',false)),',');
    if exist(fname_nob,'var')
      error('BioSystem:subsd1:variable_conflict',['function ',fname_nob,' already exists as a ',...
        'variable in the function, so please pick some other name!'])
    end
    evalc([fname_nob,'=nan;']);
    if strcmp(fname,fname_nob)
      evalc(['syms ',fname_nob,'(',str_args,')']);
    else
      evalc(['syms ',fname_nob,'(',str_args,') [2 2] matrix']);
    end
  end
  cell_char_rep=regexprep(cell_char_rep,'<strong>|</strong>','');
  cell_out=cell(size(cell_sym));
  for i=idx_reg
    cell_out{i}=str2sym(cell_char_rep{i});
  end
  cell_out(idx_symmatrix)=cell_sym(idx_symmatrix);
end
end
```
