## Supplemental Data 1 for "Principles underlying implementation of *nearly*-homeostatic biological networks": take_limit.m

```
function take_limit(obj,var,value,generic)
%takes limit of systems as a single parameter or flux to some value.
%input:
%var - cell array of variable names
%value - cell array of variable values; values can be numeric, symbolic, or character
%generic - true or false; if true, then every variable starting with the
% specified element is substituted with the corresponding value.
% false by default

%if generic variable format input, then find all matching variables/parameters
if nargin==4 && generic
  init_length=length(var);
  params=[obj.flux.keys,obj.param.keys];
  for k=init_length:-1:1
    selected=params(match(params,['^',var{k}]));
    var=[var([1:k-1,k+1:end]),selected];
    rep_values=num2cell(repmat(value{k},1,length(selected)));%cell
    value=[value([1:k-1,k+1:end]),rep_values];
  end
end

%input checking
inputvars=obj.inputvars;
value_string=cell(1,length(value));
for j=1:length(value)
  if isnumeric(value{j})
    value_string{j}=num2str(value{j});
  else
    value_string{j}=char(value{j});
  end
end
valuevars=symvar(str2sym(value_string));
valuevars=mat2cell(valuevars,1,ones(1,length(valuevars)));
for j=1:length(valuevars)
  valuevars{j}=char(valuevars{j});
end
if any(ismember(valuevars,obj.statevars))
  error(['It does not make sense to take limit of a process as a parameter',...
    ' goes to a state variable!']);
end

%take limit
value=ensure_form(value,'symbolic');
sys=obj.sys; % aliases to speed up code
flux=obj.flux;
for j=1:length(var)
  varj=str2sym(var{j});
  for k=keys(obj.sys)
    sys(k{:})=limit(obj.sys(k{:}),varj,value(j));
  end
  
  for k=keys(obj.flux)
    flux(k{:})=limit(obj.flux(k{:}),varj,value(j));
  end
end

%getting rid of corresponding parameters from param property and flux from flux property
%   and adding new parameter values
trueextras=setdiff(var,char(valuevars));
obj.prange.remove(trueextras(ismember(trueextras,obj.param.keys)));
obj.param.remove(trueextras(ismember(trueextras,obj.param.keys)));
obj.flux.remove(trueextras(ismember(trueextras,obj.flux.keys)));
for j=setdiff(obj.inputvars,inputvars)
  obj.param(j{:})=0;
end
end

function[L]=match(strings,exp)
%tells you whether each cell array element matches given string pattern
%input:
%strings: cell array of inputs OR character vector
%exp: regexp expression
%output:
%L: logical vector indicating whether a match is present for each cell
% array element

%checking for input errors
switch class(strings)
  case 'char'
    strings={strings};
  case 'cell'
    %nothing
  otherwise
    error('unexpected input is detected');
end

status=regexp(strings,exp,'start');
if length(status)<=1
  status={status};
end
L=logical(true(length(status),1));
for j=1:length(status)
  if status{j}
    L(j)=true;
  else
    L(j)=false;
  end
end
end
```
