## Supplemental Data 1 for "Principles underlying implementation of *nearly*-homeostatic biological networks": verify_mex.m

```
function[sv,sv_mex,out,out_mex]=verify_mex(obj)
%% input checking
if ~(obj.locked && obj.num_sim_only && obj.generate_mex)
  error('BioSystem:verify_mex',...
    'The system must set the following to true: locked, num_sim_only, generate_mex!')
end

%% computing function values
p=cell2mat(obj.param.values).';
x=obj.x0;
if numel(obj.inputvars)>0
  u=obj.ut(0);
  sv=obj.s_numeric_form{5}(p,x,u);
  sv_mex=obj.s_numeric_form{1}(p,x,u);
  out=obj.s_numf_output{5}(p,x,u);
  out_mex=obj.s_numf_output{1}(p,x,u);
else
  sv=obj.s_numeric_form{5}(p,x);
  sv_mex=obj.s_numeric_form{1}(p,x);
  out=obj.s_numf_output{5}(p,x);
  out_mex=obj.s_numf_output{1}(p,x);
end
pass=max(min(abs((sv-sv_mex)./sv),abs(sv-sv_mex)))<1e-12 && ...
  max(min(abs((out-out_mex)./out),abs(out-out_mex)))<1e-12;

%% trigger error if pass is false
if ~pass
  error('BioSystem:verify_mex:failed', ...
    'mex function does not produce sufficiently similar results as the original!');
end
end
```
