## Supplemental Data 1 for "Principles underlying implementation of *nearly*-homeostatic biological networks": view_solution.m

```
function[t,y,full_y]=view_solution(obj,tend,vars_disp,time_limit,trans_y,view)
%given closed loop system or a plant with input, graph solution that
% displays only specified state variables
%input:
%tend - end time of simulation
%vars_disp - cell array of variable names (state variable or output label) to display
%time_limit - OPTIONAL. simulation time limit in seconds
%trans_y - OPTIONAL. 4 element cell array of function handles. The first function
%        handle transforms the y values; the second function transforms
%        into the original y values. The third function describes major breaks 
%        to be labeled. The fourth function describes manor ticks that are not
%        labeled.
%view - OPTIONAL. logical scalar that indicates whether plot of solutions
%       all togather should be made. 
%output:
%t - column vector of time; time is translated to start at 0
%y - matrix of output; rows correspond to different times; columns
%    correspond to vars_disp
%full_y - matrix of state variables + output; rows correspond to different
%         times

if nargin<6
  if nargin<5
    if nargin<4
      time_limit=inf;
    end
    trans_y={@(y)y,@(new_y)new_y};
  end
  view=true;
end

all_vars=[obj.statevars(:);obj.output_label(:)];
if ~all(ismember(vars_disp,all_vars)) 
  error('BioSystem:view_solution:vars_disp',...
    'vars_disp must all be state variables or output labels');
end
numf_output=obj.numf_output;
no_inputvars=isempty(obj.inputvars);
[t,y]=simulate_sys(obj,[0,tend],time_limit);
o=cell(size(t,1),1); % much faster than slicing matrix on R2018a
ut=obj.ut;
for i=1:size(y,1)
  if no_inputvars
    o{i}=numf_output(y(i,:).').';
  else
    o{i}=numf_output(y(i,:).',ut(t(i))).';
  end
end
o=vertcat(o{:});
y=[y,o]; full_y=y;
[~,y_cidx]=ismember(vars_disp,all_vars);
y=y(:,y_cidx);

%time series solution
if view
  min_y=min(y(:)); max_y=max(y(:));
  plot(t,trans_y{1}(y));
  legend(vars_disp,'Interpreter','none','Color','none');
  ax=gca;
  if length(trans_y)==4
    ax.YTick=trans_y{1}(trans_y{3}(min_y,max_y));
    ax.YTickLabel=trans_y{2}(ax.YTick);
    ax.YAxis.MinorTick='on';
    ax.YAxis.MinorTickValues=trans_y{1}(trans_y{4}(min_y,max_y));
  end
  ax.XLabel.String='time';
  ax.YLabel.String='value';
end
end
```
