## Supplemental Data 1 for "Principles underlying implementation of *nearly*-homeostatic biological networks": view_solutions.m

```
function view_solutions(data,output_labels,outputs_plotted,tcut)
% INPUTS
% data - N x 2 cell array of matrices. N = number of separate simulations.
%        Each entry in first column is an M x 1 matrix; each entry in the
%        second column is an M x length(output_labels) matrix.
% output_labels - single dimensional cell array of characters
% outputs_plotted - single dimensional cell array of characters
% tcut - time range of simulations to be plotted.

n_outp=length(outputs_plotted);
for c=1:n_outp
  subplot(ceil(sqrt(n_outp)),ceil(sqrt(n_outp)),c);
  hold on;
  for s=1:size(data,1)
    c_sel_idx=find( strcmp(output_labels,outputs_plotted{c}) );
    t_sel=data{s,1}>=tcut(1) & data{s,1}<=tcut(2);
    plot(data{s,1}(t_sel),data{s,2}(t_sel,c_sel_idx(1)));
  end
  legend(string(1:size(data,1)),'Interpreter','none','Color','none');
  ax=gca;
  ax.XLabel.Interpreter='none';
  ax.XLabel.String='time';
  ax.YLabel.Interpreter='none';
  ax.YLabel.String=outputs_plotted{c};
end
end
```
