## Supplemental Data 1 for "Principles underlying implementation of *nearly*-homeostatic biological networks": BioSystem_Homeostasis.m

```
%% Simulation of symbolic biochemical system
% Given a symbolic biochemical system. This program additionally analyzes homeostasis related
% properties of the system and can try to find near-homeostasis supporting parameter sets.
%
%% Specifications
% Specifically, this class can:
%
% # generate inverse homeostasis plots for the system; generate steady
%   state curves of key system components in response to another system
%   component or perturbation.
% # generate augmented system to find near-homeostasis supporting parameter
%   sets using modified gradient descent algorithms described in outline.lyx
%
classdef BioSystem_Homeostasis < BioSystem_Base
  %% Design
  %
  % <html><h3> Dynamic properties </h3></html>
  %
  % * *output_sensitivity* computes the symbolic homeostasis indicator of the
  %   dynamical system
  % * *output_numrelsensitivity* computes the numerical relative
  %   sensitivity of the dynamical system using fast matrix inverses.
  % * *output_sensitivity_nolaplaceexp* computes the sensitivity of the dynamical system
  %   without using Laplace expansion.
  % * *compute_invhomeo* compute inverse homeostasis plots for the current
  %   dynamical system. For each system component, the production rate and degradation
  %   rate curves of that system component is plotted against each system component and
  %   perturbation. All inverse homeostasis plots for each system component
  %   will have the same YAxis scale. The user can specify lower and upper limits
  %   of system component levels to plot. The user can also specify which
  %   production curves to plot and which to not plot. They can also specify scaling
  %   factor for the production and degradation curves.
  % * *compute_sscurves* finds the steady state curve of a system component
  %   in response to another system component according to a selected subset
  %   of rate equations, at different persistent variable levels. A
  %   persistent variable may be a regulatory parameter, a fixed system
  %   component, or perturbation. Inverse homeostasis plots are also drawn
  %   for the last parameter set to show how steady state relationships can
  %   be mapped to shapes of nullcline.
  % * *compute_invhomeo_lvl2* uses *compute_sscurves* to systematically plot how one system
  %   component responds to another system component at different persistent variable levels,
  %   for each unique pair of system components.
  % * *gen_augsys4nearh_* generate augmented systems to find near-homeostasis
  %   supporting parameter sets using modified gradient descent algorithm described in
  %   homeostasis-backbone.lyx
  % * *lm_allset_f_gen* generates a stopping function that takes t,y and
  %   outputs whether the current simulation should be stopped. Stopping
  %   criteria is satisfied if output allset is fulfilled or rsenso1 doesn't change for a
  %   user-defined number of iterations. This generated function is only valid if the
  %   simulation does not change p, ut, numeric_form_full, and numf_output_full, in order to
  %   maintain maximum simulation speed. Stored information is frozen in place such that the
  %   same stop condition can be regenerated again.
  % * *find_homeostasis* finds steady state of dynamical system at
  %   different multiple sets of parameters. For each parameter, whether
  %   generated at random or supplied, *find_dose_response* is called to find
  %   steady steady state that that parameter set; various metrics important to
  %   homeostasis are also computed.
  %
  methods
    function [obj]=BioSystem_Homeostasis(dx,flux,param,x0_map,prange,xrange)
      % constructor method that is the same as BioSystem_Base
      if nargin<=5
        xrange=containers.Map2(dx.keys,repmat({[-Inf,Inf]},1,dx.Count));
        if nargin<=4
          if param.Count>=1
            prange=containers.Map2(param.keys,repmat({[-10,10]},1,param.Count));
          else
            prange=containers.Map2;
          end
        end
      end
      obj@BioSystem_Base(dx,flux,param,x0_map,prange,xrange);
    end
    find_homeostasis(obj,in,param_range,nsamples,cutoff_time,out_file,start_x0,max_runtime);

    [sens,DxH_map,DxA_map,DuA_map]=output_sensitivity(obj,no_sub);
    [rsens,objc]=output_numrelsensitivity(obj,sens_denom_num);
    [sens,DxH,DxA,DuA,derivatives]=output_sensitivity_nolaplaceexp(obj);
    [sens,DxH,DxA,DuA]=output_sensitivity_nolaplaceexp_fullexp(obj);

    [fig]=compute_invhomeo(obj,x_range,u_range,prod_cv_sel,deg_cv_sel,...
      prod_rates,deg_rates,multipliers,plot_lims,skip_verif);
    compute_sscurves(obj,figNC,figIH,sub_sys,resp_v,indp_v,indp_v_range,resp_v_range,...
      cutoff,fixed_p,fixed_x,fixed_u,n_per_color,swap_axis,...
      prod_rates,deg_rates,multipliers,indp_v_frac,plot_invhomeo);
    compute_invhomeo_lvl2(obj,fprf,sub_sys,sub_sys_range,cutoff,fixed_p,fixed_x,fixed_u,n_per_color);
    [out] = lm_allset_f_gen(A_aug,allset_idx_in_o,~,~,rsenso1_idx_in_o,iter_diff,sens_thres)
  end
  
  %%
  % <html><h3> General operators </h3></html>
  %
  % * *findnhp_fmincon* uses the original system A with a single output, augmented system A_aug
  %   that tracks progress of discretization and fmincon to find and verify nhp, as well as
  %   output any troubleshooting graphs. The signature of this static method is the same as *findnhp*
  % * *findnhp* uses the original system A and the gradient descent based
  %   augmented system to find and verify nhp, as well as output any troubleshooting graphs.
  % * *findnhp_fminconngd* combines the fmincon and gradient descent based approaches.
  % * *findnhp_ga* uses the original system and relative sensitivity function to find
  %   near-homeostasis supporting discrete topologies via the genetic algorithm. 
  % * *findnhp_naive* uses the original system and relative sensitivity function to find
  %   near-homeostasis supporting parameter sets via the naive approach. 
  %
  % The different stages of search for near-homeostasis supporting parameters are:
  %
  % <html><ol>
  %     <li>Stage 0: a near-homeostasis supporting continuous topological parameter and parameter set
  %         is found. </li>
  %     <li>Stage 1: Every topological parameter very close to a discrete value is first discretized
  %         in a manner that satisfy additional constraints after discretization.</li>
  %     <li>Stage 2.1: Rest of the topological parameters are discretized to the closet discrete
  %         values.</li>
  %     <li>Stage 2.2a: Undiscretized topological parameters from stage 1 is discretized in manner
  %         that is biased toward the closest discrete topological parameter value. Multiple trials
  %         of this stage can be run. </li>
  %     <li>Stage 2.2b: Some undiscretized topological parameters from stage 1 is discretized in manner
  %         that is biased toward the closest discrete topological parameter value. Multiple trials
  %         of this stage can be run. </li>
  %     <li>Stage 3: Undiscretized topological parameters from stage 2.2b is discretized in a manner
  %         that is biased toward the closest discrete topological parameter value. Multiple trials
  %         of this stage can be run. </li>
  % </ol></html>
  %
  % <html><h3> Misc helpers </h3></html>
  %
  % * *fmincon_genfunc* generates relative sensitivity function and nonlinear constraint
  %   function for findnhp_fmincon based on a subset of sorted poptx. The functions are generated
  %   from the augmented system.
  % * *map_Aaug2A* maps a numeric function of A_aug (p',x',u) to a function that takes in
  %   (p,x,u,p_extra) of A or vice versa. p_extra is the vector of extra parameters of the first
  %   BioSystem object that does not appear in the second BioSystem object.
  % * *classify_findnhp* classifies a table of parameter searches into different
  %   categories and displays relative proportion of weighted searches for each category. 
  %   For each row, each combination of non-weight columns that are all true gives rise to 
  %   its own category.
  methods(Static=true)
    [out_c,summary_cat,summary_stg]=classify_findnhp(summary,weight_var,ax)
    [outfunc,pextra_keys]=map_Aaug2A(Aaug_numf,A_aug,A);
    [f_rs,f_cons]=fmincon_genfunc(A,A_aug,poptx_sub_idxinpoptx,u,mode,tm_min);
  end
end
```
