## Supplemental Data 1 for "Principles underlying implementation of *nearly*-homeostatic biological networks": classify_findnhp.m

```
function [out_c,summary_cat,summary_stg] = classify_findnhp(data,weight_var,ax)
% INPUT
% data - table with three columns being the {weight_var,iter,stage} and other columns having 
%   only ones and zeros; none of the columns can be named 'category'
% weight_var - name of column containing the weights
% ax - an axis to plot relative proportion of different categories
%
% OUTPUT
% out_c - a cell array of categories for each row of summary. For each row, each combination of
%   non-weight columns that are all true gives rise to its own category.
% summary_cat - table with columns {'category','p_weighted','p_noweight'} specifying proportion of 
%   data belonging to each category. 
% summary_stg - table with columns {'stage','num_iters'} that indicate the number of iterations that
%   has reached the indicated stage. 

%% input checking
if ismember({'category'},data.Properties.VariableNames)
  error('BioSystem:classify_findnhp:category_column',...
    'none of the columns can be named "category"!');
elseif ~all(ismember({weight_var,'iter','stage'},data.Properties.VariableNames))
  error('BioSystem:classify_findnhp:need_more_columns',...
    '{weight_var,"iter","stage"} must be three of the columns!');
end

%% computing out_c
cat_cols=setdiff(data.Properties.VariableNames,{weight_var,'iter','stage'},'stable');
for i=1:numel(cat_cols)
  data.(cat_cols{i})(isnan(data.(cat_cols{i})))=0;
  data.(cat_cols{i})=logical(data.(cat_cols{i}));
end
out_c=rowfun(@(varargin)strjoin(cat_cols(cell2mat(varargin)),' + '),data,...
  'InputVariables',cat_cols,'OutputVariableNames','category','OutputFormat','cell');
l_empty=arrayfun(@(x)isempty(x{:}),out_c);
out_c(l_empty)=repmat({' '},sum(l_empty),1);

%% computing summary_stg
summary_stg=rowfun(@(~,~)1,data,'InputVariables',{'iter','stage'},...
  'GroupingVariables',{'iter','stage'});
summary_stg=rowfun(@(iter,~)numel(iter),summary_stg,'InputVariables',{'iter','stage'},...
  'GroupingVariables',{'stage'},'OutputVariableNames',{'num_iters'});
summary_stg=sortrows(summary_stg,'stage','ascend');
summary_stg=summary_stg(:,{'stage','num_iters'});

%% plotting relative proportion of weighted searches for each category, for both number of rows 
%  and with weights
%
% computing proportion and weighted proportion of searches belonging to each category
out_c_tbl=cell2table(out_c,'VariableNames',{'category'});
out_c_tbl.weights=data.(weight_var);
out_c_tbl.noweights=ones(height(out_c_tbl),1);
total_w=sum(out_c_tbl.weights);
total=sum(out_c_tbl.noweights);
summary_cat=rowfun(@(x_w,x)deal(sum(x_w)/total_w,sum(x)/total),out_c_tbl,...
  'InputVariables',{'weights','noweights'},...
  "GroupingVariables",{'category'},...
  "OutputVariableNames",{'p_weighted','p_noweight'});
summary_cat.GroupCount=[];

% plotting stacked bar group
x=categorical({weight_var,'rows'},'Ordinal',true);
y=[summary_cat.p_weighted,summary_cat.p_noweight];
stg_count_str=sprintf('%d,',summary_stg.num_iters);
stg_count_str=['(',stg_count_str(1:end-1),')'];
bar(ax,x,y.','stacked');
text(x,[0.5,0.5],{{num2str(total_w)},{num2str(total),stg_count_str}},...
  'HorizontalAlignment','center','FontSize',8);
legend(summary_cat.category,'Location','northoutside','Interpreter','none');
ax.TickLabelInterpreter='none';
ax.YLabel.String='proportion';
end
```
