## Supplemental Data 1 for "Principles underlying implementation of *nearly*-homeostatic biological networks": compute_invhomeo.m

```
function [fig]=compute_invhomeo(obj,x_range,u_range,prod_cv_sel,deg_cv_sel,...
  prod_rates,deg_rates,multipliers,plot_lims,skip_verif)
% Compute inverse homeostasis plots of the system. The value fields of prod_rates, deg_rates, 
% multipliers, plot_lims either all have to symbolic expressioins or
% numerical functions. 
% 
% Inputs
% x_range - n steady states of the system. Numeric matrix having n columns
% u_range - n perturbations of the system. Numeric matrix having n columns
% prod_cv_sel - a obj.sys.Count by obj.sys.Count + numel(obj.inputvars)
%  logical matrix indicating whether the corresponding production rate
%  curve will be plotted.
% deg_cv_sel - obj.sys.Count by obj.sys.Count + numel(obj.inputvars)
%  logical matrix indicating whether the corresponding degradation rate
%  curve will be plotted.
% prod_rates - production rate of the system. A container map having the fields
%  obj.statevars. Each value of either is a symbolic expression OR a numerical functions that 
%  takes (p,x,u) of obj. There could be more keys than required, so only the relavant keys will be 
%  used. 
% deg_rates - degradation rate of the system; a container map having the fields
%  obj.statevars. Each value of either is a symbolic expression OR a numerical functions that 
%  takes (p,x,u) of obj 
% multipliers - multipliers for both production rate and degradation rate of
%  the system; a container map having fields obj.statevars. Each value of either is a symbolic 
%  expression OR a numerical functions that  takes (p,x,u) of obj 
% plot_lims - upper and lower limits of each system component and perturbation. 
%  A container map haing fields obj.statevars and obj.inputvars. Each value of either is a symbolic 
%  expression OR a numerical functions that  takes (p,x) or (p,u) of obj. For state variable limits, 
%  no perturbation are present and the elementwise maximum of x will be fed into the numeric function. 
%  For input variable limits, no state variables are present and the elementwise maximum of u will be
%  fed into the numeric function.
% skip_verif - logical. Whether or not to skip verification of whether
%  prod_rate - deg_rate = A_vec. This option is ONLY valid when prod_rates
%  and deg_rates are both symbolic expressions. 
% Output
% fig - figure containing the inverse homeostasis plot

clf;
fig=gcf;
set(fig,'defaulttextInterpreter','latex');

% preliminaries
n_ss=size(x_range,2);
pvs=obj.param.keys;
p_curr=cell2mat(obj.param.values.');
svs=obj.statevars;
svs_n=numel(svs);
ivs=obj.inputvars;
ivs_n=numel(ivs);

if isa(prod_rates(svs{1}),'sym')
  % substitution of prod_rates and deg_rates
  prod_rates=BioSystem.fullexp(prod_rates,obj.flux);
  deg_rates=BioSystem.fullexp(deg_rates,obj.flux);
  multipliers=BioSystem.fullexp(multipliers,obj.flux);
  plot_lims=BioSystem.fullexp(plot_lims,obj.flux);

  % verification of prod_rates and deg_rates
  if ~skip_verif
    obj_vecf=obj.vecfun_form;
    tmp1=prod_rates.values(svs);
    tmp2=deg_rates.values(svs);
    obj_diff=[tmp1{:}].'-[tmp2{:}].';
    if ~isequal(simplify(obj_vecf-obj_diff),sym(zeros(obj.sys.Count,1)))
      error('BioSystem:compute_invhomeo:vecf_proddegrates',...
        'Production rates - degradation rates does not equal to vecfun_form!');
    end
  end

  % turn prod_rates, deg_rates, multipliers, plot_lims into numeric forms
  for i=svs
    sv=i{:};
    prod_rates(sv)=matlabFunction(prod_rates(sv),'Vars',...
      {sym(pvs).',sym(svs).',sym(ivs).'});
    deg_rates(sv)=matlabFunction(deg_rates(sv),'Vars',...
      {sym(pvs).',sym(svs).',sym(ivs).'});
    multipliers(sv)=matlabFunction(multipliers(sv),'Vars',...
      {sym(pvs).',sym(svs).',sym(ivs).'});
    plot_lims(sv)=matlabFunction(plot_lims(sv),'Vars',...
      {sym(pvs).',sym(svs).'});
  end
  for i=ivs
    iv=i{:};
    plot_lims(iv)=matlabFunction(plot_lims(iv),'Vars',...
      {sym(pvs).',sym(ivs).'});
  end
end

% plot inverse homeostasis plots
for r=1:svs_n
  multiplier=multipliers(svs{r});
  prod_rate=prod_rates(svs{r});
  m_prod_rate=@(p,x,u)multiplier(p,x,u)*prod_rate(p,x,u);
  deg_rate=deg_rates(svs{r});
  m_deg_rate=@(p,x,u)multiplier(p,x,u)*deg_rate(p,x,u);
  for c=1:(svs_n+ivs_n)
    subplot(svs_n,svs_n+ivs_n,c+(r-1)*(svs_n+ivs_n));
    hold on;
    if c<=svs_n
      % inverse homeostasis plots as a function of different state
      % variables
      plot_xlim=plot_lims(svs{c});
      xlim=plot_xlim(p_curr,max(x_range,[],2));
      for j=1:n_ss
        if prod_cv_sel(r,c); tmp='-'; else; tmp='none'; end
        fplot(@(xc)m_prod_rate(p_curr,vec_replace(x_range(:,j),c,xc),u_range(:,j)),...
          xlim,'Color','b','LineStyle',tmp);
        if deg_cv_sel(r,c); tmp='-'; else; tmp='none'; end
        fplot(@(xc)m_deg_rate(p_curr,vec_replace(x_range(:,j),c,xc),u_range(:,j)),...
          xlim,'Color','r','LineStyle',tmp);
        if prod_cv_sel(r,c) || deg_cv_sel(r,c); tmp='-'; else; tmp='none'; end
        l=line(repmat(x_range(c,j),1,2),...
          [0,m_prod_rate(p_curr,x_range(:,j),u_range(:,j))],...
          'Color','k','LineStyle',tmp);
      end
    else
      % inverse homeostasis plots as a function of different input
      % variables
      plot_xlim=plot_lims(ivs{c-svs_n});
      xlim=plot_xlim(p_curr,max(u_range,[],2));
      for j=1:n_ss
        if prod_cv_sel(r,c); tmp='-'; else; tmp='none'; end
        fplot(@(uc)m_prod_rate( p_curr,x_range(:,j),...
          vec_replace(u_range(:,j),c-svs_n,uc) ),...
          xlim,'Color','b','LineStyle',tmp);
        if deg_cv_sel(r,c); tmp='-'; else; tmp='none'; end
        fplot(@(uc)m_deg_rate( p_curr,x_range(:,j),...
          vec_replace(u_range(:,j),c-svs_n,uc) ),...
          xlim,'Color','r','LineStyle',tmp);
        if prod_cv_sel(r,c) || deg_cv_sel(r,c); tmp='-'; else; tmp='none'; end
        l=line(repmat(u_range(c-svs_n,j),1,2),...
          [0,m_prod_rate(p_curr,x_range(:,j),u_range(:,j))],...
          'Color','k','LineStyle',':','LineStyle',tmp);
      end
    end
    % set up dual axis: one for production rate, one for degradation
    % rate.
    ax1=gca;
    if xlim(1)~=xlim(2)
      ax1.XLim=xlim;
    end
    if ax1.YLim(1)<0 && ax1.YLim(2)>0
      ax1.YLim(1)=max(ax1.YLim(1),-ax1.YLim(2)/10);
    end
    if c<=svs_n && r==svs_n
      ax1.XLabel.String=svs{c};
    elseif c>svs_n && r==svs_n
      ax1.XLabel.String=ivs{c-svs_n};
    end
    if c==1
      ax1.YLabel.String={['rate of ',svs{r}],'production'};
    end
    ax1.YColor='b';
    ax2=axes('Position',ax1.Position,'Color','none',...
      'XAxisLocation','top','XColor','k',...
      'YAxisLocation','right','YColor','r',...
      'XLim',ax1.XLim,'YLim',ax1.YLim);
    if c==svs_n+ivs_n
      ax2.YLabel.String={['rate of ',svs{r}],'degradation'};
    else
      ax2.YTick=[];
    end
    if r>1
      ax2.XTick=[];
    end
  end
end
% make Y-axis be the same across each row; new nested loop is necessary to
% prevent subsequent rows of graphs from reajusting right axis. 
drawnow;
for r=1:svs_n
  ylim=[Inf,-Inf];
  for c=1:(svs_n+ivs_n)
    if prod_cv_sel(r,c)
      ylim(1)=min(ylim(1),fig.Children(end-2*(c-1+(r-1)*(svs_n+ivs_n))).YLim(1));
      ylim(2)=max(ylim(2),fig.Children(end-2*(c-1+(r-1)*(svs_n+ivs_n))).YLim(2));
    end
    if deg_cv_sel(r,c)
      ylim(1)=min(ylim(1),fig.Children(end-1-2*(c-1+(r-1)*(svs_n+ivs_n))).YLim(1));
      ylim(2)=max(ylim(2),fig.Children(end-1-2*(c-1+(r-1)*(svs_n+ivs_n))).YLim(2));
    end
  end
  for c=1:(svs_n+ivs_n)
    fig.Children(end-2*(c-1+(r-1)*(svs_n+ivs_n))).YLim=...
      [ylim(1),1.1*(ylim(2)-ylim(1))+ylim(1)];
    fig.Children(end-1-2*(c-1+(r-1)*(svs_n+ivs_n))).YLim=...
      fig.Children(end-2*(c-1+(r-1)*(svs_n+ivs_n))).YLim;
    fig.Children(end-1-2*(c-1+(r-1)*(svs_n+ivs_n))).Position=...
      fig.Children(end-2*(c-1+(r-1)*(svs_n+ivs_n))).Position;
  end
end
end
```
