## Supplemental Data 1 for "Principles underlying implementation of *nearly*-homeostatic biological networks": compute_invhomeo_lvl2.m

```
function compute_invhomeo_lvl2(obj,fprf,sub_sys,sub_sys_range,...
  cutoff,fixed_p,fixed_x,fixed_u,n_per_color)
% Systematically plot how one system component responds to another system component at 
% different persistent variable levels, for each unique pair of system components. 
% INPUTS
% obj - BioSystem object
% fprf - character vector specify graphical output prefix. 
% sub_sys - a cell array of characters that specify components of the
%   subsystem where a steady state response curve will be generated for each pair of system
%   components in sub_sys_range.keys, while all other state variables in sub_sys_range.keys are fixed 
% sub_sys_range - a map container of two element vector specifying the plot range of each 
%   system component in sub_sys.
% cutoff - time range to determine steady state. 
% fixed_p - fixed parameter sets; matrix; each column is a set of
%   parameters to be fixed
% fixed_x - fixed system component levels; matrix having same number of 
%   columns as fixed_p; each column is a set of system components to be 
%   fixed, with the appropriate row that vary and appropriate row that will
%   be the response variable. The entries in rows of the sub-system will be
%   used as starting points of simulation. 
% fixed_u - fixed perturbation levels; matrix having the same number of
%   columns as fixed_p; each column is a set of perturbations to be fixed
% n_per_color - number of consecutive fixed columns to have the same color

%% setting up necessary data
sv=obj.statevars;
figNC=figure('Position',[0 0 250 250]);
[~,sub_sys_idx]=ismember(sub_sys,sv);
[~,sub_sys_2plot_idx]=ismember(sub_sys_range.keys,sv);
sub_sys_attach_idx=setdiff(sub_sys_idx,sub_sys_2plot_idx);
xsubs_idx=combnk(sub_sys_2plot_idx,2); % N by 2 matrix

for ver=1:size(xsubs_idx,1)
  %% plot steady state response curve for each pair of system components
  sv_4sscurves=sv([sub_sys_attach_idx,xsubs_idx(ver,1)]);
  clf(figNC);
  compute_sscurves(obj,figNC,nan,sv_4sscurves,sv{xsubs_idx(ver,1)},sv{xsubs_idx(ver,2)},...
    sub_sys_range(sv{xsubs_idx(ver,2)}),sub_sys_range(sv{xsubs_idx(ver,1)}),...
    cutoff,fixed_p,fixed_x,fixed_u,n_per_color,false,...
    [],[],[],[],false);
  for i=1:numel(figNC.Children(1).Children)
    if isa(figNC.Children(1).Children(i),'matlab.graphics.chart.primitive.Line')
      figNC.Children(1).Children(i).LineStyle='--';
    end
  end
  sv_4sscurves=sv([sub_sys_attach_idx,xsubs_idx(ver,2)]);
  compute_sscurves(obj,figNC,nan,sv_4sscurves,sv{xsubs_idx(ver,2)},sv{xsubs_idx(ver,1)},...
    sub_sys_range(sv{xsubs_idx(ver,1)}),sub_sys_range(sv{xsubs_idx(ver,2)}),...
    cutoff,fixed_p,fixed_x,fixed_u,n_per_color,true,...
    [],[],[],[],false);
  exportgraphics(figNC,sprintf('%s-ss%d.pdf',fprf,ver));
end
end
```
