## Supplemental Data 1 for "Principles underlying implementation of *nearly*-homeostatic biological networks": compute_sscurves.m

```
function compute_sscurves(obj,figNC,figIH,sub_sys,resp_v,indp_v,indp_v_range,resp_v_range,...
  cutoff,fixed_p,fixed_x,fixed_u,n_per_color,swap_axis,...
  prod_rates,deg_rates,multipliers,indp_v_range_inv,plot_invhomeo)
% Finds the steady state curve of a system component in response to another 
% system component. Plots fixed_x. Also plot the relevant inverse homeostasis plot having
% only resp_v and indp_v as x-axis at the last set of parameters. 
%
% For the inverse homeostasis plots, choose smaller of the intended plot range or response range
% for the dependent variable. 
% 
% INPUTS
% obj - BioSystem object
% fig_NC - figure handle for nullclines
% sub_sys - a cell array of characters that specify components of the
%   subsystem OR a locked BioSystem object whose statevars are components
%   of the subsystem, output is resp_v, and input is indp_v. 
% resp_v - character vector specifying the system component that will
%   respond
% indp_v - character vector specifying the system component that will act as
%   the independent variable. 
% indp_v_range, resp_v_range - range of independent variable and dependent variable to plot. 
%   two element numeric vector
% cutoff - time range to determine steady state. 
% fixed_p - fixed parameter sets; matrix; each column is a set of
%   parameters to be fixed
% fixed_x - fixed system component levels; matrix having same number of 
%   columns as fixed_p; each column is a set of system components to be 
%   fixed, with the appropriate row that vary and appropriate row that will
%   be the response variable. The entries in rows of the sub-system will be
%   used as starting points of simulation. Values here will also be used as initial states. 
% fixed_u - fixed perturbation levels; matrix having the same number of
%   columns as fixed_p; each column is a set of perturbations to be fixed
% n_per_color - number of consecutive fixed columns to have the same color
% swap_axis - if false, then indp_v is x-axis. 
% prod_rates, deg_rates, multipliers - for compute_invhomeo, see relevant
%   documentation. 
% indp_v_range_inv - 2 element numeric vector; range of indp_v to plot vertical
%   dotted lines in the inverse homeostasis plot. 
% plot_invhomeo - OPTIONAL logical scalar that specify whether the level 1 inverse homeostasis 
%   plot will be generated. If false, then prod_rates, deg_rates, multipliers, indp_v_range_inv
%   could all be []. 

if nargin<=18
  plot_invhomeo=true;
end

%% setting up the subsystem such that only indp_v is the input
if ~isa(sub_sys,'BioSystem')
  obj_sub=obj.copy;
  obj_sub.xrange=containers.Map(sub_sys,obj_sub.xrange.values(sub_sys));
  obj_sub.x0_map=containers.Map(sub_sys,obj_sub.x0_map.values(sub_sys));
  obj_sub.sys=containers.Map(sub_sys,obj_sub.sys.values(sub_sys));
  obj_sub.output=sym(resp_v);
  obj_sub_mparam=setdiff([obj.statevars,obj.inputvars],...
    [{indp_v,resp_v},obj_sub.statevars]); % more parameters for sub sys: fixed states and pert
  obj_sub.prange=[obj_sub.prange;
    containers.Map(obj_sub_mparam,repmat({[-inf,inf]},size(obj_sub_mparam)))];
  obj_sub.param=[obj_sub.param;
    containers.Map(obj_sub_mparam,ones(size(obj_sub_mparam)))];
  if strcmp('extra_flux',obj_sub.flux.keys)
    error('BioSystem:compute_sscurves:extra_flux',...
      'extra_flux cannot be a flux in the original system!');
  else
    obj_sub.flux('extra_flux')=sym(indp_v);
  end
  obj_sub.locked=true;
else
  obj_sub=sub_sys;
end
[~,subsys_parind_inorig]=ismember(obj_sub.param.keys,...
  [obj.param.keys,obj.statevars,obj.inputvars]);
[~,subsys_svind_inorig]=ismember(obj_sub.statevars,obj.statevars);
[~,indpv_idx_inorig]=ismember(indp_v,obj.statevars);
[~,respv_idx_inorig]=ismember(resp_v,obj.statevars);
pxukeys_sub=[obj_sub.param.keys,obj_sub.statevars,obj_sub.inputvars];
[~,p_idxinAsubpxu]=ismember(obj.param.keys,pxukeys_sub);
[~,x_idxinAsubpxu]=ismember(obj.statevars,pxukeys_sub);
[~,u_idxinAsubpxu]=ismember(obj.inputvars,pxukeys_sub);

%% simulating the subsystem to find and plot steady state response curves of resp_v vs indp_v
set(0,'CurrentFigure',figNC);
ax=gca;
hold on;
fixed_pxu=[fixed_p;fixed_x;fixed_u];
n_ind_v_nums=1e2;
ind_v_nums=sort([indp_v_range_inv,...
  logspace(log10(indp_v_range(1)),log10(indp_v_range(2)),n_ind_v_nums)]);
ind_v_nums=ind_v_nums(~isnan(ind_v_nums));
for c=1:size(fixed_p,2)
  obj_sub.param=containers.Map(obj_sub.param.keys,...
    num2cell( fixed_pxu(subsys_parind_inorig,c) ));
  obj_sub.x0=fixed_x(subsys_svind_inorig,c);
  % find_dose_response curves
  usub=ind_v_nums.'; % column vector
  [resp,~,o,o_fluct]=find_dose_response(obj_sub,usub,cutoff,2);
  % ss response curves
  l_ss=min_sn(o_fluct,abs(o_fluct./o))<1e-3;
  lin_color=repmat( 0.80 - 0.80/(ceil(size(fixed_p,2)/n_per_color)-1)*(ceil(c/n_per_color)-1) , 1 , 3 );
  if isnan(lin_color(1))
    lin_color=zeros(1,3);
  end
  if swap_axis
    plot(o(l_ss),ind_v_nums(l_ss),'Color',lin_color);
    scatter(fixed_x(respv_idx_inorig,c),fixed_x(indpv_idx_inorig,c),'x','MarkerEdgeColor',lin_color);
  else
    plot(ind_v_nums(l_ss),o(l_ss),'Color',lin_color);
    scatter(fixed_x(indpv_idx_inorig,c),fixed_x(respv_idx_inorig,c),'x','MarkerEdgeColor',lin_color);
  end
end

% putting axes on log scale and labeling the axes
ax.XScale='log';
ax.YScale='log';
if swap_axis
  ax.XLabel.String=resp_v;
  ax.YLabel.String=indp_v;
  ax.XLim=resp_v_range;
  ax.YLim=indp_v_range;
else
  ax.XLabel.String=indp_v;
  ax.YLabel.String=resp_v;
  ax.XLim=indp_v_range;
  ax.YLim=resp_v_range;
end

if plot_invhomeo
  %% compute inverse homeostasis plot for the sub system evaluated at the last
  %  fixed set of parameters
  % - How to plot inverse homeostasis plot such that data is NOT compressed to one tiny region of graph?
  % - - assume independent variable range is the same.
  % - - for dependent variable, choose smaller of the intended plot range or response range.
  obj_sub.param=containers.Map(obj_sub.param.keys,...
    num2cell( fixed_pxu(subsys_parind_inorig,end) ));
  l_invplot=true(obj_sub.sys.Count,obj_sub.sys.Count+1);
  sel_ul_range=[find(usub>=indp_v_range_inv(1),1,'first');
    find(usub>=indp_v_range_inv(2),1,'first')];
  obj_sub_svs=obj_sub.statevars;
  plot_lims=containers.Map2;
  for j=1:numel(obj_sub_svs)
    plot_lims(obj_sub_svs{j})=sym([min(resp(:,j)),max(resp(:,j))]);
  end
  plot_lims(indp_v)=sym(indp_v_range);
  resp_v_resprange=plot_lims(resp_v);
  plot_lims(resp_v)=sym([max([resp_v_range(1),resp_v_resprange(1)]),...
    min([resp_v_range(2),resp_v_resprange(2)])]);
  if ~isa(prod_rates(indp_v),'sym')
    prod_rates=[containers.Map; prod_rates];
    deg_rates=[containers.Map; deg_rates];
    multipliers=[containers.Map; multipliers];
    for k=plot_lims.keys
      key=k{:};
      if ismember(key,obj_sub.statevars)
        prod_rates(key)=sys2subsys_gen(prod_rates(key),p_idxinAsubpxu,x_idxinAsubpxu,u_idxinAsubpxu);
        deg_rates(key)=sys2subsys_gen(deg_rates(key),p_idxinAsubpxu,x_idxinAsubpxu,u_idxinAsubpxu);
        multipliers(key)=sys2subsys_gen(multipliers(key),p_idxinAsubpxu,x_idxinAsubpxu,u_idxinAsubpxu);
        plot_lims(key)=matlabFunction(plot_lims(key),...
          'vars',{sym(obj_sub.param.keys).',sym(obj_sub.statevars).'});
      else
        plot_lims(key)=matlabFunction(plot_lims(key),...
          'vars',{sym(obj_sub.param.keys).',sym(indp_v).'});
      end
    end
  end
  set(0,'CurrentFigure',figIH);
  clf;
  compute_invhomeo(obj_sub,resp(sel_ul_range,:).',usub(sel_ul_range,:).',...
    l_invplot,l_invplot,prod_rates,deg_rates,multipliers,plot_lims,false);
end
end

function outf = sys2subsys_gen(func,p_idxinAsubpxu,x_idxinAsubpxu,u_idxinAsubpxu)
% Convert a function that takes (p,x,u) of the system to a function that
% takes (p_sub,x_sub,u_sub) of the sub system. 

outf=@func_gen;

  function out=func_gen(p_sub,x_sub,u_sub)
    pxu_sub=[p_sub;x_sub;u_sub];
    p=pxu_sub(p_idxinAsubpxu);
    x=pxu_sub(x_idxinAsubpxu);
    u=pxu_sub(u_idxinAsubpxu);
    out=func(p,x,u);
  end
end
```
