## Supplemental Data 1 for "Principles underlying implementation of *nearly*-homeostatic biological networks": find_homeostasis.m

```
function find_homeostasis(obj,in,param_range,nsamples,cutoff_time,out_file,start_x0,max_runtime)
%search for parameter regimes for which homeostasis is obtained;
%   directional derivative (sens) of each output coordinate is evaluated as input changes
%   from in(k,:) to in(k+1,:) for each k=1:(size(in,1)-1). Compute also sensitivity
%   gradients that is meaningful only when one input component is changed between 
%   consecutive inputs. Locks the process before simulations and unlocks after. 
%input:
%in - numeric matrix; each row correspons to an input.
%param_range - container map; {'<param1>',[<from>,<to>][;...]} OR
%   table of parameter values to simulate. If table of parameter values, column names
%   must correspond one to one to parameters.
%nsamples - number of parameter samples to evaluate relative sensitivity
%cutoff_time - [<start>,<end>];
%out_file - string; name of output file; no need to include suffix
%start_x0 - Table of starting values. Each row corresponds a parameter. OPTIONAL.
%max_runtime: maximum run time to find steady state. OPTIONAL. 1s by default. 
%output:
%file of parameter, input and sensitivity. Each row contains a simulation
% for an unique parameter and input combination. Sensitivity is computed for all inputs
% except the last.

%input checking
if isempty(obj.inputvars) || isempty(obj.output)
  error('find_homeostasis:MissingInputOutput',...
    'input and output of the process must be defined!');
elseif size(in,2)~=length(obj.inputvars)
  error('find_homeostasis:IncorrectInputChange',...
    'incorrect input change is specified!');
elseif exist([out_file,'.csv'],'file')
  error('find_homeostasis:FileConflict',[out_file,' already exists!']);
end
if nargin==6
  start_x0=repmat(obj.x0.',nsamples,1);
  max_runtime=1;
elseif nargin==7
  start_x0=start_x0{:,obj.statevars};
  max_runtime=1;
elseif nargin==8
  start_x0=start_x0{:,obj.statevars};
end

%% final preparation for simulation
obj.locked=true;
paramset=gen_params(obj,param_range,nsamples);
out_length=length(obj.output); sv_length=numel(obj.statevars); in_length=numel(obj.inputvars);
in_total=size(in,1); 
dvu=strcat({'dv_'},obj.inputvars); % assumed to be normal
out_sens=obj.output_sensitivity; out_sens=out_sens*sym(dvu).'; % Y x 1 function
out_sens_j=jacobian(out_sens,sym([obj.statevars,obj.inputvars])); % Y x (N + M) function
out_sens_j=matlabFunction(out_sens_j,'vars',...
  {sym(obj.param.keys).',sym(obj.statevars).',sym(obj.inputvars).',sym(dvu).'});
out_sens=matlabFunction(out_sens,'vars',...
  {sym(obj.param.keys).',sym(obj.statevars).',sym(obj.inputvars).',sym(dvu).'});
eig_ss=@(x,u) eig(obj.jacobn(x,u));
det_ss=@(x,u) prod(eig_ss(x,u));
out_sens=@(x,u,dvu) out_sens(cell2mat(obj.param.values).',x,u,dvu)./det_ss(x,u);
out_sens_j=@(x,u,dvu) out_sens_j(cell2mat(obj.param.values).',x,u,dvu);
%%output file preparation
outvars=strcat('o',cellfun(@num2str,num2cell(1:length(obj.output)),'UniformOutput',false));
out_file=[out_file,'.csv']; f=fopen(out_file,'w');
fprintf(f,'%s,%s,%s,%s,%s,%s,',strjoin(obj.param.keys,','),strjoin(obj.inputvars,','),...
  strjoin(obj.statevars,','),strjoin(strcat({'fluct_'},obj.statevars),','),...
  strjoin(outvars,','),strjoin(strcat({'fluct_'},outvars),','));
fprintf(f,'%s',strjoin([strcat('sens_',outvars),strcat('t_sens_',outvars),{'det'},...
  strjoin(strcat({'eig_'},arrayfun(@num2str,1:length(obj.statevars),...
  'UniformOutput',false)),',')],','));
for j=1:numel(outvars)
  fprintf(f,',%s',strjoin(strcat({'psi_'},outvars(j),{'_'},[obj.statevars,obj.inputvars]),','));
end
fprintf(f,'\n');
%simulation
for j=1:size(paramset,1)
  %steady state function simulation
  obj.param=containers.Map(obj.param.keys,paramset(j,:));
  obj.x0=start_x0(j,:);
  [ss,fluct,out,out_fluct]=obj.find_dose_response(in,cutoff_time,max_runtime);
  %compute measure of homeostasis
  %%for each output coordinate, compute directional derivative of that
  %%coordinate with respect to input.
  for k=1:in_total %compute directional derivative of output with respect to input
    %compute various simulated results
    eiga=NaN([1,length(obj.statevars)]);  eiga=eiga+eiga*1i; deta=NaN;
    sens=NaN(1,out_length);  t_sens=sens; t_sens_j=NaN(out_length,sv_length+in_length);
    if all(isfinite(ss(k,:))) && all(all(isfinite(obj.jacobn(ss(k,:).',in(k,:).'))))
      eiga=eig_ss(ss(k,:).',in(k,:).').';
      deta=det_ss(ss(k,:).',in(k,:).');
      if k<in_total
        out_pair=[out(k,:);out(k+1,:)];in_pair=[in(k,:);in(k+1,:)];
        in_pair_d=in_pair(2,:)-in_pair(1,:);   in_pair_n=norm(in_pair_d);
        for r=1:out_length
          %for each output coordinate, compute directional derivative
          sens(r)=(out_pair(2,r)-out_pair(1,r))/in_pair_n;
        end
        t_sens(:)=out_sens(ss(k,:).',in(k,:).',in_pair_d.'/in_pair_n);
        t_sens_j(:)=out_sens_j(ss(k,:).',in(k,:).',in_pair_d.'/in_pair_n);
      end
    end

    %write results to file
    lin=[sprintf('%.20e,',paramset(j,:)),sprintf('%.20e,',in(k,:)),...
      sprintf('%.20e,',ss(k,:)),sprintf('%.20e,',fluct(k,:)),...
      sprintf('%.20e,',out(k,:)),sprintf('%.20e,',out_fluct(k,:)),...
      sprintf('%.20e,',sens),sprintf('%.20e,',t_sens),...
      sprintf('%.20e,',deta),sprintf('%.20e+%.20ei,',[real(eiga);imag(eiga)])];
    for r=1:out_length
      lin=[lin,sprintf('%.20e,',t_sens_j(r,:))];
    end
    lin=regexprep(lin,'\+-','-');
    lin=lin(1:(end-1));fprintf(f,'%s\n',lin);
  end
end
fclose(f);
obj.locked=false;
end

function paramset=gen_params(obj,param_range,nsamples)
%% parameter generation
if isa (param_range,'containers.Map')
  if param_range.length~=obj.param.length || ~all(strcmp(param_range.keys,obj.param.keys))
    error('find_homeostasis:RangeKeys','range keys must be the same as property param!');
  else
    param_range=prange_convert(param_range);
    paramset=rand_param(param_range,nsamples);
  end
elseif isa(param_range,'table')
  if width(param_range)~=obj.param.length || ...
      ~all(ismember(obj.param.keys,param_range.Properties.VariableNames))
    error('find_homeostasis:paramtable',...
      'input parameter table must correspond to listed parameters exactly!');
  elseif height(param_range)==nsamples &&...
      width(param_range)==length(obj.param.keys)
    paramset=table2array(param_range(:,obj.param.keys));
  else
    error('find_homeostasis:paramdimension',...
      'input parameter table has the wrong dimension!');
  end
else
  error('find_homeostasis:paramclass','param_range must be one of two classes');
end
end
```
