## Supplemental Data 1 for "Principles underlying implementation of *nearly*-homeostatic biological networks": fmincon_genfunc.m

```
function [f_rs,f_cons]=fmincon_genfunc(A,A_aug,poptx_sub_idxinpoptx,u,mode,tm_min)
% Generate output relative sensitivity for fmincon
% INPUT
% A - original BioSystem object
% A_aug - augmented BioSystem object. Each state variable of A_aug must compute neg gradient of
%   relative sensitivity squared. The outputs must contain norm(rsenso1), the non-linear constraints
%   stcat('A_',A.statevars) and gradient of the non-linear constraints
%   strcat('Avec',A.statvars,'_',poptx). A_aug.flux('rsenso1_min_multiplier_rsenso1') must be
%   heaviside_fast(norm(rsenso1)-sens_thres*sens_thres_rsmultiplier)
% poptx_sub_idxinpoptx - index for sorted poptx that are currently being optimized;
%   column vector
% u - column vector
% mode - Character vector. If contains "sqrt_sumsqsys constraint", then the constraint function 
%   will output sqrt_sumsqsys rather than the default rate equations themselves. 
% tm_min - timetracker object that keeps track of time and number of iterations in the search process. 
%   If the current iteration does not change for many function calls (100*numel(poptx)), then the 
%   objective function will try to crash fmincon. 
% OUTPUT
% f_rs - output relative sensitivity for fmincon. Takes in poptx_sub and gives two outputs: the
%   first one is relative sensitivity value, the second one is a column vector of gradients
%   with respect to poptx_sub. 
% f_cons - constraint function that takes in poptx_sub and gives four outputs: 
%   dummy, const, dummy, grad. 

numel_Aaugx=numel(A_aug.statevars);
Aaug_numf=A_aug.numeric_form;
Aaug_numfout=A_aug.numf_output;
heaviside_fast=nan;
var1=nan;
norm_num=@norm;
norm=nan;
syms norm(var1) heaviside_fast(var1)
if A_aug.flux('rsenso1_min_multiplier_rsenso1')~=...
    str2sym('heaviside_fast(norm(rsenso1)-sens_thres*sens_thres_rsmultiplier)')
  error('BioSystem_Homeostasis:fmincon_genfunc',...
    'rsenso1_min_multiplier_rsenso1 expression not correct!');
end

% Compare current iteration with previter only when evals_spreviter reach a certain limit. If current
% iteration > previter, update previter. Otherwise, return nan for all future evals to try to crash
% fmincon. 
prevtmiter=0; 
evals_s_prevtmiter=0;
try2crash=false;

% computing information for f_rs_actual
[~,normrsenso1_idx_in_o]=ismember('norm(rsenso1)',A_aug.output_label);
if normrsenso1_idx_in_o==0
  error('BioSystem:fmincon_genfunc','norm(rsenso1) must be present in output of A_aug!');
end
n_poptx=A_aug.sys.Count;
poptx_subf_idxinpoptx=setdiff(1:n_poptx,poptx_sub_idxinpoptx);
poptx0=A_aug.x0;
sens_thres_act=A_aug.param('sens_thres')*A_aug.param('sens_thres_rsmultiplier');

% computing information for f_cons_actual
mode_sumsqsys_l=~isempty(regexp(mode,"sqrt_sumsqsys constraint",'once'));
Asv=A.statevars;
poptx_keys=A_aug.statevars;
[~,cons_idxino]=ismember(strcat('A_',Asv.'),A_aug.output_label);
[poptx_sub_j,sv_i]=ndgrid(poptx_sub_idxinpoptx,1:numel(Asv));
[~,grad_idxino]=ismember(strcat('Avec',Asv(sv_i),'_',poptx_keys(poptx_sub_j)),...
  A_aug.output_label);
if any(cons_idxino==0,"all") || any(grad_idxino==0,'all')
  error('BioSystem:fmincon_genfunc',...
    'strcat("A_",sv) and strcat("Avec",sv,"_",px) must be present in output of A_aug!');
end

% output functions
f_rs=@f_rs_actual;
f_cons=@f_cons_actual;

  function [out,grad] = f_rs_actual (poptx_sub)
    poptx=nan(n_poptx,1);
    poptx(poptx_sub_idxinpoptx)=poptx_sub;
    poptx(poptx_subf_idxinpoptx)=poptx0(poptx_subf_idxinpoptx);
    out=max(extract_farray(Aaug_numfout(poptx,u),normrsenso1_idx_in_o),sens_thres_act,'includenan');
    if nargout>1
      grad=Aaug_numf(poptx,u);
      grad=-extract_farray(grad,poptx_sub_idxinpoptx)/(2*out);
    end

    % determine whether to try to crash fmincon
    evals_s_prevtmiter=evals_s_prevtmiter+1;
    if evals_s_prevtmiter==100*numel_Aaugx
      if tm_min.curr_iter>prevtmiter
        prevtmiter=tm_min.curr_iter;
        evals_s_prevtmiter=0;
      else
        try2crash=true;
      end
    end
    if try2crash
      out=out.*nan;
      if nargout>1
        grad=grad.*nan;
      end
    end
  end

  function [dummy1,out,dummy2,grad] = f_cons_actual (poptx_sub)
    poptx=nan(n_poptx,1);
    poptx(poptx_sub_idxinpoptx)=poptx_sub;
    poptx(poptx_subf_idxinpoptx)=poptx0(poptx_subf_idxinpoptx);
    out_Aaug=Aaug_numfout(poptx,u);
    out=extract_farray(out_Aaug,cons_idxino);
    if nargout>2
      grad=extract_farray(out_Aaug,grad_idxino);
      if mode_sumsqsys_l
        grad=2.*grad*out./(2*norm_num(out));
      end
    end
    if mode_sumsqsys_l
      out=norm_num(out);
    end
    dummy1=[];
    dummy2=[];
  end
end
```
