## Supplemental Data 1 for "Principles underlying implementation of *nearly*-homeostatic biological networks": BioSystem_Homeostasis.html

Simulation of symbolic biochemical system 

### Simulation of symbolic biochemical system

Given a symbolic biochemical system. This program additionally analyzes homeostasis related properties of the system and can try to find near-homeostasis supporting parameter sets.

#### Contents

- Specifications
- Design

#### Specifications

Specifically, this class can:

1. generate inverse homeostasis plots for the system; generate steady state curves of key system components in response to another system component or perturbation.
2. generate augmented system to find near-homeostasis supporting parameter sets using modified gradient descent algorithms described in homeostasis-backbone.lyx

##### General operators

- **findnhp\_fmincon** uses the original system A with a single output, augmented system A\_aug that tracks progress of discretization and fmincon to find and verify nhp, as well as output any troubleshooting graphs. The signature of this static method is the same as **findnhp**
- **findnhp** uses the original system A and the gradient descent based augmented system to find and verify nhp, as well as output any troubleshooting graphs.
- **findnhp\_fminconngd** combines the fmincon and gradient descent based approaches.

The different stages of search for near-homeostasis supporting parameters are:

1. Stage 0: a near-homeostasis supporting continuous topological parameter and parameter set
   is found.
2. Stage 1: Every topological parameter very close to a discrete value is first discretized
   in a manner that satisfy additional constraints after discretization.
3. Stage 2.1: Rest of the topological parameters are discretized to the closet discrete
   values.
4. Stage 2.2a: Undiscretized topological parameters from stage 1 is discretized in manner
   that is biased toward the closest discrete topological parameter value. Multiple trials
   of this stage can be run.
5. Stage 2.2b: Some undiscretized topological parameters from stage 1 is discretized in manner
   that is biased toward the closest discrete topological parameter value. Multiple trials
   of this stage can be run.
6. Stage 3: Undiscretized topological parameters from stage 2.2b is discretized in a manner
   that is biased toward the closest discrete topological parameter value. Multiple trials
   of this stage can be run.

##### Misc helpers

- **fmincon\_genfunc** generates relative sensitivity function and nonlinear constraint function for findnhp\_fmincon based on a subset of sorted poptx. The functions are generated from the augmented system.
- **map\_Aaug2A** maps a numeric function of A\_aug (p',x',u) to a function that takes in (p,x,u,p\_extra) of A or vice versa. p\_extra is the vector of extra parameters of the first BioSystem object that does not appear in the second BioSystem object.
- **classify\_findnhp** classifies a table of parameter searches into different categories and displays relative proportion of weighted searches for each category. For each row, each combination of non-weight columns that are all true gives rise to its own category.

Published with MATLAB® R2022b
