## Supplemental Data 1 for "Principles underlying implementation of *nearly*-homeostatic biological networks": lm_allset_f_gen.m

```
function [out] = lm_allset_f_gen(A_aug,...
  allset_idx_in_o,~,~,rsenso1_idx_in_o,iter_diff,sens_thres)
% Generate allset_f to detect local minimum and stop parameter search. The output
% function assumes that A_aug.param.values are fixed. 
% Inputs:
% output selection indices must be positive integers
% iter_diff - the same criteria has to be met at least iter_diff search
%   iterations apart AND rsenso1 difference is sufficiently small. 
% sens_thres - threshold for relative sensitivity; 
%   norm(rsenso1)<1.05*sens_thres*sens_thres_rsmultiplier for iter_diff will 
%   trigger local minimum. 
% Outputs:
% function @(t,y) that output status=1 or 0 to stop simulation or not.

A_aug_numf_output_wrapped=A_aug.numf_output_full_wrapped;
A_aug_ut=A_aug.ut;
p=cell2mat(A_aug.param.values.');
sqrtssq_idx_in_o=1; % place holder
norm_rate_p_idx_in_o=1; % place holder
sens_thres_rsmultiplier=A_aug.param('sens_thres_rsmultiplier');
optimValues=struct('iteration',nan,'fval',nan);
lm_f_fmincon=lm_f_fmincon_gen(iter_diff,sens_thres*sens_thres_rsmultiplier);

% output function
out=@allset_f;

  function status = allset_f(t,y)
    x=extract_farray(A_aug_numf_output_wrapped(p,y,A_aug_ut(t)),...
      [allset_idx_in_o,sqrtssq_idx_in_o,norm_rate_p_idx_in_o,rsenso1_idx_in_o]);
    optimValues.iteration=A_aug.lastsimindex;
    optimValues.fval=x(end);
    
    if x(1)
      status=true; 
    else
      status=false;
    end
    status=status || lm_f_fmincon([],optimValues,'iter');
  end
end
```
