## Supplemental Data 1 for "Principles underlying implementation of *nearly*-homeostatic biological networks": map_Aaug2A.m

```
function [outfunc,pextra_keys] = map_Aaug2A(Aaug_numf,A_aug,A)
% maps a numeric function of A_aug (p',x',u) to a function that takes in 
% (p,x,u,pextra) of A.
% INPUT:
% Aaug_numf - numeric function of (p',x',u)
% A_aug - Augmented BioSystem of A
% A - Original BioSystem
% OUTPUT:
% outfunc - numeric function of (p,x,u,pextra); same output as Aaug_numf
% pextra_keys - names of parameters in pextra; cell array of characters. 

% computing location of p,x in p',x' and pextra
for px={'param','sys'}
  px_a=px{:};
  [px_idxinAaugp_l.(px_a),px_idxinAaugp.(px_a)]=ismember(A.(px_a).keys,A_aug.param.keys);
  px_idxinAaugp_no0.(px_a)=px_idxinAaugp.(px_a)(px_idxinAaugp_l.(px_a));
  [px_idxinAaugx_l.(px_a),px_idxinAaugx.(px_a)]=ismember(A.(px_a).keys,A_aug.sys.keys);
  px_idxinAaugx_no0.(px_a)=px_idxinAaugx.(px_a)(px_idxinAaugx_l.(px_a));
end
pextra_keys=extract_farray(A_aug.param.keys,~ismember(A_aug.param.keys,[A.param.keys,A.sys.keys]));
[pextra_idxinAaugp_l,pextra_idxinAaugp]=ismember(pextra_keys,A_aug.param.keys);
pextra_idxinAaugp_no0=pextra_idxinAaugp(pextra_idxinAaugp_l);

% default lengths of A_aug.param and A_aug.x0
n_pAaug=A_aug.param.Count;
n_xAaug=A_aug.sys.Count;

% output of function
outfunc=@outfunc_act;

  function out = outfunc_act(p,x,u,pextra)
    % initialize pAaug and xAaug
    pAaug=nan(n_pAaug,1);
    xAaug=nan(n_xAaug,1);

    % fill in pAaug and xAaug
    pAaug(px_idxinAaugp_no0.param)=p(px_idxinAaugp_l.param);
    pAaug(px_idxinAaugp_no0.sys)=x(px_idxinAaugp_l.sys);
    pAaug(pextra_idxinAaugp_no0)=pextra(pextra_idxinAaugp_l);
    xAaug(px_idxinAaugx_no0.param)=p(px_idxinAaugx_l.param);
    xAaug(px_idxinAaugx_no0.sys)=x(px_idxinAaugx_l.sys);

    % compute output
    out=Aaug_numf(pAaug,xAaug,u);
  end
end
```
