## Supplemental Data 1 for "Principles underlying implementation of *nearly*-homeostatic biological networks": output_numrelsensitivity.m

```
function [rsens,objc] = output_numrelsensitivity(obj,sens_denom_num)
% Determine relative sensitivity of the system output with respect to perturbation
%  D H(g(u))=-DxH (DxA)^-1 DuA / sens_denom_num
% INPUT:
% sens_denom_num - normalize sensitivity fuction; takes in p,x,u; outputs
% numel(h) x numel(u) matrix. 
% output:
% rsens - takes in p,x,u
% objc - a copy of the BioSystem that details that is in rsens. The calling code must accept this 
%  for rsens to be evaluated. 

%% checking further input requirements
if isempty(obj.output) 
    error('output of the system must be set');
end

%% computing numerical relative sensitivity using automatic differentiation.  
[sens,DxH_a,DxA_a,DuA_a,derivatives]=output_sensitivity_nolaplaceexp(obj);
objc=obj.copy;
new_fluxkeys=[{'senso1','DxH','DxA','DuA'},derivatives.keys];
if any(objc.flux.isKey(new_fluxkeys))
  error('the system''s flux already contains one of %s!',strjoin(new_fluxkeys,', '));
end
objc.flux=[ obj.flux ; containers.Map2(new_fluxkeys,[{sens,DxH_a,DxA_a,DuA_a},derivatives.values]) ];
objc.output_label='senso1';
objc.output=sym('senso1');
objc.locked=true;
numf_output_full=objc.numf_output_full;
rsens=@(p,x,u)numf_output_full(p,x,u)./sens_denom_num(p,x,u);
end
```
