## Supplemental Data 1 for "Principles underlying implementation of *nearly*-homeostatic biological networks": output_sensitivity.m

```
function [sens,DxH_map,DxA_map,DuA_map]= output_sensitivity(obj,no_sub)
% Determine numerator of steady state sensitivity of output with respect to input
%  D H(g(u))=-DxH C^T DuA
% INPUT:
% no_sub - logical; if true, then sens uses keys of DxH_map, DxA_map,
%  DuA_map
% output:
% sens - -DxH C^T DuA; symbolic matrix; consisted only of parameters and
%  system components by default.
% DxH_map - map container of DxH
% DxA_map - map container of DxA
% DuA_map - map container of DuA

if nargin==1
  no_sub=false;
end

%checking further input requirements
if isempty(obj.output) 
    error('output of the system must be set');
end
vecf=obj.vecfun_form; 
sv=obj.statevars;
iv=obj.inputvars;

DxA=jacobian(vecf,sym(sv));
DuA=jacobian(vecf,sym(iv));
out=containers.Map2({'place holder'},{obj.output});
out=BioSystem.fullexp(out,obj.flux);
out=out('place holder');
DxH=jacobian(out,sym(sv));

% sens=-DxH*cof(DxA).'*DuA; consume too much memory
DxH_rep=sym('DxH',size(DxH));
DxA_rep=sym('DxA',size(DxA));
DuA_rep=sym('DuA',size(DuA));
sens=-DxH_rep*cof(DxA_rep).'*DuA_rep;

% substituted output_sensitivity
DxH_map=containers.Map2(arrayfun(@char,DxH_rep,'UniformOutput',false),sym2cell(DxH));
DxA_map=containers.Map2(arrayfun(@char,DxA_rep,'UniformOutput',false),sym2cell(DxA));
DuA_map=containers.Map2(arrayfun(@char,DuA_rep,'UniformOutput',false),sym2cell(DuA));
if ~no_sub
  sens=subs(sens,DxH_rep,DxH);
  sens=subs(sens,DuA_rep,DuA);
  sens=subs(sens,DxA_rep,DxA);
end
end
```
