## Supplemental Data 1 for "Principles underlying implementation of *nearly*-homeostatic biological networks": output_sensitivity_nolaplaceexp.m

```
function [sens,DxH_a,DxA_a,DuA_a,derivatives] = output_sensitivity_nolaplaceexp(obj)
% Determine sensitivity of the system output with respect to perturbation
%  D H(g(u))=-DxH (DxA)^-1 DuA. Assumes obj.output is a column vector.  
% output:
% sens - symbolic expression
% DxH_a - symbolic expression
% DxA_a - symbolic expression
% DuA_a - symbolic expression
% derivatives - map container of variable names and scalar symbolic expressions. 

inv=nan;
syms DxH DxA DuA 2 matrix
syms inv(DxA) 2 matrix keepargs

%% checking further input requirements
if isempty(obj.output) 
    error('output of the system must be set');
end
ratesSym=cell2sym(obj.sys.values).'; 
rates_names=strcat('rates_',obj.statevars.');
outSym=obj.output;
out_n=numel(outSym);
sv=obj.statevars;
svSym=sym(sv);
sv_n=numel(sv);
iv=obj.inputvars;
ivSym=sym(iv);
iv_n=numel(iv);

%% 
[DxA_a,derivatives]=reverseAutoDiff(obj.flux,ratesSym,rates_names,svSym);
DxA_a=reshape(DxA_a,[sv_n,sv_n]);
[DuA_a,tmp]=reverseAutoDiff(obj.flux,ratesSym,rates_names,ivSym);
DuA_a=reshape(DuA_a,[sv_n,iv_n]);
derivatives=[derivatives;tmp];
[DxH_a,tmp]=reverseAutoDiff(obj.flux,outSym,obj.output_label,svSym);
DxH_a=reshape(DxH_a,[out_n,sv_n]);
derivatives=[derivatives;tmp];
sens=-DxH*inv(DxA)*DuA; 
end
```
