## Supplemental Data 1 for "Principles underlying implementation of *nearly*-homeostatic biological networks": output_sensitivity_nolaplaceexp_fullexp.m

```
function [sens,DxH_a,DxA_a,DuA_a] = output_sensitivity_nolaplaceexp(obj)
% Determine sensitivity of the system output with respect to perturbation
%  D H(g(u))=-DxH (DxA)^-1 DuA 
% output:
% sens - symbolic expression
% DxH_a - symbolic expression
% DxA_a - symbolic expression
% DuA_a - symbolic expression

inv=nan;
syms DxH DxA DuA 2 matrix
syms inv(DxA) 2 matrix keepargs
%checking further input requirements
if isempty(obj.output) 
    error('output of the system must be set');
end
vecf=obj.vecfun_form; 
sv=obj.statevars;
iv=obj.inputvars;

DxA_a=jacobian(vecf,sym(sv));
DuA_a=jacobian(vecf,sym(iv));
out=containers.Map2({'place holder'},{obj.output});
out=BioSystem.fullexp(out,obj.flux);
out=out('place holder');
DxH_a=jacobian(out,sym(sv));

sens=-DxH*inv(DxA)*DuA; 
end
```
