## Supplemental Data 1 for "Principles underlying implementation of *nearly*-homeostatic biological networks": BioSystemTests_Base.m

```
%%  Unit testing framework for BioSystem class
% It will ensure that major components of BioSystem class is fully
% functional under implementation changes. A method is considered a major component
% if it is user intended OR it is a major function within a hiararchy.
classdef BioSystemTests_Base < matlab.unittest.TestCase
  properties
    BioSys
    exp_sol
  end
  
  methods(TestMethodSetup)
    function setup(tc) % run for each test
      sys={'x1','f3+b*x2';'x2','c*x1+d*x2'};
      flux={'f1','a*x1';'f2','f1';'f3','f2';'o1','x1'};
      param={'a',-5;'b',1;'c',4;'d',-2};
      x0_map={'x1',1;'x2',2};
      tc.BioSys=BioSystem(cell2map(sys),cell2map(flux),...
        cell2map(param),cell2map(x0_map));
      tc.BioSys.output='o1';
      tc.BioSys.output_label={'o1'};
      tc.exp_sol=@(t)3/5.*[1;4]*exp(-t)-2/5.*[-1;1]*exp(-6.*t);%t must be row vector
    end
  end
  methods (Test)
    function vecfun_form(tc)
      vf=tc.BioSys.vecfun_form;
      vf_exp=str2sym({'a*x1+b*x2';'c*x1+d*x2'});
      tc.verifyEqual(vf,vf_exp);
    end
    function sol_locked(tc)
      %% attempted modifications when sol_locked is true
      tc.BioSys.locked=true;
      tc.BioSys.sol_locked=true;
      verifyTrue(tc,tc.BioSys.sys.locked_a);
      verifyTrue(tc,tc.BioSys.flux.locked_a);
      verifyTrue(tc,tc.BioSys.param.locked_a);  
      verifyTrue(tc,tc.BioSys.x0_map.locked_a); 
      verifyTrue(tc,tc.BioSys.xrange.locked_a);
      verifyTrue(tc,tc.BioSys.prange.locked_a);
      verifyError(tc,@mod_sys,'BioSystem:sys');
      verifyError(tc,@mod_sys2,'Map2:alocked');
      verifyError(tc,@mod_sys3,'Map2:alocked');
      verifyError(tc,@mod_sys4,'Map2:klocked');
      verifyError(tc,@mod_flux,'BioSystem:flux');
      verifyError(tc,@mod_param,'BioSystem:param');
      verifyError(tc,@mod_output,'BioSystem:output');
      verifyError(tc,@mod_output_label,'BioSystem:output_label');
      verifyError(tc,@mod_x0_map,'BioSystem:x0_map');
      verifyError(tc,@mod_xrange,'BioSystem:xrange');
      verifyError(tc,@mod_prange,'BioSystem:prange');
      verifyError(tc,@mod_ut,'BioSystem:ut');
      verifyError(tc,@mod_int_tol,'BioSystem:int_tol');
      verifyError(tc,@mod_num_sim_only,'BioSystem:num_sim_only');
      verifyError(tc,@mod_crash_sim,'BioSystem:crash_sim');
      function mod_sys(); tc.BioSys.sys=containers.Map({'x1','x2'},{1,2}); end
      function mod_sys2(); tc.BioSys.sys('x1')=1; end
      function mod_sys3(); tc.BioSys.sys('x3')=1; end
      function mod_sys4(); tc.BioSys.sys.remove('x1'); end
      function mod_flux(); tc.BioSys.flux=containers.Map({'x1','x2'},{1,2}); end
      function mod_param(); tc.BioSys.param=containers.Map(); end
      function mod_output; tc.BioSys.output=sym('x1'); end
      function mod_output_label; tc.BioSys.output_label={'x1'}; end
      function mod_x0_map; tc.BioSys.x0_map=containers.Map(); end
      function mod_xrange; tc.BioSys.xrange=containers.Map(); end
      function mod_prange; tc.BioSys.prange=containers.Map(); end
      function mod_ut; tc.BioSys.ut=@(t)1; end
      function mod_int_tol; tc.BioSys.int_tol=-6; end
      function mod_num_sim_only; tc.BioSys.num_sim_only=true; end
      function mod_crash_sim; tc.BioSys.crash_sim=true; end

      %% see if changing time frame to a subset extracts the already stored solution. 
      [t1,y1]=tc.BioSys.simulate_sys([0,1000]);
      tc.verifyEqual(t1([1,end]),[0;1000]);
      tc.verifyEqual(y1,tc.exp_sol(t1.').','RelTol',0.002,'AbsTol',0.002);
      tc.BioSys.sol_locked=true;
      [t2,y2]=tc.BioSys.simulate_sys([10,100]);
      [~,idx_t2int1]=ismember(t2,t1);
      tc.verifyTrue(t2(1)>=10 && t2(end)<=100);
      tc.verifyTrue(all(idx_t2int1));
      tc.verifyEqual(y2,y1(idx_t2int1,:));
      % current_sol extraction
      tc.verifyEqual(size(tc.BioSys.current_sol),size([t1,y1].'));
      tc.verifyError(@mod_currentsol,'MATLAB:class:noSetMethod');
      
      %% attempted modifications when sol_locked is false but locked is true
      tc.BioSys.sol_locked=false;
      verifyTrue(tc,tc.BioSys.sys.locked_a);
      verifyTrue(tc,tc.BioSys.flux.locked_a);
      verifyTrue(tc,tc.BioSys.param.locked_k);  verifyTrue(tc,~tc.BioSys.param.locked_a);
      verifyTrue(tc,tc.BioSys.x0_map.locked_k); verifyTrue(tc,~tc.BioSys.x0_map.locked_a);
      verifyTrue(tc,tc.BioSys.xrange.locked_k); verifyFalse(tc,tc.BioSys.xrange.locked_a);
      verifyTrue(tc,tc.BioSys.prange.locked_k); verifyFalse(tc,tc.BioSys.prange.locked_a);
      verifyError(tc,@mod_sys,'BioSystem:sys');
      verifyError(tc,@mod_flux,'BioSystem:flux');
      verifyError(tc,@mod_param2,'Map2:klocked');
      verifyError(tc,@mod_output,'BioSystem:output');
      verifyError(tc,@mod_output_label,'BioSystem:output_label');
      verifyError(tc,@mod_x0_map,'BioSystem:x0_map');
      verifyError(tc,@mod_xrange,'BioSystem:xrange');
      verifyError(tc,@mod_prange,'BioSystem:prange');
      function mod_param2(); tc.BioSys.param.remove('a'); end
      function mod_currentsol(); tc.BioSys.current_sol(1,1)=1; end
    end
    function locked(tc)
      %% verify that modification of system before locking doesn't cause the
      %  wrong function to be locked in place.
      verifyEqual(tc,size(tc.BioSys.numeric_form([1;2])),[2,1]);
      tc.BioSys.convert_to_process({'a'});
      tc.BioSys.locked=true;
      verifyEqual(tc,size(tc.BioSys.numeric_form([1;2],5)),[2,1]);

      %% testing for properties to resist change
      verifyTrue(tc,tc.BioSys.sys.locked_a);
      verifyTrue(tc,tc.BioSys.flux.locked_a);
      verifyTrue(tc,tc.BioSys.param.locked_k);  verifyTrue(tc,~tc.BioSys.param.locked_a);
      verifyTrue(tc,tc.BioSys.x0_map.locked_k); verifyTrue(tc,~tc.BioSys.x0_map.locked_a);
      verifyTrue(tc,tc.BioSys.xrange.locked_k); verifyTrue(tc,~tc.BioSys.xrange.locked_a);
      verifyTrue(tc,tc.BioSys.prange.locked_k); verifyTrue(tc,~tc.BioSys.prange.locked_a);
      verifyError(tc,@mod_sys,'BioSystem:sys');
      verifyError(tc,@mod_sys2,'Map2:alocked');
      verifyError(tc,@mod_sys3,'Map2:alocked');
      verifyError(tc,@mod_sys4,'Map2:klocked');
      verifyError(tc,@mod_flux,'BioSystem:flux');
      verifyError(tc,@mod_param,'BioSystem:param');
      verifyError(tc,@mod_x0_map,'BioSystem:x0_map');
      verifyError(tc,@mod_xrange,'BioSystem:xrange');
      verifyError(tc,@mod_prange,'BioSystem:prange');
      verifyError(tc,@mod_output,'BioSystem:output');
      verifyError(tc,@mod_output_label,'BioSystem:output_label');
      verifyError(tc,@mod_deriv_pairs,'BioSystem:deriv_pairs');
      verifyError(tc,@mod_numsimonly,'BioSystem:num_sim_only');
      verifyError(tc,@mod_generatemex,'BioSystem:generate_mex');
      function mod_sys(); tc.BioSys.sys=containers.Map({'x1','x2'},{1,2}); end
      function mod_sys2(); tc.BioSys.sys('x1')=1; end
      function mod_sys3(); tc.BioSys.sys('x3')=1; end
      function mod_sys4(); tc.BioSys.sys.remove('x1'); end
      function mod_flux(); tc.BioSys.flux=containers.Map({'x1','x2'},{1,2}); end
      function mod_param()
        tc.BioSys.param=containers.Map({'b','c','e'},tc.BioSys.param.values);
      end
      function mod_output; tc.BioSys.output=sym('x1'); end
      function mod_output_label; tc.BioSys.output_label={'x1'}; end
      function mod_x0_map; tc.BioSys.x0_map=containers.Map(); end
      function mod_xrange; tc.BioSys.xrange=containers.Map(); end
      function mod_prange; tc.BioSys.prange=containers.Map(); end
      function mod_deriv_pairs; tc.BioSys.deriv_pairs{1}='a'; end
      function mod_numsimonly; tc.BioSys.num_sim_only=true; end
      function mod_generatemex; tc.BioSys.generate_mex=true; end

    tc.BioSys.locked=false;
    verifyTrue(tc,~tc.BioSys.sys.locked_a);
    verifyTrue(tc,~tc.BioSys.flux.locked_a);
    verifyTrue(tc,~tc.BioSys.param.locked_k);
    verifyTrue(tc,~tc.BioSys.x0_map.locked_k);
    verifyTrue(tc,~tc.BioSys.xrange.locked_a);
    verifyTrue(tc,~tc.BioSys.prange.locked_a);
    end
    function simulate_sys(tc)
      %% simulate_sys
      tc.BioSys.iter_check=2; % testing continued expansion of stored solution
      tc.BioSys.num_sim_only=false;
      tc.BioSys.odesolver=@ode23t; 
      [t,y]=tc.BioSys.simulate_sys([0,100]);
      tc.verifyEqual(y,tc.exp_sol(t.').','RelTol',0.002,'AbsTol',0.002);
      
      tc.BioSys.num_sim_only=true;
      tc.BioSys.generate_mex=true;
      %% when num_sim_only is true and numeric forms are manually generated;
      %  and when generate_mex is true
      %%% make flux a matrix
      tc.BioSys.flux('rates')=repmat([tc.BioSys.sys('x1');tc.BioSys.sys('x2')],1,2);
      extract_farray=nan; x=sym('x'); ind=sym('ind');
      syms extract_farray(x,ind)
      tc.BioSys.sys('x1')=str2sym('extract_farray(rates,1)');
      tc.BioSys.sys('x2')=str2sym('extract_farray(rates,2)');
      tc.BioSys.locked=true;
      tc.BioSys.verify_mex;
      [t,y]=tc.BioSys.simulate_sys([0,100]);
      tc.BioSys.locked=false;
      tc.verifyEqual(y,tc.exp_sol(t.').','RelTol',0.002,'AbsTol',0.002);
      tc.BioSys.sys('x1')=tc.BioSys.sys('x1')+str2sym('e-2*f');
      tc.BioSys.ut=@(t) [t;0.5*t];
      tc.BioSys.locked=true;
      [t0,y0]=tc.BioSys.simulate_sys([0,100]);
      tc.BioSys.locked=false;
      tc.verifyEqual(y0,tc.exp_sol(t0.').','RelTol',0.002,'AbsTol',0.002);
      
      %% output test
      tc.BioSys.locked=true;
      for i=1:numel(t)
        tc.verifyEqual(tc.BioSys.numf_output(y(i,:).',tc.BioSys.ut(t(i))),...
          y(i,1),'RelTol',0.002,'AbsTol',0.002);
      end
      tc.BioSys.locked=false;
      
      %% verify error to parameters, states, and inputs do not conflict with
      % each other and with numeric_form output to result in confusion
      tc.BioSys.param('x1')=1;
      tc.verifyError(@()tc.BioSys.numeric_form,'BioSystem:gen_numf');
      tc.BioSys.param.remove('x1');
      tc.BioSys.param('input')=1;
      tc.verifyError(@()tc.BioSys.numeric_form,'BioSystem:gen_numf');
      tc.BioSys.param.remove('input');
            
      %% time limit test 
      tc.BioSys.iter_check=2;
      [t,~]=tc.BioSys.simulate_sys([0,100],0);
      tc.verifyEqual(size(t),[2,1]);
      
      %% xrange test 
      tc.BioSys.int_tol=-3;
      abs_tol=tc.BioSys.int_tol+tc.BioSys.int_tol_absdiff(1);
      tc.BioSys.param=cell2map({'a',-1;'b',0;'c',0;'d',-1});
      tc.BioSys.xrange=cell2map({'x1',[0.2216,Inf];'x2a',[0.8818,Inf]});
      tc.verifyError(@get_xr,'BioSystem:getxrange');
      tc.BioSys.xrange=cell2map({'x1',[-Inf,Inf];'x2',[-Inf,Inf]});
      tc.BioSys.locked=true;
      [~,y]=tc.BioSys.simulate_sys([0,1]);
      tc.verifyGreaterThanOrEqual(y,repmat([0.3677,0.7356],size(y,1),1));
      tc.verifyLessThanOrEqual(y(end,:),[0.3677,0.7356]+1e-3);

      % crash_sim test to see if simulation crashes once solution is out of range
      tc.BioSys.crash_sim=true;
      tc.BioSys.xrange=cell2map({'x1',[0.4,Inf];'x2',[0.7356,Inf]});
      tc.verifyError(@exceed,'BioSystem:simulate_sys:traj');
      function exceed()
        tc.BioSys.simulate_sys([0,1]);
      end

      % numf_full_wrapper test to see if solution stops near boundary
      tc.BioSys.crash_sim=false;
      tc.BioSys.xrange=cell2map({'x1',[0.4,Inf];'x2',[0.7356,Inf]});
      tc.BioSys.sol_locked=true;
      tc.BioSys.simulate_sys([0,1e12]);
      y=tc.BioSys.current_sol(2:end,:).';
      tc.BioSys.sol_locked=false;
      tc.verifyLessThan(y(end,:),[0.4,0.7356]);
      tc.verifyGreaterThan(y(end,:),[0.4,0.7356]-1.1*10^tc.BioSys.int_tol);
      y_diff_from_lr=y-[0.4,0.7356];
      tc.verifyNotEqual(sum(y_diff_from_lr(:,1)<0),sum(y_diff_from_lr(:,2)<0));
      function get_xr()
        tc.BioSys.xrange;
      end
      %%% output test
      tc.BioSys.locked=false;
      tc.BioSys.output=str2sym({'x1';'x2'});
      numf_out_full_wrapped=tc.BioSys.numf_output_full_wrapped;
      p=cell2mat(tc.BioSys.param.values.');
      ut=tc.BioSys.ut;
      tc.verifyEqual(numf_out_full_wrapped(p,[0.3;0.6],ut(0)),[0.4;0.7356]);
      
      tc.BioSys.locked=true;
      %% allset_f test
      i=0;
      tc.BioSys.allset_f=@allset_f;
      [t,y_sol]=tc.BioSys.simulate_sys([0,100]);
      verifyEqual(tc,numel(t),10);
      verifyLessThan(tc,max(t),100);
      function status = allset_f (~,~)
        i=i+1;
        if i==10
          status=1;
        else
          status=0;
        end
      end
      
      %% lastsimindex test
      tc.BioSys.allset_f=@allset_f_lastsimindex;
      [~,y_sol_2]=tc.BioSys.simulate_sys([0,100]);
      verifyEqual(tc,y_sol,y_sol_2);
      function status = allset_f_lastsimindex(~,y)
        lastsimindex=tc.BioSys.lastsimindex;
        if lastsimindex<size(y_sol,1) && all(y_sol(lastsimindex,:)==y.')
          % error in lastsimindex results in premature termination
          status=0;
        else
          status=1;
        end
      end
    end
    function simulate_sys_myode15s(tc)
      tc.BioSys.locked=true;     

      % error estimation based on halving step size and solutions derived
      % from Euler's method
      [t2,y2]=tc.BioSys.simulate_sys_myode15s_halft([0,1e4],inf);
      tc.verifyEqual(y2,tc.exp_sol(t2.').','RelTol',0.05,'AbsTol',0.05);
      
      % error estimation based on solution difference between Euler's method and
      % Heun's method
      [t3,y3]=tc.BioSys.simulate_sys_myode15s_eulerheuns([0,1e4],inf);
      tc.verifyEqual(y3,tc.exp_sol(t3.').','RelTol',0.0005,'AbsTol',0.0005);
    end
    function view_solution(tc)
      figure('Name','view_solution test');
      [t,y,full_y]=tc.BioSys.view_solution(100,{'x2','o1','x1'});
      exp_sol=tc.exp_sol(t.').';
      tc.verifyEqual(y,exp_sol(:,[2,1,1]),'RelTol',0.002,'AbsTol',0.002);
      tc.verifyEqual(full_y,exp_sol(:,[1,2,1]),...
        'RelTol',0.002,'AbsTol',0.002);
      
      figure('Name','view_solution scale test');
      log10m=cell(1,4);
      [log10m{:}]=log10m_trans(10);
      tc.verifyEqual(log10m{1}([-10,0,10,100,1000]),...
        [-log10(100+1),0,log10(100+1),log10(1001),log10(10001)]);
      tc.verifyEqual(log10m{2}(log10m{1}([-10,0,10,100,1000])),...
        [-10,0,10,100,1000],'RelTol',1e-8,'AbsTol',1e-8);
      tc.BioSys.view_solution(100,{'x2','o1','x1'},10,log10m);
      set(gca,'YLim',[0,log10(100+1)]);
    end
    function view_solutions(tc)
      figure('Name','view_solutions test');
      [t,y]=tc.BioSys.simulate_sys([0,100]);
      BioSystem.view_solutions({t,y;t,y+0.1},tc.BioSys.statevars,...
        tc.BioSys.statevars,[0,10]);
    end
    function find_dose_response(tc)
      tc.BioSys.convert_to_process({'a'});
      tc.BioSys.locked=true;
      x0=tc.BioSys.x0;
      
      in=(-5:0.1:-4).';
      ss_func_exp=repmat([0,0],length(in),1);
      [ss_func,ss_fluct]=tc.BioSys.find_dose_response(in,[2000,4000]);
      tc.verifyEqual(ss_func,ss_func_exp,'RelTol',0.0001,'AbsTol',0.0001);
      tc.verifyEqual(ss_fluct,zeros(size(ss_fluct)),'RelTol',0.0001,'AbsTol',0.0001);
      tc.verifyEqual(tc.BioSys.x0,x0);
      
      % testing for warning cases
      tc.BioSys.ut=@(t)in(1,:).';
      [t,y]=tc.BioSys.simulate_sys([0,5]);
      tc.BioSys.iter_check=numel(t)+1;
      tc.BioSys.ut=@(t)in(1,:);
      tc.BioSys.sol_locked=true;
      [ss_func]=tc.verifyWarning(@exp_insufsimtime_prevcutoff,...
        'BioSystem:find_steadystate:insufficientsimtime');
      ss_t=tc.BioSys.current_sol(1,find(tc.BioSys.current_sol(1,:)>5,1,'first'));
      tc.BioSys.sol_locked=false;
      [t,y]=tc.BioSys.simulate_sys([0,ss_t]);
      tc.verifyEqual(ss_func,y(end,:),'AbsTol',1e-16);
      tc.BioSys.iter_check=1;
      [ss_func,ss_fluct]=tc.verifyWarning(@exp_insufsimtime,...
        'BioSystem:find_steadystate:insufficientsimtime');
      verifyTrue(tc,all(all(isnan(ss_func))));
      verifyTrue(tc,all(all(isnan(ss_fluct))));
      tc.verifyWarning(@exp_ltimewin,'BioSystem:find_steadystate');
      function [ss,fluct]=exp_insufsimtime_prevcutoff()
        [ss,fluct]=tc.BioSys.find_steadystate([0,5],1e-16);
      end
      function [ss,fluct]=exp_insufsimtime()
        [ss,fluct]=tc.BioSys.find_dose_response(in(1:2,:),[2000,4000],0);
      end
      function exp_ltimewin()
        tc.BioSys.find_dose_response(in(1:2,:),[0.1,0.2]);
      end
    end
    function find_dose_response2(tc)
      dx={'x','-x-d';'y','-d-2*y'};
      flux={'o1','x+y'};params={'dummy',0};x0_map={'x',1;'y',1};
      A=BioSystem(cell2map(dx),cell2map(flux),cell2map(params),cell2map(x0_map));
      A.output='o1';
      
      [ss,ss_fluct,o,o_fluct]=find_dose_response(A,[0.5,0.6,0.7].',[2000,2500]);
      ss_exp=[-0.5,-0.25;-0.6,-0.3;-0.7,-0.35];
      o_exp=[-0.75;-0.9;-1.05];
      tc.verifyEqual(ss,ss_exp,'RelTol',0.0001,'AbsTol',0.0001);
      tc.verifyEqual(o,o_exp,'RelTol',0.0001,'AbsTol',0.0001);
      tc.verifyEqual(ss_fluct,zeros(size(ss_fluct)),'RelTol',0.0001,'AbsTol',0.0001);
      tc.verifyEqual(o_fluct,zeros(size(o_fluct)),'RelTol',0.0001,'AbsTol',0.0001); 
    end
    function eval_eig(tc)
      tc.BioSys.num_sim_only=false;
      tc.verifyEqual(eig([-5,1;4,-2]),tc.BioSys.eval_eig([1;1]));
      tc.BioSys.num_sim_only=true;
      tc.verifyEqual(eig([-5,1;4,-2]),tc.BioSys.eval_eig([1;1]));
    end
    function satisfy_constrs(tc)
      L=tc.BioSys.satisfy_constrs({'a<b';'b>c'});
      tc.verifyEqual(L,[true;false])
    end
    function get_feasible(tc)
      tc.BioSys.get_feasible({'a>b'},...
        cell2map({'a',[0,10];'b',[0,10];'c',[0,10];'d',[0,10]}));
      tc.verifyTrue(tc.BioSys.satisfy_constrs({'a>b'}));
      
      b=tc.BioSys.param('b');c=tc.BioSys.param('c');d=tc.BioSys.param('d');
      tc.BioSys.get_feasible({'a>c'},cell2map({'a',[0,10]}));
      tc.verifyTrue(tc.BioSys.satisfy_constrs({'a>c'}));
      tc.verifyEqual(tc.BioSys.param('b'),b);
      tc.verifyEqual(tc.BioSys.param('c'),c);
      tc.verifyEqual(tc.BioSys.param('d'),d);
      
      inequalities=tc.BioSys.gen_constrs({'a>b'});
      tc.BioSys.get_feasible(inequalities,...
        cell2map({'a',[0,10];'b',[0,10];'c',[0,10];'d',[0,10]}));
      
      %new feature: parameter specification allow direct index to be passed in. 
      tc.verifyTrue(tc.BioSys.satisfy_constrs({'a>b'}));
      tc.BioSys.get_feasible(inequalities,cell2map({1,[0,10];2,[0,10];3,[0,10];4,[0,10]}));
      tc.verifyTrue(tc.BioSys.satisfy_constrs({'a>b'}));
      tc.BioSys.get_feasible(inequalities,cell2map({1,[0,10]}));
      tc.verifyTrue(tc.BioSys.satisfy_constrs({'a>b'}));
      b=tc.BioSys.param('b');c=tc.BioSys.param('c');d=tc.BioSys.param('d');
      tc.verifyEqual(tc.BioSys.param('b'),b);
      tc.verifyEqual(tc.BioSys.param('c'),c);
      tc.verifyEqual(tc.BioSys.param('d'),d);
    end
    function get_feasible_sw(tc)
      inequalities=tc.BioSys.gen_constrs({'a>b';'c>d'}); steps={1,{1,2},Inf;2,{3,4},Inf};
      tc.BioSys.param('a')=tc.BioSys.param('b')-1; tc.BioSys.param('c')=tc.BioSys.param('d')-1;
      tc.verifyTrue(all(~inequalities(cell2mat(tc.BioSys.param.values).')));
      
      tc.BioSys.get_feasible_sw(inequalities,...
        cell2map({1,[0,10];2,[0,10];3,[0,10];4,[0,10]}),steps);
      tc.verifyTrue(all(inequalities(cell2mat(tc.BioSys.param.values).')));
    end
    
    function get_feasibles(tc)
      range=cell2map({'a',[0,10];'b',[0,10];'c',[0,10];'d',[0,10]});
      
      tc.BioSys.get_feasibles({'a>b'},range,10,'test1.csv');
      params=readtable('test1.csv');
      verifyGreaterThan(tc,params.a,params.b);
      
      params_old=params;
      tc.BioSys.get_feasibles({'a>b'},range,params(:,'a'),'test2.csv');
      params=readtable('test2.csv');
      verifyGreaterThan(tc,params.a,params.b);
      verifyEqual(tc,params.a,params_old.a);
      
      tc.verifyError(@()tc.BioSys.get_feasibles({'a>b'},...
        range,params(:,'a'),'test3.csv',{1,{1},Inf}),'BioSystem:get_feasibles');
      tc.BioSys.get_feasibles({'a>b'},range,params(:,'a'),'test3.csv',{1,{2},Inf});
      params=readtable('test3.csv');
      verifyGreaterThan(tc,params.a,params.b);
      verifyEqual(tc,params.a,params_old.a);
      delete('test1.csv','test2.csv','test3.csv');
    end
    function get_infeasible(tc)
      tc.BioSys.get_infeasible({'a<b';'b<c'},'b','up',0.1,10000);
      tc.verifyFalse(all(tc.BioSys.satisfy_constrs({'a<b';'b<c'})));
    end
    function get_infeasible_almost(tc)
      tc.BioSys.get_infeasible({'a<b';'b<c'},'b','up',0.1,10000,true);
      tc.verifyTrue(all(tc.BioSys.satisfy_constrs({'a<b';'b<c'})));
    end
    function take_limit(tc)
      vecfun_form=str2sym({'x1-(a-e)*x2';'c*x1+d*x2'});
      param=[tc.BioSys.param;containers.Map()];
      param.remove({'b'});
      param('e')=0;
      tc.BioSys.take_limit({'a','b'},{'1','-a+e'});
      tc.verifyEqual(tc.BioSys.vecfun_form,vecfun_form);
      tc.verifyEqual(map2cell(tc.BioSys.param),...
        map2cell(param));
    end
    function replace1(tc)
      vecfun_form=tc.BioSys.vecfun_form;
      vecfun_form=subs(vecfun_form,str2sym('a'),str2sym('b+x1'));
      param=[tc.BioSys.param;containers.Map()];
      param.remove({'a'});
      tc.BioSys.replace('a','b+x1');
      tc.verifyEqual(tc.BioSys.vecfun_form,vecfun_form);
      tc.verifyEqual(map2cell(tc.BioSys.param),...
        map2cell(param));
      tc.BioSys.replace('x2',sym('x1'));
      tc.verifyEqual(tc.BioSys.vecfun_form,str2sym('x1*(b+x1)+b*x1'));
      tc.verifyEqual(numel(tc.BioSys.x0),1);
    end
    function replace2(tc)
      tc.verifyError(@()tc.BioSys.replace('a','f1'),'replace:ClassIncompatibility');
    end
    function convert_to_process(tc)
      vecfun_form=tc.BioSys.vecfun_form;
      vecfun_form=vecfun_form(1);
      param=[tc.BioSys.param;containers.Map()];
      param.remove({'a','c','d'});
      tc.BioSys.convert_to_process({'x2','a'});
      tc.verifyEqual(tc.BioSys.vecfun_form,vecfun_form);
      tc.verifyTrue(all(~ismember(tc.BioSys.param.keys,'a')));
      tc.verifyEqual(tc.BioSys.inputvars,{'a','x2'});
    end
    function apply_conservation(tc)
      tc.BioSys.replace('a','-c');
      tc.BioSys.replace('b','-d');
      tc.BioSys.apply_conservation({'x1',1;'x2',1},'T','x1');
      tc.verifyEqual(tc.BioSys.vecfun_form,str2sym('c*(T-x2)+d*x2'))
      tc.verifyEqual(tc.BioSys.param.keys,{'T','c','d'});
    end
    function copytest(tc)
      cpObj=copy(tc.BioSys);
      tc.verifyEqual(map2cell(cpObj.param),map2cell(tc.BioSys.param));
      cpObj.param('a')=cpObj.param('a')-1;
      tc.verifyNotEqual(map2cell(cpObj.param),map2cell(tc.BioSys.param));
    end
    function combine(tc)
      o1=tc.BioSys;
      o2=copy(tc.BioSys);
      o3=copy(tc.BioSys); 
      o3.xrange.remove('x1'); o3.x0_map.remove('x1'); o3.sys.remove('x1');
      o3.xrange('x3')=[0,10]; o3.x0_map('x3')=0; o3.sys('x3')=str2sym('-x3+x2'); 
      o4=copy(tc.BioSys); o4.sys('x1')=sym('1');
      
      properties={'param','flux','sys','x0_map','xrange','prange'};
      o1_o2=BioSystem.combine(o1,o2);
      for p_l=properties 
        verifyEqual(tc,o1_o2.(p_l{:}).keys,o1.(p_l{:}).keys); 
        verifyEqual(tc,o1_o2.(p_l{:}).values,o1.(p_l{:}).values); 
      end
      
      o1_o3=BioSystem.combine(o1,o3);
      o1_o3_exp=copy(o1); 
      o1_o3_exp.xrange('x3')=[0,10]; 
      o1_o3_exp.x0_map('x3')=0; 
      o1_o3_exp.sys('x3')=str2sym('-x3+x2'); 
      verifyEqual(tc,o3.sys.keys,{'x2','x3'});
      for p_l=properties 
        verifyEqual(tc,o1_o3.(p_l{:}).keys,o1_o3_exp.(p_l{:}).keys); 
        verifyEqual(tc,o1_o3.(p_l{:}).values,o1_o3_exp.(p_l{:}).values); 
      end
      
      verifyError(tc,@()BioSystem.combine(o1,o4),'combine:conflict');
    end
    convert_to_acceleration_test(tc);
    XNetworks(tc);
    utilities_test(tc);
    reverseAutoDiff_test(tc);
  end
end
```
