## Supplemental Data 1 for "Principles underlying implementation of *nearly*-homeostatic biological networks": convert_to_acceleration_test.m

```
function convert_to_acceleration_test(tc)
sv={'x1','x2'};
tc.BioSys.output=sym(sv.');
tc.BioSys.output_label=sv.';
newSys=tc.BioSys.copy;
convert_to_acceleration(newSys,tc.BioSys.statevars,'f');
verifyEqual(tc,newSys.sys('x1'),sym('x1_v'));
verifyEqual(tc,newSys.xrange('x1_v'),[-inf,inf]);
verifyEqual(tc,newSys.sys('x1_v'),tc.BioSys.sys('x1')-str2sym('f*x1_v'));
verifyEqual(tc,newSys.inputvars,tc.BioSys.inputvars);
verifyEqual(tc,tc.BioSys.x0_map.values,newSys.x0_map.values(sv));
newSys2=tc.BioSys.copy;
newSys2.sys('f')=0;
verifyError(tc,@()convert_to_acceleration(newSys2,sv,'f'),...
  'BioSystem:convert_to_acceleration:statevars');

% acceleration vectors results in vastly higher fluctuation around steady
% state
[x_sys,x_sys_fluct]=find_steadystate(tc.BioSys,[500,600],2);
[~,~,x_nsys,x_nsys_fluct]=find_steadystate(newSys,[500,600],2);
verifyGreaterThan(tc,x_nsys_fluct./1e9,x_sys_fluct);
verifyEqual(tc,x_sys,x_nsys,'AbsTol',1e-4);
end
```
