## Supplemental Data 1 for "Principles underlying implementation of *nearly*-homeostatic biological networks": reverseAutoDiff_test.m

```
function reverseAutoDiff_test(tc)
% Define symbolic variables
syms z1 z2 V real
syms h1 h2 f1 f2 real

% Create a container map for definitions
defs = containers.Map('KeyType','char','ValueType','any');

% Define relationships without substituting intermediates, outputs, and inputs
defs('h1') = V*z1^2;
defs('h2') = sin(z2);
defs('f1') = z1*z2*h1*h2;
defs('f2') = h2;
defs('f3') = f2;
outputs = [V*f1*f2 + f1^3*(f2^2) + z1*z2,1;
  f1 + f2^2,1; 
  V^2*f1*f2,1];
outputs_label={'o1','o4';'o2','o5';'o3','o6'};
inputs = [z1, z2];

% Compute the Jacobian and verify
[dX_dZ, derivatives, topoOrder] = reverseAutoDiff(defs, outputs, outputs_label, inputs);
dX_dZ_full = subs(dX_dZ,sym(derivatives.keys),sym(derivatives.values));
dX_dZ_full = subs(dX_dZ_full,sym(defs.keys),sym(defs.values));
for i=1:100
  dX_dZ_full_new=subs(dX_dZ_full,sym(derivatives.keys),sym(derivatives.values));
  dX_dZ_full_new=subs(dX_dZ_full_new,sym(defs.keys),sym(defs.values));
  if isequal(dX_dZ_full_new,dX_dZ_full)
    break;
  else
    dX_dZ_full=dX_dZ_full_new;
  end
end
defs_full=BioSystem.fullexp(defs,defs);
dX_dZ1_full_exp=diff(subs(outputs,sym(defs_full.keys),cell2sym(defs_full.values)),inputs(1));
verifyTrue(tc,isequal(simplify(dX_dZ_full(:,:,1)-dX_dZ1_full_exp),sym(zeros(3,2))));

% additional verifications: 
% - cyclic dependency detection
% - topoOrder does not contain 'f3', 
% - derivatives does not contain dh1_d_V,
% - derivative of outputs(2,1) with respect to z1 should NOT involve f2 at all
testdep=cell2map({'x1',{'x2','x3'};'x2',{'x1'};'x3','x4'});
verifyError(tc,@()topologicalSort(testdep,testdep.keys),'topologicalSort:cyclic_dependencies');
verifyTrue(tc,~ismember('f3',topoOrder));
verifyTrue(tc,~ismember('dl_h1_r_dl_V_r',derivatives.keys));
verifyTrue(tc,ismember('dl_h1_r_dl_z1_r',derivatives.keys));
verifyTrue(tc,isempty(regexp(char(dX_dZ(2,1,1)),'dl_o2_r_dl_f2_r','once')));
verifyTrue(tc,~isempty(regexp(char(dX_dZ(2,1,1)),'dl_o2_r_dl_f1_r','once')));
end
```
