## Supplemental Data 1 for "Principles underlying implementation of *nearly*-homeostatic biological networks": utilities_test.m

```
function utilities_test(tc)
% Unit testing of certain utilities

%% ref1_array
for c={'ref1_array','ref1_array_struct'}
  rng(1,'twister');
  m=rand(3.15e3);
  m_c=m;
  eval(['m_r1a=',c{:},'(size(m));']);
  m_r1a.mod(':',':',m);
  tic; m(1)=1; t_m=toc;
  tic; for i=1:100; m(i)=i; end; t_m_nocopy=toc/100;
  tic; for i=1:100; m_r1a.mod(i,i); end; t_mr1a=toc/100;
  verifyLessThan(tc,t_mr1a,t_m/100);
  verifyEqual(tc,m_r1a.size(),size(m));
  verifyEqual(tc,m_r1a.read(':',':'),m);
  m_r1a.subset(1:2,1:2);
  verifyEqual(tc,m_r1a.read(':',':'),m(1:2,1:2));
end

%% convert2_absthres
verifyEqual(tc,convert2_absthres([0.1,0.1],[2,0.1]),[0.2,0.1]);

%% smoothstep_shifted, smoothstep_shifted_d1, smoothstep_std, smoothstep_std_d1
verifyEqual(tc,smoothstep_shifted([1,2,3.5,5,6],4,3.5,[2,5]),[0,0,0.5,1,1]); % smoothstep_shifted
verifyLessThan(tc,smoothstep_shifted(0.6167-0.15,4,0.6167,[0,1]),0.15); % 0.6167+-0.15
verifyGreaterThan(tc,smoothstep_shifted(0.6167+0.15,4,0.6167,[0,1]),0.85);
verifyEqual(tc,smoothstep_shifted([0,1],4,0.6,[0,1]),[0,1]); % boundary values
% smoothstep_shifted_d1
verifyEqual(tc,diff(smoothstep_shifted([0.6,0.6+1e-6],4,0.6,[0,1]))/1e-6,... 
  smoothstep_shifted_d1(0.6,4,0.6,[0,1]),'AbsTol',1e-6,'RelTol',1e-6);
verifyEqual(tc,smoothstep_shifted_d1([-1,2],4,0.6,[0,1]),[0,0]);
% smoothstep_shifted_Rnoband
verifyGreaterThan(tc,smoothstep_shifted_Rnoband(4,0.6167,1),1-1e-3);
verifyEqual(tc,smoothstep_shifted_Rnoband(4,1.25,1.05),0);
verifyEqual(tc,smoothstep_shifted_Rnoband(4,1.25,1.5),1);
% smoothstep_shifted_d1_Rnoband
verifyEqual(tc,smoothstep_shifted_d1_Rnoband(4,0.6167,0.6),...
  diff(smoothstep_shifted_Rnoband(repmat(4,1,2),repmat(0.6167,1,2),[0.6,0.6+1e-8]))/1e-8,...
  'AbsTol',1e-6,'RelTol',1e-6);
verifyEqual(tc,smoothstep_shifted_d1_Rnoband(4,1.1,1.15),...
  diff(smoothstep_shifted_Rnoband(repmat(4,1,2),repmat(1.1,1,2),[1.15,1.15+1e-8]))/1e-8,...
  'AbsTol',1e-6,'RelTol',1e-6);
verifyEqual(tc,smoothstep_shifted_d1_Rnoband(4,1.4,1.5),...
  diff(smoothstep_shifted_Rnoband(repmat(4,1,2),repmat(1.4,1,2),[1.5,1.5+1e-8]))/1e-8,...
  'AbsTol',1e-6,'RelTol',1e-6);

%% shifted_identity, shifted_identity_d1, shifted_identity_Rnoband, shifted_identity_dx_Rnoband test
% shifted_identity
verifyEqual(tc,shifted_identity(0.6,0.6,[0,1],1e-2),0.6,'RelTol',1e-15);
verifyEqual(tc,shifted_identity((1-1e-2+0.6)/2,0.6,[0,1],1e-2),0.8,'RelTol',1e-15);
verifyEqual(tc,shifted_identity((1e-2+0.6)/2,0.6,[0,1],1e-2),0.3,'RelTol',1e-15);
verifyEqual(tc,shifted_identity(1-1e-3,0.6,[0,1],1e-2),1);
verifyEqual(tc,shifted_identity(1e-3,0.6,[0,1],1e-2),0);
verifyEqual(tc,shifted_identity(1.89,1.6,[1,2],1e-1),1/0.8*(0.89-0.1)+1,'RelTol',1e-15);
% shifted_identity_d1
for v=[1e-3,0.4,0.8,1-1e-3]
  verifyEqual(tc,shifted_identity_dx(v,0.6,[0,1],1e-2),...
    (shifted_identity(v+1e-6,0.6,[0,1],1e-2)-shifted_identity(v,0.6,[0,1],1e-2))/1e-6,'RelTol',1e-9);
end
% shifted_identity_Rnoband
verifyEqual(tc,shifted_identity_Rnoband((1-1e-2+0.6)/2,0.6,1),0.8,'RelTol',1e-15);
verifyEqual(tc,shifted_identity_Rnoband((1+1e-2+1.6)/2,1.6,1),1.3,'RelTol',1e-15);
verifyEqual(tc,shifted_identity_Rnoband(0,1.6,1),1,'RelTol',1e-15);
verifyEqual(tc,shifted_identity_Rnoband((1-1e-2+0.6)/2,0.6,0),(1-1e-2+0.6)/2);
% shifted_identity_dx_Rnoband
verifyEqual(tc,shifted_identity_dx_Rnoband((1-1e-2+0.6)/2,0.6,0),1);

%% symvarfun and subsd1 
% - also test whether subsd1 also converts diff(f(x,y),x) into that standard D([1],f)(x,y) form.
% - also test whether diff(f(x,y),x) + diff(f(y,x),x) gets converted into two expressions instead of
%   one.
M=nan;
inv=nan;
fv=nan;
syms M [2 2] matrix
syms inv(M) [2 2] matrix keepargs
syms fv(x,y) [2 1] matrix keepargs
cell_sym={str2sym('diff(func(g(x,h(z)),y,z),y)+diff(2*func(y,g(x,h(z)),z),y)+2*diff(f(x,y),x)'),...
  [str2sym('diff(g(2*x,3*x),x)');str2sym('k(x,f(x,y))')],...
  inv(M)*diff(fv(sym('y'),sym('y')))};
d1pairs={'D([2],func)','dfuncdy2'; 'D([1],func)','dfuncdy1';
  'D([1],f)','dfdx'; 'D([1],g)','dgdx'; 'D([2],g)','dgdy'};
cell_char={char(cell_sym{1}),char(cell_sym{2}),evalc('disp(cell_sym{3})')};
% symvarfun input check
verifyError(tc,@()symvarfun(cell_char),'symvarfun:input_check');
% symvarfun check
cell_char=regexprep(cell_char,{'\n','\s+'},{'',''});
func_info=symvarfun(cell_char);
verifyEqual(tc,height(func_info),9);
verifyEqual(tc,func_info.numargs('f'),2);
verifyEqual(tc,func_info.numargs('func'),3);
verifyEqual(tc,func_info.numargs('k'),2);
verifyEqual(tc,func_info.numargs('<strong>inv</strong>'),1);
% d1pairs check
verifyError(tc,@()BioSystem.subsd1({str2sym('fname(x)')},d1pairs),'BioSystem:subsd1:variable_conflict');
cell_sym_out=BioSystem.subsd1(cell_sym,d1pairs);
verifyEqual(tc,cell_sym_out{1},str2sym('dfuncdy2(g(x,h(z)),y,z)+2*dfuncdy1(y,g(x,h(z)),z)+2*dfdx(x,y)'))
end
```
