## Supplemental Data 1 for "Principles underlying implementation of *nearly*-homeostatic biological networks": XNetworks.m

```
function XNetworks(tc)
rng(1,'twister');

%%  Testing for GeneticNetwork
% expected expressions
A=GeneticNetwork(...
  {'uplinear','none','none','none'; ...
  'up','none','none','none';
  'down','none','none','none'; ...
  'unknown','none','none','none'},...
  {'normal','linear','zeroorder','zeroorder'});
exp_vecfun123=str2sym(...
  {'V1*(b1_1 - (x1^n1_1*(b1_1 - 1))/Kd1_1) - (Vd1*x1^dn1)/(dKd1^dn1 + x1^dn1)'; 
  'V2*(b2_1 - (x1^n2_1*(b2_1 - 1))/(Kd2_1^n2_1 + x1^n2_1)) - (Vd2*x2^dn2)/dKd2'; 
  'V3*(b3_1 - (Kd3_1^n3_1*(b3_1 - 1))/(Kd3_1^n3_1 + x1^n3_1)) - Vd3'});
verifyEqual(tc,A.vecfun_form(1:3),exp_vecfun123);
verifyEqual(tc,A.xrange('x1'),[-1.1e-4,inf]);
A.output=str2sym('s4_1_act');
A.output_label={'s4_1_act'};

% testing gen_proddegrates
Ac=A.copy;
Ac.convert_to_process({'x1'});
Ac.prange('x1')=[0,10];
Ac.param('x1')=1;
[~,numf_prod,~,numf_deg]=Ac.gen_proddegrates();
p_rand=rand(Ac.param.Count,1);
x_rand=rand(Ac.sys.Count,1);
numf_net_exp=Ac.numeric_form_full;
for i=1:Ac.sys.Count
  x=Ac.statevars{i};
  numf_prod_x=numf_prod(x);
  numf_deg_x=numf_deg(x);
  numf_net_x=@(p,x)numf_prod_x(p,x)-numf_deg_x(p,x);
  verifyEqual(tc,numf_net_x(p_rand,x_rand),extract_farray(numf_net_exp(p_rand,x_rand),i),'AbsTol',1e-8);
  verifyTrue(tc,numf_net_x(p_rand,x_rand)~=0);
end

% topo_drefs=[0,0.75,1.5]; % corresponding to tanh based general form
topo_drefs=[0,1,2];

% testing GeneticNetwork.discretize_bf
verifyEqual(tc,GeneticNetwork.discretize_bf(0.1*topo_drefs(end)),topo_drefs(1));
verifyEqual(tc,GeneticNetwork.discretize_bf(0.3*topo_drefs(end)),topo_drefs(1));
verifyEqual(tc,GeneticNetwork.discretize_bf(0.35*topo_drefs(end)),topo_drefs(2));
verifyEqual(tc,GeneticNetwork.discretize_bf(0.65*topo_drefs(end)),topo_drefs(2));
verifyEqual(tc,GeneticNetwork.discretize_bf(0.7*topo_drefs(end)),topo_drefs(3));

% s to s_act's derivatives: softstep edition
sma=@(x)softstep(6,0.6167,x)+softstep(6,2-0.6167,x);
x=0:1e-5:2;
sm=arrayfun(sma,x);
dsmdx=diff(sm)/1e-5;
max_dsmdx=max(dsmdx);
dsmdx_minatmid=min(dsmdx(x>=0.5 & x<=1.5));
verifyGreaterThan(tc,dsmdx,0);
verifyLessThan(tc,dsmdx_minatmid,max_dsmdx/10);
verifyGreaterThan(tc,dsmdx_minatmid,max_dsmdx/20);

% s to s_act: check to see if discretization allows true discrete topology to be reached and only
% nearly discrete topology to be reached otherwise. 
Ac=A.copy;
Ac.prange('dobj_s4_1_dval_1')=[0,2];
Ac.param('dobj_s4_1_dval_1')=0;
Ac.prange('dobj_s4_1_dval_2')=[0,2];
Ac.param('dobj_s4_1_dval_2')=1;
Ac.prange('dobj_s4_1_dval_3')=[0,2];
Ac.param('dobj_s4_1_dval_3')=2;
Ac.locked=true;
verifyEqual(tc,Ac.param('s4_1_don'),1);
Ac.param('s4_1')=0;
verifyEqual(tc,Ac.numf_output(A.x0),0);
Ac.param('s4_1')=1;
verifyEqual(tc,Ac.numf_output(A.x0),1);
Ac.param('s4_1')=2;
verifyEqual(tc,Ac.numf_output(A.x0),2);
Ac.gd_reset(Ac,{'s4_1'});
Ac.param('s4_1')=0; 
verifyGreaterThan(tc,Ac.numf_output(A.x0),0);
Ac.param('s4_1')=1; 
verifyEqual(tc,Ac.numf_output(A.x0),1);
Ac.param('s4_1')=2; 
verifyLessThan(tc,Ac.numf_output(A.x0),2);


% unknown function properties test
Ac=A.copy;
take_limit(Ac,{'n4_1'},{1});
sk=[0.9 1].*topo_drefs(end)/3; 
test_band=[3e-1 5e-3]; 
sn=[6 5e2];  
% test_band=[3e-1]; 
% sn=[3]; 
for i=1:numel(test_band)
  Ac.flux('sn')=sn(i);
  Ac.flux('sk')=sk(i);
  test_map=containers.Map({'fH4_1'},{Ac.flux('fH4_1')});
  test_map_fullexp=BioSystem.fullexp(test_map,Ac.flux);
  test_numf=matlabFunction(test_map_fullexp('fH4_1'));

  % softstep tanh based topology parametrization
  verifyTrue(tc,test_numf(1,topo_drefs(1),0,0)<1e-2); % near discrete topology at discrete topo param
  verifyTrue(tc,test_numf(1,topo_drefs(1),0,1e6)>0.9999);
  verifyTrue(tc,test_numf(1,topo_drefs(2),0,0)>0.99);
  verifyTrue(tc,test_numf(1,topo_drefs(2),0,1e6)>0.99);
  verifyTrue(tc,test_numf(1,topo_drefs(3),0,1e12)<1e-2);
  sk_true=fzero(@(s)test_numf(1,s,0,0)-0.5,[0,1]);
  verifyTrue(tc,test_numf(1,sk_true-test_band(i)/2,0,0)<0.22); % testing sharpness of topo transition
  verifyTrue(tc,test_numf(1,sk_true+test_band(i)/2,0,0)>0.80);
  sk_true=fzero(@(s)test_numf(1,s,0,1e12)-0.5,[1,2]);
  verifyTrue(tc,test_numf(1,sk_true-test_band(i)/2,0,1e12)>0.80);
  verifyTrue(tc,test_numf(1,sk_true+test_band(i)/2,0,1e12)<0.22);
  verifyTrue(tc,test_numf(1,topo_drefs(3),0,1e12)<1e6);
end

%% Testing for MNetwork
A=MNetwork({'yes','no','linear'},{'no','yes','no'});
exp_vecfun=str2sym(['(V1_1*x1*(1 - x1)^n1_1)/((1 - x1)^n1_1 + Kd1_1^n1_1) - ',...
  '(Vd1_2*x1^dn1_2*x2)/(dKd1_2^dn1_2 + x1^dn1_2) + (V1_3*x3*(1 - x1)^n1_3)/Kd1_3']);
verifyEqual(tc,A.vecfun_form,exp_vecfun);
verifyEqual(tc,A.xrange('x1'),[-1.1e-4,1+1.1e-4]);

%% Testing for RNetwork
% verify whether both modes generate expected rates, prod rates, and degradation rates when 
% coefficients are integers
for modes_c={'normal','general2'}
  mode=modes_c{:};
  %%% x1 (f) + x2 (s) -> x3 (l); nothing -> x2
  sv2x=@(sv)[sv(1);sv(3);sv(2)]; % f,l,s -> 
  x2sv=@(x)[x(1);x(3);x(2)]; % x -> f,l,s
  smr=[[1;1;0],[0;0;0]];
  A=RNetwork(smr,[[0;0;1],[0;1;0]],mode,{'f','s','l'}); 
  A.prep4_sma;
  verifyEqual(tc,A.sma.r,smr) % sma test
  A.convert_to_process({'gfp_k','gfp_n'});
  krf1=2;
  krf2=1;
  A.param('krf1')=krf1;
  A.param('krf2')=krf2;
  numf_A=A.numeric_form;
  numf_A_exp=@(x)[-krf1*x(1)*x(2); -krf1*x(1)*x(2)+krf2; krf1*x(1)*x(2)];
  verifyEqual( tc , sv2x(numf_A(x2sv([3;2;4]),[2/3;4])) , numf_A_exp([3;2;4]) );
  %%% change smr and see whether the reaction network changes as expected
  A.flux('smr_1_1')=2; % 2 x1 + x2 -> x3; nothing -> x2
  numf_A=A.numeric_form;
  numf_A_exp=@(x)[-2*krf1*x(1)^2*x(2); -krf1*x(1)^2*x(2)+krf2; krf1*x(1)^2*x(2)];
  verifyEqual( tc , sv2x(numf_A(x2sv([3;2;4]),[2/3;4])) , numf_A_exp([3;2;4]) );
  verifyEqual( tc , sv2x(numf_A(x2sv([0;2;4]),[2/3;4])) , numf_A_exp([0;2;4]) );
  verifyEqual( tc , sv2x(numf_A(x2sv([2;0;4]),[2/3;4])) , numf_A_exp([2;0;4]) );
  %%% verify whether prod rate - deg rate = actual rate
  [~,numf_prod,~,numf_deg]=gen_proddegrates(A);
  convert_to_process(A,{'x1'});
  p=cell2mat(A.param.values).';
  numf_prod_v=numf_prod.values;
  numf_prod_vec=@(x,u)cellfun(@(f)f(p,x,u),numf_prod_v).';
  numf_deg_v=numf_deg.values;
  numf_deg_vec=@(x,u)cellfun(@(f)f(p,x,u),numf_deg_v).';
  verifyEqual( tc , ...
    numf_prod_vec(x2sv([3;2;4]),[2/3;4])-numf_deg_vec(x2sv([3;2;4]),[2/3;4]) , ...
    numf_A(x2sv([3;2;4]),[2/3;4]) );
  verifyEqual( tc , ...
    numf_prod_vec(x2sv([0;2;4]),[2/3;4])-numf_deg_vec(x2sv([0;2;4]),[2/3;4]) , ...
    numf_A(x2sv([0;2;4]),[2/3;4]) );
end

% general2 mode verifications - basic version: sm to sma mapping's derivatives
gfp_n=3;
sma=@(x)smoothstep_shifted_Rnoband(gfp_n,0.6167,x)/smoothstep_shifted_Rnoband(gfp_n,0.6167,1) - ...
  (1/smoothstep_shifted_Rnoband(gfp_n,0.6167,1)-1)*smoothstep_shifted_Rnoband(1,1.3,x) + ...
  smoothstep_shifted_Rnoband(gfp_n,2-0.6167,x);
x=0:1e-5:2;
sm=arrayfun(sma,x);
dsmdx=diff(sm)/1e-5;
max_dsmdx=max(dsmdx);
dsmdx_minatmid=min(dsmdx(x>=0.5 & x<=1.5));
verifyGreaterThan(tc,dsmdx,-1e9);
verifyLessThan(tc,dsmdx_minatmid,max_dsmdx/200);
verifyGreaterThan(tc,dsmdx_minatmid,max_dsmdx/400);

gfp_n=3;
gfp_k=0.6167;
bl_k=-8;
br_k=1;
bl_2mk=1;
br_2mk=10;
b1=@(x)(smoothstep_shifted_Rnoband(gfp_n,gfp_k,x)-smoothstep_shifted_Rnoband(gfp_n,gfp_k,bl_k))/...
  (smoothstep_shifted_Rnoband(gfp_n,gfp_k,br_k)-smoothstep_shifted_Rnoband(gfp_n,gfp_k,bl_k))...
  - ...
  ((1-smoothstep_shifted_Rnoband(gfp_n,gfp_k,bl_k))/...
  (smoothstep_shifted_Rnoband(gfp_n,gfp_k,br_k)-smoothstep_shifted_Rnoband(gfp_n,gfp_k,bl_k))-1)*...
  smoothstep_shifted_Rnoband(1,1.3,x);
b2=@(x)(smoothstep_shifted_Rnoband(gfp_n,2-gfp_k,x)-smoothstep_shifted_Rnoband(gfp_n,2-gfp_k,bl_2mk))/...
  (smoothstep_shifted_Rnoband(gfp_n,2-gfp_k,br_2mk)-smoothstep_shifted_Rnoband(gfp_n,2-gfp_k,bl_2mk));
sma=@(x)b1(x)+b2(x);
x=0:1e-5:2;
sm=arrayfun(sma,x);
b1mb2=arrayfun(@(x)b1(x)-b2(x),x);
dsmdx=diff(sm)/1e-5;
max_b1mb2=max(b1mb2);
max_dsmdx=max(dsmdx);
dsmdx_minatmid=min(dsmdx(x>=0.5 & x<=1.5));
verifyGreaterThan(tc,dsmdx,-1e9);
verifyLessThan(tc,dsmdx_minatmid,max_dsmdx/200);
verifyGreaterThan(tc,dsmdx_minatmid,max_dsmdx/300);

% general2 mode verifications: 
A=RNetwork(sym({'smr1_1';'smr2_1';'smr3_1'}),[0;1;0],'general2');
A.convert_to_process({'gfp_k','gfp_n'});
output_label={'smra_1_1';'smra_2_1';'smra_3_1';
  'smpa_1_1';'smpa_2_1';'smpa_3_1'};
A.output=str2sym(output_label);
A.output_label=output_label;
A.locked=true;
% softstep_mode='hill';
% softstep_mode='tanh';
softstep_mode='smoothstep_shifted';
flux_subbed=BioSystem.fullexp(A.flux,[containers.Map();A.flux]);
%%% smr vs smra
smra=flux_subbed('smra_1_1');
smra_num=matlabFunction(smra,'Vars',...
  {sym(A.inputvars.'),sym('smr1_1'),sym('smrst_1_1'),sym('smrd_1_1')});
smrd0_d=A.param('smrd_1_1');
%%%% different gfp_kn: what the mapped smra_num are
gfp_kn=[(2-0.15*4)/3+0.15;15];
verifyEqual(tc,smra_num(gfp_kn,0,0,0),0,'AbsTol',1e-8);
verifyLessThan(tc,smra_num(gfp_kn,gfp_kn(1)-0.15,0,0),0.02);
verifyGreaterThan(tc,smra_num(gfp_kn,gfp_kn(1)+0.15,0,0),0.96);
verifyLessThan(tc,smra_num(gfp_kn,2-gfp_kn(1)-0.15,0,0),1.04);
verifyGreaterThan(tc,smra_num(gfp_kn,2-gfp_kn(1)+0.15,0,0),1.95);
%%%% different gfp_kn: what the mapped smra_num are
if strcmp(softstep_mode,'hill')
  gfp_kn=[(2-0.15*4)/3+0.15;8];
elseif strcmp(softstep_mode,'tanh')
  gfp_kn=[(2-0.15*4)/3+0.15;6];
else
  gfp_kn=[0.6167;3];
end
verifyEqual(tc,smra_num(gfp_kn,0,0,0),0,'AbsTol',1e-3);
verifyEqual(tc,smra_num(gfp_kn,1,0,0),1,'AbsTol',2e-2);
verifyEqual(tc,smra_num(gfp_kn,2,0,0),2,'AbsTol',3e-3);
k_true=fzero(@(s)smra_num(gfp_kn,s,0,0)-0.5,[0,1]);
verifyLessThan(tc,smra_num(gfp_kn,k_true-0.16,0,0),0.22);
verifyGreaterThan(tc,smra_num(gfp_kn,k_true+0.15,0,0),0.78);
k_true=fzero(@(s)smra_num(gfp_kn,s,0,0)-1.5,[1,2]);
verifyLessThan(tc,smra_num(gfp_kn,k_true-0.15,0,0),1.22);
verifyGreaterThan(tc,smra_num(gfp_kn,k_true+0.15,0,0),1.78);
if strcmp(softstep_mode,'smoothstep_shifted')
  % ensure that smra is truly discrete when the continuous topo is discrete
  verifyEqual(tc,smra_num(gfp_kn,0,0,smrd0_d),0);
  verifyEqual(tc,smra_num(gfp_kn,1,0,smrd0_d),1);
  verifyEqual(tc,smra_num(gfp_kn,2,0,smrd0_d),2);
end
%%% fv1 rate at different smr
gfp_k=gfp_kn(1);
fv1_num=matlabFunction(flux_subbed('fv1'),'Vars',...
  {sym('krf1'),str2sym({'smr1_1';'smr2_1';'smr3_1'}),...
  str2sym({'smrd_1_1';'smrd_2_1';'smrd_3_1'; ...
  'smrst_1_1';'smrst_2_1';'smrst_3_1'}),...
  str2sym(A.statevars.'),sym(A.inputvars.')});
k_true=fzero(@(s)smra_num(gfp_kn,s,0,0)-0.5,[0,1]);
verifyEqual(tc,arrayfun( @(x)fv1_num(1,[k_true-0.15;0;0],zeros(6,1),[x;0;0],gfp_kn) , [0,1,4] ),...
  ones(1,3),'RelTol',0.26);
verifyEqual(tc,arrayfun( @(x)fv1_num(1,[k_true+0.2;0;0],zeros(6,1),[x;0;0],gfp_kn) , [0,1,4] ),...
  [0,1,4],'RelTol',0.25,'AbsTol',0.25);
k_true=fzero(@(s)smra_num(gfp_kn,s,0,0)-1.5,[1,2]);
verifyEqual(tc,arrayfun( @(x)fv1_num(1,[k_true-0.2;0;0],zeros(6,1),[x;0;0],gfp_kn) , [0,1,4] ),...
  [0,1,4],'RelTol',0.32,'AbsTol',5e-3);
verifyEqual(tc,arrayfun( @(x)fv1_num(1,[k_true+0.15;0;0],zeros(6,1),[x;0;0],gfp_kn) , [0,1,4] ),...
  [0,1,4^2],'RelTol',0.25,'AbsTol',1e-12);
if strcmp(softstep_mode,'smoothstep_shifted')
  % ensure discretization matches the true discrete topology dynamics. 
  verifyEqual(tc,fv1_num(1,[0;0;0],ones(6,1)*smrd0_d,[0;0;0],gfp_kn),1);
  verifyEqual(tc,fv1_num(1,[0;0;0],ones(6,1)*smrd0_d,[2;0;0],gfp_kn),1);
  verifyEqual(tc,fv1_num(1,[1;0;0],ones(6,1)*smrd0_d,[0;0;0],gfp_kn),0);
  verifyEqual(tc,fv1_num(1,[1;0;0],ones(6,1)*smrd0_d,[2;0;0],gfp_kn),2);
  verifyEqual(tc,fv1_num(1,[2;0;0],ones(6,1)*smrd0_d,[0;0;0],gfp_kn),0);
  verifyEqual(tc,fv1_num(1,[2;0;0],ones(6,1)*smrd0_d,[2;0;0],gfp_kn),4,'RelTol',1e-15);
  verifyNotEqual(tc,fv1_num(1,[1.05;0;0],ones(6,1)*smrd0_d,[0;0;0],gfp_kn),0);
  verifyEqual(tc,fv1_num(1,[1.05;0;0],ones(6,1)*smrd0_d,[0;0;0],gfp_kn),0,'AbsTol',5e-3);
end
%%% fvrx_1 rate at different x
fvr1_1_num=matlabFunction(flux_subbed('fvr1_1'),'Vars',...
  {sym('krf1'),str2sym({'smr1_1';'smr2_1';'smr3_1'}),...
  str2sym({'smrd_1_1';'smrd_2_1';'smrd_3_1'; ...
  'smrst_1_1';'smrst_2_1';'smrst_3_1'}),...
  str2sym(A.statevars.'),sym(A.inputvars.')});
fvr2_1_num=matlabFunction(flux_subbed('fvr2_1'),'Vars',...
  {sym('krf1'),str2sym({'smr1_1';'smr2_1';'smr3_1'}),...
  str2sym({'smrd_1_1';'smrd_2_1';'smrd_3_1'; ...
  'smrst_1_1';'smrst_2_1';'smrst_3_1'}),...
  str2sym(A.statevars.'),sym(A.inputvars.')});
verifyEqual(tc,fvr2_1_num(1,[gfp_k-0.15;1;1],zeros(6,1),[0;1;1],gfp_kn),...
  fv1_num(1,[gfp_k-0.15;1;1],zeros(6,1),[0;1;1],gfp_kn),...
  'RelTol',2e-2,'AbsTol',2e-2);
verifyEqual(tc,fvr2_1_num(1,[gfp_k-0.15;1;1],zeros(6,1),[0;1;1],gfp_kn),1,'RelTol',0.5);
verifyEqual(tc,fvr1_1_num(1,[gfp_k-0.15;1;1],zeros(6,1),[0;1;1],gfp_kn),0);
%%% bounded on second derivative
fv1_dxdx_num=matlabFunction(diff(flux_subbed('fv1'),sym('x1'),2),'Vars',...
  {sym('krf1'),str2sym({'smr1_1';'smr2_1';'smr3_1'}),...
  str2sym({'smrd_1_1';'smrd_2_1';'smrd_3_1'; ...
  'smrst_1_1';'smrst_2_1';'smrst_3_1'}),...
  str2sym(A.statevars.'),sym(A.inputvars.')});
verifyLessThan(tc,abs(fv1_dxdx_num(1,[1.001;0;0],zeros(6,1),[0;0;0],gfp_kn)),1.25e-2);

% verify that stoichoimetric matrix constraints are satisfied
verifyEqual(tc,RNetwork.sm_rules_satisfied([1,1],[-1,1]),false);
verifyEqual(tc,RNetwork.sm_rules_satisfied([0,1],[-1,1]),true);

% verify that sm matrix that satisfy GK condition in paper also satisfy
% condition coded here. 
%%% all GK's conditions
sm=[0 -1 -1 1; 1 0 1 -1];
smr=zeros(size(sm));
smr(1,2)=1;
sm_prod=sm;
sm_prod(1,2)=0;
smr(2,3)=1;
sm_prod(2,3)=2;
[l,l_r,l_c,l_q,qT,qTS1to2]=RNetwork.sm_gk_satisfied(smr,sm_prod,2);
verifyEqual(tc,qT(1),qT(2),'AbsTol',1e-12);
verifyEqual(tc,qTS1to2,qT*sm(:,1:2));
verifyTrue(tc,l);
%%% without qTS1to2 having opposite signs AND qTS1to2 both having negative
%%% signs
smr_new=smr;
smr_new(1,1)=2;
sm_prod_new=smr_new-sm_prod;
[l,l_r,l_c,l_q,qT,qTS1to2]=RNetwork.sm_gk_satisfied(smr_new,sm_prod,2);
verifyFalse(tc,l_q);
verifyLessThan(tc,qTS1to2,0); 
%%% when m>2, only two of the columns need to equal at 2:end
[l,l_r,l_c,l_q,qT]=RNetwork.sm_gk_satisfied(smr(:,[1,3,2,4]),sm_prod(:,[1,3,2,4]),3);
verifyTrue(tc,l);
%%% only l_q satisfied
sm=[0 0 0 0 1 -1  0;
    1 0 0 0 0  0 -1;
    0 1 0 0 0  0 -1;
    0 0 1 -1 0 0  0];
[l,l_r,l_c,l_q,qT]=RNetwork.sm_gk_satisfied(zeros(size(sm)),sm,2);
verifyTrue(tc,l_q);
verifyFalse(tc,l);
sm=[0 0 0 0 1 -1  0;
    1 0 0 0 0  0 -1;
    0 1 0 1 0  0 -1;
    0 0 1 0 0  0  0];
[l,l_r,l_c,l_q,qT]=RNetwork.sm_gk_satisfied(zeros(size(sm)),sm,2);
verifyFalse(tc,l_q);

% see if sm_gk_gen generated stoichoimetric matrices is as expected
%%% customization section
n_sv=6;
n_rxns=10;
%%% 
sm_react0=nan(n_sv,n_rxns+n_sv);
sm_subm=[ones(1,n_sv);zeros(n_sv/2-1,n_sv)];
sm_react0(1:n_sv/2,(end-n_sv+1):end)=sm_subm; 
sm_prod0=nan(n_sv,n_rxns+n_sv);
sm_prod0(1:n_sv/2,(end-n_sv+1):end)=sm_subm;
%%% both-bimolecular mode
rng_seed=2;
rng(rng_seed);
[smr,sm_prod]=RNetwork.sm_gk_gen(n_sv,n_rxns+n_sv,4,sm_react0,sm_prod0,'both-bimolecular');
%%% subm maintained?
verifyEqual(tc,smr(1:n_sv/2,end-n_sv+1:end),sm_subm); 
verifyEqual(tc,sm_prod(1:n_sv/2,end-n_sv+1:end),sm_subm); 
%%% bimolecular constraint satisfied
for c=1:n_rxns
  verifyLessThanOrEqual(tc,sum(smr(smr(:,c)~=0,c)),2);
  verifyLessThanOrEqual(tc,sum(sm_prod(sm_prod(:,c)~=0,c)),2);
end
%%% sm_gk_satisfied
l=RNetwork.sm_gk_satisfied(smr,sm_prod,4);
verifyTrue(tc,l);
%%% both mode
rng(rng_seed);
[smr,sm_prod,iter_b]=RNetwork.sm_gk_gen(n_sv,n_rxns+n_sv,4,sm_react0,sm_prod0,'both');
%%% qT only mode
rng(rng_seed);
[smr,sm_prod,iter_q]=RNetwork.sm_gk_gen(n_sv,n_rxns+n_sv,4,sm_react0,sm_prod0,'qT only');
[l,lr,lc,lq,qT]=RNetwork.sm_gk_satisfied(smr,sm_prod,4);
verifyTrue(tc,lq);
verifyFalse(tc,l);
verifyGreaterThanOrEqual(tc,iter_b,iter_q);
%%% none mode
rng(rng_seed);
[smr,sm_prod,iter_n]=RNetwork.sm_gk_gen(n_sv,n_rxns+n_sv,4,sm_react0,sm_prod0,'none:+');
[l,lr,lc,lq,qT]=RNetwork.sm_gk_satisfied(smr,sm_prod,4);
verifyFalse(tc,lq);
verifyEqual(tc,iter_n,1);
end
```
