## Supplemental Data 1 for "Principles underlying implementation of *nearly*-homeostatic biological networks": BioSystemTests_Homeostasis.m

```
%%  Unit testing framework for BioSystem class
% It will ensure that major components of BioSystem class is fully
% functional under implementation changes. A method is considered a major component
% if it is user intended OR it is a major function within a hiararchy.
classdef BioSystemTests_Homeostasis < matlab.unittest.TestCase
  properties
    BioSys
    exp_sol
  end
  
  methods(TestMethodSetup)
    function setup(tc) % run for each test
      sys={'x1','f3+b*x2';'x2','c*x1+d*x2'};
      flux={'f1','a*x1';'f2','f1';'f3','f2';'o1','x1'};
      param={'a',-5;'b',1;'c',4;'d',-2};
      x0_map={'x1',1;'x2',2};
      tc.BioSys=BioSystem(cell2map(sys),cell2map(flux),...
        cell2map(param),cell2map(x0_map));
      tc.BioSys.output='o1';
      tc.BioSys.output_label={'o1'};
      tc.exp_sol=@(t)3/5.*[1;4]*exp(-t)-2/5.*[-1;1]*exp(-6.*t);%t must be row vector
    end
  end
  
  methods(Test)
    function output_sensitivity(tc)
      tc.BioSys.sys('x1')=tc.BioSys.sys('x1')+'u';
      tc.BioSys.flux('o1')='x1';
      tc.BioSys.output='o1';
      sens=tc.BioSys.output_sensitivity;
      syms a b c d
      sens_exp=-[1,0]*[d,-c;-b,a].'*[1;0];
      tc.verifyEqual(sens,sens_exp,'RelTol',0.0001,'AbsTol',0.0001);
      
      tc.BioSys.sys.remove('x2');
      tc.BioSys.sys('x1')=2*sym('x1')+'u';
      sens=tc.BioSys.output_sensitivity;
      tc.verifyEqual(sym(-1),sens);
      [sens,DxH_map,DxA_map,DuA_map]=tc.BioSys.output_sensitivity(true);
      sens=subs(sens,str2sym(DxH_map.keys),cell2sym(DxH_map.values));
      sens=subs(sens,str2sym(DxA_map.keys),cell2sym(DxA_map.values));
      sens=subs(sens,str2sym(DuA_map.keys),cell2sym(DuA_map.values));
      tc.verifyEqual(sym(-1),simplify(sens));
    end
    output_numrelsensitivity(tc);
    output_sensitivity_nolaplaceexp(tc);

    function find_homeostasis(tc)
      %setting up BioSystem object
      dx={'x','a1*u-b1*x-k*x*y';'y','a2*u-k*x*y'};
      flux={'o1','y'};
      params={'a1',1;'a2',1';'b1',1;'k',1};
      x0_map={'x',1;'y',1};
      A=BioSystem(cell2map(dx),cell2map(flux),...
        cell2map(params),cell2map(x0_map));
      A.output='o1';
      %finding parameter region of homeostasis
      param_range={'a1',[1,11];'a2',[1,11];'b1',[1,11];'k',[1,11]};
      find_homeostasis(A,[0.5,0.6,0.7].',cell2map(param_range),10,[1000,2000],'test1');
      t=readtable('test1.csv');
      find_homeostasis(A,[0.5,0.6,0.7].',unique(t(:,{'a1','a2','b1','k'})),...
        10,[1000,2000],'test2');
      s=readtable('test2.csv');
      t=sortrows(t,{'a1','a2','b1','k','u'});
      s=sortrows(s,{'a1','a2','b1','k','u'});
      max_u=max(t.u);
      for j=1:height(t)
        if t.a1(j)>t.a2(j) && t.u(j)~=max_u && abs(t.o1(j))>1e-2 && t.fluct_o1(j)<1e-4
          tc.verifyEqual(t{j,'y'},t{j,'o1'},'RelTol',0.0001,'AbsTol',0.0001);
          tc.verifyEqual(t{j,'sens_o1'},s{j,'sens_o1'},'RelTol',0.0001,'AbsTol',0.0001);
          tc.verifyEqual(t{j,'sens_o1'}./t{j,'o1'},0,'RelTol',0.0001,'AbsTol',0.0001);
        end
      end
      
      %Feature addition: optional starting value table
      start_x0=table(ones(10,1),ones(10,1),'variableNames',{'x','y'});
      find_homeostasis(A,[0.5,0.6,0.7].',cell2map(param_range),10,[1000,2000],'test3',start_x0);
      
      %Feature addition: optional max_runtime
      start_x0=table(ones(10,1),ones(10,1),'variableNames',{'x','y'});
      find_homeostasis(A,[0.5,0.6,0.7].',cell2map(param_range),10,[1000,2000],'test4',start_x0,1);
      
      %deleting all written files
      delete('test1.csv','test2.csv','test3.csv','test4.csv');
    end

    function find_homeostasis2 (tc) %testing theoretical columns
      dx={'x','-x-d^2';'y','-y^3-d'};
      flux={'o1','x'};params={'dummy',0};x0_map={'x',1;'y',1};
      A=BioSystem(cell2map(dx),cell2map(flux),cell2map(params),cell2map(x0_map));
      A.output={'o1';'o1'};
      
      param_range={'dummy',[1,11]};
      find_homeostasis(A,[0.5,0.6,0.7].',cell2map(param_range),10,[1000,2000],'test1');
      t=readtable('test1.csv');
      
      sens_exp=[-2*0.5-0.1;-2*0.6-0.1;NaN]; t_sens_exp=sens_exp+0.1;
      t_sens_j_exp=[zeros(3,1),-12.*t{1:3,'y'}.*[0.5;0.6;0.7],-6*t{1:3,'y'}.^2]; 
      t_sens_j_exp(3,:)=NaN;
      det_exp=[-1*-3*nthroot(-0.5,3)^2;-1*-3*nthroot(-0.6,3)^2;-1*-3*nthroot(-0.7,3)^2];
      tc.verifyEqual(t{:,'sens_o1'},repmat(sens_exp,10,1),...
        'RelTol',0.0001,'AbsTol',0.0001);
      tc.verifyEqual(t{:,'sens_o2'},repmat(sens_exp,10,1),...
        'RelTol',0.0001,'AbsTol',0.0001);
      tc.verifyEqual(t{:,'t_sens_o1'},repmat(t_sens_exp,10,1),...
        'RelTol',0.0001,'AbsTol',0.0001);
      tc.verifyEqual(t{:,'t_sens_o2'},repmat(t_sens_exp,10,1),...
        'RelTol',0.0001,'AbsTol',0.0001);
      tc.verifyEqual(t{:,strcat({'psi_o1'},{'_x','_y','_d'})},...
        repmat(t_sens_j_exp,10,1),'RelTol',0.0001,'AbsTol',0.0001);
      tc.verifyEqual(t{:,strcat({'psi_o2'},{'_x','_y','_d'})},...
        repmat(t_sens_j_exp,10,1),'RelTol',0.0001,'AbsTol',0.0001);
      tc.verifyEqual(t{:,'det'},repmat(det_exp,10,1),...
        'RelTol',0.0001,'AbsTol',0.0001);
      tc.verifyEqual(t{:,'fluct_o1'},t{:,'fluct_x'},...
        'RelTol',0.0001,'AbsTol',0.0001);
      tc.verifyEqual(t{:,{'eig_1','eig_2'}},...
        repmat([det_exp/-1,[-1;-1;-1]],10,1),...
        'RelTol',0.0001,'AbsTol',0.0001);
      
      delete('test1.csv');
    end

    function hillfast_test(tc)
      verifyEqual(tc,hill_fast(-1, 0,2),0);
      verifyEqual(tc,hill_fast(0,  0,2),0);
      verifyEqual(tc,hill_fast(0.5,0,2),0.25);
      verifyEqual(tc,hill_fast(2,  0,2),1);
      verifyEqual(tc,hill_fast(2.5,0,2),1);
    end

    function compute_invhomeosscurves_test(~)
      rng(1,'twister');
      %% setting up system: x1 -> x2 -| x1 zeroth order degradation based leaky-integral controller
      action_params={'none','down';'up','none'};
      deg_params={'linear','normal'};
      A=GeneticNetwork(action_params,deg_params);
      A.output=sym('x1');
      A.output_label={'x1'};
      A.int_tol=-11;
      A.sys('x2')=A.sys('x2')-str2sym('epsilon*Vd1/dKd1*x2');
      A.prange('epsilon')=[0,1];
      A.param('epsilon')=0;
      take_limit(A,{'n','dn'},{1,1},true);
      A.sys('x1')=subs(A.sys('x1'),sym('V1'),str2sym('V1*(1-u2)'));
      A.flux('fH1_2')=subs(A.flux('fH1_2'),sym('x2'),...
        str2sym('Kiaa*(x2+x2g)/(Kaia*u+Kiaa)'));
      A.prange('x2g')=[0,1];
      A.param('x2g')=1;
      A.prange('Kiaa')=[0,10];
      A.param('Kiaa')=1;
      A.prange('Kaia')=[0,10];
      A.param('Kaia')=1;
      A.locked=true;
      u=[0,0;0,0.5];
      pars_noleak=setdiff(A.param.keys,{'epsilon','dKd2'});
      while true
        A.param('epsilon')=0;
        A.param('dKd2')=0;
        get_feasible(A,{'V2>Vd2';'V2*b2_1<Vd2'},pars_noleak);
        [x_A,x_A_f]=find_dose_response(A,u,[1e4,1e4+1e3],5);
        if ~any(isnan(x_A),'all') && all(abs(x_A_f./x_A)<=1e-3,'all')
          % at upper bound of 1e-2, up to 10% of rs_AL>0.01
          A.param('epsilon')=1e-3;
          A.param('dKd2')=1e-3;
          [x_A,x_A_f]=find_dose_response(A,u,[1e4,1e4+1e3],5);
          if ~any(isnan(x_A),'all') && all(abs(x_A_f./x_A)<=1e-3,'all')
            break;
          end
        end
      end
      
      %% testing of compute_invhomeo
      prod_rates=cell2map({'x1',str2sym('H1_1*H1_2*V1*(1-u2)');
        'x2',str2sym('H2_1*H2_2*V2')});
      deg_rates=cell2map({'x1',str2sym('D1*Vd1');
        'x2',str2sym('D2*Vd2+epsilon*Vd1/dKd1*x2')});
      multipliers=cell2map({'x1',str2sym('1/(1-u2)');'x2',str2sym('1')});
      plot_lims=cell2map({'x1',str2sym('[0,1.25*x1]');
        'x2',str2sym('[0,1.25*x2]');
        'u',str2sym('[-Kiaa/Kaia/4,Kiaa/Kaia]');
        'u2',str2sym('[-0.25,1.5*u2]')});
      figure;
      compute_invhomeo(A,x_A.',u.',...
        [false,true,false,true; true(1,3),false(1,1)],...
        [false,true,false,true; true(1,3),false(1,1)],...
        prod_rates,deg_rates,multipliers,plot_lims,false);
      
      %% testing of compute_sscurves and compute_invhomeo_lvl2 when two parameters differ by b1_2
      A.locked=false;
      A.xrange('x3')=[0,inf]; % see if more than 2 panels for the invh plots
      A.x0_map('x3')=A.x0_map('x1');
      A.sys('x3')=subs(A.sys('x1'),sym('D1'),str2sym('x3/dKd1'));
      A.xrange('x4')=[0,inf]; % implement case where dim(sub_sys)>dim(sub_sys_range.keys)
      A.x0_map('x4')=A.x0_map('x1');
      A.sys('x4')=subs(A.sys('x1'),sym('D1'),str2sym('x4/dKd1'));
      p1=A.param;
      p2=[containers.Map();p1];
      p2('b1_2')=p2('b1_2')*10;
      prod_rates('x3')=sym(0);
      deg_rates('x3')=sym(0);
      multipliers('x3')=multipliers('x1');
      plot_lims('x3')=plot_lims('x1');
      prod_rates('x4')=sym(0);
      deg_rates('x4')=sym(0);
      multipliers('x4')=multipliers('x1');
      plot_lims('x4')=plot_lims('x1');
      % converting prod/deg_rates, multipliers to numeric forms
      A_paramsk=A.param.keys;
      A_sv=A.statevars;
      A_u=A.inputvars;
      prod_rates_num=BioSystem.fullexp(prod_rates,A.flux);
      deg_rates_num=BioSystem.fullexp(deg_rates,A.flux);
      multipliers_num=BioSystem.fullexp(multipliers,A.flux);
      for k=A.statevars
        key=k{:};
        prod_rates_num(key)=matlabFunction(prod_rates_num(key),...
          'vars',{sym(A_paramsk).',sym(A_sv).',sym(A_u).'});
        deg_rates_num(key)=matlabFunction(deg_rates_num(key),...
          'vars',{sym(A_paramsk).',sym(A_sv).',sym(A_u).'});
        multipliers_num(key)=matlabFunction(multipliers_num(key),...
          'vars',{sym(A_paramsk).',sym(A_sv).',sym(A_u).'});
      end
      % testing x1 subsystem; 
      figNC=figure;
      figIH=figure;
      x2_pltrange=[1e-3,1e2*A.param('Kd1_2')];
      x1_pltrange=[1e-2,1e1];
      compute_sscurves(A,figNC,figIH,{'x1','x3'},'x1',...
        'x2',x2_pltrange,x1_pltrange,[1e3,2e3],...
        [repmat(cell2mat(p1.values.'),1,2),repmat(cell2mat(p2.values.'),1,2)],...
        [ones(1,4);1:4;ones(2,4)],repmat([[0;0],[0;0.5]],1,2),2,false,...
        prod_rates_num,deg_rates_num,multipliers_num,...
        x2_pltrange(1)+[0.25,0.75]*diff(x2_pltrange));
      % testing swapped axis
      figNC=figure;
      compute_sscurves(A,figNC,figIH,{'x1','x3'},'x1',...
        'x2',x2_pltrange,x1_pltrange,[1e3,2e3],...
        [repmat(cell2mat(p1.values.'),1,2),repmat(cell2mat(p2.values.'),1,2)],...
        [ones(1,4);[1:2,nan(1,2)];ones(2,4)],repmat([[0;0],[0;0.5]],1,2),2,true,...
        prod_rates_num,deg_rates_num,multipliers_num,...
        x2_pltrange(1)+[0.25,0.75]*diff(x2_pltrange));
      
      % testing compute_invhomeo_lvl2
      fprf='compute_invhomeo_lvl2-sscurve';
      sub_sys_range=containers.Map({'x1','x2','x3'},{x1_pltrange,x2_pltrange,x1_pltrange});
      compute_invhomeo_lvl2(A,fprf,A.statevars,sub_sys_range,[1e16,2e16],...
        [repmat(cell2mat(p1.values.'),1,2),repmat(cell2mat(p2.values.'),1,2)],...
        [ones(1,4);1:4;ones(2,4)],repmat([[0;0],[0;0.5]],1,2),2);
    end
    classify_findnhp_test(tc);
    utilities_test(tc);
  end
end
```
