## Supplemental Data 1 for "Principles underlying implementation of *nearly*-homeostatic biological networks": classify_findnhp_test.m

```
function classify_findnhp_test(tc)
% Testing classify_findnhp

%% generating summary table along with expected categories
iter=[repmat(1,1,3),repmat(2,1,3),repmat(3,1,3),repmat(4,1,3)].';
stage=[repmat([1,2,2],1,3),repmat(1,1,3)].';
iter_time=[1,2,ones(1,10)].';
search_success=[ones(1,2),zeros(1,10)].';
A_aug_x0_trivialss=[zeros(1,2),ones(1,3),zeros(1,7)].';
allset=[ones(1,6),zeros(1,6)].';
data=table(iter,stage,iter_time,search_success,A_aug_x0_trivialss,allset,...
  'VariableNames',{'iter','stage','iter_time','search_success','A_aug_x0_trivialss','allset'});
out_c_exp=[repmat({'search_success + allset'},1,2),...
  repmat({'A_aug_x0_trivialss + allset'},1,3),...
  {'allset'},repmat({' '},1,6)].';
summary_exp=table({' ','A_aug_x0_trivialss + allset','allset','search_success + allset'}.',...
  [6/13,3/13,1/13,(2+1)/13].',[6/12,3/12,1/12,2/12].',...
  'VariableNames',{'category','p_weighted','p_noweight'});
summary_stg_exp=table([1;2],[4;3],'VariableNames',{'stage','num_iters'});

%% classify_nhp and compare with expected results
figure;
ax=gca;
[out_c,summary_cats,summary_stg]=BioSystem.classify_findnhp(data,'iter_time',ax);
verifyEqual(tc,out_c,out_c_exp);
verifyEqual(tc,summary_cats,summary_exp);
verifyEqual(tc,summary_stg,summary_stg_exp);
end
```
