## Supplemental Data 1 for "Principles underlying implementation of *nearly*-homeostatic biological networks": output_numrelsensitivity.m

```
function output_numrelsensitivity(tc)
rng(1);

min_sn=nan;
syms min_sn(x,y)

tc.BioSys.sys('x1')=tc.BioSys.sys('x1')+'u';
tc.BioSys.flux('o1')='x1';
tc.BioSys.output_label={'output'};
tc.BioSys.output='o1';
A=tc.BioSys;
sens_denom=str2sym('5'); % denominator of the sensitivity function
sens_denom_num=matlabFunction(sens_denom,'Vars',...
   {sym(A.param.keys.'),sym(A.statevars.'),sym(A.inputvars.')});

%% verify computation of rsenso1
A.locked=true;
u=1;
inputs=[u;u+1e-4];
while true
  get_feasible(A,@(p)true,A.prange);
  [x,x_fluct,o1]=find_dose_response(A,inputs,[1000,2000]);
  if min(max(abs(x_fluct./x),[],'all'),max(x_fluct,[],'all'))<=1e-8
    break;
  end
end
[numf_relsens,Ac]=output_numrelsensitivity(A,sens_denom_num);
relsens_theor=numf_relsens(cell2mat(A.param.values.'),x(1,:).',u);
numf_hindct=matlabFunction(A.output_sensitivity,'Vars',...
  {sym(A.param.keys.'),sym(A.statevars.'),sym(A.inputvars.')});
numf_relsens2=@(p,x,u)numf_hindct(p,x,u)/det(A.jacobn_full(p,x,u))/5;
relsens_theor2=numf_relsens2(cell2mat(A.param.values.'),x(1,:).',u);
relsens_emp=(o1(2)-o1(1))/1e-4/5;
verifyGreaterThan(tc,1e-8,abs(relsens_theor-relsens_emp)/abs(relsens_emp));
verifyGreaterThan(tc,1e-8,abs(relsens_theor-relsens_theor2)/abs(relsens_theor2))
end
```
