## Supplemental Data 1 for "Principles underlying implementation of *nearly*-homeostatic biological networks": output_sensitivity_nolaplaceexp.m

```
function output_sensitivity_nolaplaceexp(tc)
rng(1);

min_sn=nan;
syms min_sn(x,y)

A=tc.BioSys.copy;
A.sys('x1')=A.sys('x1')+'u';
A.flux('o1')='x1';
A.output=str2sym({'o1'});
A.output_label={'o_o1'};
Asimp=A.copy;
[sens,DxH,DxA,DuA]=output_sensitivity_nolaplaceexp_fullexp(A);
A.flux('DxH')=DxH;
A.flux('DxA')=DxA;
A.flux('DuA')=DuA;
A.flux('sens')=sens;
A.output=str2sym({'o1';'sens'});
A.output_label={'o_o1';'o_sens'};

%% verify computation of rsenso1
A.num_sim_only=true;
A.locked=true;
u=1;
inputs=[u;u+1e-4];
while true
  get_feasible(A,@(p)true,A.prange);
  [x,x_fluct,o1]=find_dose_response(A,inputs,[1000,2000]);
  if min(max(abs(x_fluct./x),[],'all'),max(x_fluct,[],'all'))<=1e-8
    break;
  end
end
numf_output_full=A.numf_output_full;
numf_relsens=@(p,x,u)extract_farray(numf_output_full(p,x,u),2);
relsens_theor=numf_relsens(cell2mat(A.param.values.'),x(1,:).',u);
relsens_emp=(o1(2)-o1(1))/1e-4;
verifyGreaterThan(tc,1e-8,abs(relsens_theor-relsens_emp)/abs(relsens_emp));

%% verify equivalence between fullexp and not fullexp
[sens,DxH,DxA,DuA,derivatives]=output_sensitivity_nolaplaceexp(Asimp);
Asimp.flux('DxH')=DxH;
Asimp.flux('DxA')=DxA;
Asimp.flux('DuA')=DuA;
Asimp.flux('sens')=sens;
Asimp.flux=[Asimp.flux;derivatives];
Asimp.output=str2sym({'o1';'sens'});
Asimp.output_label={'o_o1';'o_sens'};
Asimp.locked=true;
Asimp.ut=A.ut;
Asimp.param=[ Asimp.param; containers.Map2(A.param.keys,A.param.values) ];
Asimp.x0=A.x0;
verifyEqual( tc , Asimp.numf_output(Asimp.x0,Asimp.ut(0)) , A.numf_output(A.x0,A.ut(0)) );
end
```
