## Supplemental Data 1 for "Principles underlying implementation of *nearly*-homeostatic biological networks": utilities_test.m

```
function utilities_test(tc)

%% lm_f_fmincon_gen: ensuring that stop=1 is triggered under the right conditions
lm_f=lm_f_fmincon_gen(400,0.05);
verifyEqual( tc , lm_f([],struct('iteration',1,'fval',1000),'iter') , false);
verifyEqual( tc , lm_f([],struct('iteration',402,'fval',1000),'iter') , true);
verifyEqual( tc , lm_f([],struct('iteration',1,'fval',0.05),'iter') , false);
verifyEqual( tc , lm_f([],struct('iteration',402,'fval',0.052),'iter') , true);
verifyEqual(tc , lm_f([],struct('iteration',1,'fval',1000),'iter') , false);
verifyEqual(tc , lm_f([],struct('iteration',402,'fval',1020),'iter') , false);
verifyEqual(tc , lm_f([],struct('iteration',403,'fval',1001),'iter') ,true);
verifyEqual(tc , lm_f([],struct('iteration',1,'fval',1000),'iter') , false);
verifyEqual(tc , lm_f([],struct('iteration',402,'fval',100),'iter') , false);
verifyEqual(tc , lm_f([],struct('iteration',411,'fval',100),'iter') , false);
verifyEqual(tc , lm_f([],struct('iteration',803,'fval',100.05),'iter') , false);
verifyEqual(tc , lm_f([],struct('iteration',820,'fval',100.05),'iter') , true);
end
```
