## Supplemental Data 1 for "Principles underlying implementation of *nearly*-homeostatic biological networks": discretize_bf.m

```
function dtopo = discretize_bf(topo)
% A function that discretize a given topological parameter for brute force search.
% 
% INPUT
% topo - topological parameter
% OUTPUT
% dtopo - discretized topological parameter 

% topo_drefs=[0,0.75,1.5]; % corresponding to tanh based general form
topo_drefs=[0,1,2];
weights=[0.5,1,0.5]; % ensure all topo state given equal weighting
dtopo=extract_farray(topo_drefs,...
  find( weights.*abs(topo-topo_drefs)==min(weights.*abs(topo-topo_drefs)) , 1 ));
if isempty(dtopo) && isnan(topo)
  dtopo=nan;
end
end
```
