## Supplemental Data 1 for "Principles underlying implementation of *nearly*-homeostatic biological networks": gd_discretize.m

```
function gd_discretize(A_aug,par_opt,status_d)
% Modify A_aug to ensure that dobj_key_dval1/2/3 are modified to attract topo parameter to the
% desired discrete value
% 
% INPUT
% status_d - 0 means decrease to first possible discrete value; 1 means increase to first possible
%   discrete value

%% discretization by setting dobj_par_dval_
% topo_drefs=[0,0.75,1.5]; % corresponding to tanh based general form
topo_drefs=[0,1,2];
val=A_aug.x0_map(par_opt);
A_aug.param([par_opt,'_don'])=1;
if status_d==0 && val>topo_drefs(end)
  % do nothing
  1;
elseif status_d==0 && val>topo_drefs(2)
  A_aug.param(['dobj_',par_opt,'_dval_3'])=A_aug.param(['dobj_',par_opt,'_dval_1']);
elseif status_d==0 && val<topo_drefs(2)
  A_aug.param(['dobj_',par_opt,'_dval_2'])=A_aug.param(['dobj_',par_opt,'_dval_1'])-...
    abs(A_aug.param(['dobj_',par_opt,'_dval_2'])-A_aug.param(['dobj_',par_opt,'_dval_1']));
elseif status_d==1 && val<topo_drefs(1)
  % do nothing
  1;
elseif status_d==1 && val<topo_drefs(2)
  A_aug.param(['dobj_',par_opt,'_dval_1'])=A_aug.param(['dobj_',par_opt,'_dval_3']);
elseif status_d==1 && val>topo_drefs(2)
  A_aug.param(['dobj_',par_opt,'_dval_2'])=A_aug.param(['dobj_',par_opt,'_dval_3'])+...
    abs(A_aug.param(['dobj_',par_opt,'_dval_2'])-A_aug.param(['dobj_',par_opt,'_dval_3']));
end
end
```
