## Supplemental Data 1 for "Principles underlying implementation of *nearly*-homeostatic biological networks": gd_discretize_check.m

```
function [topos_d,l_d,topos_act_fdisc] = gd_discretize_check(A_aug,paropts_topo,topovs,topos_thres)
% Return discretize topology if the right criteria are met: mapped topology is within topo_thres
% of discrete topology AND every component still has a reaction with net degradation.
% Otherwise, return the original topology.
% INPUT
% A_aug - BioSystem object. Output must contain topo_act_name; state variable must contain
%  paropt_topo
% paropts_topo - cell array of characters for names of the topological parameters
% topovs - numeric vector
% topos_thres - cell array of numeric scalar that determines the threshold the calculated
%  topology must fall under to be discretized OR contains 'up' OR contains 'down'. If contains 'up', 
%  then tries to evaluate whether the ideal higher discrete value satisfy other constraints.
% OUTPUT
% topos_d - numeric vector of either discretized topology or original topology value
% l_d - logical vector whether topology is discretized
% topos_act_fdisc - numeric vector how far is current topo act from discrete topo_act; topo_act - topo_d

% getting necessary preliminary information
% topo_drefs=[0,0.75,1.5]; % corresponding to tanh based general form
topo_drefs=[0,1,2];
x0_map=[containers.Map;A_aug.x0_map];
topos_act_name=strcat(paropts_topo,'_act');
[~,topoact_in_out_idx]=ismember(topos_act_name,A_aug.output_label);

% compute candidate discrete topology based on topo_thres
x0_map=[ x0_map ; containers.Map(paropts_topo,topovs) ];
out_Aaug=A_aug.numf_output(cell2mat(x0_map.values).',A_aug.ut(0));
topos_act=extract_farray(out_Aaug,topoact_in_out_idx);
topos_dc=nan(size(topos_act));
for k=1:numel(paropts_topo)
  if isa(topos_thres{k},'numeric')
    seldref=topo_drefs( find(abs(topos_act(k)-topo_drefs)<=topos_thres{k},1) );
    if isempty(seldref)
      topos_dc(k)=topovs(k);
    else
      topos_dc(k)=seldref;
    end
  elseif ~isempty(regexp(topos_thres{k},'up','once'))
    drefs_idx=find(topovs(k)<=topo_drefs,1,'first');
    if ~isempty(drefs_idx)
      topos_dc(k)=topo_drefs(drefs_idx);
    else
      topos_dc(k)=max(topo_drefs);
    end
  elseif ~isempty(regexp(topos_thres{k},'down','once'))
    drefs_idx=find(topovs(k)>=topo_drefs,1,'last');
    if ~isempty(drefs_idx)
      topos_dc(k)=topo_drefs(drefs_idx);
    else
      topos_dc(k)=min(topo_drefs);
    end
  end
end

% discretization of topology if all criteria are met
l_d=false(size(topovs));
topos_d=nan(size(topovs));
topos_act_fdisc=nan(size(topovs));
for k=1:numel(paropts_topo)
  if any(topos_dc(k)==topo_drefs)
    l_d(k)=true;
  else
    l_d(k)=false;
  end
  if l_d(k)
    topos_d(k)=topos_dc(k);
    topos_act_fdisc(k)=topos_act(k)-topos_d(k);
  else
    topos_d(k)=topovs(k);
    topos_act_fdisc(k)=0;
  end
end
end
```
