## Supplemental Data 1 for "Principles underlying implementation of *nearly*-homeostatic biological networks": gd_reset.m

```
function gd_reset(A_aug,pars_opt_topo)
% Reset A_aug parameters associated with pars_opt_topo
%
% INPUT
% pars_opt_topo - cell array of topological parameters where the associated parameters are reset. 

% topo_drefs=[0,0.75,1.5];
topo_drefs=[0,1,2];
for i=1:numel(pars_opt_topo)
  key=pars_opt_topo{i};
  A_aug.param([key,'_don'])=0;
  A_aug.param(['dobj_',key,'_dval_1'])=topo_drefs(1);
  A_aug.param(['dobj_',key,'_dval_2'])=topo_drefs(2);
  A_aug.param(['dobj_',key,'_dval_3'])=topo_drefs(3);
end
end
```
