## Supplemental Data 1 for "Principles underlying implementation of *nearly*-homeostatic biological networks": gen_proddegrates.m

```
function [sys_prod,numf_prod,sys_deg,numf_deg]=gen_proddegrates(obj)
% Generate production rates and degradation rates.
% INPUT
% obj - GeneticNetwork object
% OUTPUT
% sys_prod - container map of symbolic production rates. Each key is a separate state variable. 
% numf_prod - container map of numeric functions [(p,x) or (p,x,u)] of production rates.  
%  Each key is a separate state variable. 
% sys_deg - container map of symbolic degradation rates. Each key is a separate state variable. 
% numf_deg - container map of numeric functions [(p,x) or (p,x,u)] of degration rates. 
%  Each key is a separate state variable. 

%% computing required information
dim_x=obj.sys.Count;
lastn=str2double(regexprep(char(obj.sys(obj.statevars{1})),'^.*H\d+_(\d+)\*V.*$','$1'));
one2n=1:lastn;
one2n_char=arrayfun(@num2str,one2n,'UniformOutput',false);
sv=obj.statevars;
iv=obj.inputvars;
sys_prod=containers.Map;
sys_deg=containers.Map;
numf_prod=containers.Map;
numf_deg=containers.Map;

%% split up Hill function canomical form to generate symbolic production rate and degradation rate
for i=1:dim_x
  x=sv{i};
  i_char=regexprep(char(obj.sys(x)),'^.*H(\d+)_\d+\*V.*$','$1');
  sys_prod(x)=strjoin(strcat('H',i_char,'_',one2n_char),'*');
  sys_prod(x)=['V',i_char,'*',sys_prod(x)];
  sys_prod(x)=str2sym(sys_prod(x));
  sys_deg(x)=['Vd',i_char,' * D',i_char];
  sys_deg(x)=str2sym(sys_deg(x));
end

%% generate numeric form of production and degradation rates
sys_prod_full=BioSystem.fullexp(sys_prod,obj.flux);
sys_deg_full=BioSystem.fullexp(sys_deg,obj.flux);
for i=1:dim_x
  x=sv{i};
  if isempty(obj.inputvars)
    numf_prod(x)=matlabFunction(sys_prod_full(x),'Vars',{sym(obj.param.keys).',sym(sv).'});
    numf_deg(x)=matlabFunction(sys_deg_full(x),'Vars',{sym(obj.param.keys).',sym(sv).'});
  else
    numf_prod(x)=matlabFunction(sys_prod_full(x),'Vars',{sym(obj.param.keys).',sym(sv).',sym(iv).'});
    numf_deg(x)=matlabFunction(sys_deg_full(x),'Vars',{sym(obj.param.keys).',sym(sv).',sym(iv).'});
  end
end

%% ensuring GeneticNetwork's general form has not changed to make this method invalid
for i=1:dim_x
  x=sv{i};
  if ~isequal(sys_prod(x)-sys_deg(x)-obj.sys(x),sym(0))
    error('GeneticNetwork:gen_proddegrates:general_form',...
      'The general form has changed sufficiently to render this method invalid!');
  end
end
end
```
