## Supplemental Data 1 for "Principles underlying implementation of *nearly*-homeostatic biological networks": GeneticNetwork.m

```
%% Overview
% This class defines a subset of BioSystem that are only genetic
% networks of Hill function canonical form
% 
% $$ \frac{d x_i}{dt}=V_i \prod_{j=1}^{\dim(x)} H_{i,j}(x_j) - \textrm{Vd}_i D_i(x_i) $$
% 
% Initial conditions are randomly computed to be strictly positive.
%% Design
% Additional methods are added on top of BioSystem as required by findnhp and findnhp_fmincon. 
% 
% * *istrivialss*, *discretize_bf*, *gd_discretize*, *gd_reset*, *gd_discretize_check* 
%
% Additional methods are added on top of BioSystem to generate inverse homeostasis plots. 
% 
% * *gen_proddegrates* generates production rates and degradation rates of the object. 
% Both symbolic and numeric forms are generated. 
% 
classdef GeneticNetwork < BioSystem
  methods(Static=true)
    l=istrivialss(xss);
    dtopo=discretize_bf(topo);
    gd_reset(A_aug,pars_opt_topo); 
    gd_discretize(A_aug,par_opt,status_d);
    [topo_d,l_d,topoact_fdisc]=gd_discretize_check(A_aug,paropt_topo,topov,topo_thres);
    [topo_act,b1,b2,b1_nkbl,b1_nkbr,b2_nkbl,b2_nkbr]=gen_form(var,d,n,k,s);
  end
  methods %user intended methods
    [sys_prod,numf_prod,sys_deg,numf_deg]=gen_proddegrates(obj);
    function[obj]=GeneticNetwork(inflow_params,deg_params,param_range)
      %INPUT
      %action_params - {'<type>', '<type>'[,...];'<type>','<type>'[,...]}
      %   type=uplinear|up|down|unknown|none
      %   Each row represent regulation of one component by all other components.
      %deg_params - {'<type>', '<type>'[,...]}
      %   type=normal|linear|zeroorder|general
      %param_range - container map; {'<param1>',[<from>,<to>][;...]}; OPTIONAL
      %OUTPUT
      %an instance of the class, reflecting form set by action_params

      unknown_in_inflow=false;
      %input checking
      if(size(inflow_params,1)~=length(deg_params))
        error(['The number of rows of inflow_params must be the',...
          ' same as number of deg_params']);
      end
      heaviside_fast=nan;
      syms heaviside_fast(a1)

      %setting up sys and flux
      dx=containers.Map();
      flux=containers.Map();
      sk=sym('sk');
      sn=sym('sn');
      rp_to_full=struct('r','react','p','prod');
      for r=1:size(inflow_params,1)
        r_char=num2str(r);
        x=['x',r_char];
        V=sym(['V',r_char]);
        dx(x)=V;
        for c=1:size(inflow_params,2)
          c_char=num2str(c);
          H=['H',r_char,'_',c_char];fH=['fH',r_char,'_',c_char];
          b=sym(['b',r_char,'_',c_char]);
          flux(fH)=prod_action(inflow_params{r,c},r,c);
          if strcmp(inflow_params{r,c},'none')
            flux(H)=sym(fH);
          else
            flux(H)=(b+(1-b)*sym(fH));
            if strcmp(inflow_params{r,c},'unknown')
              unknown_in_inflow=true;
              if true
              % if false
                % smoothstep_shifted based topology parametrization
                topo_drefs=[0,1,2];
                s_char=['s',r_char,'_',c_char];
                s=str2sym(s_char);
                ptopo_don_sym=str2sym(['s',r_char,'_',c_char,'_don']);
                [topo_act,b1,b2,b1_nkbl,b1_nkbr,b2_nkbl,b2_nkbr]= ...
                  gen_form_sstep(char(s),ptopo_don_sym,sn,sk,s);
                b1_nkbl_name=[s_char,'_b1_nkbl'];
                flux(b1_nkbl_name)=b1_nkbl;
                b1_nkbr_name=[s_char,'_b1_nkbr'];
                flux(b1_nkbr_name)=b1_nkbr;
                b1_name=[s_char,'_b1'];
                flux(b1_name)=b1;
                b2_nkbl_name=[s_char,'_b2_nkbl'];
                flux(b2_nkbl_name)=b2_nkbl;
                b2_nkbr_name=[s_char,'_b2_nkbr'];
                flux(b2_nkbr_name)=b2_nkbr;
                b2_name=[s_char,'_b2'];
                flux(b2_name)=b2;
                topo_act_name=[s_char,'_act'];
                flux(topo_act_name)=topo_act;
              else
                % softstep tanh based topology parametrization
                % topo_drefs=[0,0.75,1.5];
                topo_drefs=[0,1,2];
                s_char=['s',r_char,'_',c_char];
                s=str2sym(s_char);
                ptopo_don_sym=str2sym(['s',r_char,'_',c_char,'_don']);
                b1_name=[s_char,'_b1'];
                flux(b1_name)=softstep(sn,sk,s)*heaviside_fast(ptopo_don_sym+1);
                b2_name=[s_char,'_b2'];
                flux(b2_name)=softstep(sn,topo_drefs(end)-sk,s)*heaviside_fast(ptopo_don_sym+1);
                topo_act_name=[s_char,'_act'];
                flux(topo_act_name)=str2sym([num2str(topo_drefs(2)),'*','(',b1_name,'+',b2_name,')']);
              end
            end
          end
          dx(x)=dx(x)*sym(H);
        end
        fD=['fD',r_char];D=['D',r_char];
        flux(fD)=deg_action(deg_params{r},r);
        flux(D)=sym(fD);
        dx(x)=dx(x)-sym(['Vd',r_char])*sym(D); 
      end
      %generate GeneticNetwork object
      obj@BioSystem(dx,flux,containers.Map,containers.Map,containers.Map);
      k_params=obj.inputvars;
      %parameter initiation
      if nargin==2
        param_range=containers.Map();
        for j=1:numel(k_params)
          if regexp(k_params{j},'^b')
            param_range(k_params{j})=[0,0.1];
          elseif regexp(k_params{j},'^V')
            param_range(k_params{j})=[0,10];
          elseif regexp(k_params{j},'^s\d+_\d+$')
            param_range(k_params{j})=[0,topo_drefs(end)];
          elseif regexp(k_params{j},'^s\d+_\d+_don$')
            param_range(k_params{j})=[1,1]; % discretizable to actual topology by default
          else
            param_range(k_params{j})=[0,10];
          end
        end
      elseif nargin==3
        param_range=ensure_form(param_range,'numeric');
        if ~all(ismember(param_range.keys,k_params))||...
            ~all(ismember(k_params,param_range.keys))
          error('GeneticNetwork:param_range',...
            'param_range keys must be the same as parameters');
        end
      end
      if unknown_in_inflow % maintain compatibility with code that assumes no unknown inflow params. 
        param_range('sn')=[3,3];
        param_range('sk')=[0.6167,0.6167];
      end
      obj.param=containers.Map(k_params,rand_param(prange_convert(param_range),1));
      obj.prange=param_range;
      %%%% 10x than absolute tolerance
      obj.xrange=containers.Map(obj.statevars,repmat({[-1.1e-4,Inf]},1,obj.sys.length));
      obj.x0=rand(length(obj.statevars),1)+0.1;
    end
  end
end
function[f]=prod_action(type,self,j)
%construct specific regulation function of one variable on another's
%production ONLY
%   production ONLY
%inputs:
%type - uplinear|up|down|unknown|none
%self - index of variable that is self
%j - index of variable that is the agent of action
%output:
%f - symbolic expression encoding a function of x
self=num2str(self);j=num2str(j);
x=sym(['x',j]); Kd=sym(['Kd',self,'_',j]); n=sym(['n',self,'_',j]); 
b1=sym(['s',self,'_',j,'_b1']); b2=sym(['s',self,'_',j,'_b2']);
if  strcmp(type,'unknown')
  % too flat region of f vs (s,x) is predicted to hinder gradient descent, so 
  % power shouldn't be too high and Km parameter for s shouldn't be too far apart.
  % smap2=1 only if smap=1
  f=(1-b1)*x^n/(x^n+Kd^n) + (b1 - b2) + b2*Kd^n/(x^n+Kd^n);
elseif strcmp(type,'uplinear')|| strcmp(type,'up') || strcmp(type,'down')
  f=action(type);
  f=f(x,Kd,n);
elseif strcmp(type,'none')
  f=1;
else
  error('wrong type of regulation');
end
end
function[f]=deg_action(type,self)
%construct specific regulation function of one variable on another's  degradation ONLY
%inputs:
%type - normal|linear|zeroorder
%self - index of variable that is self
%output:
%f - symbolic expression encoding a function of x
self=num2str(self);
x=sym(['x',self]);Kd=sym(['dKd',self]);n=sym(['dn',self]);
if strcmp(type,'linear')
  f=action('uplinear');
  f=f(x,Kd,n);
elseif strcmp(type,'normal')
  f=action('up');
  f=f(x,Kd,n);
elseif strcmp(type,'zeroorder')
  f=1;
else
  error('degradation type is not correctly specified!');
end
end
```
