## Supplemental Data 1 for "Principles underlying implementation of *nearly*-homeostatic biological networks": GeneticNetwork.html

Overview 

### Overview

This class defines a subset of BioSystem that are only genetic networks of Hill function canonical form

Initial conditions are randomly computed to be strictly positive.

#### Contents

- Design

#### Design

Additional methods are added on top of BioSystem as required by findnhp and findnhp\_fmincon.

- **istrivialss**, **discretize\_bf**, **gd\_reset**, **gd\_discretize\_check**

Additional methods are added on top of BioSystem to generate inverse homeostasis plots.

- **gen\_proddegrates** generates production rates and degradation rates of the object. Both symbolic and numeric forms are generated.

Published with MATLAB® R2022b
