## Supplemental Data 1 for "Principles underlying implementation of *nearly*-homeostatic biological networks": istrivialss.m

```
function l=istrivialss(xss)
% A function that returns a logical value indicating whether the supplied steady state is trivial
%
% Unit tested by gen_augsys4nearh_prsenso1min_novs_test
% 
% INPUTS
% xss - vector
% OUTPUT
% logical scalar
l=...
  xss(1)<1e-3 || ... % eliminating trivial steady state
  sqrt(sum(xss(2:end).^2))<1e-5 || ... % eliminating trivial steady state part II
  any(xss<-1e-5) || ... % to prevent ss from fluctuating near absolute boundary
  0;
end
```
