## Supplemental Data 1 for "Principles underlying implementation of *nearly*-homeostatic biological networks": f_regulate.m

```
function[f]=f_regulate(type,self,j)
%construct regulatory fractional function of one variable acting on another.
%inputs:
%type - uplinear|up|downlinear|down|unknown|none
%self - index of variable that is self
%j - index of variable that is the agent of action
%output:
%f - symbolic expression
charj=num2str(j); charself=num2str(self); 
charx=sym(['x',charself]); symx=sym(charx);
Kd=sym(['Kd',charself,'_',charj]);n=sym(['n',charself,'_',charj]);
dKd=sym(['dKd',charself,'_',charj]);dn=sym(['dn',charself,'_',charj]);
if strcmp(type,'unknown')
  f=sym(['fH',charself','_',charj,'(x',charself,')']);
elseif strcmp(type,'uplinear')|| strcmp(type,'up')
  f=action(type);
  f=f(1-symx,Kd,n);
elseif strcmp(type,'down')
  f=action('up');
  f=f(symx,dKd,dn);
elseif strcmp(type,'downlinear')
  f=action('uplinear');
  f=f(symx,dKd,dn);
elseif strcmp(type,'none')
  f=1;
else
  error('wrong type of regulation');
end
end
```
