## Supplemental Data 1 for "Principles underlying implementation of *nearly*-homeostatic biological networks": MNetwork.m

```
% Metabolic Networks
%% Specification
% This class defines the subclass of metabolic networks. This class allows members with
% arbitrary number of nodes and inputs. Each node or input can regulate a node through
% activation, repression, null.
classdef MNetwork < BioSystem
  methods
    function obj = MNetwork(up_params,rep_params)
      %Level of each component denote active fraction of that component.
      %INPUTS
      %up_params - Up-regulation topogical matrix.
      % {'<type>','<type>'[,...];'<type>','<type>'[,...]};
      % type=yes|no|linear; Each row represent regulation of one component by all other
      % components.
      %rep_params - Down-regulation topological matrix. same format and size as action_params.
      
      %% input checking
      if ~all(size(up_params)==size(rep_params))
        error('MetebolicNetwork:constructor','Size of the two inputs must be equal!')
      end
      
      %% actual work of generating fluxes
      [nx,nt]=size(up_params);
      dx=containers.Map(); flux=containers.Map();
      for i=1:nx
        chari=num2str(i);  charx=['x',chari];
        dx(charx)=sym(0);
        for j=1:nt %regulation of xi by xj
          charj=num2str(j); symax=sym(['x',charj]);
          H=['H',chari,'_',charj];fH=['fH',chari,'_',charj];V=sym(['V',chari,'_',charj]);
          if ~strcmp(up_params{i,j},'no')
            if strcmp(up_params{i,j},'yes')
              flux(fH)=MNetwork.f_regulate('up',i,j);
            elseif strcmp(up_params{i,j},'linear')
              flux(fH)=MNetwork.f_regulate('uplinear',i,j);
            end
            flux(H)=symax*V*fH;
          else
            flux(H)=sym(0);
          end
          D=['D',chari,'_',charj];fD=['fD',chari,'_',charj];Vd=sym(['Vd',chari,'_',charj]);
          if ~strcmp(rep_params{i,j},'no')
            if strcmp(rep_params{i,j},'yes')
              flux(fD)=MNetwork.f_regulate('down',i,j);
            elseif strcmp(rep_params{i,j},'linear')
              flux(fD)=MNetwork.f_regulate('downlinear',i,j);
            end
            flux(D)=symax*Vd*fD;
          else
            flux(D)=sym(0);
          end
          dx(charx)=dx(charx)+H-D;
        end
      end
      
      %% generate BioSystem object
      obj@BioSystem(dx,flux,containers.Map,containers.Map);
      k_params=obj.inputvars;
      prange=containers.Map();
      for k=k_params
        p=k{:};
        if regexp(p,'x')
          prange(p)=[0.1,1];
        elseif regexp(p,'V')
          % preventing sumsqsys minimization from attaining non-steady states
          prange(p)=[2e-3,2];
        else
          prange(p)=[0,2];
        end
      end
      obj.param=containers.Map(k_params,rand_param(prange_convert(prange),1));
      obj.prange=prange;
      obj.xrange=containers.Map(obj.statevars,repmat({[-1.1e-4,1+1.1e-4]},1,obj.sys.length));
      obj.x0=rand(length(obj.statevars),1);
    end
  end
  methods (Static) %helpers
    f=f_regulate(type,self,j);
  end
end
```
