## Supplemental Data 1 for "Principles underlying implementation of *nearly*-homeostatic biological networks": gen_proddegrates.m

```
function [sys_prod,numf_prod,sys_deg,numf_deg]=gen_proddegrates(obj)
% Generate production rates and degradation rates as long as rate equation formula in terms of fluxes 
% has not been changed.
%
% INPUT
% obj - RNetwork object
% OUTPUT
% sys_prod - container map of symbolic production rates. Each key is a separate state variable. 
% numf_prod - container map of numeric functions [(p,x) or (p,x,u)] of production rates.  
%  Each key is a separate state variable. 
% sys_deg - container map of symbolic degradation rates. Each key is a separate state variable. 
% numf_deg - container map of numeric functions [(p,x) or (p,x,u)] of degration rates. 
%  Each key is a separate state variable. 

%% computing required information
dim_x=obj.sys.Count;
sv=obj.statevars;
iv=obj.inputvars;
smra_k=obj.flux.keys;
smra_k=smra_k(...
  cellfun(@(x)~isempty(regexp(x,'^smra_\d+_\d+$','once')),smra_k,'UniformOutput',true));
n_rxns=numel(smra_k)/dim_x;
heaviside_fast=nan;
syms heaviside_fast(x)
sys_prod=containers.Map;
sys_deg=containers.Map;
numf_prod=containers.Map;
numf_deg=containers.Map;

%% split up Hill function canomical form to generate symbolic production rate and degradation rate
for i=1:dim_x
  x=sv{i};
  i_char=regexprep(char(obj.sys(x)),'^.*smpa_(\d+)_\d+.*$','$1');
  sys_prod(x)=sym(0);
  sys_deg(x)=sym(0);
  for j=1:n_rxns
    j_char=num2str(j);
    smpa_ij=['smpa_',i_char,'_',j_char];
    smra_ij=['smra_',i_char,'_',j_char];
    fv_j=['fv',j_char];
    fvr_ij=['fvr',i_char,'_',j_char];
    addrate_prod=['heaviside_fast(',smpa_ij,'-',smra_ij,') * ',...
      '(',smpa_ij,'*',fv_j,' - ',smra_ij,'*',fvr_ij,')'];
    addrate_deg=['- (1-heaviside_fast(',smpa_ij,'-',smra_ij,')) * ',...
      '(',smpa_ij,'*',fv_j,' - ',smra_ij,'*',fvr_ij,')'];
    sys_prod(x)=sys_prod(x)+str2sym(addrate_prod);
    sys_deg(x)=sys_deg(x)+str2sym(addrate_deg);
  end
end

%% generate numeric form of production and degradation rates
sys_prod_full=BioSystem.fullexp(sys_prod,obj.flux);
sys_deg_full=BioSystem.fullexp(sys_deg,obj.flux);
for i=1:dim_x
  x=sv{i};
  if isempty(obj.inputvars)
    numf_prod(x)=matlabFunction(sys_prod_full(x),'Vars',{sym(obj.param.keys).',sym(sv).'});
    numf_deg(x)=matlabFunction(sys_deg_full(x),'Vars',{sym(obj.param.keys).',sym(sv).'});
  else
    numf_prod(x)=matlabFunction(sys_prod_full(x),'Vars',{sym(obj.param.keys).',sym(sv).',sym(iv).'});
    numf_deg(x)=matlabFunction(sys_deg_full(x),'Vars',{sym(obj.param.keys).',sym(sv).',sym(iv).'});
  end
end
end
```
