## Supplemental Data 1 for "Principles underlying implementation of *nearly*-homeostatic biological networks": RNetwork.html

### Contents

- Specification and design

### Specification and design

- **\*sm\_rules\_satisfied**\* determines whether reaction and product stoichiometric matricies violate constraints such as for zero-order degradation
- **\*sm\_gk\_satisfied**\* determine whether the provided reaction and product stoichiometric matrices satisfy Gupta and Khammash (2022)'s condition for perfect adaptation.
- **\*sm\_gk\_gen**\* generates a new reactant and product stoichiometric matrix based on provided number of state variables and number of reactions

Additional methods are added on top of BioSystem as required by findnhp and findnhp\_fmincon.

- **istrivialss**, **discretize\_bf**, **gd\_reset**, **gd\_discretize\_check**

Additional methods are added on top of BioSystem to generate inverse homeostasis plots.

- **gen\_proddegrates** generates production rates and degradation rates of the object. Both symbolic and numeric forms are generated.

Published with MATLAB® R2022b
