## Supplemental Data 1 for "Principles underlying implementation of *nearly*-homeostatic biological networks": prep4_sma.m

```
function prep4_sma(obj)
% Modifies obj such that sma can be determined: appending sma from flux to the output.
for rp={'r','p'}
  rp_a=rp{:};
  smrpa_regexp=regexp(obj.flux.keys,['^sm',rp_a,'a_\d+_\d+$'],'once');
  smrpa_regexp=cellfun(@(x)~isempty(x),smrpa_regexp,'UniformOutput',true);
  smrpa=extract_farray(obj.flux.keys,smrpa_regexp);
  smrpa=smrpa(:);
  smrpa_nondup=setdiff(smrpa,obj.output_label);
  obj.output_label=[obj.output_label;smrpa_nondup];
  obj.output=[obj.output;str2sym(smrpa_nondup)];
end
end
```
