## Supplemental Data 1 for "Principles underlying implementation of *nearly*-homeostatic biological networks": RNetwork.m

```
% This class defines a subset of BioSystem that are only reaction networks.
classdef RNetwork < BioSystem
  %% Specification and design
  % * **sm_rules_satisfied** determines whether reaction and product stoichiometric matricies 
  % violate constraints such as for zero-order degradation
  % * **sm_gk_satisfied** determine whether the provided reaction and product stoichiometric 
  % matrices satisfy Gupta and Khammash (2022)'s condition for perfect adaptation. 
  % * **sm_gk_gen** generates a new reactant and product stoichiometric matrix based on
  % provided number of state variables and number of reactions
  % 
  % Additional static methods are added on top of BioSystem as required by findnhp and findnhp_fmincon.
  %
  % * *istrivialss*, *discretize_bf*, *gd_discretize*, *gd_reset*, *gd_discretize_check*
  %
  % Additional methods are added on top of BioSystem to generate inverse homeostasis plots.
  %
  % * *gen_proddegrates* generates production rates and degradation rates of the object.
  % Both symbolic and numeric forms are generated.
  %
  % Additional methods
  % 
  % * *prep4_sma* modifies the object such that sma can be determined. 

  properties (Dependent)
    % A struct of two fields, react and prod, that puts numeric smra and smpa into 
    % matrix form. To retrieve this property, the output of object must contain all smra_i_j 
    % and smpa_i_j
    sma
  end
  methods
    function obj = RNetwork(smr,smp,mode,sv_custom)
      % Generate a reaction network from stoichiometric matrices. Fluxes
      % are all scalars to allow the gradient descent algorithm to find near-homeostasis
      % supporting parameter sets.
      % INPUT
      % smr - stoichoimetric matrix for the reactants; symbolic or numeric;
      %   each column specifies coefficients of reactants for a distinct reaction. 
      %   If a matrix element is a symbolic variable, then that variable becomes a parameter of the
      %   reaction network
      % smp - stoichoimetric matrix for the products; symbolic or numeric;
      %   each column specifies coefficients of products for a distinct reaction
      % mode - OPTIONAL with 'normal' as the default. 'normal'|'general2'.
      %   'general2' assumes that entries in the reactant and product stoichoimetric
      %   matrices is 0, 1, or 2.
      % sv_custom - OPTIONAL; a cell array of characters that customize state variables; the default
      %   variable order maybe different from supplied, so stoichiometric matrix should be
      %   regenerated. If this input is specified, one should carefully inspect the fluxes to make
      %   sure that elements of this input does not conflict with intended parameters or flux.keys. 
      % OUTPUT
      % A - reaction network object that a subclass of BioSystems, where fluxes are
      % - smr/pa_i_j = var_b1 + var_b2 are the actual stoichoimetric coefficients 
      %   that is a function of smr/ps_i_j. Linking parameters together should be
      %   done here: var_b1 = var2_b1. 
      % - smr/ps_i_j is a function of r/p, 
      % - smr/pd_i_j = 0,1 is a parameter that indicates whether nearly discrete 
      %   smr/p_i_j maps to discrete smr/ps_i_j. The default parameter range
      %   for random parameter generation is 0. 
      % - when smr/pd_i_j is 1, smr/pst_i_j is a parameter ensures 
      %   smr/p_i_j = smr/pst_start_i_j gets mapped to smr/ps_i_j = smr/pst_i_j.
      % - smr/p_i_j is taken directly from the constructor. Possibly varying topological
      %   parameter must take the form smr/pi_j. 
      % - fv as the flux vector,
      % - fvr matrix for row wise dot product with smra such that the rate of xi is
      %   smr/pa(i,:)*fv(:) - smr/pa(i,:)*fvr(i,:).'.
      % - - fvr(2,c).b3 = [b1(1,c),1,b1(2,c),...,b1(dim(x),c)]
      % smr/pa is the same as smr/p_ when the mode is normal. 
      %
      % Code simplification on 2023-10-27

      if nargin<=3
        sv_custom=strcat('x',arrayfun(@num2str,1:size(smr,1),'UniformOutput',false));
        if nargin<=2
          mode='normal';
        end
      end

      extract_fmatrix=nan;
      min=nan;
      max=nan;
      smoothstep_shifted_Rnoband=nan;
      smoothstep_shifted_d1_Rnoband=nan;
      shifted_identity_Rnoband=nan;
      shifted_identity_dx_Rnoband=nan;
      syms extract_fmatrix(M,x,y) min(x,y) max(x,y) 
      syms smoothstep_shifted_Rnoband(Ns,Ks,s) smoothstep_shifted_d1_Rnoband(Ns,Ks,s)
      syms shifted_identity_Rnoband(x,x0,shifted) shifted_identity_dx_Rnoband(x,x0,shifted)

      %%%% checking inputs
      if any(size(smr)~=size(smp))
        error('RNetwork:smr_or_smp_size',...
          'smr must have the same size as smp!')
      end

      %%%% construct the BioSystems object
      % set up flux rates: for each reaction flux, there is a flux rate for each component such that
      % the degradation rate of that component is zero if the level of that component is zero.
      % - flux: must be scalars such that vecfun_form can be called when computing A_aug
      flux=containers.Map2();
      [r,c]=ndgrid(1:size(smr,1),1:size(smr,2));
      r_c_char=strcat(arrayfun(@num2str,r,'UniformOutput',false),'_',...
        arrayfun(@num2str,c,'UniformOutput',false));
      fv_name=strcat('fv',arrayfun(@num2str,1:size(smr,2),'UniformOutput',false));
      fv_name_sym=sym(fv_name);
      fvr_name=strcat('fvr',r_c_char);
      fvr_name_sym=sym(fvr_name);
      [c,r]=ndgrid(1:size(smr,2),1:size(smr,1));
      sv=sv_custom;
      sv_sym=str2sym(sv);
      mas_sv_name=strcat('mas',arrayfun(@num2str,c,'UniformOutput',false),'_',sv(r));
      mas_sv_name_sym=sym(mas_sv_name);
      masr_sv_name=strcat('masr',arrayfun(@num2str,c,'UniformOutput',false),'_',sv(r));
      masr_sv_name_sym=sym(masr_sv_name);

      % set up stoichoimetric matrix and sm_shifted for both reactants and products
      rp_to_full=struct('r','r','p','p');
      for rp={'r','p'}
        rp_a=rp{:};
        sm_rc_name.(rp_a)=strcat('sm',rp_a,'_',r_c_char);
        sm_rc_name_sym.(rp_a)=sym(sm_rc_name.(rp_a));
        smd_rc_name.(rp_a)=strcat('sm',rp_a,'d_',r_c_char);
        smd_rc_name_sym.(rp_a)=sym(smd_rc_name.(rp_a));
        smst_rc_name.(rp_a)=strcat('sm',rp_a,'st_',r_c_char);
        smst_rc_name_sym.(rp_a)=sym(smst_rc_name.(rp_a));
        sms_rc_name.(rp_a)=strcat('sm',rp_a,'s_',r_c_char);
        sms_rc_name_sym.(rp_a)=sym(sms_rc_name.(rp_a));
        sma_rc_name.(rp_a)=strcat('sm',rp_a,'a_',r_c_char);
        sma_rc_name_sym.(rp_a)=sym(sma_rc_name.(rp_a));
        b1_nkbl_name.(rp_a)=strcat(sms_rc_name.(rp_a),'_b1_nkbl');
        b1_nkbr_name.(rp_a)=strcat(sms_rc_name.(rp_a),'_b1_nkbr');
        b1_name.(rp_a)=strcat(sms_rc_name.(rp_a),'_b1');
        b1_name_sym.(rp_a)=sym(b1_name.(rp_a));
        b2_nkbl_name.(rp_a)=strcat(sms_rc_name.(rp_a),'_b2_nkbl');
        b2_nkbr_name.(rp_a)=strcat(sms_rc_name.(rp_a),'_b2_nkbr');
        b2_name.(rp_a)=strcat(sms_rc_name.(rp_a),'_b2');
        b2_name_sym.(rp_a)=sym(b2_name.(rp_a));
        b3_name.(rp_a)=strcat(sms_rc_name.(rp_a),'_b3');
        b3_name_sym.(rp_a)=sym(b3_name.(rp_a));
      end

      % filling in reaction fluxes for smp for normal mode and general2 and stoichoimetric 
      % matrices, as well as mas_sv and masreac_sv
      gfp_k_sym=sym('gfp_k');
      gfp_n_sym=sym('gfp_n');
      flux('extra_flux')=str2sym('gfp_n+gfp_k');
      for c=1:size(smr,2)
        c_char=num2str(c);
        for r=1:size(smr,1)
          flux(sm_rc_name.r{r,c})=smr(r,c);
          flux(sm_rc_name.p{r,c})=smp(r,c);
          if r==1
            flux(fv_name{c})=str2sym(['krf',c_char]);
          end
          if strcmp(mode,'normal') % normal mode
            flux(sma_rc_name.r{r,c})=sm_rc_name_sym.r(r,c);
            flux(sma_rc_name.p{r,c})=sm_rc_name_sym.p(r,c);
            flux(mas_sv_name{c,r})=sv_sym(r)^sma_rc_name_sym.r(r,c);
            flux(fv_name{c})=flux(fv_name{c})*mas_sv_name_sym(c,r);
          elseif strcmp(mode,'general2') % general2 mode
            for rp={'r','p'}
              rp_a=rp{:};
              % actual stoichoimatrix coefficient as a function of user inputted topological
              % parameter
              flux(sms_rc_name.(rp_a){r,c})=...
                shifted_identity_Rnoband(sm_rc_name_sym.(rp_a)(r,c),...
                ... smst_rc_name_sym.(rp_a)(r,c),smd_rc_name_sym.(rp_a)(r,c));
                smst_rc_name_sym.(rp_a)(r,c),0);
              [topo_act,b1,b2,b1_nkbl,b1_nkbr,b2_nkbl,b2_nkbr]=gen_form_sstep(...
                sms_rc_name.(rp_a){r,c},smd_rc_name_sym.(rp_a)(r,c),...
                gfp_n_sym,gfp_k_sym,sms_rc_name_sym.(rp_a)(r,c));
              flux(b1_nkbl_name.(rp_a){r,c})=b1_nkbl;
              flux(b1_nkbr_name.(rp_a){r,c})=b1_nkbr;
              flux(b1_name.(rp_a){r,c})=b1;
              flux(b2_nkbl_name.(rp_a){r,c})=b2_nkbl;
              flux(b2_nkbr_name.(rp_a){r,c})=b2_nkbr;
              flux(b2_name.(rp_a){r,c})=b2;
              flux(sma_rc_name.(rp_a){r,c})=topo_act;
            end
            flux(mas_sv_name{c,r})=...
              ... % General forms, where b1 in [0,1] and b2 in [0,1]
              ... % b3=b1 for production flux, b3=1 for reaction flux such that degradation
              ... % rate of x(r) is zero when x(r)=0
              (1-b2_name_sym.r(r,c))*((sv_sym(r)+1)^b1_name_sym.r(r,c)-b3_name_sym.r(r,c)) + ...
              b2_name_sym.r(r,c)*sv_sym(r)^2;
              ... ... (sv_sym(r)+1)^(b1_name_sym.r(r,c)+b2_name_sym.r(r,c)) - ...
              ... ... (b3_name_sym.r(r,c)-b2_name_sym.r(r,c)) - ...
              ... ... b2_name_sym.r(r,c)*(2*sv_sym(r)+1);
              ... ... (1-b3_name_sym.r(r,c))*(1-b1_name_sym.r(r,c)) + ...
              ... ... sv_sym(r)*b1_name_sym.r(r,c)^1.55*(1-b2_name_sym.r(r,c)) + ...
              ... ... sv_sym(r)^2*b2_name_sym.r(r,c)^1.55;
            flux(masr_sv_name{c,r})=subs(flux(mas_sv_name{c,r}),b3_name_sym.r(r,c),1);
            flux(mas_sv_name{c,r})=subs(flux(mas_sv_name{c,r}),b3_name_sym.r(r,c),b1_name_sym.r(r,c));
            flux(fv_name{c})=flux(fv_name{c})*mas_sv_name_sym(c,r);
          else
            error('RNetworks:mode','The selected mode does not exist!');
          end
        end
      end

      % computing fluxes fvr
      for c=1:size(smr,2)
        for r=1:size(smr,1)
          if strcmp(mode,'general2')
            flux(fvr_name{r,c})=subs(flux(fv_name{c}),mas_sv_name_sym(c,r),masr_sv_name_sym(c,r));
          else
            flux(fvr_name{r,c})=fv_name_sym(c);
          end
        end
      end

      % dx and xrange
      dx=containers.Map2();
      xrange=containers.Map2();
      for i=1:size(smr,1)
        svi=sv{i};
        dx(svi)=sma_rc_name_sym.p(i,:)*fv_name_sym.' - ...
          sma_rc_name_sym.r(i,:)*fvr_name_sym(i,:).';
        xrange(svi)=[-1.1e-4,inf]; % ensure there is an open set containing desired xrange
      end
      obj@BioSystem(dx,flux,containers.Map2,containers.Map2);
      obj.xrange=xrange;

      % param and prange
      k_params=obj.inputvars;
      prange=containers.Map2(k_params,repmat({[0,2]},size(k_params)));
      for k=k_params
        p=k{:};
        if ~isempty(regexp(p,'^krf','once'))
          prange(p)=[0,10];
        elseif ~isempty(regexp(p,'^sm[a-z]+st_','once'))
          prange(p)=[0,10];
        elseif ~isempty(regexp(p,'^sm[a-z]+d_','once'))
          prange(p)=[1,1]; % default to enable true discretization, unless true shifted_identity is on
        end
      end
      prange('gfp_n')=[100,100]; % do not use inf, as inf/(inf+1)==nan
      prange('gfp_k')=[2/3,2/3];
      obj.prange=prange;
      obj.param=containers.Map2(k_params,rand_param(prange_convert(prange),1));

      % x0
      obj.x0=rand(length(obj.statevars),1);
      obj.num_sim_only=true; % hopefully reduce numerical inaccuracy

      % important simulation parameters
      obj.crash_sim=true; % general form can result in complex numbers otherwise.
    end
    function [value]=get.sma(obj)
      % compute n_sv and n_rxns
      smra_regexp=regexp(obj.output_label,'^smra_\d+_\d+$','once'); % output_label is faster than flux
      smra_regexp=cellfun(@(x)~isempty(x),smra_regexp,'UniformOutput',true);
      n_sv=regexprep(extract_farray(obj.output_label,smra_regexp),'^.*_(\d+)_\d+$','$1');
      n_sv=max(cellfun(@str2double,n_sv));
      n_rxns=sum(smra_regexp)/n_sv;
      if n_rxns~=ceil(n_rxns)
        error('RNetwork:sma','smra and smpa must each have n_sv by n_rxns terms');
      end

      % fill in sma.r and sma.p as matrices
      if ~isempty(obj.inputvars)
        out_obj=obj.numf_output(obj.x0,obj.ut(0));
      else
        out_obj=obj.numf_output(obj.x0);
      end
      [r,c]=ndgrid(1:n_sv,1:n_rxns);
      for rp={'r','p'}
        rp_a=rp{:};
        sma_names.(rp_a)=strcat(['sm',rp_a,'a_'],...
          arrayfun(@(x)num2str(x),r,'UniformOutput',false),'_',...
          arrayfun(@(x)num2str(x),c,'UniformOutput',false));
        [~,sma_IdxInOut.(rp_a)]=ismember(sma_names.(rp_a),obj.output_label);
        if any(sma_IdxInOut.(rp_a)==0)
          error('RNetwork:sma','smra and smpa must each have n_sv by n_rxns terms');
        end
        value.(rp_a)=extract_farray(out_obj,sma_IdxInOut.(rp_a));
      end
    end
    prep4_sma(obj);
  end
  methods(Static=true)
    function [topos_d,l_d,topos_act_fdisc] = gd_discretize_check(A_aug,parsopt_topo,topovs,topos_thres)
      % Return discretize topology if the right criteria are met: mapped topology is within topo_thres 
      % of discrete topology AND every component still has a reaction with net degradation.
      % Otherwise, return the original topology.
      % 
      % - Wider smp - smr gap resulted in 10/41 rather than 10/74 searches being successful
      % (2023-11-09)
      % 
      % INPUT
      % A_aug - BioSystem object. Output must contain topo_act for all topo parameters; 
      %  state variable must contain paropt_topo
      % parsopt_topo - cell array of character vector of the names of the topological parameters
      % topovs - numeric vector
      % topos_thres - cell array that determines the threshold the calculated
      %  topology must fall under to be discretized OR 'up' OR 'down' OR 'measure'. 
      %  If 'up', then tries to evaluate whether the ideal higher discrete value satisfy other 
      %  constraints. If 'up/down-measure', only topo_act_fdisc is computed and returned. 
      % OUTPUT
      % topos_d - numeric array; either discretized topology or original topology value
      % l_d - logical array; whether topology is discretized
      % topos_act_fdisc - how far is current topo act from discrete topo_act; topo_act - topo_d

      % getting necessary preliminary information
      topo_drefs=[0,1,2];
      x0_map_orig=A_aug.x0_map;
      x0_map=[containers.Map;A_aug.x0_map];
      topos_act_name=regexprep(parsopt_topo,'^([a-z_]+)(\d)','$1a_$2');
      [~,toposact_in_out_idx]=ismember(topos_act_name,A_aug.output_label);

      % compute candidate discrete topology based on topo_thres
      x0_map=[x0_map; containers.Map(parsopt_topo,topovs)];
      out_Aaug=A_aug.numf_output(cell2mat(x0_map.values).',A_aug.ut(0));
      topos_act=extract_farray(out_Aaug,toposact_in_out_idx);
      topos_dc=nan(size(topos_act));
      for k=1:numel(parsopt_topo)
        if isa(topos_thres{k},'numeric')
          drefs_idx=find(abs(topos_act(k)-topo_drefs)<=topos_thres{k},1);
          if isempty(drefs_idx)
            topos_dc(k)=topovs(k);
          else
            topos_dc(k)=topo_drefs(drefs_idx);
          end
        elseif ~isempty(regexp(topos_thres{k},'up','once'))
          drefs_idx=find(topovs(k)<=topo_drefs,1,'first');
          if ~isempty(drefs_idx)
            topos_dc(k)=topo_drefs(drefs_idx);
          else
            topos_dc(k)=max(topo_drefs);
          end
        elseif ~isempty(regexp(topos_thres{k},'down','once'))
          drefs_idx=find(topovs(k)>=topo_drefs,1,'last');
          if ~isempty(drefs_idx)
            topos_dc(k)=topo_drefs(drefs_idx);
          else
            topos_dc(k)=min(topo_drefs);
          end
        end
      end

      % discretization of topology if all criteria are met
      x0_map=[x0_map; containers.Map(parsopt_topo,topos_dc)];
      A_aug.x0_map=x0_map;
      sma=A_aug.sma;
      topos_d=nan(size(topos_dc));
      l_d=false(size(topos_dc));
      topos_act_fdisc=nan(size(topos_dc));
      l_netdeg=all(any(sma.p-sma.r<-0.01,2)); % all comp have 1 net degradation reaction
      for k=1:numel(parsopt_topo)
        if l_netdeg && any(topos_dc(k)==topo_drefs) && ...
            (isa(topos_thres{k},'numeric') || isempty(regexp(topos_thres{k},'measure','once')))
          l_d(k)=true;
        elseif any(topos_dc(k)==topo_drefs) && ...
            isa(topos_thres{k},'char') && ~isempty(regexp(topos_thres{k},'measure','once'))
          l_d(k)=true;
        else
          l_d(k)=false;
        end
        if l_d(k)
          topos_d(k)=topos_dc(k);
          topos_act_fdisc(k)=topos_act(k)-topos_d(k);
        elseif ~l_d(k)
          topos_d(k)=topovs(k);
          topos_act_fdisc(k)=0;
        else
          error('This scenario should NOT occur!');
        end
      end

      % ensure A_aug.x0 is unmodified
      A_aug.x0_map=x0_map_orig;
    end

    function gd_reset(A_aug,pars_opt_topo)
      % A_aug.param('xnorm_thres')=100;
      % reset dobj_<par_opt>_dval_j to appropriate values
      for k=pars_opt_topo
        key=k{:};
        paropt_start=regexprep(key,'^(sm[a-z]+)(\d+_\d+)$','$1st_$2');
        paropt_disc=regexprep(key,'^(sm[a-z]+)(\d+_\d+)$','$1d_$2');
        A_aug.param(paropt_start)=0;
        A_aug.param(paropt_disc)=0;
        A_aug.param(['dobj_',key,'_dval_1'])=0;
        A_aug.param(['dobj_',key,'_dval_2'])=1;
        A_aug.param(['dobj_',key,'_dval_3'])=2;
      end
    end

    function gd_discretize(A_aug,par_opt,status_d)
      % Modify A_aug to ensure discretized topology is truly discrete depending on the topological
      % parameter.
      %
      % INPUT
      % status_d - 0 means decrease to first possible discrete value; 1 means increase to first possible
      %   discrete value
      val_orig=A_aug.x0_map(par_opt);
      paropt_start=regexprep(par_opt,'^(sm[a-z]+)(\d+_\d+)$','$1st_$2');
      paropt_disc=regexprep(par_opt,'^(sm[a-z]+)(\d+_\d+)$','$1d_$2');
      A_aug.param(paropt_start)=val_orig;
      A_aug.param(paropt_disc)=1;
      if status_d==0 && val_orig>2
        % do nothing
        1;
      elseif status_d==0 && val_orig>1
        A_aug.param(['dobj_',par_opt,'_dval_3'])=A_aug.param(['dobj_',par_opt,'_dval_1']);
      elseif status_d==0 && val_orig<1
        A_aug.param(['dobj_',par_opt,'_dval_2'])=A_aug.param(['dobj_',par_opt,'_dval_1'])-...
          abs(A_aug.param(['dobj_',par_opt,'_dval_2'])-A_aug.param(['dobj_',par_opt,'_dval_1']));
      elseif status_d==1 && val_orig<0
        % do nothing
        1;
      elseif status_d==1 && val_orig<1
        A_aug.param(['dobj_',par_opt,'_dval_1'])=A_aug.param(['dobj_',par_opt,'_dval_3']);
      elseif status_d==1 && val_orig>1
        A_aug.param(['dobj_',par_opt,'_dval_2'])=A_aug.param(['dobj_',par_opt,'_dval_3'])+...
          abs(A_aug.param(['dobj_',par_opt,'_dval_2'])-A_aug.param(['dobj_',par_opt,'_dval_3']));
      end
    end
    function dtopo = discretize_bf(topo)
      % INPUT
      % topo - topological parameter
      % OUTPUT
      % dtopo - discretized topological parameter
      weights=[0.5,1,0.5]; % ensure all topo state given equal weighting
      dtopo=extract_farray([0,1,2],...
        find( weights.*abs(topo-[0,1,2])==min(weights.*abs(topo-[0,1,2])) , 1 ));
      if isempty(dtopo) && isnan(topo)
        dtopo=nan;
      end
    end
    function l=istrivialss(xss)
      % INPUTS
      % xss - vector
      l=sqrt(sum(xss(2:end).^2))<1e-8 || xss(1)<1e-3 || ... % trivialss criteria
        any(xss<-1e-5); % out of bound detection
    end
    function l = sm_rules_satisfied(smr,smp)
      % See if stoimetric matrix efficients rules are satisfied
      % * Rule 1: zeroth order degradation - a product entry can only be negative when 
      % reactant entry at identical position is 0. 
      % INPUT
      % smr - stoichoimetric matrix for the reactants; numeric; 
      % smp - stoichoimetric matrix for the products; numeric;
      % OUTPUT
      % l - logical scalar
      l=all(smr(smp<0)==0);
    end
    function [l,l_r,l_c,l_q,qT,qTS1tom] = sm_gk_satisfied(smr,smp,m)
      % INPUT
      % smr - stoichoimetric matrix for the reactants; numeric; 
      % smp - stoichoimetric matrix for the products; numeric;
      % m - first m reactions of smr and smp are involved in the
      %  set point. l_c is only valid for m=2
      % OUTPUT
      % l - logical scalar indicating whethter GK's conditions are satisfied
      % l_r - logical scalar indicating whether GK's rank condition is satisfied. 
      % l_c - logical scalar indicating whether GK's first condition
      %  regarding controller's reactions is satisfied
      % l_q - logical scalar indicating whether GK's second condition is satisifed
      % qT - row vector qT or matrix as stated in Gupta and Khammash (2022); 
      % qTS1tom - row vector or matrix as stated in Gupta and Khammash
      
      sm=smp-smr;
      %%%% see if rank(sm)=size(sm,1)
      l_r=(rank(sm)==size(sm,1));

      %%%% GK1: see if it is satisfied; smr(1,1)~=smr(1,2) when m=1 AND
      %  the effect of x2ss(u) to xNss(u) on qT x S x c = 0 has to be none or separable
      %  for each set point encoding parameter associated mass action.
      %
      %  The condition is generalized for m>2
      l_c1=m>2 || ... 
        ... % GK1.1: condition in paper but cannot find justification for it
        smr(1,1)~=smr(1,2);
      ij_comb=nchoosek(1:m,2); % matrix w 2 columns
      for r=1:size(ij_comb)
        % Here, I'm assuming the effect is separable for only 2 set point encoding 
        % parameter associated mass actions
        l_c=l_c1 && all(smr(2:end,ij_comb(r,1))==smr(2:end,ij_comb(r,2)));
        if l_c; break; end
      end

      %%%% GK2: see if a non-zero q exists such that qT x S(:,m+1:K) = 0 AND 
      %  qT x S(:,1:m) have opposite signs when m=2
      q=null(sm(:,m+1:end).');
      qT=q.';
      if numel(q)>=1 
        qTS1tom=qT*sm(:,1:m); % row vector if qT is only a vector
        if m>2 || ... 
            ... % GK2.2: opposite signs in qTS1to2 when m=2
            ... % Since I cannot assume that x1ss=pfi(p1/p2) for m>2, it is difficult to
            ... % generalize this condition to m>2
            any(abs(sum(qTS1tom,2))<sum(abs(qTS1tom),2)) 
          l_q=true;
        else
          l_q=false;
        end
      else
        l_q=false;
        qT=zeros(1,size(smp,1));
        qTS1tom=zeros(1,m);
      end     

      l=l_r && l_c && l_q;
    end
    function [smr,smp,iter] = sm_gk_gen(n_sv,n_rxns,m,sm_react0,sm_prod0,mode)
      % Assuming the number of reaction involved in set point is 2.
      % Currently, the maximum value of each entry in the stoichiometric
      % matrix is 2.
      % 
      % INPUT
      % n_sv - numeric scalar; number of state variables
      % n_rxns - numeric scalar; number of reactions
      % m - number of first reactions to encode the set point
      % sm_react0 - numeric matrix; nan indicate entries that can be randomly chosen
      % sm_prod0 - numeric matrix; nan indicate entries that can be randomly chosen
      % mode - 'both+rank-bimolecular'|'both+rank'|'qT only'|'none*'; 
      %  specifies which Gupta and Khammash (2022)'s criteria need to be satisfied. 
      % OUTPUT
      % smr,smp - stoichoimetric matrices
      % iter - number of random parameters sampled before suitable parameter is found
      
      % customization section
      max_smentry=2; % 2 takes account autocatalytic networks
      
      % generate feasible smr and smp
      max_smentry_rand=max_smentry+1;
      row_ids=1:n_sv;
      iter=0;
      while true
        iter=iter+1;
        smr=floor(rand([n_sv,n_rxns])*max_smentry_rand);
        smr(~isnan(sm_react0))=sm_react0(~isnan(sm_react0));
        smp=floor(rand([n_sv,n_rxns])*max_smentry_rand);
        smp(~isnan(sm_prod0))=sm_prod0(~isnan(sm_prod0));
        if ~isempty(regexp(mode,'none','once'))
          break; 
        elseif strcmp(mode,'qT only') || ~isempty(regexp(mode,'both','once'))
          if ~isempty(regexp(mode,'both','once'))
            % this block should be before bimolecular code block, bc the
            % later code block assumes modification is final. 
            if smr(1,1)==smr(1,2) && m==2
              continue;
            else
              for c=2:m
                smr(2:end,c)=smr(2:end,1);
              end
            end
            
            %
            if ~isempty(regexp(mode,'bimolecular','once'))
              for c=1:n_rxns
                % sel is used to select for entries that can be anything
                % sel selects up to two entries
                sel_rnd=randperm(n_sv);
                sel=[];
                if c<=m 
                  % sel must include 1 in first position for smr(2:end,1)=
                  % smr(2:end,2)
                  sel=[1,sel]; 
                end
                sel=unique([ sel, find( ~isnan(sm_react0(:,c)) & sm_react0(:,c)~=0 )]);
                if numel(sel)>2
                  error('RNetwork:sm_gk_gen:sm_react0',...
                    'smr already violate bimolecular constraints!')
                end
                sel_rnd=setdiff(sel_rnd,sel,'stable');
                sel=[sel,sel_rnd( 1:2-numel(sel) )];
                if c>1 && c<=m
                  sel=selr; % sel for col=c be same as col=1. 
                end
                selr=sel; % logging purpose
                smr(setdiff(row_ids,sel),c)=0;
                if sum(smr(:,c))>2
                  % eliminate one of the two entries to be zero if sum of entries > 2
                  if c<=m
                    sel_elim=2;
                  else
                    sel_elim=ceil(rand(1)*2);
                  end
                  if ~isnan(sm_react0(sel(sel_elim),c)) && sm_react0(sel(sel_elim),c)~=0
                    sel_elim=setdiff([1,2],sel_elim);
                    if ~isnan(sm_react0(sel(sel_elim),c)) && sm_react0(sel(sel_elim),c)~=0
                      continue;
                    end
                  end
                  smr(sel(sel_elim),c)=0;
                end

                sel_rnd=randperm(n_sv);
                sel=[];
                sel=unique([ find( ~isnan(sm_prod0(:,c)) & sm_prod0(:,c)~=0 ),sel ]);
                if numel(sel)>2
                  error('RNetwork:sm_gk_gen:sm_prod0',...
                    'sm_prod0 already violate bimolecular constraints!')
                end
                sel_rnd=setdiff(sel_rnd,sel,'stable');
                sel=[sel,sel_rnd( 1:2-numel(sel) )];
                smp(setdiff(row_ids,sel),c)=0;
                if sum(smp(:,c))>2
                  % eliminate one of the two entries to be zero if sum of entries > 2
                  sel_elim=ceil(rand(1)*2);
                  if ~isnan(sm_prod0(sel(sel_elim),c)) && sm_prod0(sel(sel_elim),c)~=0
                    sel_elim=setdiff([1,2],sel_elim);
                    if ~isnan(sm_prod0(sel(sel_elim),c)) && sm_prod0(sel(sel_elim),c)~=0
                      continue;
                    end
                  end
                  smp(sel(sel_elim),c)=0;
                end
              end
            end
          end
          [l,l_r,l_c,l_q,qT]=RNetwork.sm_gk_satisfied(smr,smp,m); % self-testing
          if ~isempty(regexp(mode,'both','once'))
            if l 
              break;
            end
          else
            if l_q
              break; 
            end
          end
        end
      end
    end
  end
end
```
