## Supplemental Data 1 for "Principles underlying implementation of *nearly*-homeostatic biological networks": RNetwork_old.m

```
% This class defines a subset of BioSystem that are only reaction networks.
classdef RNetwork < BioSystem
  %% Specification and design
  % * **sm_rules_satisfied** determines whether reaction and product stoichiometric matricies 
  % violate constraints such as for zero-order degradation
  % * **sm_gk_satisfied** determine whether the provided reaction and product stoichiometric 
  % matrices satisfy Gupta and Khammash (2022)'s condition for perfect adaptation. 
  % * **sm_gk_gen** generates a new reactant and product stoichiometric matrix based on
  % provided number of state variables and number of reactions
  % 
  % Additional methods are added on top of BioSystem as required by findnhp and findnhp_fmincon.
  %
  % * *istrivialss*, *discretize_bf*, *gd_discretize*, *gd_reset*, *gd_discretize_check*
  %
  % Additional methods are added on top of BioSystem to generate inverse homeostasis plots.
  %
  % * *gen_proddegrates* generates production rates and degradation rates of the object.
  % Both symbolic and numeric forms are generated.
  %

  properties (Dependent)
    % A struct of two fields, react and prod, that puts numeric sm_react_act and sm_prod_act into 
    % matrix form. To retrieve this property, the output of object must contain all sm_react_act_i_j 
    % and sm_prod_act_i_j
    sm_act
  end
  methods
    function obj = RNetwork(sm_react,sm_prod,mode,sv_custom)
      % Generate a reaction network from stoichiometric matrices. Fluxes
      % are all scalars to allow the gradient descent algorithm to find near-homeostasis
      % supporting parameter sets.
      % INPUT
      % sm_react - stoichoimetric matrix for the reactants; symbolic or numeric;
      %   each column specifies coefficients of reactants for a distinct reaction. 
      %   If a matrix element is a symbolic variable, then that variable becomes a parameter of the
      %   reaction network
      % sm_prod - stoichoimetric matrix for the products; symbolic or numeric;
      %   each column specifies coefficients of products for a distinct reaction
      % mode - OPTIONAL with 'normal' as the default. 'normal'|'general2'.
      %   'general2' assumes that entries in the reactant and product stoichoimetric
      %   matrices is 0, 1, or 2.
      % sv_custom - OPTIONAL; a cell array of characters that customize state variables; the default
      %   variable order maybe different from supplied, so stoichiometric matrix should be
      %   regenerated. If this input is specified, one should carefully inspect the fluxes to make
      %   sure that elements of this input does not conflict with intended parameters or flux.keys. 
      % OUTPUT
      % A - reaction network object that a subclass of BioSystems, where fluxes are
      % - sm_react/prod_act_i_j = b1r/pi_j + b2r/pi_j are the actual stoichoimetric coefficients 
      %   that is a function of sm_react/prod_shifted_i_j. Linking parameters together should be
      %   done here: b1r/pi_j = b1r/pk_l. 
      % - sm_react/prod_shifted_i_j is a function of sm_react/prod_i_j, 
      % - sm_react/prod_disc_i_j = 0,1 is a parameter that indicates whether nearly discrete 
      %   sm_react/prod_i_j maps to discrete sm_react/prod_shifted_i_j. The default parameter range
      %   for random parameter generation is 0. 
      % - when sm_react/prod_disc_i_j is 1, sm_react/prod_start_i_j is a parameter ensures 
      %   sm_react/prod_i_j = sm_react/prod_start_i_j gets mapped to sm_react/prod_shifted_i_j = 
      %   sm_react/prod_start_i_j.
      % - sm_react/prod_i_j is taken directly from the constructor. 
      % - fv as the flux vector,
      % - fvreact matrix for row wise dot product with sm_react_act such that the rate of xi is
      %   sm_prod_act(i,:)*fv(:) - sm_react_act(i,:)*fvreact(i,:).'.
      % - - fvreact(2,c).b3 = [b1(1,c),1,b1(2,c),...,b1(dim(x),c)]
      % sm_react/prod_act is the same as sm_react/prod_ when the mode is normal. 
      %
      % Code simplification on 2023-10-27

      if nargin<=3
        sv_custom=strcat('x',arrayfun(@num2str,1:size(sm_react,1),'UniformOutput',false));
        if nargin<=2
          mode='normal';
        end
      end

      extract_fmatrix=nan;
      min=nan;
      max=nan;
      smoothstep_shifted_Rnoband=nan;
      smoothstep_shifted_d1_Rnoband=nan;
      shifted_identity_Rnoband=nan;
      shifted_identity_dx_Rnoband=nan;
      syms extract_fmatrix(M,x,y) min(x,y) max(x,y) 
      syms smoothstep_shifted_Rnoband(Ns,Ks,s) smoothstep_shifted_d1_Rnoband(Ns,Ks,s)
      syms shifted_identity_Rnoband(x,x0,shifted) shifted_identity_dx_Rnoband(x,x0,shifted)

      %%%% checking inputs
      if any(size(sm_react)~=size(sm_prod))
        error('RNetwork:sm_react_or_sm_prod_size',...
          'sm_react must have the same size as sm_prod!')
      end

      %%%% construct the BioSystems object
      % set up flux rates: for each reaction flux, there is a flux rate for each component such that
      % the degradation rate of that component is zero if the level of that component is zero.
      % - flux: must be scalars such that vecfun_form can be called when computing A_aug
      flux=containers.Map2();
      [r,c]=ndgrid(1:size(sm_react,1),1:size(sm_react,2));
      r_c_char=strcat(arrayfun(@num2str,r,'UniformOutput',false),'_',...
        arrayfun(@num2str,c,'UniformOutput',false));
      fv_name=strcat('fv',arrayfun(@num2str,1:size(sm_react,2),'UniformOutput',false));
      fv_name_sym=sym(fv_name);
      fvreact_name=strcat('fvreact',r_c_char);
      fvreact_name_sym=sym(fvreact_name);
      [c,r]=ndgrid(1:size(sm_react,2),1:size(sm_react,1));
      sv=sv_custom;
      sv_sym=str2sym(sv);
      mas_sv_name=strcat('mas',arrayfun(@num2str,c,'UniformOutput',false),'_',sv(r));
      mas_sv_name_sym=sym(mas_sv_name);
      masreact_sv_name=strcat('masreact',arrayfun(@num2str,c,'UniformOutput',false),'_',sv(r));
      masreact_sv_name_sym=sym(masreact_sv_name);

      % set up stoichoimetric matrix and sm_shifted for both reactants and products
      sstep_nk0_name='sstep_nk0';
      sstep_nk0_name_sym=sym(sstep_nk0_name);
      sstep_nk1_name='sstep_nk1';
      sstep_nk1_name_sym=sym(sstep_nk1_name);
      sstep_n2mk1_name='sstep_n2mk1';
      sstep_n2mk1_name_sym=sym(sstep_n2mk1_name);
      sstep_n2mk2_name='sstep_n2mk2';
      sstep_n2mk2_name_sym=sym(sstep_n2mk2_name);
      rp_to_full=struct('r','react','p','prod');
      for rp={'r','p'}
        rp_a=rp{:};
        b1_name.(rp_a)=strcat('b1',rp_a,r_c_char);
        b1_name_sym.(rp_a)=sym(b1_name.(rp_a));
        b2_name.(rp_a)=strcat('b2',rp_a,r_c_char);
        b2_name_sym.(rp_a)=sym(b2_name.(rp_a));
        b3_name.(rp_a)=strcat('b3',rp_a,r_c_char);
        b3_name_sym.(rp_a)=sym(b3_name.(rp_a));
        sm_rc_name.(rp_a)=strcat('sm_',rp_to_full.(rp_a),'_',r_c_char);
        sm_rc_name_sym.(rp_a)=sym(sm_rc_name.(rp_a));
        sm_act_rc_name.(rp_a)=strcat('sm_',rp_to_full.(rp_a),'_act_',r_c_char);
        sm_act_rc_name_sym.(rp_a)=sym(sm_act_rc_name.(rp_a));
        sm_rc_disc_name.(rp_a)=strcat('sm_',rp_to_full.(rp_a),'_disc_',r_c_char);
        sm_rc_disc_name_sym.(rp_a)=sym(sm_rc_disc_name.(rp_a));
        sm_rc_shifted_name.(rp_a)=strcat('sm_',rp_to_full.(rp_a),'_shifted_',r_c_char);
        sm_rc_shifted_name_sym.(rp_a)=sym(sm_rc_shifted_name.(rp_a));
        sm_rc_start_name.(rp_a)=strcat('sm_',rp_to_full.(rp_a),'_start_',r_c_char);
        sm_rc_start_name_sym.(rp_a)=sym(sm_rc_start_name.(rp_a));
        sstep_b1p1_name.(rp_a)=strcat('sstep_b1p1_',rp_a,'_',r_c_char);
        sstep_b1p1_name_sym.(rp_a)=sym(sstep_b1p1_name.(rp_a));
        sstep_b1p2_name.(rp_a)=strcat('sstep_b1p2_',rp_a,'_',r_c_char);
        sstep_b1p2_name_sym.(rp_a)=sym(sstep_b1p2_name.(rp_a));
        sstep_b2_name.(rp_a)=strcat('sstep_b2_',rp_a,'_',r_c_char);
        sstep_b2_name_sym.(rp_a)=sym(sstep_b2_name.(rp_a));
      end

      % filling in reaction fluxes for sm_prod for normal mode and general2 and stoichoimetric 
      % matrices, as well as mas_sv and masreac_sv
      gfp_k_sym=sym('gfp_k');
      gfp_n_sym=sym('gfp_n');
      flux('extra_flux')=str2sym('gfp_n+gfp_k');
      softstep_var=smoothstep_shifted_Rnoband;
      flux(sstep_nk0_name)=softstep_var(gfp_n_sym,gfp_k_sym,sym(0));
      flux(sstep_nk1_name)=softstep_var(gfp_n_sym,gfp_k_sym,sym(1));
      flux(sstep_n2mk1_name)=softstep_var(gfp_n_sym,2-gfp_k_sym,sym(1));
      flux(sstep_n2mk2_name)=softstep_var(gfp_n_sym,2-gfp_k_sym,sym(2));
      for c=1:size(sm_react,2)
        c_char=num2str(c);
        for r=1:size(sm_react,1)
          flux(sm_rc_name.r{r,c})=sm_react(r,c);
          flux(sm_rc_name.p{r,c})=sm_prod(r,c);
          if r==1
            flux(fv_name{c})=str2sym(['krf',c_char]);
          end
          if strcmp(mode,'normal') % normal mode
            flux(sm_act_rc_name.r{r,c})=sm_rc_name_sym.r(r,c);
            flux(sm_act_rc_name.p{r,c})=sm_rc_name_sym.p(r,c);
            flux(mas_sv_name{c,r})=sv_sym(r)^sm_act_rc_name_sym.r(r,c);
            flux(fv_name{c})=flux(fv_name{c})*mas_sv_name_sym(c,r);
          elseif strcmp(mode,'general2') % general2 mode
            for rp={'r','p'}
              rp_a=rp{:};
              % actual stoichoimatrix coefficient as a function of user inputted topological
              % parameter
              flux(sm_rc_shifted_name.(rp_a){r,c})=...
                shifted_identity_Rnoband(sm_rc_name_sym.(rp_a)(r,c),...
                sm_rc_start_name_sym.(rp_a)(r,c),sm_rc_disc_name_sym.(rp_a)(r,c));
              flux(sstep_b1p1_name.(rp_a){r,c})=...
                (softstep_var(gfp_n_sym,gfp_k_sym,sm_rc_shifted_name_sym.(rp_a)(r,c))-...
                sstep_nk0_name_sym) / ...
                (sstep_nk1_name_sym-sstep_nk0_name_sym);
              flux(sstep_b1p2_name.(rp_a){r,c})=...
                ((1-sstep_nk0_name_sym)/(sstep_nk1_name_sym-sstep_nk0_name_sym)-1) * ...
                softstep_var(sym(1),sym(1.3),sm_rc_shifted_name_sym.(rp_a)(r,c));
              flux(sstep_b2_name.(rp_a){r,c})=...
                (softstep_var(gfp_n_sym,2-gfp_k_sym,sm_rc_shifted_name_sym.(rp_a)(r,c))-...
                sstep_n2mk1_name_sym) / ...
                (sstep_n2mk2_name_sym-sstep_n2mk1_name_sym);
              flux(b1_name.(rp_a){r,c})=sstep_b1p1_name_sym.(rp_a)(r,c)-sstep_b1p2_name_sym.(rp_a)(r,c);
              flux(b2_name.(rp_a){r,c})=sstep_b2_name_sym.(rp_a)(r,c);
              flux(sm_act_rc_name.(rp_a){r,c})=b1_name_sym.(rp_a)(r,c)+b2_name_sym.(rp_a)(r,c);
            end
            flux(mas_sv_name{c,r})=...
              ... % General forms, where b1r in [0,1] and b2r in [0,1]
              ... % b3r=b1r for production flux, b3r=1 for reaction flux such that degradation
              ... % rate of x(r) is zero when x(r)=0
              (1-b2_name_sym.r(r,c))*((sv_sym(r)+1)^b1_name_sym.r(r,c)-b3_name_sym.r(r,c)) + ...
              b2_name_sym.r(r,c)*sv_sym(r)^2;
              ... ... (sv_sym(r)+1)^(b1_name_sym.r(r,c)+b2_name_sym.r(r,c)) - ...
              ... ... (b3_name_sym.r(r,c)-b2_name_sym.r(r,c)) - ...
              ... ... b2_name_sym.r(r,c)*(2*sv_sym(r)+1);
              ... ... (1-b3_name_sym.r(r,c))*(1-b1_name_sym.r(r,c)) + ...
              ... ... sv_sym(r)*b1_name_sym.r(r,c)^1.55*(1-b2_name_sym.r(r,c)) + ...
              ... ... sv_sym(r)^2*b2_name_sym.r(r,c)^1.55;
            flux(masreact_sv_name{c,r})=subs(flux(mas_sv_name{c,r}),b3_name_sym.r(r,c),1);
            flux(mas_sv_name{c,r})=subs(flux(mas_sv_name{c,r}),b3_name_sym.r(r,c),b1_name_sym.r(r,c));
            flux(fv_name{c})=flux(fv_name{c})*mas_sv_name_sym(c,r);
          else
            error('RNetworks:mode','The selected mode does not exist!');
          end
        end
      end

      % computing fluxes fvreact
      for c=1:size(sm_react,2)
        for r=1:size(sm_react,1)
          if strcmp(mode,'general2')
            flux(fvreact_name{r,c})=subs(flux(fv_name{c}),mas_sv_name_sym(c,r),masreact_sv_name_sym(c,r));
          else
            flux(fvreact_name{r,c})=fv_name_sym(c);
          end
        end
      end

      % dx and xrange
      dx=containers.Map2();
      xrange=containers.Map2();
      for i=1:size(sm_react,1)
        svi=sv{i};
        dx(svi)=sm_act_rc_name_sym.p(i,:)*fv_name_sym.' - ...
          sm_act_rc_name_sym.r(i,:)*fvreact_name_sym(i,:).';
        xrange(svi)=[-1.1e-4,inf]; % ensure there is an open set containing desired xrange
      end
      obj@BioSystem(dx,flux,containers.Map2,containers.Map2);
      obj.xrange=xrange;

      % param and prange
      k_params=obj.inputvars;
      prange=containers.Map2(k_params,repmat({[0,2]},size(k_params)));
      for k=k_params
        p=k{:};
        if ~isempty(regexp(p,'^krf','once'))
          prange(p)=[0,10];
        elseif ~isempty(regexp(p,'^sm_\w+_start_','once'))
          prange(p)=[0,10];
        elseif ~isempty(regexp(p,'^sm_\w+_disc_','once'))
          prange(p)=[0,0];
        end
      end
      prange('gfp_n')=[100,100]; % do not use inf, as inf/(inf+1)==nan
      prange('gfp_k')=[2/3,2/3];
      obj.prange=prange;
      obj.param=containers.Map2(k_params,rand_param(prange_convert(prange),1));

      % x0
      obj.x0=rand(length(obj.statevars),1);
      obj.num_sim_only=true; % hopefully reduce numerical inaccuracy
    end
    function [value]=get.sm_act(obj)
      % compute n_sv and n_rxns
      sm_react_act_regexp=regexp(obj.output_label,'^sm_react_act_\d+_\d+$','once');
      sm_react_act_regexp=cellfun(@(x)~isempty(x),sm_react_act_regexp,'UniformOutput',true);
      n_sv=regexprep(obj.output_label(sm_react_act_regexp),'^.*_(\d+)_\d+$','$1');
      n_sv=max(cellfun(@str2double,n_sv));
      n_rxns=sum(sm_react_act_regexp)/n_sv;
      if n_rxns~=ceil(n_rxns)
        error('RNetwork:sm_act','sm_react_act and sm_prod_act must each have n_sv by n_rxns terms');
      end

      % fill in sm_act.react and sm_act.prod as matrices
      if ~isempty(obj.inputvars)
        out_obj=obj.numf_output(obj.x0,obj.ut(0));
      else
        out_obj=obj.numf_output(obj.x0);
      end
      [r,c]=ndgrid(1:n_sv,1:n_rxns);
      for rp={'react','prod'}
        rp_a=rp{:};
        sm_act_names.(rp_a)=strcat(['sm_',rp_a,'_act_'],...
          arrayfun(@(x)num2str(x),r,'UniformOutput',false),'_',...
          arrayfun(@(x)num2str(x),c,'UniformOutput',false));
        [~,sm_act_IdxInOut.(rp_a)]=ismember(sm_act_names.(rp_a),obj.output_label);
        if any(sm_act_IdxInOut.(rp_a)==0)
          error('RNetwork:sm_act','sm_react_act and sm_prod_act must each have n_sv by n_rxns terms');
        end
        value.(rp_a)=extract_farray(out_obj,sm_act_IdxInOut.(rp_a));
      end
    end
  end
  methods(Static=true)
    function [topos_d,l_d,topos_act_fdisc] = gd_discretize_check(A_aug,parsopt_topo,topovs,topos_thres)
      % Return discretize topology if the right criteria are met: mapped topology is within topo_thres 
      % of discrete topology AND every component still has a reaction with net degradation.
      % Otherwise, return the original topology.
      % 
      % - Wider sm_prod - sm_react gap resulted in 10/41 rather than 10/74 searches being successful
      % (2023-11-09)
      % 
      % INPUT
      % A_aug - BioSystem object. Output must contain topo_act for all topo parameters; 
      %  state variable must contain paropt_topo
      % parsopt_topo - cell array of character vector of the names of the topological parameters
      % topovs - numeric vector
      % topos_thres - cell array that determines the threshold the calculated
      %  topology must fall under to be discretized OR 'up' OR 'down' OR 'measure'. 
      %  If 'up', then tries to evaluate whether the ideal higher discrete value satisfy other 
      %  constraints. If 'up/down-measure', only topo_act_fdisc is computed and returned. 
      % OUTPUT
      % topos_d - numeric array; either discretized topology or original topology value
      % l_d - logical array; whether topology is discretized
      % topos_act_fdisc - how far is current topo act from discrete topo_act; topo_act - topo_d

      % getting necessary preliminary information
      topo_drefs=[0,1,2];
      x0_map_orig=A_aug.x0_map;
      x0_map=[containers.Map;A_aug.x0_map];
      topos_act_name=regexprep(parsopt_topo,'^([a-z_]+)(\d)','$1_act_$2');
      [~,toposact_in_out_idx]=ismember(topos_act_name,A_aug.output_label);

      % compute candidate discrete topology based on topo_thres
      x0_map=[x0_map; containers.Map(parsopt_topo,topovs)];
      out_Aaug=A_aug.numf_output(cell2mat(x0_map.values).',A_aug.ut(0));
      topos_act=extract_farray(out_Aaug,toposact_in_out_idx);
      topos_dc=nan(size(topos_act));
      for k=1:numel(parsopt_topo)
        if isa(topos_thres{k},'numeric')
          drefs_idx=find(abs(topos_act(k)-topo_drefs)<=topos_thres{k},1);
          if isempty(drefs_idx)
            topos_dc(k)=topovs(k);
          else
            topos_dc(k)=topo_drefs(drefs_idx);
          end
        elseif ~isempty(regexp(topos_thres{k},'up','once'))
          drefs_idx=find(topovs(k)<=topo_drefs,1,'first');
          if ~isempty(drefs_idx)
            topos_dc(k)=topo_drefs(drefs_idx);
          else
            topos_dc(k)=max(topo_drefs);
          end
        elseif ~isempty(regexp(topos_thres{k},'down','once'))
          drefs_idx=find(topovs(k)>=topo_drefs,1,'last');
          if ~isempty(drefs_idx)
            topos_dc(k)=topo_drefs(drefs_idx);
          else
            topos_dc(k)=min(topo_drefs);
          end
        end
      end

      % discretization of topology if all criteria are met
      x0_map=[x0_map; containers.Map(parsopt_topo,topos_dc)];
      A_aug.x0_map=x0_map;
      sm_act=A_aug.sm_act;
      topos_d=nan(size(topos_dc));
      l_d=false(size(topos_dc));
      topos_act_fdisc=nan(size(topos_dc));
      l_netdeg=all(any(sm_act.prod-sm_act.react<-0.01,2)); % all comp have 1 net degradation reaction
      for k=1:numel(parsopt_topo)
        if l_netdeg && any(topos_dc(k)==topo_drefs) && ...
            (isa(topos_thres{k},'numeric') || isempty(regexp(topos_thres{k},'measure','once')))
          l_d(k)=true;
        elseif any(topos_dc(k)==topo_drefs) && ...
            isa(topos_thres{k},'char') && ~isempty(regexp(topos_thres{k},'measure','once'))
          l_d(k)=true;
        else
          l_d(k)=false;
        end
        if l_d(k)
          topos_d(k)=topos_dc(k);
          topos_act_fdisc(k)=topos_act(k)-topos_d(k);
        elseif ~l_d(k)
          topos_d(k)=topovs(k);
          topos_act_fdisc(k)=0;
        else
          error('This scenario should NOT occur!');
        end
      end

      % ensure A_aug.x0 is unmodified
      A_aug.x0_map=x0_map_orig;
    end

    function gd_reset(A_aug,pars_opt_topo)
      % A_aug.param('xnorm_thres')=100;
      % reset dobj_<par_opt>_dval_j to appropriate values
      for k=pars_opt_topo
        key=k{:};
        paropt_start=regexprep(key,'^(sm_[a-z]+)(\d+_\d+)$','$1_start_$2');
        paropt_disc=regexprep(key,'^(sm_[a-z]+)(\d+_\d+)$','$1_disc_$2');
        A_aug.param(paropt_start)=0;
        A_aug.param(paropt_disc)=0;
        A_aug.param(['dobj_',key,'_dval_1'])=0;
        A_aug.param(['dobj_',key,'_dval_2'])=1;
        A_aug.param(['dobj_',key,'_dval_3'])=2;
      end
    end

    function gd_discretize(A_aug,par_opt,status_d)
      % Modify A_aug to ensure discretized topology is truly discrete depending on the topological
      % parameter.
      %
      % INPUT
      % status_d - 0 means decrease to first possible discrete value; 1 means increase to first possible
      %   discrete value
      val_orig=A_aug.x0_map(par_opt);
      paropt_start=regexprep(par_opt,'^(sm_[a-z]+)(\d+_\d+)$','$1_start_$2');
      paropt_disc=regexprep(par_opt,'^(sm_[a-z]+)(\d+_\d+)$','$1_disc_$2');
      A_aug.param(paropt_start)=val_orig;
      A_aug.param(paropt_disc)=1;
      if status_d==0 && val_orig>2
        % do nothing
        1;
      elseif status_d==0 && val_orig>1
        A_aug.param(['dobj_',par_opt,'_dval_3'])=A_aug.param(['dobj_',par_opt,'_dval_1']);
      elseif status_d==0 && val_orig<1
        A_aug.param(['dobj_',par_opt,'_dval_2'])=A_aug.param(['dobj_',par_opt,'_dval_1'])-...
          abs(A_aug.param(['dobj_',par_opt,'_dval_2'])-A_aug.param(['dobj_',par_opt,'_dval_1']));
      elseif status_d==1 && val_orig<0
        % do nothing
        1;
      elseif status_d==1 && val_orig<1
        A_aug.param(['dobj_',par_opt,'_dval_1'])=A_aug.param(['dobj_',par_opt,'_dval_3']);
      elseif status_d==1 && val_orig>1
        A_aug.param(['dobj_',par_opt,'_dval_2'])=A_aug.param(['dobj_',par_opt,'_dval_3'])+...
          abs(A_aug.param(['dobj_',par_opt,'_dval_2'])-A_aug.param(['dobj_',par_opt,'_dval_3']));
      end
    end
    function dtopo = discretize_bf(topo)
      % INPUT
      % topo - topological parameter
      % OUTPUT
      % dtopo - discretized topological parameter
      weights=[0.5,1,0.5]; % ensure all topo state given equal weighting
      dtopo=extract_farray([0,1,2],...
        find( weights.*abs(topo-[0,1,2])==min(weights.*abs(topo-[0,1,2])) , 1 ));
    end
    function l=istrivialss(xss)
      % INPUTS
      % xss - vector
      l=sqrt(sum(xss(2:end).^2))<1e-8 || xss(1)<1e-3 || ... % trivialss criteria
        any(xss<-1e-5); % out of bound detection
    end
    function l = sm_rules_satisfied(sm_react,sm_prod)
      % See if stoimetric matrix efficients rules are satisfied
      % * Rule 1: zeroth order degradation - a product entry can only be negative when 
      % reactant entry at identical position is 0. 
      % INPUT
      % sm_react - stoichoimetric matrix for the reactants; numeric; 
      % sm_prod - stoichoimetric matrix for the products; numeric;
      % OUTPUT
      % l - logical scalar
      l=all(sm_react(sm_prod<0)==0);
    end
    function [l,l_r,l_c,l_q,qT,qTS1tom] = sm_gk_satisfied(sm_react,sm_prod,m)
      % INPUT
      % sm_react - stoichoimetric matrix for the reactants; numeric; 
      % sm_prod - stoichoimetric matrix for the products; numeric;
      % m - first m reactions of sm_react and sm_prod are involved in the
      %  set point. l_c is only valid for m=2
      % OUTPUT
      % l - logical scalar indicating whethter GK's conditions are satisfied
      % l_r - logical scalar indicating whether GK's rank condition is satisfied. 
      % l_c - logical scalar indicating whether GK's first condition
      %  regarding controller's reactions is satisfied
      % l_q - logical scalar indicating whether GK's second condition is satisifed
      % qT - row vector qT or matrix as stated in Gupta and Khammash (2022); 
      % qTS1tom - row vector or matrix as stated in Gupta and Khammash
      
      sm=sm_prod-sm_react;
      %%%% see if rank(sm)=size(sm,1)
      l_r=(rank(sm)==size(sm,1));

      %%%% GK1: see if it is satisfied; sm_react(1,1)~=sm_react(1,2) when m=1 AND
      %  the effect of x2ss(u) to xNss(u) on qT x S x c = 0 has to be none or separable
      %  for each set point encoding parameter associated mass action.
      %
      %  The condition is generalized for m>2
      l_c1=m>2 || ... 
        ... % GK1.1: condition in paper but cannot find justification for it
        sm_react(1,1)~=sm_react(1,2);
      ij_comb=nchoosek(1:m,2); % matrix w 2 columns
      for r=1:size(ij_comb)
        % Here, I'm assuming the effect is separable for only 2 set point encoding 
        % parameter associated mass actions
        l_c=l_c1 && all(sm_react(2:end,ij_comb(r,1))==sm_react(2:end,ij_comb(r,2)));
        if l_c; break; end
      end

      %%%% GK2: see if a non-zero q exists such that qT x S(:,m+1:K) = 0 AND 
      %  qT x S(:,1:m) have opposite signs when m=2
      q=null(sm(:,m+1:end).');
      qT=q.';
      if numel(q)>=1 
        qTS1tom=qT*sm(:,1:m); % row vector if qT is only a vector
        if m>2 || ... 
            ... % GK2.2: opposite signs in qTS1to2 when m=2
            ... % Since I cannot assume that x1ss=pfi(p1/p2) for m>2, it is difficult to
            ... % generalize this condition to m>2
            any(abs(sum(qTS1tom,2))<sum(abs(qTS1tom),2)) 
          l_q=true;
        else
          l_q=false;
        end
      else
        l_q=false;
        qT=zeros(1,size(sm_prod,1));
        qTS1tom=zeros(1,m);
      end     

      l=l_r && l_c && l_q;
    end
    function [sm_react,sm_prod,iter] = sm_gk_gen(n_sv,n_rxns,m,sm_react0,sm_prod0,mode)
      % Assuming the number of reaction involved in set point is 2.
      % Currently, the maximum value of each entry in the stoichiometric
      % matrix is 2.
      % 
      % INPUT
      % n_sv - numeric scalar; number of state variables
      % n_rxns - numeric scalar; number of reactions
      % m - number of first reactions to encode the set point
      % sm_react0 - numeric matrix; nan indicate entries that can be randomly chosen
      % sm_prod0 - numeric matrix; nan indicate entries that can be randomly chosen
      % mode - 'both+rank-bimolecular'|'both+rank'|'qT only'|'none*'; 
      %  specifies which Gupta and Khammash (2022)'s criteria need to be satisfied. 
      % OUTPUT
      % sm_react,sm_prod - stoichoimetric matrices
      % iter - number of random parameters sampled before suitable parameter is found
      
      % customization section
      max_smentry=2; % 2 takes account autocatalytic networks
      
      % generate feasible sm_react and sm_prod
      max_smentry_rand=max_smentry+1;
      row_ids=1:n_sv;
      iter=0;
      while true
        iter=iter+1;
        sm_react=floor(rand([n_sv,n_rxns])*max_smentry_rand);
        sm_react(~isnan(sm_react0))=sm_react0(~isnan(sm_react0));
        sm_prod=floor(rand([n_sv,n_rxns])*max_smentry_rand);
        sm_prod(~isnan(sm_prod0))=sm_prod0(~isnan(sm_prod0));
        if ~isempty(regexp(mode,'none','once'))
          break; 
        elseif strcmp(mode,'qT only') || ~isempty(regexp(mode,'both','once'))
          if ~isempty(regexp(mode,'both','once'))
            % this block should be before bimolecular code block, bc the
            % later code block assumes modification is final. 
            if sm_react(1,1)==sm_react(1,2) && m==2
              continue;
            else
              for c=2:m
                sm_react(2:end,c)=sm_react(2:end,1);
              end
            end
            
            %
            if ~isempty(regexp(mode,'bimolecular','once'))
              for c=1:n_rxns
                % sel is used to select for entries that can be anything
                % sel selects up to two entries
                sel_rnd=randperm(n_sv);
                sel=[];
                if c<=m 
                  % sel must include 1 in first position for sm_react(2:end,1)=
                  % sm_react(2:end,2)
                  sel=[1,sel]; 
                end
                sel=unique([ sel, find( ~isnan(sm_react0(:,c)) & sm_react0(:,c)~=0 )]);
                if numel(sel)>2
                  error('RNetwork:sm_gk_gen:sm_react0',...
                    'sm_react already violate bimolecular constraints!')
                end
                sel_rnd=setdiff(sel_rnd,sel,'stable');
                sel=[sel,sel_rnd( 1:2-numel(sel) )];
                if c>1 && c<=m
                  sel=sel_react; % sel for col=c be same as col=1. 
                end
                sel_react=sel; % logging purpose
                sm_react(setdiff(row_ids,sel),c)=0;
                if sum(sm_react(:,c))>2
                  % eliminate one of the two entries to be zero if sum of entries > 2
                  if c<=m
                    sel_elim=2;
                  else
                    sel_elim=ceil(rand(1)*2);
                  end
                  if ~isnan(sm_react0(sel(sel_elim),c)) && sm_react0(sel(sel_elim),c)~=0
                    sel_elim=setdiff([1,2],sel_elim);
                    if ~isnan(sm_react0(sel(sel_elim),c)) && sm_react0(sel(sel_elim),c)~=0
                      continue;
                    end
                  end
                  sm_react(sel(sel_elim),c)=0;
                end

                sel_rnd=randperm(n_sv);
                sel=[];
                sel=unique([ find( ~isnan(sm_prod0(:,c)) & sm_prod0(:,c)~=0 ),sel ]);
                if numel(sel)>2
                  error('RNetwork:sm_gk_gen:sm_prod0',...
                    'sm_prod0 already violate bimolecular constraints!')
                end
                sel_rnd=setdiff(sel_rnd,sel,'stable');
                sel=[sel,sel_rnd( 1:2-numel(sel) )];
                sm_prod(setdiff(row_ids,sel),c)=0;
                if sum(sm_prod(:,c))>2
                  % eliminate one of the two entries to be zero if sum of entries > 2
                  sel_elim=ceil(rand(1)*2);
                  if ~isnan(sm_prod0(sel(sel_elim),c)) && sm_prod0(sel(sel_elim),c)~=0
                    sel_elim=setdiff([1,2],sel_elim);
                    if ~isnan(sm_prod0(sel(sel_elim),c)) && sm_prod0(sel(sel_elim),c)~=0
                      continue;
                    end
                  end
                  sm_prod(sel(sel_elim),c)=0;
                end
              end
            end
          end
          [l,l_r,l_c,l_q,qT]=RNetwork.sm_gk_satisfied(sm_react,sm_prod,m); % self-testing
          if ~isempty(regexp(mode,'both','once'))
            if l 
              break;
            end
          else
            if l_q
              break; 
            end
          end
        end
      end
    end
  end
end
```
