## Supplemental Data 1 for "Principles underlying implementation of *nearly*-homeostatic biological networks": readme

BioSystem Suite installation instructions: 
- add the BioSystem Suite folder and its subfolder "utilities" to Matlab (>=2022)'s search path
- use showdemo @BioSystem/BioSystem to get a technical overview of the program. 
- If the installation is successful, run(BioSystemTests_Base,'utilities_test') should run successfully. 
