## Supplemental Data 1 for "Principles underlying implementation of *nearly*-homeostatic biological networks": Map2.m

```
classdef Map2 < containers.Map
  % Extension of containers.Map that allows locking of contents inside containers.Map 
  % handle. There are two types of locks. When key lock is on, this class will prevent changing
  % key contents. When all lock is on, this class will prevent change in any content of the
  % map container.
  % Design
  % * lock all should trigger key locking as well as lock of rest of
  %   contents; unlocking unlocks rest of the contents as well 
  % * When key lock is on, an intended set of keys is recorded. If container map is modified,
  %   new keys are compared to the intended set. Error is triggered when the two sets differ.
  %   There two methods for container.Map to modify its content: *remove* and *subsasg*.
  properties (Dependent)
    locked_k % status of key lock
    locked_a % status of overall lock
  end
  properties (Hidden,Access=private)
    s_locked_k=false
    s_locked_a=false
  end
  properties (Hidden)
    keys_intd % intended set of keys when key lock is on
  end
  
  methods
    function obj = Map2(varargin)
      obj=obj@containers.Map(varargin{:});
    end 
    function value=get.locked_k(obj)
      value=obj.s_locked_k;
    end
    function set.locked_k(obj,value)
      if value
        obj.keys_intd=obj.keys;
      else
        obj.s_locked_a=false;
      end
      obj.s_locked_k=value;
    end
    function value=get.locked_a(obj)
      value=obj.s_locked_a;
    end
    function set.locked_a(obj,value)
      obj.locked_k=value;
      obj.s_locked_a=value;
    end
    function varargout=remove(varargin)
      obj=varargin{1};
      if obj.locked_k; error('Map2:klocked','Map keys are locked!'); end
      varargout=cell(1,nargout);
      [varargout{1:end}]=remove@containers.Map(varargin{:});
    end
    function varargout=subsasgn(varargin)
      obj=varargin{1};
      varargout=cell(1,nargout);
      if strcmp(varargin{2}.type,'.') % taking into account map2.KeyType='any'
        [varargout{:}]=builtin('subsasgn',varargin{:});
      else
        if obj.locked_a % faster if placed first
          error('Map2:alocked','All of map container is locked!');
        elseif obj.locked_k && ...
            (obj.length~=length(obj.keys_intd) || ...
            ~all(strcmp(obj.keys,obj.keys_intd)))
          error('Map2:klocked','Map keys are locked!');
        end
        [varargout{1:end}]=subsasgn@containers.Map(varargin{:});
      end
    end
    function varargout=vertcat(varargin)
      varargout=cell(1,nargout);
      [varargout{:}]=vertcat@containers.Map(varargin{:});
      if nargout>0
        varargout{1}.locked_a=false;
        varargout{1}.locked_k=false;
      end
    end
  end
end
```
