## Supplemental Data 1 for "Principles underlying implementation of *nearly*-homeostatic biological networks": action.m

```
function[f]=action(type)
%generate generic regulation function based on the type of specified regulation
%INPUTS
%type - character; uplinear | up | down
%OUTPUTS
%f - symbolic function of three variables [x,Kd,n]. 
% * uplinear: 1/K*x
% * up:positive hill function mapping to [0,1)
% * down: repression hill function mapping to [0,1)

syms x V Kd n
if strcmp(type,'uplinear')
  f(x,Kd,n)=1/Kd*x^n;
elseif strcmp(type,'up')
  f(x,Kd,n)=x^n/(Kd^n+x^n);
elseif strcmp(type,'down')
  f(x,Kd,n)=Kd^n/(Kd^n+x^n);
else
  error('action:action type must be correctly specified!')
end
end
```
