## Supplemental Data 1 for "Principles underlying implementation of *nearly*-homeostatic biological networks": action_to_fname.m

```
function filename=action_to_fname(action_params)
%convert an action parameter to a file name
%Inputs:
%action_param - cell array of regulatory parameters 
%    {'<action1>', '<action2>'[;'<action2>', '<action4>']}
%Outputs:
%filename -  character array. '<action1>_<action2>-<action3>_<action4>'
rows=cell(size(action_params,1),1);
for j=1:length(rows)
    rows{j}=strjoin(action_params(j,:),'_');
end
filename=strjoin(rows.','-');
end
```
