## Supplemental Data 1 for "Principles underlying implementation of *nearly*-homeostatic biological networks": cell2map.m

```
function[map]=cell2map(c)
%converts an n x 2 cell array to container map2. The first
%   column are names of container Map2. The second column are
%   values of container Map2
if isa(c,'cell')
    s=size(c);
    if length(s)==2&& s(2)==2
        map=containers.Map2(c(:,1),c(:,2));
    else
        map=containers.Map2();
    end
else
    error('input must be a cell array with n rows and 2 columns!')
end
end
```
