## Supplemental Data 1 for "Principles underlying implementation of *nearly*-homeostatic biological networks": cof.m

```
function cof=cof(a)
% 
%     cof=cof(A) generates matrix of cofactor values for an M-by-N matrix
%     A    : an M-by-N matrix
% 
%     Example: Find the cofactor matrix for A.
% 
%     A = 1     3     1
%         1     1     2
%         2     3     4
% 
%     >>cof=cof(A)
% 
%     cof =
% 
%         -2     0     1
%         -9     2     3
%          5    -1    -2
%
%
%   Angelica Ochoa
%   Boston University
%  
%   Last Modified: 8-Feb-2012

%% Check Input Argument
%----------------------
if isempty(a)
    error(message('cof:EmptyMatrix'));
end

%% Algorithm 
%-----------
[r, c] = size(a);    %determine size of input
if isa(a,'sym')
    m=sym(ones(r,c));
elseif isa(a,'num')
    m = ones(r,c);      %preallocate r x c cofactor matrix 
else
    error('format of input is incorrect!');
end
a_temp=a;           %create temporary matrix equal to input
for i = 1:r
    for k = 1:c
        a_temp(i,:)=[];   %remove ith row
        a_temp(:,k)=[];   %remove kth row
        m(i,k) = ((-1)^(i+k))*det(a_temp);  %compute cofactor element
        a_temp=a;   %reset elements of temporary matrix to input elements
    end  
end

cof=m;  %return cofactor matrix as output variable

end
```
