## Supplemental Data 1 for "Principles underlying implementation of *nearly*-homeostatic biological networks": compute_dependencies.m

```
% Given a container map of expressions (defs). For each key, determine all symbols that are used to 
% compute that key.
% INPUTS
% defs - a map container of {'name',possibly non-scalar symbolic expression}
% OUTPUT
% dependencies - a map container where values are all the symbols in a column cell array of characters 
%   that are used to compute the key. 
function [dependencies]=compute_dependencies(defs)
defVars = defs.keys;
dependencies = containers.Map('KeyType','char','ValueType','any');
for i = 1:numel(defVars)
  v = defVars{i};
  expr = defs(v);
  depNames = symvar_all(char_allsyms(expr));
  dependencies(v) = depNames;
end
end

function out = char_allsyms(s)
% find character form of symbolic expression, including symmatrix or
% symfunmatrix
%%% sym.disp and symbolicmatrix.disp but not symvar_all takes the longest time in 
%%% order_fluxvars
if isa(s,'sym')
  out=char(s);
else 
  out=evalc('disp(s)');
end
end 
```
