## Supplemental Data 1 for "Principles underlying implementation of *nearly*-homeostatic biological networks": convert2_absthres.m

```
function [absthres] = convert2_absthres(thres,y)
% Convert a thres vector that is the same size as y to absolute threshold vector based on values of
% y. 

abs_y=abs(y);
thres=abs(thres);
absthres=max(thres.*abs_y,thres);
end
```
