## Supplemental Data 1 for "Principles underlying implementation of *nearly*-homeostatic biological networks": dropnan.m

```
function [out] = dropnan(x,rep)
% Suppose x is a number, then rep is a number also. If x is not nan, then 
% dropnan is the identity function. Otherwise, rep value is the output. 
% Suppose x is a sym, then rep has to be a sym also. 
if isa(x,'sym')
  out=privBinaryOp(x,sym('dropnan'),'symobj::map',char(rep));
elseif ~isnan(x)
  out=x;
else
  out=rep;
end
end
```
