## Supplemental Data 1 for "Principles underlying implementation of *nearly*-homeostatic biological networks": ensure_form.m

```
function out =ensure_form(in,form)
%ensure that the input is a specific data format.
%INPUTS:
%in - input container map or lowest class array whose index is capable of
%   separating elements assumed to be the same class. 
%   For example, if characters are used, the
%   appropriate array to use to denote different elements would be
%   cell array. Matrix is used wherever possible; no cell array of numbers.
%form - 'numeric' or 'symbolic' or 'character'
%OUTPUTS
%out - container map subclass or lowest (matrix<cell) array whose index is capable of
% separating elements. If in is a container map, then out may be an alias of in.
if isa(in,'containers.Map')
  switch form
    case 'symbolic'
      out=containers.Map2(in.keys,cell(1,in.Count));
      for k=keys(in)
        if isa(in(k{:}),'char')
          out(k{:})=str2sym(in(k{:}));
        elseif ~isempty(regexp(class(in(k{:})),'sym','once'))
          out(k{:})=in(k{:});
        else
          out(k{:})=sym(in(k{:}));
        end
      end
      if isa(in,'containers.Map2')
        out.locked_a=in.locked_a;
        out.locked_k=in.locked_k;
      end
    case 'numeric'
      if strcmp(in.ValueType,'double')
        out=in;
      else
        out=containers.Map2();
        for k=keys(in)
          out(k{:})=double(in(k{:}));
        end
      end
    case 'character'
      if strcmp(in.ValueType,'char')
        out=in;
      else
        out=containers.Map2();
        for k=keys(in)
          out(k{:})=char(in(k{:}));
        end
      end
    otherwise
      error('form specification is not correct');
  end
else
  switch form
    case 'symbolic'
      if isa(in,'char') || isa(in,'cell')
        try
          out=str2sym(in);
        catch
          out=sym(in);
        end
      elseif ~isempty(regexp(class(in),'sym','once'))
        out=in;
      else
        out=sym(in);
      end
    case 'numeric'
      out=double(in);
    case 'character'
      out=cell(size(in));
      for j=1:numel(in)
        out{j}=char(in(j));
      end
    otherwise
      error('form specification is not correct');
  end
end
```
