## Supplemental Data 1 for "Principles underlying implementation of *nearly*-homeostatic biological networks": fname_to_action.m

```
function action_params=fname_to_action(filename)
%parse filename into action parameters
%Inputs:
%filename -  character array. '<action1>_<action2>-<action3>_<action4>'
%Outputs:
%action_param - cell array of regulatory parameters 
%   {'<action1>', '<action2>'[;'<action2>', '<action4>']}

filename=regexprep(filename,'\.csv','');
row=strsplit(filename,'-').';
col=strsplit(row{1},'_');
action_params=cell(length(row),length(col));
for j=1:length(row)
    action_params(j,:)=strsplit(row{j},'_');
end
end
```
