## Supplemental Data 1 for "Principles underlying implementation of *nearly*-homeostatic biological networks": func2mex.m

```
function [mex_f]=func2mex(func_f,argspecs)
% Converts a local function to a mex file using Matlab Coder. Unit tested within BioSystemTest_Base
% INPUTS
% func_f - function file name without suffix or directory prefix; the file should be present in 
%   the working directory
% argspecs - a character vector of input specifications for the function specified by func_f. Must
%   be of the form '{...}' to be put directly after -args. 
% 
% OUTPUTS
% mex_f - function file name without .mex suffix or directory prefix

%% input checking
if ~exist(func_f,"file")
  error('func2mex:func_f','file not found!');
end

%% setting up for code generation 
mex_f=new_fname('.',[regexprep(func_f,'\.','_'),'_'],'');
mex_f=regexprep(mex_f,'^\./','');
mex_config=coder.config('mex');
mex_config.TargetLang="C++";
mex_config.InlineBetweenUserFunctions='Readability';
mex_config.InlineBetweenUserAndMathWorksFunctions='Readability';
mex_config.InlineBetweenMathWorksFunctions='Readability';
codegen_instr=['codegen -config mex_config -o ',mex_f,' ',func_f,' -args ',argspecs];

%% mex file generation
eval(codegen_instr);
end
```
