## Supplemental Data 1 for "Principles underlying implementation of *nearly*-homeostatic biological networks": gen_form_sstep.m

```
function [topo_act,b1,b2,b1_nkbl,b1_nkbr,b2_nkbl,b2_nkbr] = gen_form_sstep(var,d,n,k,s)
% Computes the actual topology (topo_act, b1, b2) based on supplied inputs. See the mnanuscript for
% more details. 
%
% Tested by the GeneticNetwork's constructor and therefore XNetworks. 
%
% INPUTS
% var - character vector; name of topological variable that is used to compute the actual topology 
%   for; only used if s is symbolic.  
% d,n,k,s - symbolic or numeric scalar
% OUTPUTS
% topo_act, b1, b2, b1_nkbl, b1_nkbr, b2_nkbl, b2_nkbr - symbolic or numeric scalar; if the output
%   is symbolic, recomputation is minimized using new symbolic variables form by [var,'_b1_nkbl']. 

if isa(s,'sym')
  zero_var=sym(0);
  neight_var=sym(-8);
  one_var=sym(1);
  onep3_var=sym(1.3);
  ten_var=sym(10);
  two_var=sym(2);
else
  zero_var=0;
  neight_var=-8;
  one_var=1;
  onep3_var=1.3;
  ten_var=10;
  two_var=2;
end
b1_nkbl=heaviside_fast(d-0.5)*smoothstep_shifted_Rnoband(n,k,zero_var) + ...
  (1-heaviside_fast(d-0.5))*smoothstep_shifted_Rnoband(n,k,neight_var);
b1_nkbr=smoothstep_shifted_Rnoband(n,k,one_var);
b2_nkbl=smoothstep_shifted_Rnoband(n,2-k,one_var);
b2_nkbr=heaviside_fast(d-0.5)*smoothstep_shifted_Rnoband(n,2-k,two_var) + ...
  (1-heaviside_fast(d-0.5))*smoothstep_shifted_Rnoband(n,2-k,ten_var);
if isa(s,'sym')
  b1_nkbl_name=sym([var,'_b1_nkbl']);
  b1_nkbr_name=sym([var,'_b1_nkbr']);
  b2_nkbl_name=sym([var,'_b2_nkbl']);
  b2_nkbr_name=sym([var,'_b2_nkbr']);
else
  b1_nkbl_name=b1_nkbl;
  b1_nkbr_name=b1_nkbr;
  b2_nkbl_name=b2_nkbl;
  b2_nkbr_name=b2_nkbr;
end
b1=(smoothstep_shifted_Rnoband(n,k,s)-b1_nkbl_name)/(b1_nkbr_name-b1_nkbl_name) - ...
  ((1-b1_nkbl_name)/(b1_nkbr_name-b1_nkbl_name)-1)*smoothstep_shifted_Rnoband(one_var,onep3_var,s);
b2=(smoothstep_shifted_Rnoband(n,2-k,s)-b2_nkbl_name)/(b2_nkbr_name-b2_nkbl_name);
if isa(s,'sym')
  b1_name=sym([var,'_b1']);
  b2_name=sym([var,'_b2']);
else
  b1_name=b1;
  b2_name=b2;
end
topo_act=b1_name+b2_name;
end
```
