## Supplemental Data 1 for "Principles underlying implementation of *nearly*-homeostatic biological networks": genp_mn.m

```
function d = genp_mn( T )
%x_N+1 is the perturbation. 
%x_N+1+1 to x_N+1+N denote basal activator concentration.
%x_N+1+N+1 to x_N+1+N+N denote basal repressor concentration.
%INPUTs
%T - N x N topological cell array of regulatory interactions. ij-th entry represents the
% regulation of xi by xj. Each entry can either 'up'|'down'|'none'
%OUTPUTS
%d - a process of class Metabolic network, ready for steady state simulations. Same format as
% in Ma et al.

%% computing input for MNetwork
N=size(T,1); U_idx=N+1;
IF=cell(N,N+1+2*N); OF=cell(N,N+1+2*N); %[1,2,3,in,basals]
IF(:,U_idx:end)={'no'}; OF(:,U_idx:end)={'no'}; IF{1,U_idx}='yes';
for j=1:numel(T)
  if strcmp(T{j},'up')
    IF{j}='yes'; OF{j}='no';
  elseif strcmp(T{j},'down')
    IF{j}='no'; OF{j}='yes';
  elseif strcmp(T{j},'none')
    IF{j}='no'; OF{j}='no';
  else
    error('simulation:genp_mn','unexpected regulation type encountered!'); 
  end
end
for r=1:size(T,1)
  if ~any(strcmp(IF(r,1:U_idx),'yes'))
    IF{r,U_idx+r}='yes';
  end
  if ~any(strcmp(OF(r,1:U_idx),'yes'))
    OF{r,U_idx+N+r}='yes';
  end
end

%% build network
d=MNetwork(IF,OF); nnodes=d.sys.Count; charnp1=num2str(nnodes+1); 
take_limit(d,{'dn','n'},{1,1},true);
d.prange.remove(['x',charnp1]);
d.param.remove(['x',charnp1]); 
d.output_label={'o1'};
d.output='x3';
end
```
