## Supplemental Data 1 for "Principles underlying implementation of *nearly*-homeostatic biological networks": GN_eg.m

```
function[A]=GN_eg(topo)
% Specialized transcriptional network. 
% INPUTS
% topo - character vector denote topology of the network 
% <u|up|d|down|n|none|un|unknown>_...
% OUTPUTS
% GeneticNetwork object of N nodes, with u and Vd1 being the input and x1 being output. 
topo=fname_to_action(topo);
nnodes=size(topo,1);
reg_map=cell2map({'u','up';'up','up';'n','none';'none','none';
  'd','down';'down','down';'un','unknown';'unknown','unknown'});
A=GeneticNetwork(reg_map.values(topo),repmat({'normal'},[1,nnodes]));
A.flux('H1_1')=A.flux('H1_1')*sym('u');
A.prange.remove('Vd1');
A.param.remove('Vd1');
A.output='x1';
A.output_label={'o1'};
%%% replace is much faster than take_limit
% take_limit(A,{'n','dn'},{1,1},true);
params=A.param.keys;
for i=1:numel(params)
  if ~isempty(regexp(params{i},'^dn|^n', 'once'))
    A.replace(params{i},'1');
  end
end
end
```
