## Supplemental Data 1 for "Principles underlying implementation of *nearly*-homeostatic biological networks": heaviside_fast.m

```
function y=heaviside_fast(x)
% Fast version of heaviside that assumes heaviside(0)=1
% INPUT
% x - numeric or symbolic
% OUTPUT
% y - double precision number
if isa(x,'sym')
  heaviside_fast=nan;
  syms heaviside_fast(a1)
  y=heaviside_fast(x);
else
  y=double(x>=0);
end
end
```
