## Supplemental Data 1 for "Principles underlying implementation of *nearly*-homeostatic biological networks": hill_fast.m

```
function [y] = hill_fast(x,thres,band)
% The function is 0 below thres, grows linearly from 0 to 1 between
% x=[thres,thres+band], and stays at 1 beyond thres+band. 
% INPUT
% x - number
% thres - number
% band - positive number
x_b_thres_p_band=x-(thres+band);
x_b_thres=max(x-thres,0);
% x >=thres && x <=thres_p_band
y=heaviside_fast(x_b_thres.*-x_b_thres_p_band).*x_b_thres./band+...
  ...% x>thres_p_band rather than x>=thres_p_band
  (1-heaviside_fast(-x_b_thres_p_band)); % 
end
```
