## Supplemental Data 1 for "Principles underlying implementation of *nearly*-homeostatic biological networks": liebracket.m

```
function ad_fng = liebracket(f,g,x,n)
% LIEBRACKET  (Nth order iterated Lie bracket of f and g)
%
% Usage:
% ad_fng = liebracket(f,g,x,n)
%
% Input:
% f = symbolic vector field of length l1 (first operand)
% g = symbolic vector field of length l1 (second operand)
% x = symbolic vector of variables
% n = order of the Lie Bracket
%
% Output:
% ad_fng =  [ g    [f,g]  [f,[f,g]] [f,[f,[f,g]]] .... ]
%        n=   0      1         2         3
%             g    adf_g   adf2_g      adf3_g
%
%
% Note: define the symbolic variables as 
%       x1 =sym(x1,'real')
%       in order to get rid of the conjugates
%       (only if they are reals).
%
% Author: Atakan Varol
% Date: 03.23.2006

ad_fng = sym(zeros(length(f),n+1));
ad_fng(:,1) = g;

if n>0
    for t = 2:n+1
        ad_fng(:,t) = jacobian(ad_fng(:,t-1) ,x)*f - jacobian(f,x)*ad_fng(:,t-1);
    end
end

ad_fng = expand(ad_fng);
end
```
