## Supplemental Data 1 for "Principles underlying implementation of *nearly*-homeostatic biological networks": liederivative.m

```
function[l_fnh]=liederivative(h,f,x,n)
%finds lie derivative of scalar with respect along direction of vector
% field
%input:
%h - N to 1 scale field
%f - N to N vector field
%x - column vector of variables of length N to specify the domain
%n - number of iterative derivatives to take. 
l_fnh=h;
for i=1:n
    l_fnh=simplify(jacobian(l_fnh,x)*f);
end
end
```
