## Supplemental Data 1 for "Principles underlying implementation of *nearly*-homeostatic biological networks": lm_f_fmincon_gen.m

```
function [outf]=lm_f_fmincon_gen(iter_diff,sens_thres)
% Generates an output function for fmincon that takes in (x,optimValues,status) and outputs
% whether or not the search should be stopped. Only iteration and fval fields of optimValues
% will be used. The function automatically detects start of a new search based on
% optimValues.iteration=1, so it is always valid. 
%
% INPUTS
% iter_diff - If rsenso1 diff is large for a number of iterations after iter_diff from firstmet,
%   then update the first eligible iteration to the latest iteration. Otherwise, if rsenso1 diff
%   is small for one iter_diff later iteration, pause simulation.
% sens_thres - threshold for relative sensitivity
% OUTPUTS
% a function that takes ~, optimValues as a struct with iter and fval fields, status; returns true
% or false.

lmfirstmet_g_iterdiff_max=min(ceil(iter_diff/100),10);

% resettable parameters
firstmet=1;
firstmet_objv=0;
lmfirstmet_g_iterdiff=0;

outf=@output_f;

  function stop = output_f(~,optimValues,status)
    stop=false;
    if strcmp(status,'iter')
      lastsimindex=optimValues.iteration;
      objv=optimValues.fval;

      % detect rsenso1 fluctuation near sens_thres
      if abs(objv)<1.05*sens_thres
        rsenso1_absdiff_thres=sens_thres;
      else
        rsenso1_absdiff_thres=min(1e-3,sens_thres);
      end

      % see whether simulation should be pasued.
      if lastsimindex==1
        firstmet=lastsimindex;
        firstmet_objv=objv;
        lmfirstmet_g_iterdiff=0;
        lastsimindex_stored=0;
      elseif lastsimindex-firstmet>iter_diff
        lmfirstmet_g_iterdiff=lastsimindex-(iter_diff+firstmet);
        objvs=[firstmet_objv,objv];
        if abs(diff(log10(abs(objvs))))<1e-3 || abs(diff(objvs))<rsenso1_absdiff_thres
          stop=true;
        elseif lmfirstmet_g_iterdiff>lmfirstmet_g_iterdiff_max
          firstmet=lastsimindex;
          firstmet_objv=objv;
          lmfirstmet_g_iterdiff=0;
        end
      end
    end
  end
end
```
