## Supplemental Data 1 for "Principles underlying implementation of *nearly*-homeostatic biological networks": log10m_trans.m

```
function [forward, reverse, major_b, minor_b] = log10m_trans(Frac)
% Generates log10m transformation functions
% INPUT
% Frac - numeric scalar
% OUTPUT
% forward - @(y)sign(y).*log10(Frac.*abs(y)+1)
% reverse - @(new_y)sign(new_y).*(10.^abs(new_y)-1)./Frac}
% major_b - a function handle that takes minimum (arg 1) and maximum (arg 2) 
%           of graphical range and generates major breaks in original scale
%           that will be labeled. 
% minor_b - a function handle that takes minimum (arg 1) and maximum (arg 2)
%           of graphical range and generates minor breaks in original scale 
%           that will not be labeled. 
forward=@(y)sign(y).*log10(Frac.*abs(y)+1);
reverse=@(new_y)sign(new_y).*(10.^abs(new_y)-1)./Frac;
major_b=@major_bf;
minor_b=@minor_bf;

  function b=major_bf(min_y,max_y)
    b=breaks(min_y,max_y);
  end
  function b=minor_bf(min_y,max_y)
    [~,b]=breaks(min_y,max_y);
  end
  function [major_breaks,minor_breaks] = breaks(min_y,max_y)
    %% Find log10F range
    min_log10F=log10(abs(min_y*Frac));
    min_log10F=max([min_log10F,0]);
    min_log10F=floor( sign(min_y)*min_log10F );
    max_log10F=log10(abs(max_y*Frac));
    max_log10F=max([max_log10F,0]);
    max_log10F=ceil( sign(max_y)*max_log10F );
    % log10F range
    log10F_range=min_log10F:max([floor((max_log10F-min_log10F)/10),1]):max_log10F;
    
    %% generate linear scale minor breaks when log10F range does not completely
    %% fall outside of log10F range of (-1,1)
    if max(log10F_range)<=-1 || min(log10F_range)>=1
      minor_breaks=[];
    else
      max_linear=min([ 10*Frac^-1,max([abs(min_y),abs(max_y)]) ]);
      minor_breaks=0:max_linear/10:max_linear;
      minor_breaks=unique(sort([-minor_breaks,minor_breaks]));
      minor_breaks=minor_breaks(minor_breaks>=min_y & minor_breaks<=max_y);
    end
    
    %% generate log10F breaks for log10F range outside of (-1,1)
    log10F_breaks=log10F_range(log10F_range>=1 | log10F_range<=-1);
    major_breaks=sign(log10F_breaks).*10.^abs(log10F_breaks)./Frac;
    
    %% Supplement log10F breaks with minor breaks if needed
    if isempty(major_breaks) || ...
        max(major_breaks)<=min(minor_breaks) || ...
        min(major_breaks)>=max(minor_breaks)
      major_breaks=[major_breaks,minor_breaks];
    end
    major_breaks=unique(sort(major_breaks));
  end
end
```
