## Supplemental Data 1 for "Principles underlying implementation of *nearly*-homeostatic biological networks": map2cell.m

```
function[c]=map2cell(map)
%converts a container map to n x 2 cell array.
if ~isa(map,'containers.Map')
    error('must have containers.Map as input!')
end
ks=keys(map);
vs=values(map,ks);
c=[ks;vs].';
end
```
