## Supplemental Data 1 for "Principles underlying implementation of *nearly*-homeostatic biological networks": map2map2.m

```
function[map2]=map2map2(map)
% convert containers.Map to upgraded version Map2. If start out with Map2, do not change. 
if strcmp(class(map),'containers.Map')
  if map.length==0
    map2=containers.Map2();
  else
    map2=containers.Map2(map.keys,map.values);
  end
elseif strcmp(class(map),'containers.Map2')
  map2=map;
else
  error('util:map2map2','wrong class input!');
end
end
```
