## Supplemental Data 1 for "Principles underlying implementation of *nearly*-homeostatic biological networks": new_fname.m

```
function n=new_fname(dir,prefix,suffix)
% generate a file name with the right suffix in dir that does not already
% exist
% Parameters
% dir - 1xn character vector; 
% prefix - 1xn character vector
% suffix - 1xn character vector
% Returns
% n - file name
pid=feature('getpid');
i=0;
while exist([dir,'/',prefix,num2str(pid),'_',num2str(i),suffix],'file')
  i=i+1;
end
n=[dir,'/',prefix,num2str(pid),'_',num2str(i),suffix];
if isempty(dir); n=regexprep(n,'^/',''); end
end
```
