## Supplemental Data 1 for "Principles underlying implementation of *nearly*-homeostatic biological networks": next_permutation.m

```
function[vec]=next_permutation(vec,mod)
%given x in Z mod j1 times Z mod j2 times ... times Z mod jN space.
% find the next member with a lower order. 
% if no lower number can be found, returns the original member. 
% e.g. the next_permutation of ([2,1,1,0],[3,3,3,3]) is [2,1,0,2]
%input: 
%vec: vector of numbers
%mod: number that specify which finite ring
%output
%vec: next member of a lower order
if ~isequal(vec,zeros(size(vec)))
    n=numel(vec);
    vec(n)=vec(n)-1;

    for j=fliplr(1:n)
        if borrow(vec(j))
            vec(j)=mod(j)-1;
            vec(j-1)=vec(j-1)-1;
        else
            break
        end    
    end
end
end

function[TF]=borrow(scalar)
if scalar<0
    TF=1;
else
    TF=0;
end
end
```
