## Supplemental Data 1 for "Principles underlying implementation of *nearly*-homeostatic biological networks": numTostr.m

```
function[out]=numTostr(in)
%convert a numeric to characters. If input is not numeric, return original input as output. 
%INPUT
%in - array of expressions
%OUTPUT
%out - cell array of strings.
if isnumeric(in)
    out=num2str(in);
else
    out=in;
end
end
```
