## Supplemental Data 1 for "Principles underlying implementation of *nearly*-homeostatic biological networks": parsim_hs.m

```
function parsim_hs(genp,topos,I,np,subdir)
% Parallel simulation of topologies for understanding of robust homeostasis
% INPUTS
% genp - Function that takes a character vector and output a BioSystem process
% topos - cell array of strings denoting various topologies
% I - matrix of inputs to simulate. Each row is a separate input. 
% np - number of parameters to simulate.
% subdir - character vector specifying sub-directory to output simulation files to. 
% OUTPUTS
% Simulations files - [subdir,'/',topo,'.csv'] and [subdir,'/',topo,'.log']
evalc_w=@(A,s)evalc(s);
parfor j=1:length(topos)
  topo=topos{j};
  fname=[subdir,'/',topo];
  fname_log=[fname,'.log'];
  if ~exist([fname,'.csv'],'file')
    A=genp(topo);
    fid=fopen(fname_log,'w');
    fprintf(fid,[evalc_w(A,'disp(A.vecfun_form);'),'\n']);
    fprintf(fid,[evalc_w(A,'disp(A);'),'\n']);
    fclose(fid);
    find_homeostasis(A,I,A.prange,np,[5000,6000],fname);
  end
end
```
