## Supplemental Data 1 for "Principles underlying implementation of *nearly*-homeostatic biological networks": prange_convert.m

```
function[prange]=prange_convert(prange)
%convert parameters from container map to matrix
%inputs:
%prange - container map; {'<param1>',[<from>,<to>][;...]}
%outputs:
%prange - matrix with 2 rows. Each column corresponds to a different
%   parameter. The order of parameters is the same as the order of keys in the input. 
prange=prange.values(prange.keys);
prange=vertcat(prange{:}).'; %2 x p matrix
end
```
