## Supplemental Data 1 for "Principles underlying implementation of *nearly*-homeostatic biological networks": rand_param.m

```
function[paramset]=rand_param(param_range,n,rndstream)
%Given parameter range in matrix form, generate n low-discrepency
%parameters. If only 1 random number is requested, that number is generated
%based on default seed value. If more than 1 random number is requested,
%those numbers are generated using sobolset.
%Inputs:
%param_range - matrix; first row contains lower limits and second
%   row contains upper limits
%n - number of random parameters requested
%rndstream - OPTIONAL random number stream to draw random numbers from
%Outputs:
%paramset - matrix; each row contains a set of parameters

if nargin<=2
  rndstream=RandStream.getGlobalStream;
end

%input checking
[r,p]=size(param_range);
if r~=2
  error('rand_param:WrongInput','check documentation!');
end
%actual work
if n==1
  paramset=rand(rndstream,1,p);
elseif n>1
  set=sobolset(p,'Skip',1e3,'Leap',1e2);
  paramset=net(set,n);
end
paramset=paramset.*repmat(abs(param_range(2,:)-param_range(1,:)),n,1)+...
  repmat(param_range(1,:),n,1);
end
```
