## Supplemental Data 1 for "Principles underlying implementation of *nearly*-homeostatic biological networks": ref1_array.m

```
classdef ref1_array < handle
  % create a numeric array object where the reference count for the underlying data is 1. 
  %
  % Unit tested by BioSystem_Base's simulate_sys method. 

  properties (Access=private)
    array % underlying storage for the ref1_array object
  end

  methods
    
    function obj = ref1_array(varargin)
      % INPUTS
      % number of elements in each dimension starting from dimension 1

      obj.array = nan(varargin{:},'double');
    end

    function out = read(obj,varargin)
      % Read array
      % Inputs
      % varargin - array indices

      out=obj.array(varargin{:});
    end

    function mod(obj,varargin)
      % Modify array
      % Inputs
      % varargin - array indices followed by replacement array
      
      rep=varargin{end};
      obj.array(varargin{1:end-1})=rep;
    end
    
    function subset(obj,varargin)
      % Subsetting array
      % Inputs
      % varargin - array indices to update the array to. 

      obj.array=obj.array(varargin{:});
    end

    function size_m=size(obj)
      % Get the dimension of the array

      size_m=size(obj.array);
    end

  end

  methods(Hidden) %% internal methods - hiding non-sealed handle class methods
    function varargout=addlistener(varargin)
      varargout=cell(1,nargout);
      [varargout{1:end}]=addlistener@handle(varargin{:});
    end
    function varargout=eq(varargin)
      varargout=cell(1,nargout);
      [varargout{1:end}]=eq@handle(varargin{:});
    end
    function varargout=findobj(varargin)
      varargout=cell(1,nargout);
      [varargout{1:end}]=findobj@handle(varargin{:});
    end
    function varargout=findprop(varargin)
      varargout=cell(1,nargout);
      [varargout{1:end}]=findprop@handle(varargin{:});
    end
    function varargout=ge(varargin)
      varargout=cell(1,nargout);
      [varargout{1:end}]=ge@handle(varargin{:});
    end
    function varargout=gt(varargin)
      varargout=cell(1,nargout);
      [varargout{1:end}]=gt@handle(varargin{:});
    end
    function varargout=le(varargin)
      varargout=cell(1,nargout);
      [varargout{1:end}]=le@handle(varargin{:});
    end
    function varargout=lt(varargin)
      varargout=cell(1,nargout);
      [varargout{1:end}]=lt@handle(varargin{:});
    end
    function varargout=ne(varargin)
      varargout=cell(1,nargout);
      [varargout{1:end}]=ne@handle(varargin{:});
    end
    function varargout=notify(varargin)
      varargout=cell(1,nargout);
      [varargout{1:end}]=notify@handle(varargin{:});
    end
  end
end
```
