## Supplemental Data 1 for "Principles underlying implementation of *nearly*-homeostatic biological networks": ref1_array_struct.m

```
function [methods_struct] = ref1_array_struct (varargin)
% create a numeric array object where the reference count for the underlying data is 1.
array=nan(varargin{:},'double');
methods_struct=struct('read',@read,'mod',@mod,'subset',@subset,'size',@sizeof);

  function out = read(varargin)
    % Read array
    % Inputs
    % varargin - array indices

    out=array(varargin{:});
  end

  function mod(varargin)
    % Modify array
    % Inputs
    % varargin - array indices followed by replacement array

    rep=varargin{end};
    array(varargin{1:end-1})=rep;
  end

  function subset(varargin)
    % Subsetting array
    % Inputs
    % varargin - array indices to update the array to.

    array=array(varargin{:});
  end

  function size_m=sizeof()
    % Get the dimension of the array

    size_m=size(array);
  end

end
```
