## Supplemental Data 1 for "Principles underlying implementation of *nearly*-homeostatic biological networks": reverseAutoDiff.m

```
function [dX_dZ, derivatives, topoOrder] = reverseAutoDiff(defs, outputs, outputs_label, inputs)
% reverseAutoDiff computes the Jacobian of the output array w.r.t. input variables
% using a reverse-mode differentiation approach. Intermediate variables that are not involved in
% outputs are not included in the differentiation process. The reverse differentiation approach also
% minimizes recomputation of intermediate variables. 
%
% INPUTS:
%   defs    : containers.Map from variable names (char) to scalar symbolic expressions
%   outputs : symbolic array
%   outputs_label: cell array of character vector having the same size as outputs
%   inputs  : 1D array of symbolic inputs
%
% OUTPUT:
%   dX_dZ   : symbolic array of partial derivatives, where each slice along the last dimension
%     correspond to a single scalar variable. 
%   derivatives: containers.Map of scalar derivatives of different scalar variables.
%   topoOrder: order of def.keys from least dependent to most dependent

inputNames = arrayfun(@char, inputs, 'UniformOutput', false);
numInputs = length(inputs);
sizeOutputs=size(outputs);


% Compute dependencies, topological order, and defs.keys that are involved in the computation of 
% outputs
dependencies = [compute_dependencies(defs);
  compute_dependencies(containers.Map( outputs_label(:) , num2cell(outputs(:)) ))];
start_keys = intersect(defs.keys,symvar(char(outputs)));
topoOrder = topologicalSort(dependencies,start_keys);
topoOrder=[topoOrder;outputs_label(:)];

% ensure outputs_label is not named as one of its dependencies
allDep=dependencies.values;
if numel(allDep)>0
  allDep=vertcat(allDep{:});
end
if any(ismember(outputs_label,allDep))
  error('reverseAutoDiff:name_conflict',...
    'outputs_label {%s} should NOT contain any variables in its definition!',...
    strjoin(outputs_label,','));
end

% Initialize sensitivities dX/dv: varName -> zeros(sizeOutputs)
sensitivities = containers.Map('KeyType','char','ValueType','any');
allVars = unique([topoOrder; inputNames(:)]);
for i = 1:numel(allVars)
  sensitivities(allVars{i}) = sym(zeros(sizeOutputs));
end

% Initialize sensitivities to dX_dXi
for o = 1:numel(outputs)
  outName = outputs_label{o};
  sensArray = sensitivities(outName);
  sensArray(o) = sym(1);
  sensitivities(outName) = sensArray;
end

% A map to store intermediate derivatives dv/dw
derivatives = containers.Map('KeyType','char','ValueType','any');

% Reverse-mode backprop
revOrder = flip(topoOrder); 
for i = 1:numel(revOrder)
  v = revOrder{i};
  sens_v = sensitivities(v); % dX/dv
  if ismember(v,outputs_label)
    idx=find(strcmp(v,outputs_label),1);
    expr=outputs(idx);
    varsInExpr = symvar(expr);
  else
    expr = defs(v);
    varsInExpr=sym(dependencies(v));
  end
  for j = 1:numel(varsInExpr)
    wSym = varsInExpr(j);
    wName = char(wSym);
    if ~sensitivities.isKey(wName)
      continue;
    end

    % Compute dv/dw if not already stored
    dv_dw_name = ['dl_', v, '_r_dl_', wName,'_r'];
    if ~derivatives.isKey(dv_dw_name)
      dv_dw = diff(expr, wSym);
      derivatives(dv_dw_name) = dv_dw;
    end

    % Update sensitivity: dX/dw += dX/dv * dv/dw
    sens_w = sensitivities(wName);
    sens_w = sens_w + sens_v .* sym(dv_dw_name);
    sensitivities(wName) = sens_w;
  end
end

% Extract final dX/dZ
dX_dZ = sym(zeros([sizeOutputs,numInputs]));
nonlast_dims = repmat({':'},1,ndims(dX_dZ)-1);
for j = 1:numInputs
  inName = inputNames{j};
  if sensitivities.isKey(inName)
    dX_dZ(nonlast_dims{:}, j) = sensitivities(inName);
  else
    dX_dZ(nonlast_dims{:}, j) = sym(0);
  end
end
end
```
