## Supplemental Data 1 for "Principles underlying implementation of *nearly*-homeostatic biological networks": routh.m

```
function RA=routh(poli,epsilon);

%ROUTH   Routh array.
%   RA=ROUTH(R,EPSILON) returns the symbolic Routh array RA for 
%   polynomial R(s). The following special cases are considered:
%   1) zero first elements and 2) rows of zeros. All zero first 
%   elements are replaced with the symbolic variable EPSILON
%   which can be later substituted with positive and negative 
%   small numbers using SUBS(RA,EPSILON,...). When a row of 
%   zeros is found, the auxiliary polynomial is used.
%
%	Examples:
%
%	1) Routh array for s^3+2*s^2+3*s+1
%
%		>>syms EPS
%		>>ra=routh([1 2 3 1],EPS)
%		ra =
%
% 		   1.0000    3.0000
% 		   2.0000    1.0000
% 		   2.5000         0
% 		   1.0000         0
%
%	2) Routh array for s^3+a*s^2+b*s+c
%	
%		>>syms a b c EPS;
%		>>ra=routh([1 a b c],EPS);
%		ra =
% 
%		[          1,          b]
%		[          a,          c]
%		[ (-c+b*a)/a,          0]
%		[          c,          0]
%
% 
%   Author:Rivera-Santos, Edmundo J.
%  
%

if(nargin<2),
	fprintf('\nError: Not enough input arguments given.');
	return
end

dim=size(poli);		%get size of poli		

coeff=dim(2);				%get number of coefficients
RA=sym(zeros(coeff,ceil(coeff/2)));	%initialize symbolic Routh array 

for i=1:coeff,
	RA(2-rem(i,2),ceil(i/2))=poli(i); %assemble 1st and 2nd rows
end

rows=coeff-2;		%number of rows that need determinants
index=zeros(rows,1);	%initialize columns-per-row index vector

for i=1:rows,
	index(rows-i+1)=ceil(i/2); %form index vector from bottom to top
end

for i=3:coeff,				%go from 3rd row to last
	if(all(RA(i-1,:)==0)),		%row of zeros
			fprintf('\nSpecial Case: Row of zeros detected.');
			a=coeff-i+2;		%order of auxiliary equation
			b=ceil(a/2)-rem(a,2)+1; %number of auxiliary coefficients
			temp1=RA(i-2,1:b);	%get auxiliary polynomial
			temp2=a:-2:0;		%auxiliry polynomial powers
			RA(i-1,1:b)=temp1.*temp2;	%derivative of auxiliary
	elseif(RA(i-1,1)==0),		%first element in row is zero
			fprintf('\nSpecial Case: First element is zero.');
			RA(i-1,1)=epsilon;	%replace by epsilon
	end
				%compute the Routh array elements
	for j=1:index(i-2),	
		RA(i,j)=-det([RA(i-2,1) RA(i-2,j+1);RA(i-1,1) RA(i-1,j+1)])/RA(i-1,1);
	end
end

	
	
```
