## Supplemental Data 1 for "Principles underlying implementation of *nearly*-homeostatic biological networks": shifted_identity.m

```
function [out,doutdx] = shifted_identity(x,x0,x0_discretes,band)
% When x0 outside of 5*band: shift_identify(x0)=x0, but is x0_discretes(1) at x=x0_discretes(1)+band 
% and x0_discrete(2) at x0_discretes(2) at x=x0_discretes(2)-band. 
% When x0 is inside of 5*band: shift_identity(x0) no longer has to be x0. 
% INPUT
% x - numerical scalar
% x0 - numerical scalar
% x0_discretes - 2 element vector; first element is assumed to be smaller than second element. 
% band - numerical scalar. 
idx_closestdisc=find(abs(x0-x0_discretes)==min(abs(x0-x0_discretes)),1);
x0_inbband=abs(x0-x0_discretes(idx_closestdisc))<5*band;
if ~x0_inbband
  if x>=x0
    doutdx=(x0_discretes(2)-x0)*(x0_discretes(2)-x0-band)^-1;
    out=doutdx*(x-x0)+x0;
  else
    doutdx=(x0-x0_discretes(1))*(x0-x0_discretes(1)-band)^-1;
    out=doutdx*(x-x0)+x0;
  end
else
  doutdx=(x0_discretes(2)-x0_discretes(1))*(x0_discretes(2)-x0_discretes(1)-2*band)^-1;
  out=doutdx*(x-x0_discretes(1)-band)+x0_discretes(1);
end
if x<x0_discretes(1)+band
  doutdx=0;
  out=x0_discretes(1);
elseif x>x0_discretes(2)-band
  doutdx=0;
  out=x0_discretes(2);
end
end
```
