## Supplemental Data 1 for "Principles underlying implementation of *nearly*-homeostatic biological networks": shifted_identity_Rnoband.m

```
function [out,doutdx] = shifted_identity_Rnoband(x,x0,shifted)
% If shifted is true, then call shifted_identity with 1e-2 band and appropriate discrete topo 
% parameter values based on x0. Otherwise, acts like the identity function.
if isa(x,'sym')
  shifted_identity_Rnoband=nan;
  syms shifted_identity_Rnoband(arg1,arg2,arg3)
  out=shifted_identity_Rnoband(x,x0,shifted);
  doutdx=nan;
else
  if shifted
    x0_discretes=[floor(x0),ceil(x0)];
    if x0_discretes(1)==x0_discretes(2) && (x0_discretes(1)==0 || x0_discretes(1)==1)
      x0_discretes(2)=x0_discretes(2)+1;
    elseif x0_discretes(1)==x0_discretes(2) && x0_discretes(1)==2
      x0_discretes(1)=x0_discretes(1)-1;
    end
    [out,doutdx]=shifted_identity(x,x0,x0_discretes,1e-2);
  else
    out=x;
    doutdx=1;
  end
end
end
```
