## Supplemental Data 1 for "Principles underlying implementation of *nearly*-homeostatic biological networks": smoothstep_shifted.m

```
function [out] = smoothstep_shifted(x,N,Ks,band)
% Compute output of the variably compressed smoothstep function 
% out = S_1( (softstep(N,Ks,x)-softstep(N,Ks,band(1)))/(softstep(N,Ks,band(2))-softstep(N,Ks,band(1)) )
% INPUTS
% x - input column array
% N - column array passed to softstep 
% Ks - column array passed to softstep; should be within band
% band - out=0 at band(:,1), out=1 at band(:,2)
if true
  out_softstep_x=softstep(N,Ks,x);
  out_softstep_b1=softstep(N,Ks,band(:,1));
  out_softstep_b2=softstep(N,Ks,band(:,2));
else
  % JIT compiler makes this option only negligibly faster. 
  out_softstep_x= 0.5 + 0.5 .* tanh(N .* x - N .* Ks);
  out_softstep_b1= 0.5 + 0.5 .* tanh(N .* band(:,1) - N .* Ks);
  out_softstep_b2= 0.5 + 0.5 .* tanh(N .* band(:,2) - N .* Ks);
end
y=(out_softstep_x-out_softstep_b1)./(out_softstep_b2-out_softstep_b1);
out=smoothstep_std(y,2); % N=2 ensures continuity of second derivative
end
```
