## Supplemental Data 1 for "Principles underlying implementation of *nearly*-homeostatic biological networks": smoothstep_shifted_d1.m

```
function [out] = smoothstep_shifted_d1(x,N,Ks,band)
% Compute derivative of smoothstep_shifted
% INPUTS
% x - input column array
% N - column array passed to softstep 
% Ks - column array passed to softstep; should be within band
% band - out=0 at band(:,1), out=1 at band(:,2)

out_softstep_x=softstep(N,Ks,x);
out_softstep_b1=softstep(N,Ks,band(:,1));
out_softstep_b2=softstep(N,Ks,band(:,2));
dy_band=out_softstep_b2-out_softstep_b1;
y=( out_softstep_x - out_softstep_b1 ) ./ dy_band;
out=smoothstep_std_d1(y,2) .* softstep_d1(N,Ks,x)./dy_band;
end
```
