## Supplemental Data 1 for "Principles underlying implementation of *nearly*-homeostatic biological networks": smoothstep_shifted_d1_Rnoband.m

```
function [out] = smoothstep_shifted_d1_Rnoband(Ns,Ks,s)
% Compute output of the derivative of variably compressed smoothstep function. 
% out = smoothstep_shifted_d1(s,Ns,Ks,[|-8|0,1.1]|[1.1,1.5]|[1,2|10]) depending on value of Ks 
% (<=1, <=1.3, or otherwise).
% All inputs are scalars. 
if Ks<=1
  out=smoothstep_shifted_d1(s,Ns,Ks,[-8,1.1]);
elseif Ks<=1.3
  out=smoothstep_shifted_d1(s,Ns,Ks,[1.1,1.5]);
else
  out=smoothstep_shifted_d1(s,Ns,Ks,[1,10]); 
end
end
```
