## Supplemental Data 1 for "Principles underlying implementation of *nearly*-homeostatic biological networks": smoothstep_shifted_Rnoband.m

```
function [out] = smoothstep_shifted_Rnoband(Ns,Ks,s)
% Compute output of the variably compressed smoothstep function 
% out = smoothstep_shifted(s,Ns,Ks,[-8|0,1.1]|[1.1,1.5]|[1,2|10]) depending on value of Ks
% (<=1, <=1.3, or otherwise).
% All inputs are scalars. 
if isa(s,'sym')
  smoothstep_shifted_Rnoband=nan;
  syms smoothstep_shifted_Rnoband(x,y,z)
  out=smoothstep_shifted_Rnoband(Ns,Ks,s);
elseif Ks<=1
  out=smoothstep_shifted(s,Ns,Ks,[-8,1.1]);
elseif Ks<=1.3
  out=smoothstep_shifted(s,Ns,Ks,[1.1,1.5]);
else
  out=smoothstep_shifted(s,Ns,Ks,[1,10]); 
end
end
```
