## Supplemental Data 1 for "Principles underlying implementation of *nearly*-homeostatic biological networks": smoothstep_std.m

```
function [out] = smoothstep_std(x,N)
% Compute output of the standard generalized smoothstep function of degree N
% out = S_N(x)
% INPUTS
% x - input array 
% N - order of generalized smoothstep. Supported values of N are [1].
if N==1
  out=-2.*x.^3 + 3.*x.^2;
elseif N==2
  out=6.*x.^5 - 15.*x.^4 + 10.*x.^3;
end
xl0=x<0;
xg1=x>1;
xinb=~xl0 & ~xg1;
out=xinb.*out+xg1;
end
```
