## Supplemental Data 1 for "Principles underlying implementation of *nearly*-homeostatic biological networks": smoothstep_std_d1.m

```
function [out] = smoothstep_std_d1(x,N)
% Compute output of the standard generalized smoothstep function of degree N
% out = S_N(x)
% INPUTS
% x - input array 
% N - order of generalized smoothstep
if N==1
  out=-6.*x.^2 + 6.*x;
elseif N==2
  out=30.*x.^4 - 60.*x.^3 + 30.*x.^2;
end
xl0=x<0;
xg1=x>1;
xinb=~xl0 & ~xg1;
out=xinb.*out;
end
```
