## Supplemental Data 1 for "Principles underlying implementation of *nearly*-homeostatic biological networks": softstep.m

```
function out = softstep(ns,Ks,s)
% Computes the tanh based softstep function. 
% softstep = 0.5 + 0.5 .* tanh(ns .* s - ns .* K)
% INPUTS
% ns, Ks, s - numeric array or symbolic scalar
out = 0.5 + 0.5 .* tanh(ns .* s - ns .* Ks);
end
```
