## Supplemental Data 1 for "Principles underlying implementation of *nearly*-homeostatic biological networks": softstep_d1.m

```
function out = softstep_d1(ns,Ks,s)
% Computes the derivative of tanh based softstep function. 
% INPUTS
% ns, Ks, s - numeric array or symbolic scalar
out = 0.5 .* (1-tanh(ns .* s-ns .* Ks).^2) .* ns;
end
```
