## Supplemental Data 1 for "Principles underlying implementation of *nearly*-homeostatic biological networks": softstep_hill.m

```
function out = softstep_hill(ns,Ks,s)
% Computes the hill function based softstep function. 
%   softstep = 1-Ks.^ns./(Ks.^ns+s.^ns). 
% This general form's symbolic deritive only has one additive term. 

% INPUTS
% ns, Ks, s - numeric array or symbolic scalar
out = 1-Ks.^ns./(Ks.^ns+s.^ns);
end
```
