## Supplemental Data 1 for "Principles underlying implementation of *nearly*-homeostatic biological networks": softstep_hill_inv.m

```
function out = softstep_hill_inv(ns,Ks,smap)
% Computes the hill function based softstep function. 
%   softstep = (-(Ks.^ns.*smap)./(smap - 1)).^(1./ns). 
% This general form's symbolic deritive only has one additive term. 

% INPUTS
% ns, Ks, smap - numeric array or symbolic scalar
out = (-(Ks.^ns.*smap)./(smap - 1)).^(1./ns);
end
```
