## Supplemental Data 1 for "Principles underlying implementation of *nearly*-homeostatic biological networks": sub_4_param.m

```
function sub_first =sub_4_param(first,second,dependencies)
% Substitute variables in the first by the second map container. This function
% skips all symmatrix entries, as second.keys contains non-symmatrix
% expressions to be substituted.
%
% Main usages: BioSystem.fullexp, replace in BioSystem
% 
% Input:
% first: map container of symbolic expression or characters OR
%   symbolic array OR cell array of strings
% second - map container; ('<name>',value). Value to be substituted is
%   compatible with function "subs".
% dependencies - OPTIONAL map container; keys are first.keys, each value is a cell array of
%   characters specifying the underlying variables that are used define the key. 
% Output:
% sub_first: map container that is substituted OR symbolic matrix

% convert ea value of dependencies to a cell array scalar symbolic expressions
if isa(second,'containers.Map')
  second_ks_vec=str2sym(second.keys);
  if nargin<=2 && isa(first,'containers.Map')
    dependencies=containers.Map(first.keys,repmat({num2cell(second_ks_vec)},1,first.Count));
  end
end
dependencies=containers.Map(dependencies.keys,...
  cellfun(@(v)str2sym_ip(v),dependencies.values,'UniformOutput',false));

% substitutions: if first is a map container, must have 1 sub for each key, in case a value is
% symbolic matrix
first=ensure_form(first,'symbolic');
second=ensure_form(second,'symbolic');
if isa(first,'containers.Map')
  sub_first=containers.Map();
  second_ks=num2cell(second_ks_vec);
  second_vs=second.values;
  first_ks=first.keys;
  for j=first_ks
    if isa(first(j{:}),'sym')
      if nargin<=2 && ~strcmp(j{:},first_ks{1})
        second_ks_rel_idx=true(size(second_ks_vec)); % for code speedup
      else
        second_ks_rel_idx=ismember(second_ks_vec,dependencies(j{:}));
      end
      sub_first(j{:})=subs(first(j{:}),second_ks(second_ks_rel_idx),second_vs(second_ks_rel_idx));
    else
      sub_first(j{:})=first(j{:});
    end
  end
elseif isa(first,'sym') || isa(first,'cell')
  sub_first=subs(first,second.keys,second.values);
else
  error('input format is incorrect');
end
end

function out_cell=str2sym_ip(in_cell)
if isempty(in_cell) || isa(in_cell{1},'sym')
  out_cell=in_cell;
else
  out_cell=num2cell(str2sym(in_cell));
end
end
```
