## Supplemental Data 1 for "Principles underlying implementation of *nearly*-homeostatic biological networks": sym_to_loformula.m

```
function[loformula]=sym_to_loformula(symexp)
%Convert a single symbolic expression to LibreOffice formula
%inputs:
%symexp - a single symbolic expression; length must be 1
%outputs:
%loformula - character string that represent LibreOffice formula version of  symexp

%input checking
if length(symexp)>1
    error('sym_to_loformula: input only accepts a single symbolic expression!')
end

loformula=char(symexp);
%convert multiplicative operator to spaces
loformula=regexprep(loformula,'*',' ');
%adding space to variables
loformula=regexprep(loformula,'(\W)(\w+)(\W)','$1 $2 $3 ');
loformula=regexprep(loformula,'(\W)(\w+)','$1 $2 ');
%convert all slashes to fractions
loformula=regexprep(loformula,'/',' over ');
%convert to proper subscriptions
loformula=regexprep(loformula,'(\D)(\d+)_(\d+)(\D)','$1_{$2 $3}$4');
loformula=regexprep(loformula,'([a-zA-Z])(\d+) ','$1_$2 ');
%convert double equality sign to single equality
loformula=regexprep(loformula,'==','=');
%convert multiple single spaces to one single space
loformula=regexprep(loformula,'\s+',' ');
end
```
