## Supplemental Data 1 for "Principles underlying implementation of *nearly*-homeostatic biological networks": symvar_all.m

```
function [C] = symvar_all(S)
% same input and output as symvar, but S can now include symmatrix and
% symfunmatrix
str=regexprep(S,{'<strong>','</strong>','[ \f\t\v]+'},{'','',''});
C=symvar(str);
funcs=symvarfun({str}); % D([1],f) still allows symvar to return f as symbolic variable. 
C=setdiff(C,funcs.funcname);
end
```
