## Supplemental Data 1 for "Principles underlying implementation of *nearly*-homeostatic biological networks": symvarfun.m

```
function [func_info] = symvarfun(expr)
% Finds all functions and their arguments within supplied symbolic expression. This function will 
% also extract f from D([1],f). 
% INPUT
% expr - cell array character vectors
% OUTPUT
% func_info - table with columns {'funcname','GroupCount','expridx','matchidx','numargs'} and row
%   names func_info.funcname

%% Ensure no spacing or newline in expr
if any(cellfun(@(x)~isempty(x),regexp(expr,'[ \f\t\v]+'),'UniformOutput',true),'all')
  error('symvarfun:input_check','expr should NOT have spacing!')
end

%% find all forward and reverse bracket, compute number of unclosed brackets at each 
%  position within expr, find locations of all commas
idx_fbr=regexp(expr,'\(');
idx_rbr=regexp(expr,'\)');
idx_commas=regexp(expr,',');
count_ubr=cell(size(idx_fbr));
for i=1:numel(expr)
  count_ubr{i}=zeros(size(expr{i}));
  count_ubr{i}(idx_fbr{i})=1;
  count_ubr{i}(idx_rbr{i})=-1;
  count_ubr{i}=cumsum(count_ubr{i});
end

%% find all functions, their matching positions, and which expression was matched into table func_info
expr4_func_1='[^\w]*?((?:<strong>)*\w+(?:</strong>)*)(';
expr4_func_2='D\(\[\d+\],(\w+)\)\(';
[tok_func_1,tok_func_1_idx]=regexp(expr,expr4_func_1,'tokens','tokenExtents');
[tok_func_2,tok_func_2_idx]=regexp(expr,expr4_func_2,'tokens','tokenExtents');
tok_func=cell(size(tok_func_1));
tok_func_idx=cell(size(tok_func_1_idx));
for i=1:numel(expr)
  tok_func{i}=[tok_func_1{i},tok_func_2{i}];
  tok_func_idx{i}=[tok_func_1_idx{i},tok_func_2_idx{i}];
end
func_info=arrayfun(...
  @(i)table(ones(flip(size(tok_func{i})))*i,[tok_func{i}{:}].',tok_func_idx{i}.'),...
  1:numel(tok_func),'UniformOutput',false);
func_info=vertcat(func_info{:});
func_info.Properties.VariableNames={'expridx','funcname','matchidx'};
if height(func_info)>=1
  func_info=rowfun(@(expridx,matchidx)deal(expridx(1),matchidx(1,:)),func_info,...
    'InputVariables',{'expridx','matchidx'},...
    'GroupingVariables',{'funcname'},...
    'OutputVariableNames',{'expridx','matchidx'});
  func_info.Properties.RowNames=func_info.funcname;
else
  func_info.GroupCount=zeros(0);
  func_info.Properties.RowNames={};
end

%% add to the func_info table number of arguments for each funcname
func_info.numargs=nan(height(func_info),1);
for i=1:height(func_info)
  row=table2cell(func_info(i,{'funcname','expridx','matchidx'}));
  [funcname,expridx,matchidx]=deal(row{:});
  position_fbr=matchidx(end)+1;
  br_ulevel=count_ubr{expridx}(position_fbr);
  position_rbr=find(count_ubr{expridx}==br_ulevel-1);
  position_rbr=position_rbr(find(position_rbr>=position_fbr,1));
  if isempty(position_rbr)
    position_rbr=numel(expr{expridx});
  end
  position_commas=idx_commas{expridx};
  func_info.numargs(i)=1 + sum( position_commas>position_fbr & position_commas<position_rbr & ...
    count_ubr{expridx}(position_commas)==br_ulevel ); 
end
end
```
