## Supplemental Data 1 for "Principles underlying implementation of *nearly*-homeostatic biological networks": timetracker.m

```
% Timing device that can be used to keep track of sequence of tasks. 
%% Specification
% * able to start, pause, and reset timer. 
% * allow checking of time elapsed
% * allow resetting of timer
% * allow tracking of iterations
%% Design
% * Store whether the timer is paused or not. 
% * Store amount of time elapsed after latest pause: b4pause
% * When timer is *start*ed and when *paused* is true, store starttime. Otherwise, 
%   do nothing.
% * *elapsed* checks total elapsed time; paused status play a role.
% * When timer is *pause*d, compute and store total elapsed time. 
% * Timer can be *reset*. 
% * Allow increased iteration by 1 as well as resetting of iterations.  


classdef timetracker < matlab.mixin.Copyable
  
  properties
    paused=true
  end
  properties (Hidden, Access=protected)
    b4pause=0
    starttime
    iter_count=0
  end
  
  methods
    function start (obj)
      if obj.paused
        obj.paused=false;
        obj.starttime=tic;
      end
    end
    function t=elapsed(obj)
      if obj.paused
        t=obj.b4pause;
      elseif ~obj.paused && ~isempty(obj.starttime)
        t=obj.b4pause+toc(obj.starttime);
      else
        error('timetracker:elapse','timetracker has never started!');
      end
    end
    function pause (obj)
      if ~obj.paused
        obj.b4pause=obj.elapsed;
        obj.paused=true;
      end
    end
    function reset(obj)
      obj.starttime=[];
      obj.paused=true;
      obj.b4pause=0;
    end
    function value=inc_iter(obj)
      obj.iter_count=obj.iter_count+1;
      value=obj.iter_count;
    end
    function reset_iter(obj)
      obj.iter_count=0;
    end
    function value=curr_iter(obj)
      value=obj.iter_count;
    end
  end
  methods (Static)
    function runtests
      tm=timetracker;
      tm.start;
      tic;
      pause(1);
      tm.start;
      pause(1);
      tm.pause; % timer paused
      elapsed=toc;
      pause(1);
      if abs(tm.elapsed-elapsed)>1e-3
        error('timetracker:test','Failed basic testing!'); 
      end
      tm.start;
      pause(1);
      % timer not paused
      if abs(tm.elapsed-elapsed-1)>1e-2
        error('timetracker:test','Failed basic testing!'); 
      end
      tm.reset;
      tm.start;
      pause(0.5);
      % timer not paused
      if abs(tm.elapsed-0.5)>5e-3; error('timetracker:test','Failed basic testing!'); end;
      for i=1:100
        tm.inc_iter;
        if tm.curr_iter~=i; error('timetracker:test','Failed basic testing!'); end
      end
      if tm.curr_iter~=100; error('timetracker:test','Failed basic testing!'); end
      tm.reset_iter;
      if tm.curr_iter~=0; error('timetracker:test','Failed basic testing!'); end
    end
  end
end
```
