## Supplemental Data 1 for "Principles underlying implementation of *nearly*-homeostatic biological networks": topologicalSort.m

```
function order = topologicalSort(dependencies,start_keys)
% topologicalSort determines a topological order for dependencies from least dependent to 
% the most dependent, for a starting set of keys. A variable is resolved if its dependencies are 
% less dependent than that variable. 
% INPUTS
% dependencies - a container map of variable name to symbols which that variable depends upon. Each 
%   values is a column cell array of characters. 
% start_keys - a subset of dependencies; column cell array of characters
% OUTPUTS
% order - column cell array of characters

start_keys=start_keys(:);

%% computing the subset of dependencies that start_keys depends upon
depKeys_rel = start_keys(:);
idx=0;
while numel(depKeys_rel)>0
  idx=idx+1;
  v=depKeys_rel{idx};
  depNames=dependencies(v);
  defVars_rel_new=setdiff(intersect(depNames,dependencies.keys),depKeys_rel);
  if isempty(defVars_rel_new) && idx==numel(depKeys_rel)
    break;
  else
    depKeys_rel=[depKeys_rel;defVars_rel_new];
  end
end

%% perform topological ordering of depKeys_rel
unresolved = depKeys_rel;
order = {}; % from least dependent to the most dependent
changed = true;

while ~isempty(unresolved) && changed
  changed = false;
  for i = numel(unresolved):-1:1
    v = unresolved{i};
    reqs = dependencies(v);
    reqs_in_defs = reqs(ismember(reqs, depKeys_rel));

    if all(ismember(reqs_in_defs, order)) % also true when reqs_in_defs is empty
      order{end+1} = v; %#ok<AGROW>
      unresolved(i) = [];
      changed = true;
    end
  end
end
if ~isempty(unresolved)
  error('topologicalSort:cyclic_dependencies','Cyclic or unresolved dependencies detected.');
end
order=order(:);
end
```
