## Supplemental Data 1 for "Principles underlying implementation of *nearly*-homeostatic biological networks": update_J.m

```
function J = update_J (ref_J,dx,drate,mode)
% Jacobian estimation via different methods including Broyden's update
if strcmp(mode,'Broy')
  %%% Broyden
  J=norm(dx)^-2*(drate-ref_J*dx)*(dx.') + ref_J;
elseif strcmp(mode,'BFGS')
  %%% BFGS update: unable to meet int_tol
  J=(drate.'*dx)^-1*drate*(drate.') - ...
    (dx.'*ref_J*dx)^-1*ref_J*dx*(dx.')*(ref_J.') + ref_J;
elseif strcmp(mode,'fmincon')
  %%% fmincon: very slow
  D=fmincon(@(D)obj_f(D,ref_J),D0,[],[],[],[],[],[],...
    @(D)cons_f(D,ref_J,dx,drate),opts);
  J=D + ref_J;
else
  error('This scenario should not occur!');
end
end
```
