## Supplemental Data 1 for "Principles underlying implementation of *nearly*-homeostatic biological networks": vec_replace.m

```
function in = vec_replace(in,coord,rep)
% replace certain coordinates of in with rep.
% Inputs
% in - vector
% coord - coordinates of in to replace with rep; linear index only!
% rep
% Output
% in but with coordinates coord replaced by rep.
in(coord)=rep;
end
```
