## Supplemental Data 1 for "Principles underlying implementation of *nearly*-homeostatic biological networks": VisKer.m

```
% Generate numerical kernels of supplied set of functions, and then visualize them and their
% desired intersections in 3D space  
%% INPUTS
% * *fs* is cell array of functions that makes in a 3 element column vector and output a
% scalar. 
% * *thres* is a vector of thresholds for different functions to be considered almost at zero.
% Same dimension as *fs*
% * *colors* is a cell array of colors of kernels for different functions. Same dimension as
% *fs* 
% * *prange* is a 3 x 2 matrix of plot ranges. First column vector specifies lower range; 
% second column vector specifies upper range. 
% * *intsect* is a 2 x N matrix of indices to compute and plot kernel intersections for. Empty
% matrix for no intersection to be computed. 
% * *x_ss* is 3 x N matrix of points to plot; each column is a separate point to plot.
% OPTIONAL.  
% OUTPUTS
% * *h* is a cell array of MatLab figure handles

function [h] = VisKer (fs,thres,colors,prange,intsect,x_ss)
thres=abs(thres);
h=cell(1,numel(fs)+size(intsect,2)+1);
for j=1:numel(fs)
  if j==2; hold on; end;
  [x,y,z]=find_ker(fs{j},thres(j),prange);
  h{j}=scatter3(x,y,z,36,colors{j},'.');
end
if ~isempty(intsect)
  for j=1:size(intsect,2)
    sel=cell(1,2);
    [sel{:}]=ismember(...
      table(h{intsect(1,j)}.XData.',h{intsect(1,j)}.YData.',h{intsect(1,j)}.ZData.'),...
      table(h{intsect(2,j)}.XData.',h{intsect(2,j)}.YData.',h{intsect(2,j)}.ZData.'));
    for k=1:2
      sel_rep=find(~sel{k}); sel_rep=sel_rep(1);
      if k==1
        x=h{intsect(k,j)}.XData; y=h{intsect(k,j)}.YData; z=h{intsect(k,j)}.ZData;
        x=x(sel{k}); y=y(sel{k}); z=z(sel{k});
        h{numel(fs)+j}=scatter3(x,y,z,36,'k','X');
      else
        sel{k}=sel{k}(sel{k}~=0);
      end
      h{intsect(k,j)}.XData(sel{k})=h{intsect(k,j)}.XData(sel_rep);
      h{intsect(k,j)}.YData(sel{k})=h{intsect(k,j)}.YData(sel_rep);
      h{intsect(k,j)}.ZData(sel{k})=h{intsect(k,j)}.ZData(sel_rep);
    end
  end
end
if nargin==6;
  h{end}=scatter3(x_ss(1,:),x_ss(2,:),x_ss(3,:),72,'k','O','filled');
end
hold off;
ax=gca;
ax.XLim=prange(1,:);ax.YLim=prange(2,:); ax.ZLim=prange(3,:); 
end
%Test code
%VisKer({@(xy)xy(3),@(yz)yz(1),@(xz)xz(2)},repmat(2.5e-2,1,3),...
% {'r','g','b'},[-1,1;-1,1;-1,1],[1,2,3;2,3,1],[[0;0;0],[1;1;1]]);

function [x1,x2,x3,v]=find_ker(f,thres,prange)
% Find kernel of f numerical by finding all coordinates x,y,z below threshold
% INPUTS
% f - scalar valued function of column vector [x;y;z]
% OUTPUTS
% x, y, z, v - column vector of near-kernel; v is value of f evaluated at x,y,z. 

x_range=prange(1,:); y_range=prange(2,:); z_range=prange(3,:);
[x1,x2,x3]=meshgrid(linspace(x_range(1),x_range(2),50),...
  linspace(y_range(1),y_range(2),50),...
  linspace(z_range(1),z_range(2),50));
v=zeros(size(x1));
for j=1:numel(x1)
  v(j)=f([x1(j);x2(j);x3(j)]);
end
x1=x1(:); x2=x2(:); x3=x3(:); v=v(:);
sel=abs(v)<=thres;
x1=x1(sel); x2=x2(sel); x3=x3(sel); v=v(sel);
end
```
