## Supplemental Data 1 for "Principles underlying implementation of *nearly*-homeostatic biological networks": demo_T7LacI_invhomeo.m

```
%%% Inverse homeostasis plots for the zeroth order degradation motif
%% start with a clean workspace
clear;
clc;
close all;
rng(1,'twister');

%% customizable section
% - Select mode between 'invhomeo' or 'nullclines'. If invhomeo, then inverse homeostasis plot
% of the system displayed and perturbation range permitting near-homeostasis is validated.
% If nullclines, then nullclines are displayed for each change in parameter set. 
% - Select mode between 'change_u1' or 'change_u2'. If mode contains 'nullclines', 'change_u1' is
% automatically assumed. 
% - mode can further contain 'greaterleak', where integration leakiness is higher
% - mode can further contain 'largerK12', where K12 is larger
% - mode containing nullclines can further contain 'linearscale', where linear rather than log scale
%   is used. 'linearscale_uselog' plots the same set of parameters on the log scale. 
% mode='invhomeo-change_u1-greaterleak'; 
% mode='invhomeo-change_u1-greaterleak';
mode='nullclines-change_u1'; 
%
% p_custom is a series of parameter sets that represents progress to
% near-homeostasis.
p_custom{1}=cell2map(...
  {'Kaia',rand_param([0;10],1);
  'Kd1_2',rand_param([0;10],1);
  'Kd2_1',rand_param([0;10],1);
  'Kiaa',rand_param([0;10],1);
  'V1',rand_param([0;10],1);
  'V2',rand_param([0;10],1);
  'Vd1',rand_param([0;10],1);
  'Vd2',rand_param([0;10],1);
  'b1_2',rand_param([0;0.1],1); 
  'b2_1',rand_param([0;0.1],1);
  'dKd1',rand_param([0;10],1);
  'dKd2',rand_param([0;10],1)*1e-1;
  'x2g',rand_param([0;1],1)*0});
p_custom{2}=[containers.Map();p_custom{1}];
p_custom{end}('Kd2_1')=p_custom{end}('Kd2_1')*1e4;
p_custom{3}=[containers.Map();p_custom{2}];
p_custom{end}('V2')=p_custom{end}('V2')*16; 
if ~isempty(regexp(mode,'greaterleak','once'))
  p_custom{end}('V2')=p_custom{end}('V2')*2.5;
  p_custom{end}('Vd2')=p_custom{end}('Vd2')*2.5;
else
  p_custom{end}('V2')=p_custom{end}('V2')*25;
  p_custom{end}('Vd2')=p_custom{end}('Vd2')*25;
end
p_custom{4}=[containers.Map();p_custom{3}];
p_custom{end}('Vd2')=p_custom{end}('Vd2')*1.4;
p_custom{5}=[containers.Map();p_custom{4}];
if ~isempty(regexp(mode,'linearscale','once')) 
  p_custom{end}('Vd2')=p_custom{end}('Vd2')/5;
  p_custom{end}('V2')=p_custom{end}('V2')/1.5;
end
p_custom{6}=[containers.Map();p_custom{5}];
p_custom{7}=[containers.Map();p_custom{6}];
if ~isempty(regexp(mode,'largerK12','once'))
  p_custom{end}('Kd1_2')=p_custom{end}('Kd1_2')*10;
else
  p_custom{end}('Kd1_2')=p_custom{end}('Kd1_2')/10*0.5;
end
if ~isempty(regexp(mode,'change_u1','once'))
  p_custom{end}('V1')=p_custom{end}('V1')*10;
else
  p_custom{end}('V1')=p_custom{end}('V1')*80;
end
p_custom{end}('b1_2')=p_custom{end}('b1_2')*1e-2;
syms max_sn(x,y)
plot_lims_invh=cell2map({...
  'x1',str2sym('[0,18.4405]');
  'x2',str2sym('[0,30]');
  'u',str2sym('[-Kiaa/Kaia,Kiaa/Kaia*10]');
  'u2',str2sym('[0,1]')});
if ~isempty(regexp(mode,'largerK12','once'))
  plot_lims_invh('u2')=str2sym('[0.95,1]');
end
if ~isempty(regexp(mode,'change_u2','once'))
   plot_lims_invh('x2')=str2sym('[2,30]');
end
x1_range=5*[1/1.47,1];
x1_NCpltrange=[2e-3*x1_range(1),20*x1_range(1)];
x1_NCpltrange_inv=x1_range([2,2]);
if ~isempty(regexp(mode,'linearscale','once'))
  x2_NCpltrange=[1e-2,2e2];
else
  x2_NCpltrange=[1e-2,1e2];
end

%% generate system and components required for inverse homeostasis plots
A=gen_T7LacI();
A_vec=A.vecfun_form;
A.param=p_custom{end};
p=cell2mat(A.param.values.');
s2_x2_it_x1=solve(A_vec(2),sym('x2'));
s2_x2_it_x1_num=matlabFunction(s2_x2_it_x1,...
  'Vars',{sym(A.param.keys).',sym(A.statevars).',sym(A.inputvars).'});
sel_vec=s2_x2_it_x1_num(p,[x1_range(2);nan],nan(2,1)); 
sel_vec=(sel_vec>=0).';
s1_u2_it_x1x2=solve(A_vec(1),sym('u2'));
s1_u2_it_x1x2_num=matlabFunction(s1_u2_it_x1x2,...
  'Vars',{sym(A.param.keys).',sym(A.statevars).',sym(A.inputvars).'});
s1_u1_it_x1x2=solve(A_vec(1),sym('u'));
s1_u1_it_x1x2_num=matlabFunction(s1_u1_it_x1x2,....
  'Vars',{sym(A.param.keys).',sym(A.statevars).',sym(A.inputvars).'});
x1_it_u_implicit=subs(A_vec(1),sym('x2'),s2_x2_it_x1(sel_vec)); % implicit u to x1ss map
Aopen=A.copy;
Aopen.sys('x2')=sym(0);
A.locked=true;
Aopen.locked=true;
prod_rates=cell2map({'x1',str2sym('H1_1*H1_2*V1*(1-u2)');
  'x2',str2sym('H2_1*H2_2*V2')});
prod_rates_open=cell2map({'x1',str2sym('H1_1*H1_2*V1*(1-u2)');
  'x2',str2sym('0')});
deg_rates=cell2map({'x1',str2sym('D1*Vd1');
  'x2',str2sym('D2*Vd2+Vd1/dKd1*x2')});
deg_rates_open=cell2map({'x1',str2sym('D1*Vd1');
  'x2',str2sym('0')});
multipliers=cell2map({'x1',str2sym('1');'x2',str2sym('1')});

%% computing relevant upper and lower ranges for the plots based on supplied x1_range and 
%  last set of parameters
x2_range=[sel_vec*s2_x2_it_x1_num(p,[x1_range(1);nan],nan(2,1)),...
  sel_vec*s2_x2_it_x1_num(p,[x1_range(2);nan],nan(2,1))];
x2_NCpltrange_inv=x2_range([2,2]);
  
if ~isempty(regexp(mode,'nullclines','once'))
  %% plot nullclines and inverse homeostasis plots for consecutive parameter changes
  Kiaa=A.param('Kiaa');
  Kaia=A.param('Kaia');
  if ~isempty(regexp(mode,'change_u1','once'))
    u=[[0;0],[9*Kiaa/Kaia;0]]; % each column is a separate perturbation level
  else
    u=[[0;0.85],[0;0]];
  end
  iter_firstout=6; % point at which to optimize controller rate parameters vs output rate p
  for i=1:numel(p_custom)
  % for i=(iter_firstout-1):numel(p_custom)
  % for i=[iter_firstout-1,numel(p_custom)]
  % for i=5
    % creating array of figure handles
    tmp=['IH_',num2str(i)];
    figIH{i}=findall(groot,'Type','figure','Name',tmp);
    if isempty(figIH{i})
      figIH{i}=figure;
    end
    clf(figIH{i});
    set(figIH{i},...
      'Name',tmp,...
      'defaultAxesTickLabelInterpreter','latex',...
      'defaultTextInterpreter','latex',...
      'defaultAxesTickLength',[0.04,0.04]);
    figIH{i}.Position=[0 0 446.0000 136.4000];
    tmp=['NC_',num2str(i)];
    figNC{i}=findall(groot,'Type','figure','Name',tmp);
    if isempty(figNC{i})
      figNC{i}=figure;
    end
    clf(figNC{i});
    set(figNC{i},...
      'Name',tmp,...
      'defaultAxesTickLabelInterpreter','latex',...
      'defaultTextInterpreter','latex',...
      'DefaultAxesFontSize',14,...
      'defaultAxesTickLength',[0.04,0.04]);
    figNC{i}.Position=[0 0 500 480];
    if i<iter_firstout
      % focusing on optimizing controller parameters
      compute_sscurves(A,figNC{i},figIH{i},{'x2'},'x2',...
        'x1',x1_NCpltrange,x2_NCpltrange,[1e3,2e3],...
        [cell2mat(p_custom{max(i-1,1)}.values.'),cell2mat(p_custom{i}.values.')],...
        repmat([nan;1],1,2),repmat(u(:,1),2),1,false,...
        prod_rates,deg_rates,multipliers,x1_NCpltrange_inv);
      % changing figNC appearance
      [tmp_XTick,IA]=unique([figNC{i}.Children.XTick,x1_range],'last');
      tmp_XTickL=extract_farray( ...
        [ cellfun(@(x)num2str(x),num2cell(figNC{i}.Children.XTick),'UniformOutput',false),...
        {'$x_O^L$','$x_O^U$'} ] , ...
        IA );
      figNC{i}.Children.XTick=tmp_XTick;
      figNC{i}.Children.XTickLabel=tmp_XTickL;
      figNC{i}.Children.Color='none';
      % changing figIH appearance
      tmp_XTick=[0,p_custom{i}('Kd2_1'),x1_range(2)];
      tmp_XTickL={'0','$K_{CO}$','$x_O^U$'};
      [~,tmp_XTickI]=sort(tmp_XTick);
      figIH{i}.Children(end-2).XTick=tmp_XTick(tmp_XTickI);
      figIH{i}.Children(end-2).XTickLabel=tmp_XTickL(tmp_XTickI);
      figIH{i}.Children(end-2).XTickLabelRotation=90;
      delete(figIH{i}.Children(end-2).Children(end-[1,4]));
      figIH{i}.Children(end-2).YTick=[0,0.5*p_custom{i}('V2')];
      figIH{i}.Children(end-2).YTickLabel={'0','$\phantom{pppppp} 0.5 V_C$'};
      figIH{i}.Children(end-1).YTick=[0,p_custom{i}('Vd2')];
      figIH{i}.Children(end-1).YTickLabel={'0','$\hat{V}_C$'};
      figIH{i}.Children(end).XTick=[0,10*p_custom{i}('dKd2')];
      figIH{i}.Children(end).XTickLabel={'0','$\phantom{ppp}10 \hat{K}_C$'};
      figIH{i}.Children(end).YAxis.Label.String{1}='rate of $x_C$';
      figIH{i}.Children(1).YAxis.Label.String{1}='rate of $x_C$';
      figIH{i}.Children(end).XAxis.Label.String='$x_C$';
      figIH{i}.Children(2).XAxis.Label.String='$x_O$';
    elseif i>=iter_firstout
      % focusing on optimizing output parameters
      compute_sscurves(A,figNC{i},figIH{i},{'x1'},'x1',...
        'x2',x2_NCpltrange,x1_NCpltrange,[1e3,2e3],...
        [repmat(cell2mat(p_custom{max(i-1,iter_firstout)}.values.'),1,2),...
        repmat(cell2mat(p_custom{i}.values.'),1,2)],...
        repmat([1;nan],1,4),repmat(u,1,2),2,true,...
        prod_rates,deg_rates,multipliers,x2_NCpltrange_inv);
      % changing figIH appearance
      delete(figIH{i}.Children(end-2).Children(end-[1,4]));
      figIH{i}.Children(end-2).YTick=...
        [0,p_custom{i}('V1')*(1-u(4))*[p_custom{i}('b1_2'),1]];
      figIH{i}.Children(end-2).YTickLabel=...
        {'0','$V_O (1-u_2^U) b_{OC}$','$V_O (1-u_2^U)$'};
      figIH{i}.Children(end-2).XTick=20*p_custom{i}('Kd1_2');
      figIH{i}.Children(end-2).XTickLabel={'$20 K_{OC}$'};
      figIH{i}.Children(end).YAxis.Label.String{1}='rate of $x_O$';
      figIH{i}.Children(1).YAxis.Label.String{1}='rate of $x_O$';
      figIH{i}.Children(end).XAxis.Label.String='$x_O$'; % not the same as i<iter_firstout
      figIH{i}.Children(2).XAxis.Label.String='$x_C$';
      % altering figNC appearance
      [tmp_XTick,IA]=unique([figNC{i}.Children.XTick,x1_range],'last');
      tmp_XTickL=extract_farray( ...
        [ cellfun(@(x)num2str(x),num2cell(figNC{i}.Children.XTick),'UniformOutput',false),...
        {'$x_O^L$','$x_O^U$'} ] , ...
        IA );
      figNC{i}.Children.XTick=tmp_XTick;
      figNC{i}.Children.XTickLabel=tmp_XTickL;
      for j=1:numel(figNC{i}.Children.Children)
        if isa(figNC{i}.Children.Children(j),'matlab.graphics.chart.primitive.Line')
          figNC{i}.Children.Children(j).LineStyle='--';
        end
      end
      figNC{i}.Children.Color='none';
      % add bounded box and merge figure for optimized controller nullcline into current figNC
      patch(figNC{i}.Children,'XData',[x1_range,flip(x1_range)],...
        'YData',[x2_range([1,1]),x2_range([2,2])],'FaceColor','none');
      copyobj(figNC{iter_firstout-1}.Children,figNC{i});
    end

    % changing axis labels for nullcline and inverse homeostasis plots
    figNC{i}.Children(end).XAxis.Label.String='$x_O$';
    figNC{i}.Children(end).YAxis.Label.String='$x_C$';

    % change figNC{i} to linear scale if necessary
    if ~isempty(regexp(mode,'linearscale','once')) && ~~isempty(regexp(mode,'linearscale_uselog','once'))
      for j=1:numel(figNC{i}.Children)
        figNC{i}.Children(j).XScale='linear';
        figNC{i}.Children(j).XLim(2)=1.5*x1_range(2);
        figNC{i}.Children(j).YScale='linear';
      end
    end
    
    % aligning figIH axes
    drawnow;
    for j=1:(numel(figIH{i}.Children)/2)
      figIH{i}.Children(2*j).Position(4)=figIH{i}.Children(2*j).Position(4)*0.9;
      figIH{i}.Children(2*j-1).Position=figIH{i}.Children(2*j).Position;
    end
  end

  %% compare performance of closed and open loop system by simulation and plot the outputs
  % simulation
  resp=find_dose_response(A,u',[1e4,1e4+1e3],5);
  Aopen.x0_map('x2')=resp(1,2);
  Aopen.param=containers.Map(A.param.keys,A.param.values);
  resp_open=find_dose_response(Aopen,u',[1e4,1e4+1e3],5);
  % plot change in steady state for both open and closed loop system onto nullcline plot
  set(0,'currentfigure',figNC{end});
  quiver(resp(1,1),resp(1,2),...
    resp(2,1)-resp(1,1),resp(2,2)-resp(1,2),...
    'off','Color','blue','LineWidth',2,'MaxHeadSize',0.4);
  quiver(resp_open(1,1),resp_open(1,2),...
    resp_open(2,1)-resp_open(1,1),resp_open(2,2)-resp_open(1,2),...
    'off','Color','cyan','LineWidth',2,'MaxHeadSize',0.4);
  % plot closed and open loop system steady states in output vs u graph
  tmp='Bar';
  figB=findall(groot,'Type','figure','Name',tmp);
  if isempty(figB)
    figB=figure;
  end
  clf(figB);
  set(figB,'Name',tmp,'defaultAxesTickLabelInterprete','latex',...
    'DefaultAxesFontSize',14,'defaultAxesTickLength',[0.04,0.04]);
  figB.Position(3:4)=[196 190];
  set(0,'currentfigure',figB);
  colororder({'blue','cyan'});
  bar(categorical(repmat({'uL';'uU'},1,2)),[resp(:,1),resp_open(:,1)]);
  legend({'feedback','open loop'},'location','northoutside');
  ax=gca;
  ax.XLabel.String='u_2';
  ax.YLabel.String='x_O';
  ax.YTick=[0,x1_range];
  ax.YTickLabel={'0','$x_O^L$','$x_O^U\phantom{x_O}$'};
  if ~isempty(regexp(mode,'change_u1','once'))
    ax.XTickLabel={'$u_1^L$','$u_1^U$'};
  else
    ax.XTickLabel={'$u_2^U$','$u_2^L$'};
  end
elseif ~isempty(regexp(mode,'invhomeo','once'))
  %% computing and validating steady states at lower and upper limits of perturbation for the 
  % last parameter set
  A.param=p_custom{end};
  if ~isempty(regexp(mode,'change_u2','once'))
    u_range=[0,0];
    u_range_open=u_range;
    % smaller x1ss corresponds to larger u2
    u2_range=[s1_u2_it_x1x2_num(p,[x1_range(1);x2_range(1)],[u_range(1);nan]),...
      s1_u2_it_x1x2_num(p,[x1_range(2);x2_range(2)],[u_range(2);nan])];
    u2_range_open=[s1_u2_it_x1x2_num(p,[x1_range(1);x2_range(1)],[u_range(1);nan]),...
      s1_u2_it_x1x2_num(p,[x1_range(2);x2_range(1)],[u_range(2);nan])];
  else
    u2_range=[0,0];
    u2_range_open=u2_range;
    u_range=[s1_u1_it_x1x2_num(p,[x1_range(1);x2_range(1)],[nan;u2_range(1)]),...
      s1_u1_it_x1x2_num(p,[x1_range(2);x2_range(2)],[nan;u2_range(2)])];
    u_range_open=[s1_u1_it_x1x2_num(p,[x1_range(1);x2_range(1)],[nan;u2_range(1)]),...
      s1_u1_it_x1x2_num(p,[x1_range(2);x2_range(1)],[nan;u2_range(2)])];
  end
  resp=find_dose_response(A,[u_range.',u2_range.'],[1e4,1e4+1e3],5);
  Aopen.x0_map('x2')=resp(1,2);
  Aopen.param=containers.Map(A.param.keys,A.param.values);
  resp_open=find_dose_response(Aopen,[u_range_open.',u2_range_open.'],[1e4,1e4+1e3],5);
  assert(all(abs(sort(resp(:,1))-x1_range.')<1e-10));
  assert(all(abs(sort(resp_open(:,1))-x1_range.')<1e-10));

  %% plot inverse homeostasis plots for both the closed and open loop system
  i=1;
  fig=findall(groot,'Type','figure','Name',sprintf('invhomeoplots-%d',i));
  if isempty(fig)
    fig=figure('Name',sprintf('invhomeoplots-%d',i),'defaultAxesTickLabelInterprete','latex',...
      'defaultAxesTickLength',[0.02,0.02]);
    fig.Position=[0 0 733 225];
  end
  set(0,"CurrentFigure",fig);
  if i==1 % compute_invhomeo for the closed loop system. 
    compute_invhomeo(A,[x1_range;x2_range],[u_range;u2_range],...
      [false,true,true,true; true, true,false(1,2)],...
      [true,true, true,true; false,true,false(1,2)],...
      prod_rates,deg_rates,multipliers,plot_lims_invh,false);
  else
    compute_invhomeo(Aopen,[x1_range;resp_open(:,2).'],[u_range_open;u2_range_open],...
      [false,true,true,true; true, true,false(1,2)],...
      [true,true, true,true; false,true,false(1,2)],...
      prod_rates_open,deg_rates_open,multipliers,plot_lims_invh,false);
  end
  % adding necessary tick labels
  fig.Children(end-2).XTick=30*p_custom{end}('Kd1_2');
  fig.Children(end-2).XTickLabel={'$30 K_{OC}$'};
  fig.Children(end-4).XTick=5*p_custom{end}('Kiaa')/p_custom{end}('Kaia');
  fig.Children(end-4).XTickLabel={'$5 \frac{K_{iaa}}{K_{aia}}$'};
  fig.Children(end-8).XTick=[0,p_custom{end}('Kd2_1')];
  fig.Children(end-8).XTickLabel={'0','$K_{CO}$'};
  fig.Children(end-8).YTick=[0,0.5*p_custom{end}('Vd2'),0.5*p_custom{end}('V2')];
  fig.Children(end-8).YTickLabel={'0','$0.5 \hat{V}_C$','$0.5 V_C$'};
  fig.Children(end-10).XTick=[0,40*p_custom{end}('dKd2')];
  fig.Children(end-10).XTickLabel={'0','$40 \hat{K}_C$'};
  fig.Children(1).YLabel.String{1}='rate of $x_C$';
  fig.Children(8).YLabel.String{1}='rate of $x_C$';
  fig.Children(9).YLabel.String{1}='rate of $x_O$';
  fig.Children(16).YLabel.String{1}='rate of $x_O$';
  fig.Children(2).XLabel.String='$u_2$';
  fig.Children(4).XLabel.String='$u_1$';
  fig.Children(6).XLabel.String='$x_C$';
  fig.Children(8).XLabel.String='$x_O$';
  % ensure overlapping axis have the same position
  drawnow;
  for i=1:(numel(fig.Children)/2)
    fig.Children(2*i-1).Position=fig.Children(2*i).Position;
  end
end
```
