## Supplemental Data 1 for "Principles underlying implementation of *nearly*-homeostatic biological networks": demo_T7LacI_leakinessreduction.m

```
close all; clear;clc;
rng(1,'twister');

%% Demonstration for publication: near integral control is not sufficient
%  for near homeostasis
%
% Code simplification on 2023-12-01, 2024-06-21
optimal_nonctrlleak=true; % true or false;
% mode includes one of {'h_only',''} AND one of {'change_u1','chane_u2'}
% AND one of {'' (non-controller parameter optimization) , 'optim_include_V2K21b21'}
mode='change_u1'; % runs through more code 
% mode='h_only-change_u1'; % runs through more code

%% construct T7LacI system, open loop system and the perfect controller
A=gen_T7LacI();
A.num_sim_only=true;
A.ss_iter_limit=1;
A.prange('rndtracker')=[0,1];
A.param('rndtracker')=0.5;
AL=A.copy;
AL.flux('fD2')=1;
AL.sys('x2')=subs(AL.sys('x2'),sym('Vd1'),0);
Aopen=A.copy;
convert_to_process(Aopen,{'x2'});
Aopen.prange('x2')=[0,inf];
Aopen.param('x2')=0;
A.locked=true;
AL.locked=true;
pars_fixed={'Kiaa','Kaia'};
pars_nopt={'rndtracker','Vd1','dKd1'}; % prevent optim_nonctrlleak from changing leakiness
pars_opt=setdiff(A.param.keys,[pars_fixed,pars_nopt]);
if ~isempty(regexp(mode,'change_u1','once')) % ensure perturbation always increase
  u=[0,0;9*A.param('Kiaa')/A.param('Kaia'),0]; 
else
  u=[0,0;0,0.55];
end

%% inverse homeostasis metrics in numeric form
% s1_x1_it_u takes forever to compute. 
A_vec=A.vecfun_form;
s2_x1_it_x2=solve(subs(A_vec(2),str2sym('Vd1/dKd1'),sym('k_dil')),sym('x1')); 
s2_x2_it_x1=solve(A_vec(2),sym('x2'));
s2_x2_it_x1_num=matlabFunction(s2_x2_it_x1,'Vars',...
  {sym(A.param.keys).',sym(A.statevars).',sym(A.inputvars).'});
s1_x2_it_x1u=solve(A_vec(1),sym('x2'));
s1_x2_it_x1u_num=matlabFunction(s1_x2_it_x1u,'Vars',...
  {sym(A.param.keys).',sym(A.statevars).',sym(A.inputvars).'});
s1_u2_it_x1x2u=solve(A_vec(1),sym('u2'));
s1_u2_it_x1x2u_num=matlabFunction(s1_u2_it_x1x2u,'Vars',...
  {sym(A.param.keys).',sym(A.statevars).',sym(A.inputvars).'});
s1_u1_it_x1x2u=solve(A_vec(1),sym('u'));
s1_u1_it_x1x2u_num=matlabFunction(s1_u1_it_x1x2u,'Vars',...
  {sym(A.param.keys).',sym(A.statevars).',sym(A.inputvars).'});
s1_x1_it_x2u=solve(A_vec(1),sym('x1'));
s1_x1_it_x2u_num=matlabFunction(s1_x1_it_x2u,'Vars',...
  {sym(A.param.keys).',sym(A.statevars).',sym(A.inputvars).'});
AL_vec=AL.vecfun_form;
x1_set=solve(AL_vec(2),sym('x1'));
x1_set_num=matlabFunction(x1_set,'Vars',{sym(A.param.keys.')});

%% computing arguments for fmincon
% fmincon objective (sum of positive distance from the desired u range for closed loop system),
% constraint (lower range permitting near-homeostasis matches that start for closed loop AND output 
% steady state of open loop system after perturbation is sufficiently high AND lower steady state 
% range is not too small) and options
pars_opt_ctrlleak={'Vd2','dKd2'};
if ~isempty(regexp(mode,'optim_include_V2K21b21','once'))
else
  pars_opt_ctrlleak=[pars_opt_ctrlleak,{'V2','Kd2_1','b2_1'}];
end
pars_opt_ctrlleak=sort(pars_opt_ctrlleak);
pars_opt_nctrlleak=setdiff(pars_opt,pars_opt_ctrlleak);
pars_fixednoptoptctrlleakoptnctrlleak=[pars_fixed,pars_nopt,pars_opt_ctrlleak,pars_opt_nctrlleak];
[~,p_idxinnoptoptctrlleakoptnctrlleak]=ismember(A.param.keys,pars_fixednoptoptctrlleakoptnctrlleak);
p2pA=@(p_new)extract_farray(p_new,p_idxinnoptoptctrlleakoptnctrlleak);
[fmincon_obj_u1,fmincon_cons_u1]=leakinessred_fmincon_ui_gen(p2pA,u,1,...
  true,x1_set_num,s1_u1_it_x1x2u_num,s1_x1_it_x2u_num,s2_x2_it_x1_num);
[fmincon_obj_u2,fmincon_cons_u2]=leakinessred_fmincon_ui_gen(p2pA,u([2,1],:),2,...
  false,x1_set_num,s1_u2_it_x1x2u_num,s1_x1_it_x2u_num,s2_x2_it_x1_num);
if ~isempty(regexp(mode,'change_u1','once'))
  fmincon_obj=fmincon_obj_u1;
  fmincon_cons=fmincon_cons_u1;
else
  fmincon_obj=fmincon_obj_u2;
  fmincon_cons=fmincon_cons_u2;
end
p_opt_nctrlleak_bnds=cell2mat(A.prange.values(pars_opt_nctrlleak).'); % 2 column matrix
p_opt_nctrlleak_bnds(ismember(pars_opt_nctrlleak,{'V2','Vd2'}),2)=1e4;
opts=optimoptions('fmincon');
opts.MaxFunctionEvaluations=1e4;
opts.MaxIterations=1e4;
opts.Display='iter-detailed';
opts.ConstraintTolerance=1e-8;
opts.StepTolerance=1e-16;

%% simulations: for each set of leakiness parameters, at each integration leakiness, 1) find 
%  a perturbation sensitive open loop system and if applicable, find optimal non-leakiness
%  parameters, 2) simulate corresponding perfect integral controller and see perfect adaptation is
%  attained under the right mode. 
int_leakiness=[1e-1,3.15e-2,1e-2,1e-3]; 
numsims_pars=33; 
A.locked=false;
AL.locked=false;
output_parfor={};
parfor c=1:3
% for c=1
  Ac=A.copy;
  ALc=AL.copy;
  Aopenc=Aopen.copy;
  Ac.locked=true;
  ALc.locked=true;
  Aopenc.locked=true;
  Aopenc_params_nox2=setdiff(Aopenc.param.keys,{'x2'});
  outputsvars_leak=arrayfun( @(l)strcat({'Aopen_x1fold_','o_AL_','rs_AL_','o_A_','rs_A_'},num2str(l)) ,...
    int_leakiness , 'UniformOutput' , 0 );
  outputsvars_leak=[outputsvars_leak{:}];
  outputs=table('Size',[numsims_pars,Ac.param.Count+1+numel(outputsvars_leak)],...
    'VariableTypes',repmat({'double'},1,Ac.param.Count+1+numel(outputsvars_leak)),...
    'VariableNames',[strcat(Ac.param.keys,'_lastleak'),{'x2_lastleak'},outputsvars_leak]);
  outputs{:,:}=nan;

  rndstream=RandStream('twister','Seed',c);
  rng(c,'twister'); % separate randstream and separate global stream for each parfor iteration
  i=1;
  while i<=numsims_pars % for each set of leakiness parameters
    % generating a set of leakiness parameters.
    get_feasible(Ac,@(p)true,pars_nopt);
    get_feasible(Ac,{'V2>Vd2';'V2*b2_1<Vd2'},pars_opt);
    dKd2=Ac.param('dKd2');
    V2=Ac.param('V2');
    Vd2=Ac.param('Vd2');
    Kdil=Ac.param('Vd1')/Ac.param('dKd1');
    l_cur=max([dKd2/5,Kdil/V2,Kdil/Vd2]);

    % at each integration leakiness; lowest leak first to speed up code when mode contains h_only
    l_goodleakprog=true; % whether progress through different leakiness is satisfactory
    for l=sort(int_leakiness,'ascend') 
      % set controller leakiness
      Ac.param('dKd2')=dKd2/l_cur*l;
      Ac.param('V2')=V2*l_cur/l;
      Ac.param('Vd2')=Vd2*l_cur/l;
      p0_fixed=cell2mat(Ac.param.values(pars_fixed)).';
      p0_nopt=cell2mat(Ac.param.values(pars_nopt)).';
      p0_opt_ctrlleak=cell2mat(Ac.param.values(pars_opt_ctrlleak)).';
      p_opt_nctrlleak=cell2mat(Ac.param.values(pars_opt_nctrlleak)).';

      % Searching and checking open loop system and simulate perfect integral controller. 
      % - When optimal_nonctrlleak==true, fmincon first finds a set of non-controller leakiness 
      %   parameters that satisfy the inequality constraints in fmincon_cons before the main
      %   optimization. The first round of optimization is necessary because the interior point 
      %   algorithm assumes that the starting point must be in the inequality constrainted feasible 
      %   region.
      % - When optimal_nonctrlleak==false, random set of non-leakiness parameters are generated. 
      for tries=1:10
        % whether open loop condition is being satisfied and perfect controller is homeostatic when
        % required. 
        l_goodtry=true; 

        % searching for satisfactory open loop and simulate closed loop as well
        if optimal_nonctrlleak
          mat_poptnctrlleak=nan(numel(p_opt_nctrlleak),20);
          mat_obj=nan(1,20);
          for m=1:4
            p0_opt_nctrlleak=rand_param(p_opt_nctrlleak_bnds.',1,rndstream).';
            p0_opt_nctrlleak(strcmp(pars_opt_nctrlleak,'V2'))=Ac.param('V2');
            [p_opt_nctrlleak,cons_obj,eflag_pre]=fmincon(...
              @(p_opt_nonctrll)...
              sum(max(fmincon_cons(p0_fixed,p0_nopt,p0_opt_ctrlleak,p_opt_nonctrll),[0,0])),...
              ...
              p0_opt_nctrlleak,[],[],[],[],p_opt_nctrlleak_bnds(:,1),p_opt_nctrlleak_bnds(:,2),[],opts);
            if (eflag_pre==1 || eflag_pre==2) && cons_obj<=0
              [p_opt_nctrlleak,uobj,eflag]=fmincon(...
                @(p_opt_nonctrll)fmincon_obj(p0_fixed,p0_nopt,p0_opt_ctrlleak,p_opt_nonctrll),...
                p_opt_nctrlleak,[],[],[],[],p_opt_nctrlleak_bnds(:,1),p_opt_nctrlleak_bnds(:,2),...
                @(p_opt_nonctrll)fmincon_cons(p0_fixed,p0_nopt,p0_opt_ctrlleak,p_opt_nonctrll),opts);
              if uobj<norm(u(2,:)-u(1,:)) && (eflag==1 || eflag==2)
                mat_poptnctrlleak(:,m)=p_opt_nctrlleak;
                mat_obj(1,m)=uobj;
                if uobj<1e-8
                  break;
                end
              end
            end
          end
          if ~isempty(find(mat_obj==min(mat_obj,[],"all","omitnan"),1))
            p_opt_nctrlleak=mat_poptnctrlleak(:,find(mat_obj==min(mat_obj,[],"all","omitnan"),1));
            Ac.param=[Ac.param;containers.Map2(pars_opt_nctrlleak,p_opt_nctrlleak)];
          else
            l_goodtry=l_goodtry && false;
          end
        else
          get_feasible(Ac,@(p)true,pars_opt_nctrlleak);
        end
        [cineq,ceq,x1l,x2l]=fmincon_cons(p0_fixed,p0_nopt,p0_opt_ctrlleak,p_opt_nctrlleak);
        Ac.x0=[x1l;x2l];
        [x_A,x_A_f]=find_dose_response(Ac,u,[1e15,2e15],1);
        if all(min(abs(x_A_f./x_A),x_A_f)<=1e-3)
          Ac.x0=x_A(1,:);
          outputs.x2_lastleak(i)=x_A(1,2);
          outputs{i,strcat('o_A_',num2str(l))}=x_A(1);
          %%% simulate open loop system to compute rs_A
          Aopenc.param('x2')=Ac.x0_map('x2');
          Aopenc.param=[Aopenc.param;
            containers.Map2(Aopenc_params_nox2,Ac.param.values(Aopenc_params_nox2))];
          [x_Aopen,x_Aopen_f]=find_dose_response(Aopenc,u,[1e15,2e15],1);
          assert(abs(x_Aopen(1,1)-x_A(1,1))<1e-4);
          outputs.rndtracker(i)=Ac.param('rndtracker');
          outputs.(strcat('rs_A_',num2str(l)))(i)=...
            log10(x_A(2,1)/x_A(1,1))/log10(x_Aopen(2,1)/x_Aopen(1,1));
          outputs.(strcat('Aopen_x1fold_',num2str(l)))(i)=10^abs(log10(x_Aopen(2,1)/x_Aopen(1,1)));
          if optimal_nonctrlleak && l_goodtry && i==1
            %%% validation: u(1,:) results in x1_set_num AND open loop is perturbation sensitive
            assert( outputs.(strcat('Aopen_x1fold_',num2str(l)))(i)>=2 );
            if ~isempty(regexp(mode,'change_u1','once'))
              assert( abs(0.9*x1_set_num(cell2mat(Ac.param.values).')-x_A(1,1))/x_A(1,1)<1e-3 );
            else
              assert( abs(0.9*x1_set_num(cell2mat(Ac.param.values).')-x_A(2,1))/x_A(2,1)<1e-3 );
            end
          end
          if outputs.(strcat('Aopen_x1fold_',num2str(l)))(i)>=2
            l_goodtry=l_goodtry && true;
          else
            l_goodtry=l_goodtry && false;
          end
        else
          l_goodtry=l_goodtry && false;
        end

        % simulate perfect integral controller
        if l_goodtry
          ALc.param=containers.Map2(ALc.param.keys,Ac.param.values(ALc.param.keys));
          ALc.x0=[ max(0,x1_set_num(p2pA([p0_fixed;p0_nopt;p0_opt_ctrlleak;p_opt_nctrlleak]))) ;
            ... % sometimes, a low leak controller might not result in perfect-homeostasis due to change
            ... % in controller toward negative values.
            max( 0 , s1_x2_it_x1u_num(p2pA([p0_fixed;p0_nopt;p0_opt_ctrlleak;p_opt_nctrlleak]),...
            x1_set_num(p2pA([p0_fixed;p0_nopt;p0_opt_ctrlleak;p_opt_nctrlleak])),u(1,:).') ) ];
          [x_AL,x_AL_f]=find_dose_response(ALc,u,[1e15,2e15],1);
          if all(min(abs(x_AL_f./x_AL),x_AL_f)<=1e-3)
            %%% verify that AL is also perfectly homeostatic
            x_AL_2=find_dose_response(ALc,u,[1e15,2e15],1);
            assert(all(abs(x_AL(:,1)-x_AL_2(:,1))<1e-10));
            %%% simulate open loop system to compute rs_AL
            outputs{i,['o_AL_',num2str(l)]}=x_AL(1);
            Aopenc.param('x2')=x_AL(1,2);
            Aopenc.param=[Aopenc.param;
              containers.Map2(Aopenc_params_nox2,Ac.param.values(Aopenc_params_nox2))];
            [x_Aopen,x_Aopen_f]=find_dose_response(Aopenc,u,[1e15,2e15],1);
            assert(abs(x_Aopen(1,1)-x_AL(1,1))<1e-8);
            outputs{i,['rs_AL_',num2str(l)]}=(x_AL(2,1)-x_AL(1,1))/(x_Aopen(2,1)-x_Aopen(1,1));
          else
            if ~isempty(regexp(mode,'h_only','once'))
              l_goodtry=l_goodtry && false;
            end
          end
        end

        % exit if try is good
        if l_goodtry
          break;
        end
      end

      % stop progressing through more leakiness if progress is not good
      l_goodleakprog=l_goodleakprog && l_goodtry;
      if ~l_goodleakprog
        break;
      end
    end
    outputs{i,strcat(Ac.param.keys,'_lastleak')}=cell2mat(Ac.param.values);

    % proceed to next i
    if l_goodleakprog
      fprintf('core %d, iteration %d is complete...\n',c,i);
      i=i+1;
    end
  end
  outputs_parfor{c}=outputs;
end
outputs_parfor=vertcat(outputs_parfor{:});

%% for each integration leakiness and each of near-integral controller and perfect integral controller,
%  label each simulation to a class of near-homeostasis
cols_toplot=[strcat('rs_A_',arrayfun(@num2str,int_leakiness,'UniformOutput',0)),...
  strcat('rs_AL_',arrayfun(@num2str,int_leakiness,'UniformOutput',0))];
cats_ord={'<=1/6','>1/6 & <=1/3','>1/3 & <=2/3','>2/3'};
cats_map=cell2map({cats_ord{1},[-1e-6,1/6];
  cats_ord{2},[1/6,1/3];
  cats_ord{3},[1/3,2/3]; ...
  cats_ord{4},[2/3,inf]});
for cols=cols_toplot
  col=cols{:};
  for k=cats_map.keys
    k=k{:};
    range=cats_map(k);
    l=abs(outputs_parfor.(col))>range(1) & abs(outputs_parfor.(col))<=range(2);
    outputs_parfor.(strcat(col,'_cat'))(l)=repmat({k},sum(l),1);
  end
  l=isnan(outputs_parfor.(col));
  outputs_parfor.(strcat(col,'_cat'))(l)=repmat({'no steady state'},sum(l),1);
end

%% for each category, compute the proportion of simulations for a given integration leakiness 
%  belonging to that category: cats_prop's each column corresponds to a
%  different integration leakiness
cats=categorical(outputs_parfor{:,strcat(cols_toplot,'_cat')},...
  [cats_ord,{'no steady state'}],'ordinal',true);
cats_count=countcats(cats,1);
cats_prop=cats_count./sum(cats_count,1);

%% compute individual parameter's leakiness 
outputs_parfor.('dKd2/5')=outputs_parfor.dKd2_lastleak/5;
outputs_parfor.('kdil/V2')=outputs_parfor.Vd1_lastleak./...
  outputs_parfor.dKd1_lastleak./outputs_parfor.V2_lastleak;
outputs_parfor.('kdil/Vd2')=outputs_parfor.Vd1_lastleak./...
  outputs_parfor.dKd1_lastleak./outputs_parfor.Vd2_lastleak;

%% plot proportion of steady state reaching parameter sets that belong to 
%  different degrees of near-homeostasis. 
fig=figure;
fig.Position([3,4])=[364,166];
h=bar(cats_prop.','stacked','FaceColor','flat');
set(h,{'DisplayName'},categories(cats));
for k = 1:size(cats_prop,1)
    h(k).CData = [0.5,0.5,1-(k-1)*1/6];
end
lgd=legend('location','eastoutside','FontSize',9);
lgd.Title.Interpreter="latex";
lgd.Title.String=['$\frac{\log(x_O^\textrm{cl,ss}(u^L)/x_O^\textrm{cl,ss}(u^U))}',...
  '{\log(x_O^\textrm{ol,ss}(u^L)/x_O^\textrm{ol,ss}(u^U))}$'];
ax=gca;
ax.FontSize=12;
cols_toplot_labels=[ arrayfun(@(a,b)num2str(a,b{:}),int_leakiness,...
  repmat({'%.3g'},size(int_leakiness)),'UniformOutput',0) , ...
  arrayfun(@(a,b)['$',num2str(a,b{:}),'\rightarrow 0$'],int_leakiness,...
  repmat({'%.3g'},size(int_leakiness)),'UniformOutput',0) ];
ax.XTickLabel=cols_toplot_labels;
ax.TickLabelInterpreter='latex';
ax.XTickLabelRotation=45;
ax.XLabel.String={'integration leakiness'};
ax.YLabel.String='proportion';
exportgraphics(fig,...
  ['demo-reduce_leakiness-optimnonctrlleak_',num2str(optimal_nonctrlleak),'-',mode,'.pdf']);

%% plot the distribution of individual parameter's leakiness at max leakiness
fig=figure;
fig.Position([3,4])=[225,225];
hold on;
for k={'dKd2/5','kdil/V2','kdil/Vd2'}
  kexp=k{:};
  [f,xi]=ksdensity(outputs_parfor.(kexp),...
    'support',[min(outputs_parfor.(kexp))-1e-4,max(outputs_parfor.(kexp))+1e-4]);
  plot(xi,f/max(f));
end
legend({'$\hat{K}_2/5$','$k_{\textrm{dil}}/V_2$','$k_{\textrm{dil}}/\hat{V}_2$'},...
  'Interpreter','latex');
ax=gca;
ax.FontSize=12;
ax.XLabel.String='value';
ax.YLabel.String='normalized density';
exportgraphics(fig,...
  ['demo-reduce_leakiness-optimnonctrlleak_',num2str(optimal_nonctrlleak),...
  '-',mode,'-ind_leakiness.pdf']);
save(['demo-reduce_leakiness-optimnonctrlleak_',num2str(optimal_nonctrlleak),'-',mode,'.mat'],...
  'outputs_parfor');
```
