## Supplemental Data 1 for "Principles underlying implementation of *nearly*-homeostatic biological networks": gen_T7LacI.m

```
function A=gen_T7LacI(intmf)
% construct symbolic version of the T7 LacI system
% Inputs:
% intm - feedback onto existing T7 LacI system? logical
% Outputs:
% A - a BioSystems object
if nargin<1
  intmf=false;
end
action_params={'none','down';'up','none'};
deg_params={'linear','normal'};
A=GeneticNetwork(action_params,deg_params);
A.output=sym('x1');
A.output_label={'x1'};
A.int_tol=-11;
A.sys('x2')=A.sys('x2')-str2sym('Vd1/dKd1*x2');
take_limit(A,{'n','dn'},{1,1},true);
A.sys('x1')=subs(A.sys('x1'),sym('V1'),str2sym('V1*(1-u2)'));
A.flux('fH1_2')=subs(A.flux('fH1_2'),sym('x2'),str2sym('Kiaa*(x2+x2g)/(Kaia*u+Kiaa)'));
A.prange('x2g')=[0,1];
A.param('x2g')=1;
A.prange('Kiaa')=[0,10];
A.param('Kiaa')=1;
A.prange('Kaia')=[0,10];
A.param('Kaia')=1;
if intmf
  A.sys('x2')=subs(A.sys('x2'),sym('H2_1'),str2sym('H2_1*H2_3'));
  A.flux('H2_3')=str2sym('b2_3+(1-b2_3)*x3/(x3+Kd2_3)');
  A.xrange('x3')=[0,inf];
  A.x0_map('x3')=1;
  A.sys('x3')=str2sym('V3*(b3_1+(1-b3_1)*x1/(x1+Kd3_1))-Vd3*x3/(dKd3+x3)-Vd1/dKd1*x3');
  A.prange=[A.prange;
    containers.Map2({'b2_3','Kd2_3','V3','b3_1','Kd3_1','Vd3','dKd3'},...
    {[0,0.1],[0,10],[0,10],[0,0.1],[0,10],[0,10],[0,10]})];
  A.param=[A.param;
    containers.Map2({'b2_3','Kd2_3','V3','b3_1','Kd3_1','Vd3','dKd3'},...
    {0.1,10,10,0.1,10,10,10})];
end
end
```
