## Supplemental Data 1 for "Principles underlying implementation of *nearly*-homeostatic biological networks": leakinessred_fmincon_ui_gen.m

```
function [func_obj,func_cons] = leakinessred_fmincon_ui_gen(p2pA,u,i,u_firstislower,...
  x1_set_num,s1_ui_it_x1x2u_num,s1_x1_it_x2u_num,s2_x2_it_x1_num)
% Generates the objective function and constraint function for fmincon
% covers the upper range.
%
% INPUTS
% p2pA - a function that converts [pars_fixed,pars_nopt,pars_opt_ctrlleak,pars_opt_nctrlleak] to parameters
%   for the BioSystem object
% u - a numeric matrix of two perturbations, where each perturbation occupies a row. x1 steady state 
%   is assumed to increase from first to second perturbation. 
% u_firstislower - logical scalar indicating whether the first perturbation is the lower perturbation
% x1_set_num - a function that takes parameters as a column vector. 
% s1_ui_it_x1x2u_num,s1_x1_it_x2u_num,s2_x2_it_x1_num - a function that takes parameters, state variables, 
%   and perturbation as column vectors. 
% 
% OUTPUTS
% func_obj - objective function for fmincon. Returns sum of positive distance from the desired 
%   u range for closed loop system
% func_cons - nonlinear constraint function for fmincon. Lower range permitting near-homeostasis matches 
%   that of u(1,:) for closed loop AND output steady state of open loop system after perturbation is 
%   sufficiently high AND lower steady state range is not too small. 

s1_ui_it_x1u_num=@(p,x,u)s1_ui_it_x1x2u_num( p , [x(1);max(s2_x2_it_x1_num(p,[x(1);nan],nan))] , u );
func_obj=@fmincon_obj;
func_cons=@fmincon_cons;

  function [out] = fmincon_obj(p0_fixed,p0_nopt,p0_opt_ctrlleak,p_opt_nonctrlleak)
    pA=p2pA([p0_fixed;p0_nopt;p0_opt_ctrlleak;p_opt_nonctrlleak]);
    x1l=0.9*x1_set_num(pA);
    x1u=0.9*1.1225*x1_set_num(pA);
    out=max( 0, (-1)^(1-u_firstislower)*(s1_ui_it_x1u_num(pA,[x1l;nan],u(1,:).')-u(1,i)) )...
      + ...
      max( 0, (-1)^(1-u_firstislower)*(u(2,i)-s1_ui_it_x1u_num(pA,[x1u;nan],u(2,:).')) );
  end
  function [c,ceq,x1l,x2l] = fmincon_cons(p0_fixed,p0_nopt,p0_opt_ctrlleak,p_opt_nonctrlleak)
     pA=p2pA([p0_fixed;p0_nopt;p0_opt_ctrlleak;p_opt_nonctrlleak]);
     x1l=0.9*x1_set_num(pA);
     x2l=max(s2_x2_it_x1_num(pA,[x1l;nan],nan));
     x1l_afterpert=s1_x1_it_x2u_num(pA,[nan;x2l],u(2,:).');
     c=[2*x1l-x1l_afterpert,-(x1l-1e-3)];
     % c=[];
     ceq = s1_ui_it_x1u_num(pA,[x1l;nan],u(1,:).') - u(1,i);
  end
end
```
