## Supplementary figures and images for "Principles underlying implementation of *nearly*-homeostatic biological networks"

### demo-reduce_leakiness-optimnonctrlleak_0-change_u1.pdf

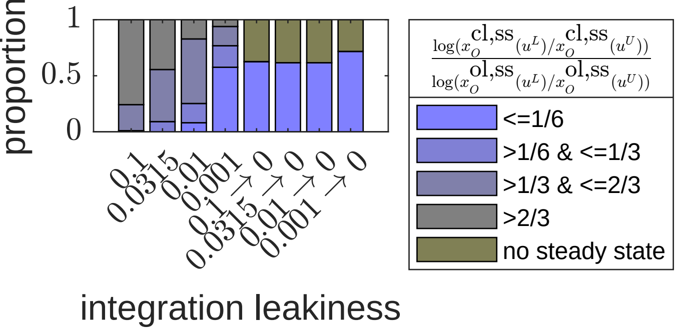

### demo-reduce_leakiness-optimnonctrlleak_0-change_u2.pdf

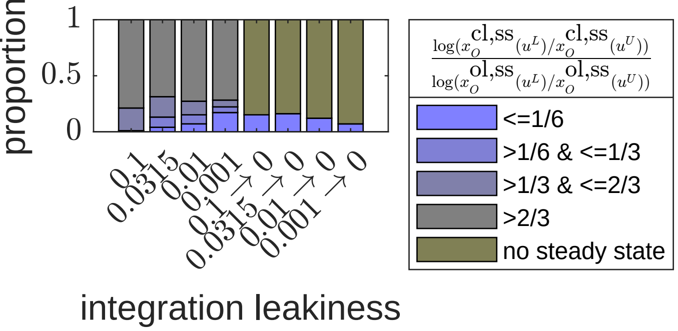

### demo-reduce_leakiness-optimnonctrlleak_0-h_only-change_u1.pdf

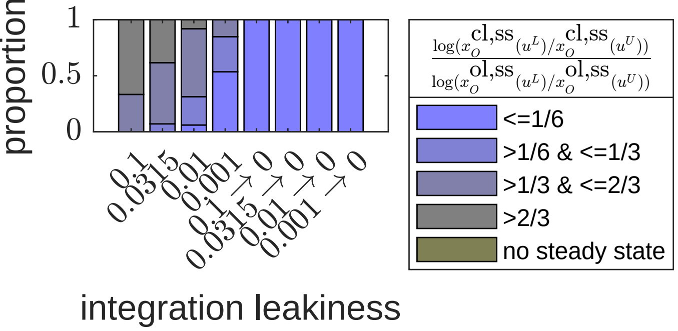

### demo-reduce_leakiness-optimnonctrlleak_0-h_only-change_u2.pdf

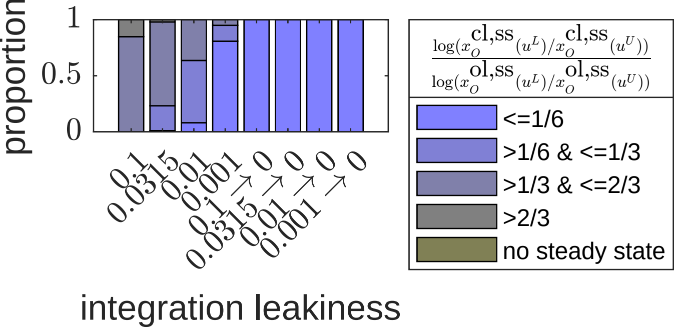

### demo-rl-oo_cl1-u1-ind_leakiness.pdf

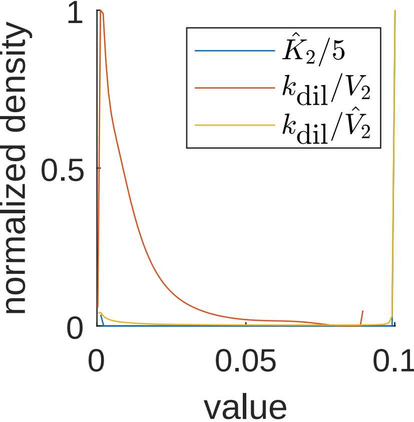

### demo-rl-oo_cl1-u1-optim_include_V2K21b21-ind_leakiness.pdf

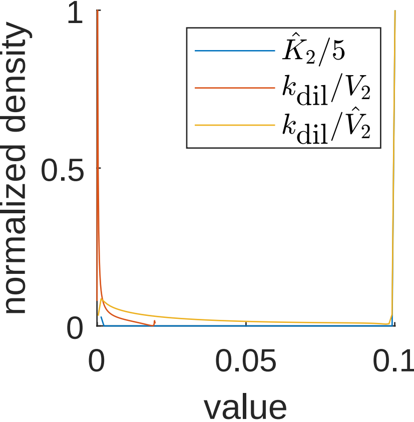

### demo-rl-oo_cl1-u1-optim_include_V2K21b21.pdf

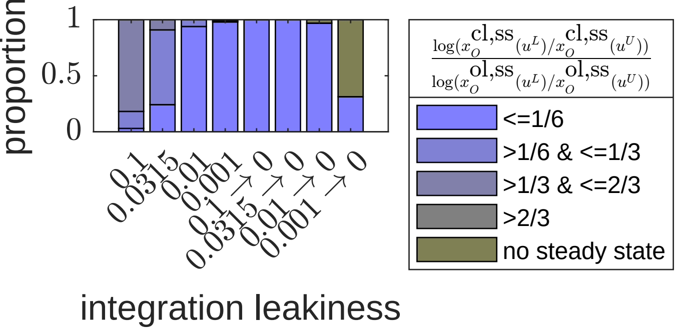

### demo-rl-oo_cl1-u1.pdf

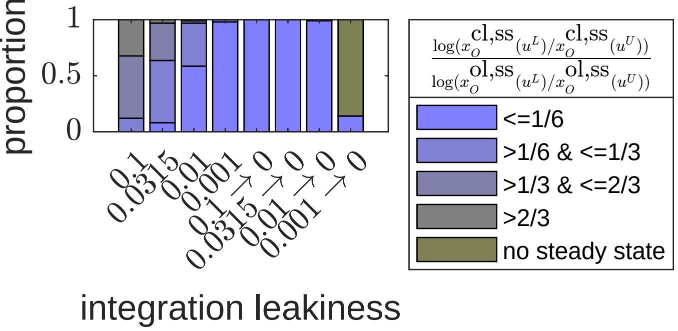

### demo-rl-oo_cl1-u2-ind_leakiness.pdf

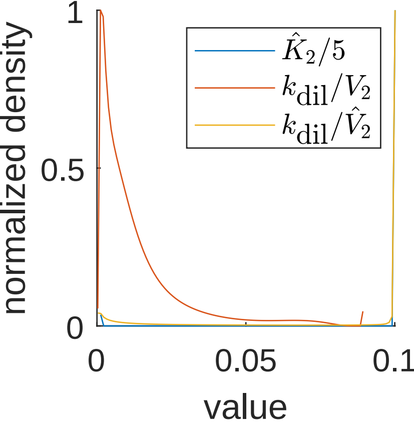

### demo-rl-oo_cl1-u2-optim_include_V2K21b21-ind_leakiness.pdf

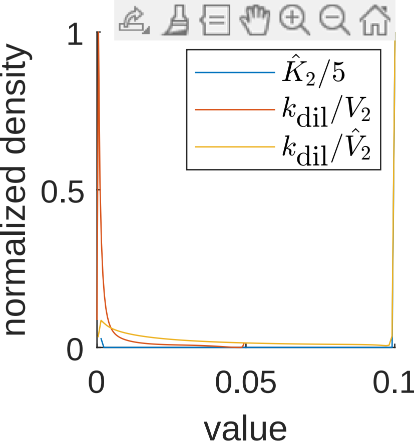

### demo-rl-oo_cl1-u2-optim_include_V2K21b21.pdf

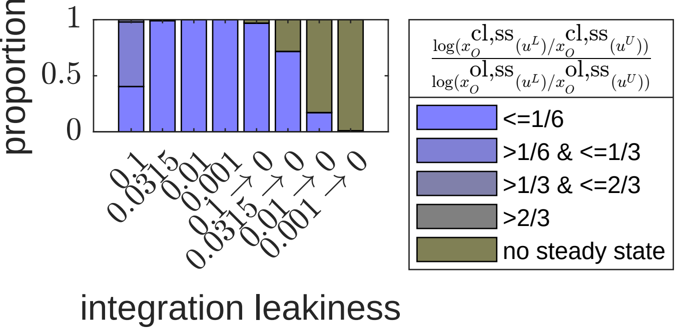

### demo-rl-oo_cl1-u2.pdf

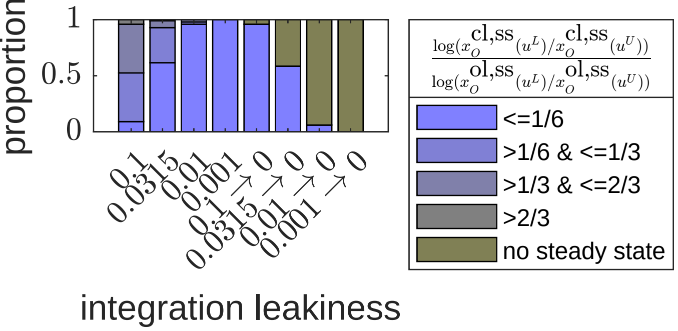

### GeneticNetwork_eq02026545178246969353.png

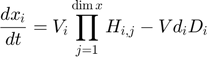

### GeneticNetwork_eq04763600937768006271.png

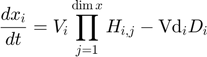

### GeneticNetwork_eq05326875898162696252.png

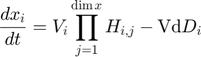

### GeneticNetwork_eq05763904313308679927.png

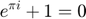

### GeneticNetwork_eq07365652609308031749.png

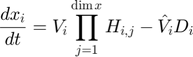

### GeneticNetwork_eq07541279522357761550.png

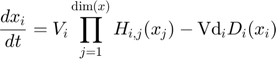

### GeneticNetwork_eq14281132615009919718.png

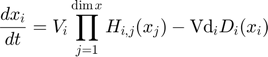

### GeneticNetwork_eq15810091842461394273.png

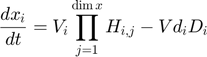

### GeneticNetwork_eq16037459538959007164.png

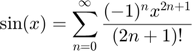

### GeneticNetwork_eq18435133195746625678.png

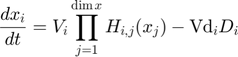
